# Impact Analysis of SARS-CoV2 on Signaling Pathways during COVID19 Pathogenesis using Codon Usage Assisted Host-Viral Protein Interactions

**DOI:** 10.1101/2020.07.29.226217

**Authors:** Jayanta Kumar Das, Subhadip Chakraborty, Swarup Roy

**Affiliations:** Department of Pediatrics, Johns Hopkins University School of Medicine, Maryland, USA; Department of Botany, Nabadwip Vidyasagar College, Nabadwip, India; Network Reconstruction & Analysis (NetRA) Lab, Department of Computer Applications, Sikkim University, Gangtok, India

**Keywords:** Protein Interaction Network, Codon usage bias, Bipartite graph, Cell signaling, Relative Synonymous Codon Usage, Centrality

## Abstract

Understanding the molecular mechanism of COVID19 disease pathogenesis helps in the rapid development of therapeutic targets. Usually, viral protein targets host proteins in an organized fashion. The pathogen may target cell signaling pathways to disrupt the pathway genes’ regular activities, resulting in disease. Understanding the interaction mechanism of viral and host proteins involved in different signaling pathways may help decipher the attacking mechanism on the signal transmission during diseases, followed by discovering appropriate therapeutic solutions.

The expression of any viral gene depends mostly on the host translational machinery. Recent studies report the great significance of codon usage biases in establishing host-viral protein-protein interactions (PPI). Exploiting the codon usage patterns between a pair of co-evolved host and viral proteins may present novel insight into the host-viral protein interactomes during disease pathogenesis. Leveraging the codon usage pattern similarity (and dissimilarity), we propose a computational scheme to recreate the host-viral protein interaction network (HVPPI). We use seventeen (17) essential signaling pathways for our current work and study the possible targeting mechanism of SARS-CoV2 viral proteins on such pathway proteins. We infer both negatively and positively interacting edges in the network. We can find a relationship where one host protein may target by more than one viral protein.

Extensive analysis performed to understand the network topologically and the attacking behavior of the viral proteins. Our study reveals that viral proteins, mostly utilize codons, rare in the targeted host proteins (negatively correlated interaction). Among non-structural proteins, NSP3 and structural protein, Spike (S) protein, are the most influential proteins in interacting multiple host proteins. In ranking the most affected pathways, MAPK pathways observe to be worst affected during the COVID-19 disease. A good number of targeted proteins are highly central in host protein interaction networks. Proteins participating in multiple pathways are also highly connected in their own PPI and mostly targeted by multiple viral proteins.

## 1. Introduction

The entire world is passing through an unprecedented pandemic situation due to a massive outbreak of Severe Acute Respiratory Syndrome Corona Virus 2 (SARS-CoV2) infected viral disease, COVID-19. SARS-CoV2, is a large enveloped coronavirus (family -*Coronaviridae*, subfamily-*Coronavirinae*) with non-segmented, single-stranded, and positive-sense RNA genomes [1], transmits rapidly through human to human contacts. The need for the hour and utmost crucial for the scientific community to understand the disease pathogenesis of SAR-CoV2 in genomics and proteomics level for the rapid development of effective drugs or vaccines to control the COVID-19. Many recent works use HVPPI as an input to elucidate potential drug targets or repurposed drug molecules [2, 3**?**, 4]. Host-pathogen protein interactions provide important insights into the molecular mechanisms of pathogenecity [5].

The host defense mechanism activates signal transduction molecules that initiate signals that activate immune effector mechanisms to protect the host from any pathogenic infections. Studies show that viral immune modulators perturb the human PPI network by targeting signaling pathways [6] to suppress the immunity in mammalian hosts [7]. To understand the molecular mechanism of pathogenicity of SARS-CoV2 during COVID19 disease, investigate the host-viral protein interactions is important. Knowledge gained through the understanding of the viral proteins interact with the host proteins involved in signaling pathways may translate into effective therapies and vaccines. We aim to study the attacking pattern of SARS-CoV2 towards its host proteins involved in signaling pathways. We focus on regulatory signaling pathways, mainly involved in signaling transduction and cellular interactions. Signal transduction focuses on molecular and functional aspects of viral interactions with host cell signaling with relevance for the anti-viral response, the viral life cycle, viral pathogenesis, and cell transformation [8]. Collectively, we consider total seventeen (17) major signaling pathways namely *TGF-beta, Jak-STAT, PI3K-Akt, MAPK, HIF-1, TNF, NF-kappa B, Cytokine-cytokine receptor interaction, Apoptosis, Th17 cell differentiation, Chemokine, Toll-like receptor, RIG-like receptor, IL-17, Insulin Signaling, mTOR*, and *Adipocytokine* signaling pathways [9, 10, 11]. Using a high throughput experimental techniques, like yeast two-hybrid (Y2H) screening and tandem-affinity purification coupled with mass spectrometry revealed miniatures the interactomes of a few model organisms so far. Finding physical interactions is always not feasible due to the expensive experimental setup. Most importantly, it is time-consuming. Experimental methods are prone to incompleteness and noise. Because of the need for rapid development of drugs or vaccines in current pandemic scenarios, the best alternative is to guess the possible physical interactions computationally based on available data sources. Considering different properties of proteins such as sequence homology, gene co-expression, and phylogenetic profiles [12, 13, 14], pairwise similarity is computed between a pair of proteins to predict a possible interaction between them. In addition to non-structural information, structural data about a pair of proteins appears to be more effective in improving prediction accuracy [15, 16, 3]. A number of *in-silico* approaches recently attempted [2, 17] to reconstruct CoV2-Host PPI. Computational methods for predicting interactions among protein molecules, *in-silico* depend largely on the merit of the data in hand and similarity measures [18] used. In reality, predicting deterministically whether two given proteins are physically interacting or not based on the similarity of different structural and non-structural features, is a challenging task due to the above crucial factors.

Few studies report the great significance of codon usage biases [19, 20] in establishing host-viral protein interactions [21]. Viral gene depends largely on the host translational machinery for their expression. Consequently, viral proteins are co-evolved with host proteins and adopt mechanisms to exploit host codon usage biases. Interestingly, highly expressed viral proteins are typically showing similar codon usage biases [22, 23] to target host proteins [24]. On the other hand, to minimize the host immune system responses, certain viral proteins adopt a mechanism to reduce viral protein expression. Few works hypothesized [25, 26] that viral proteins enrich in codons that are rare in their host genomes, and this rare codon usage helps in controlling the expression level.

In this work, we try to infer a host-viral protein interaction network leveraging the inherent correlation between codon usage biases between viral and host proteins. To the best of our knowledge, no prior work exploiting the codon usage pattern to infer host-viral PPI. We try to capture both positive and negative interactions in the host-viral PPI. We use host proteins involved in different human cellular signaling pathways by looking into their significance in disease pathogenesis. Topologically and biologically, we try to establish the relevance of the host proteins and highlight a few essential proteins in the network, which may be useful drug targets for certain reported drugs.

## 2. Methods and Materials

In this section, we discuss our proposed scheme for constructing a hostviral PPI network using codon usage patterns of host and viral proteins. To analyze the interaction mechanism of SARS-CoV2 viral proteins in host signaling pathways, we select a set of all the genes involved in few candidate signaling pathways.

### 2.1. Data acquisition and processing

Structurally, SARS-CoV2 consists of three categories of proteins, Structural, Nonstructural, and Accessory proteins. We select four (04) structural proteins, sixteen (16) non-structural proteins, and six (06) as reported in Wuhan, China [27]. The details of the viral proteins are listed in Table 1 (NCBI accession numbers for SERS-CoV2 proteins: MN908947.3, NC - 045512).

**Table 1:** SARS-CoV2 Proteins considered for host-viral PPI

| Protein category | Count | Protein Name |
| --- | --- | --- |
| Structural | 4 | Spike (S), Envelope (E), Membrane (M), Nucleocapsid (N) |
| Non Structural | 16 | Nsp1-Nsp16 |
| Accessory | 6 | Orf3a, Orf6, Orf7a, Orf7b, Orf8, Orf10 |

To collect significant host genes involved in our candidate 17 signaling pathways, we search Kyoto Encyclopedia of Genes and Genomes (KEGG) database ^1^ for the genes involved in those pathways [28]. We observe a total of 2600 genes involved in the above pathways (Supplementary-A). Out of which, we select 1313 unique genes fully engaged in those signaling pathways as a good number of genes (1274) involve in more than one pathway. We summarize our target pathways and the number of genes involved, in the Table 2.

**Table 2:** Host proteins collected from different signaling pathways

| Pathways | #Genes involved | Pathways | #Genes involved |
| --- | --- | --- | --- |
| NF-kappa B signaling pathway | 105 | Th17 cell differentiationl | 108 |
| Cytokine-cytokine receptor interaction | 295 | TGF-beta signaling pathway | 95 |
| TNF signaling pathway | 113 | Toll-like receptor signaling pathway | 105 |
| IL-17 signaling pathway | 95 | HIF-1 signaling pathway | 110 |
| RIG-I-like receptor signaling pathway | 70 | Apoptosis | 137 |
| MAPK signaling pathway | 295 | Insulin signaling pathway | 138 |
| Chemokine signaling pathway | 190 | mTOR signaling pathway | 156 |
| PI3K-Akt signaling pathway | 355 | Adipocytokine signaling pathway | 70 |
| Jak-STAT signaling pathway | 163 |  |  |

### 2.2. Computing Relative Synonymous Codon Usage (RSCU)

The genetic code describes how the 64-nucleotide triplets specify only twenty (20) different translated amino acids. These alternative codons for the same amino acids are termed as *synonymous codons*. However, most of the amino acids have at least two synonymous codons that are not used at the same frequencies in different genomes. According to the genome hypothesis proposed by Grantham et al. [29], the pattern of codon usage is speciesspecific and some way unique. Interestingly, even in the same genome, the codon usage varies significantly among genes with different expression levels [23], functions [30], and tissue-specific patterns [31]. Codon usage variation occurs may be due to natural selection and/or mutation pressure for accurate and efficient translation in various organisms [19, 20]. Differences in the frequency of occurrence on synonymous codons in coding DNA is termed as synonymous codon usage bias [32].

RSCU is one the indices for measuring codon bias independent of the amino acid composition and widely used to estimate the codon usage bias [33, 34, 35]. It may use to quantify the similarity between any two gene sequences by applying any classical proximity measure between a pair of RSCU vectors. The similarity between RSCU vectors may reflect the possible interactions between a couple of proteins in the PPI [35, 36, 34].

RSCU is the ratio between the observed number of occurrences of codons and expected during uniform usage of synonymous codons and can be calculated as follows.

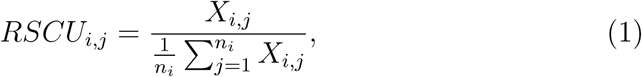

where, *X*_*i,j*_ is the number of occurrences of the *j*^*th*^ codon for the *i*^*th*^ amino acid, which is encoded by *n*_*i*_ synonymous codons. The RSCU score of a codon more than 1.0 indicates excess usage (biased) of the codon, and less than 1.0 marks poor usage of that particular codon.

We report codon usage distribution of 59 codons across 26 SARS-CoV2 proteins in Figure 1 (a) and 1313 host proteins (Figure 1 (b)) involved in our candidate signaling pathways. We observe that GGT, AGA, GCT, CCT, GTT, TCT, ACA, CTT, TTA, ACT are the highly used (median RSCU score ≥1.5 for each codon) codons in SAR-CoV2 proteins. On the other hand, CGA, AGC, ACC, CGG, CTG, CCG, ACG, GCG, TCG, GGG rarely used codons. In the case of host proteins (from 17 signaling pathways), codons such as CTG, GTG, ATC, GCC, CAG, ACC, AGC, GGC, CCC are highly used (median RSCU score ≥1.5 for each codon). The distribution margins of RSCU values of those codons are relatively wider (Figure 1 (b)). However, CCG, GTT, CGT, GCG, TCG, CAA, CTA, ATA, GTA, TTA rarely used codons in host proteins. It is worth mentioning that for SARS-CoV2 proteins, highly used codons are ending with T or A and host proteins, ending with G or C at the third position of the codons.

**Figure 1:**
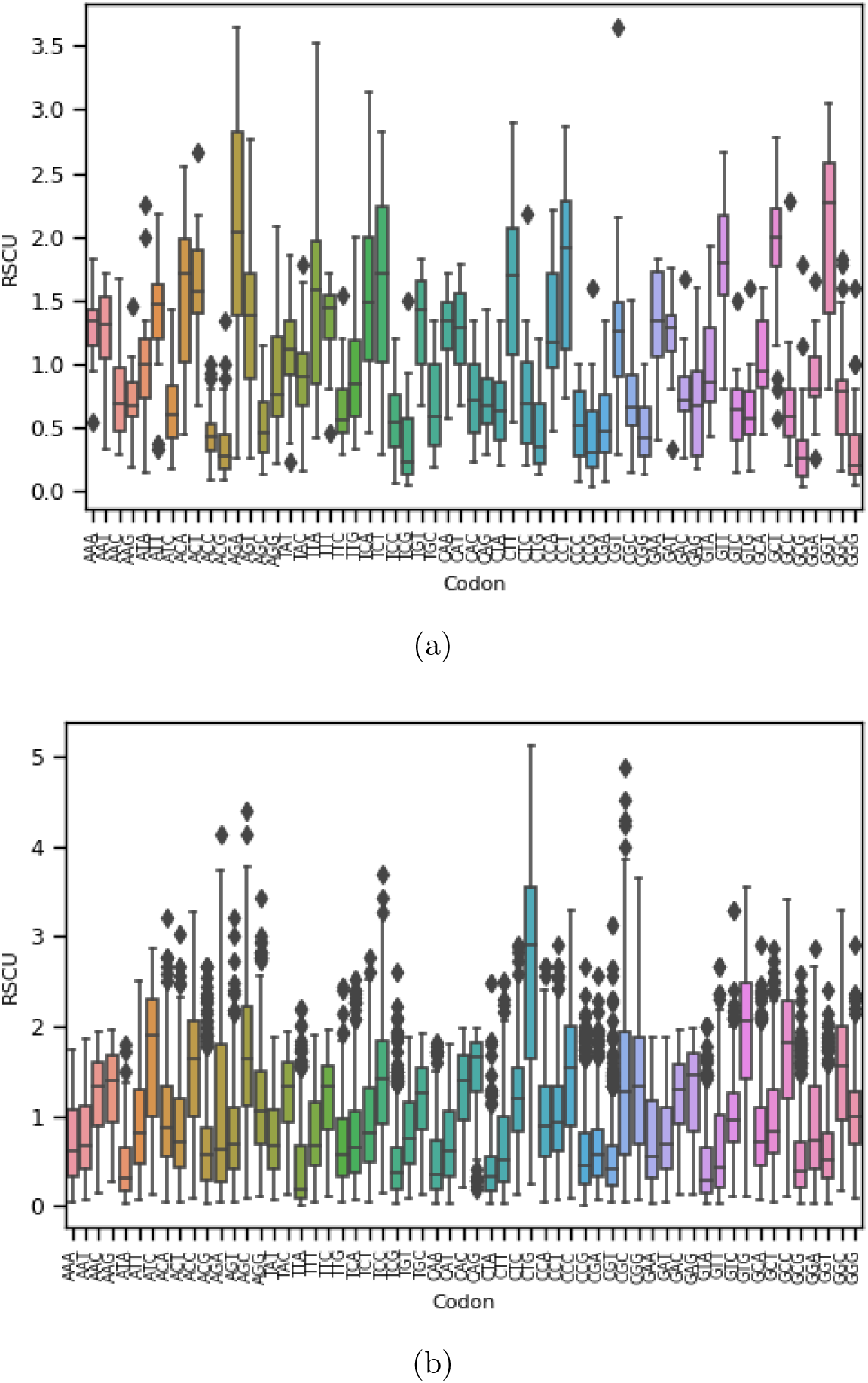
Distribution of RSCU scores for 59 codons for different (a) SARS-CoV2 proteins (b) Host proteins.

We generate a 59-dimensional RSCU feature vector for each coding protein. We consider the usage pattern of only 59 codons (out of 64 available codons). We ignore 03 stop codons and uniquely coded codons ATG and TGG coded for Met and Trp amino acids, respectively [37]. For RSCU calculation, we use *CAI* package [38] available free at ^2^. Using the feature vectors, we try to draw the similarity between host and viral proteins to form a network, as discussed next.

### 2.3. Inferring Host-Viral Protein Interaction Network

Protein-Protein Interactions (PPI) are usually studied computationally from a graph-theoretic perspective [18]. Interactions among different organisms, such as a host and its pathogen, are primarily driven by interactions among the host proteins and pathogen proteins. These interactions can also be represented as host-pathogen PPI. Host-pathogen PPI usually represented as a *bipartite graph* where any given interacting pair of nodes (proteins) does not belong to the same organism. This network essentially provides the known interactions of host proteins with pathogen proteins. On the other hand, the interactions may be guessed (or inferred) computationally based on different attributes of the target protein pairs when physical interactions are missing. The host-pathogen interaction network essentially represents a snapshot of the infection mechanism in a host cell infected by pathogens. The virus-host interactome is essential for understanding virulence factors influencing SARS-CoV2 pathogenesis [39]. Recent studies reported the use of SARS-CoV2 and host PPI networks to study the pathogenesis of SARS-CoV2 and identifying repurposed drugs [17**?**, 3, 40].

We infer interaction between two proteins using codon usage similarity. Two proteins are considered to be strongly coupled if there RSCU similarity bearing particular statistical significance.

Given *R*_*v*_ = {*x*_1_, *x*_2_, · · ·, *x*_*m*=59_} and *R*_*h*_ = {*y*_1_, *x*_2_, · · ·, *y*_*m*=59_ }, RSCU vectors of a pair of host and viral proteins respectively, proteins are strongly connected if *p* score is less than certain threshold *τ* i.e. *p*(*R*_*v*_, *R*_*h*_) *< τ*.

We use SciPy version 1.5.0 (*sipy.stats*) ^3^ for calculating Pearson correlation coefficient, which uses 2-tailed *p*-value for measuring significant relationships [41].

The presence of interaction between a viral protein *V p*_*i*_ and a host protein *Hp*_*i*_ is dependent on a significant threshold () obtained from the given RSCU-feature vectors of the host protein and the viral protein. For strong relationship we set *p <* 0.001 and *τ <* p).

Inferring the network, we consider two (02) kinds of interaction, positive and negative, between a host and viral protein. Positive interaction indicates possible similar codon usage patterns and negative interactions signifying possible rare codon usage by SARS-CoV2 proteins compared to its interacting host proteins.

Given *R*_*v*_ = {*x*_1_, *x*_2_, …, *x*_*m*=59_} and *R*_*h*_ = {*y*_1_, *x*_2_, …, *y*_*m*=59_ }, RSCU vectors of a pair of host and viral proteins, respectively, we use Pearson Correlation (*ρ*) [42] to calculate signed edge as follows.

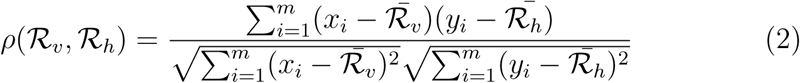

where, *x*_*i*_ *∈ R*_*v*_ and *y*_*i*_ *∈ R*_*h*_, 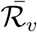 and 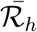 are the mean of the vectors *R*_*v*_ and *R*_*h*_ respectively. We use SciPy version 1.5.0 ^4^ for computing *ρ*.

Given a set of viral proteins, *V* = *{v*_1_, *v*_2_, *· · · v*_*n*_*}* and host proteins *H* = {*h*_1_, *h*_2_, *h*_*n*_} we can create a bipartite graph in the form of adjacency matrix using above *ρ* and *p* values as follows.

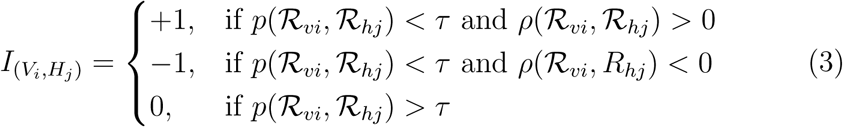

Next, we investigate the interaction mechanism of SARS-CoV2 on human signaling pathways during COVID19 disease pathogenesis.

## 3. Results and Discussion

We predict the host-viral interaction graph based on the Equation 3 involving 26 viral proteins with 1326 host proteins participating in 17 different signaling pathways. Out of 34138 (26 ×1313) maximum possible interactions, our method infers 9412 (≈36%) strong interactions where 859 distinct host proteins (≈66%) are connected to at least one viral protein. We fix *τ* = 0.001 as significant *p*-value for inferring strong edge between two proteins. Interestingly, our inferred network reveals that out of 859 host proteins, a total of 779 proteins is targeted by more than one viral protein. A snapshot of isolated networks with one (viral) to many (host) interactions is shown in Figure 3 between viral and host proteins.

**Figure 2:**
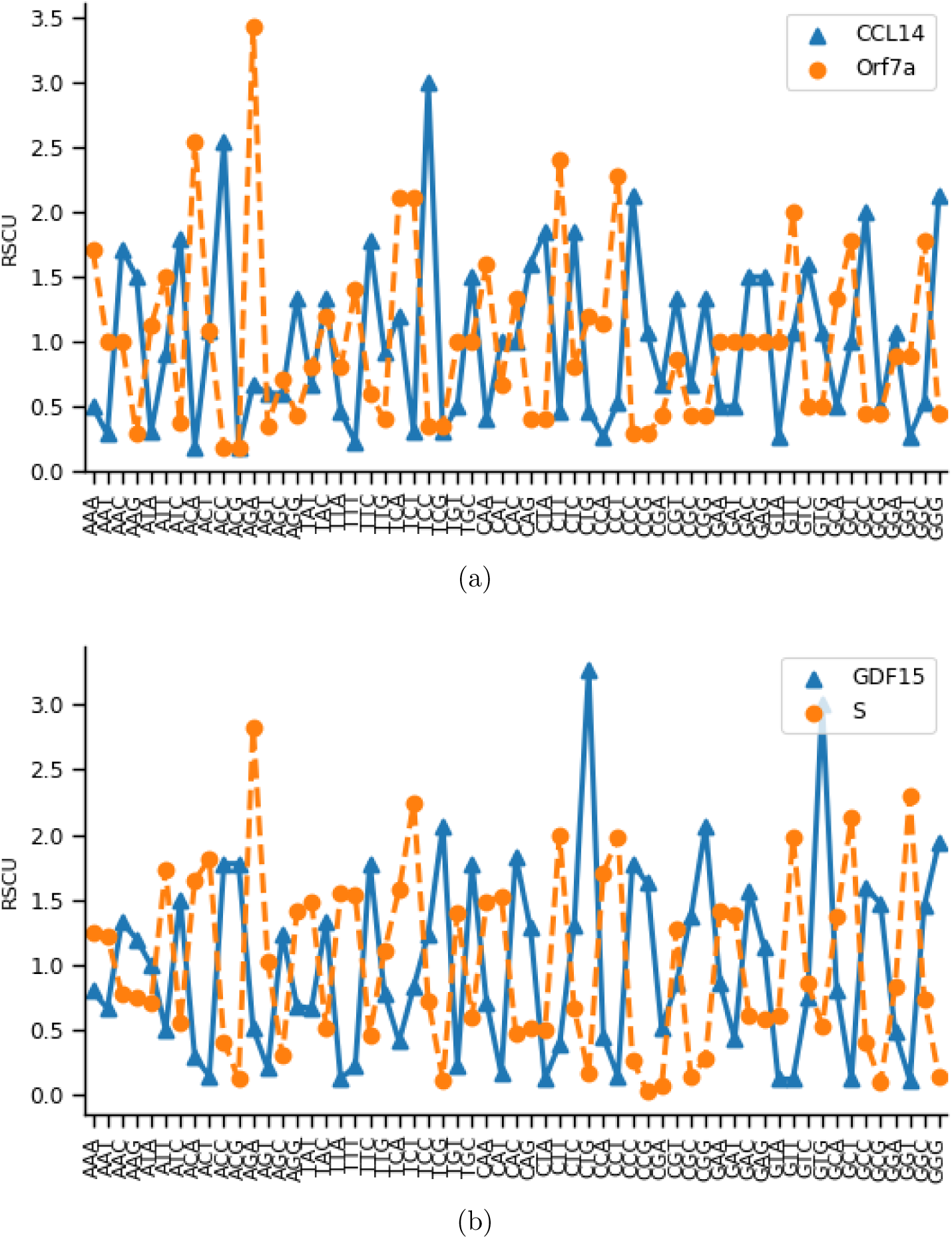
An example is showing for the viral protein and host protein codon usage RSCU pattern. x-axis is showing 59 codons and y-axis is for respective RSCU value of each codon. (a) Viral protein Orf7a that showed positive correlation (*r* = 0.58) with host protein TANK. (b) Viral protein Spike (S) that showed negative correlation (*r* = −0.73) with host protein GDF15.

**Figure 3:**
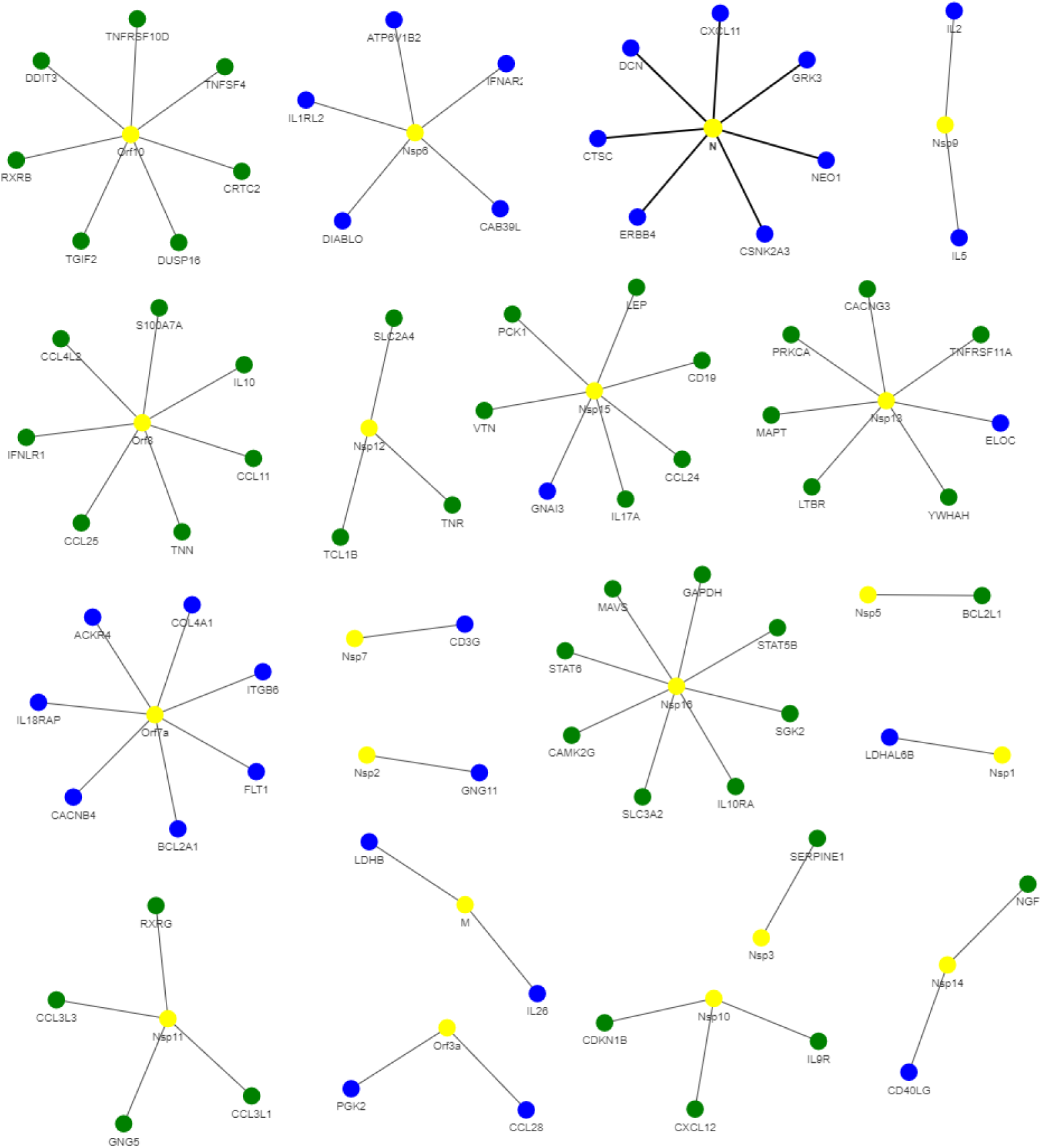
Disconnected networks showing one viral node (yellow node) to many host nodes interacting positively (blue node) and negatively (green node). Among 20 viral proteins, 09 viral proteins (Nsp1, Nsp2, Nsp6, Nsp7, Nsp9, M, N, Orf3a and Orf7a) are interacting positively by unique (one-to-one) host protein and the 08 viral proteins (Nsp3, Nsp5, Nsp10, Nsp11, Nsp12, Nsp16, Orf8 and Orf10) are interacting negatively with unique host protein. Nsp13, Nsp14 and Nsp15 involve both types of interactions.

Recent similar researches on SARS-CoV2 host proteins interaction [2] produce only viral protein oriented star-like topology and unable to report any host protein oriented multiple interactions. We report a list of such highly connected host proteins with the viral proteins (at least 15) in Table 3. Many (viral) to one (host) interactions are also reported as Supplementary material (Supplementary-B).

**Table 3:**
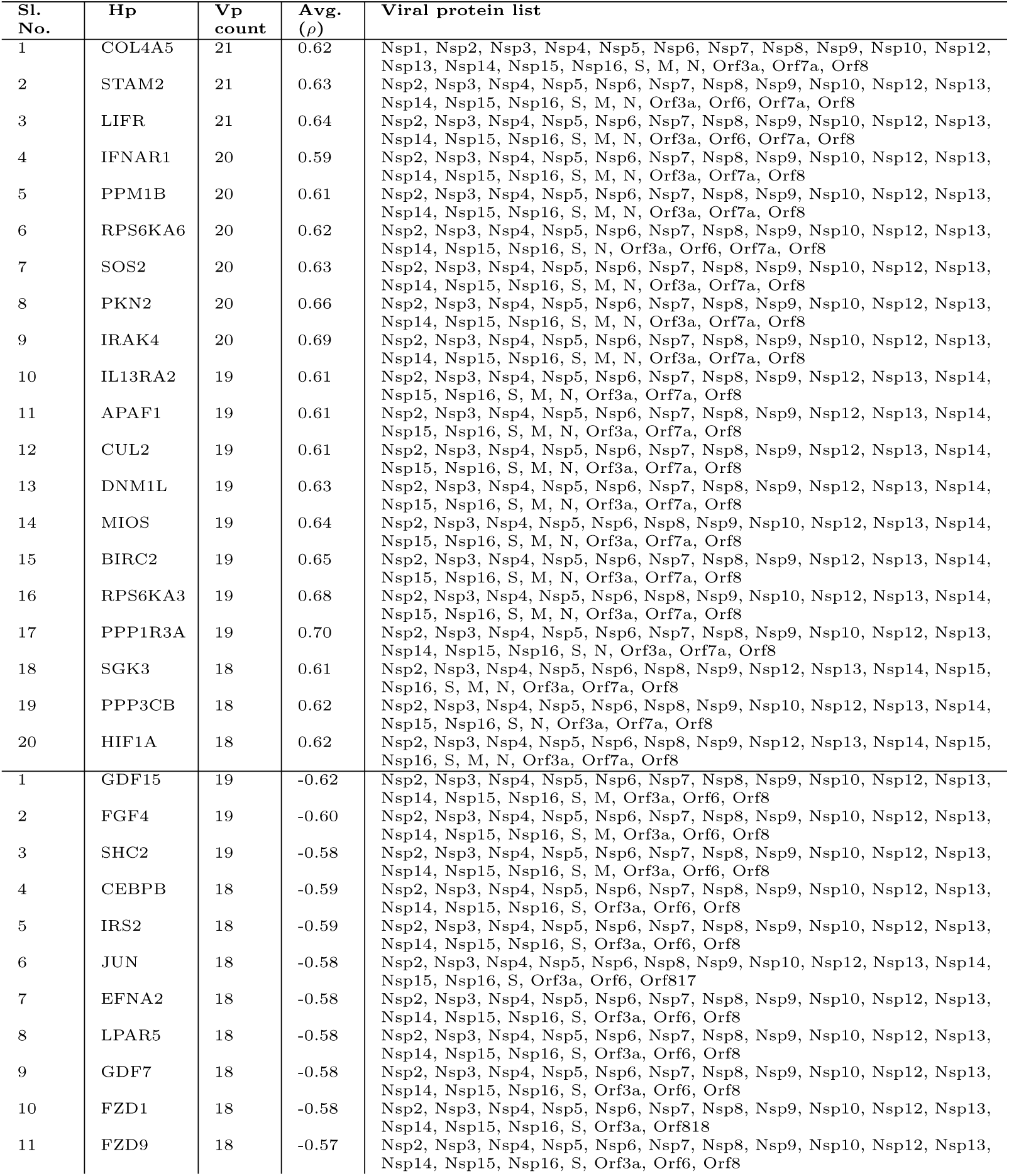
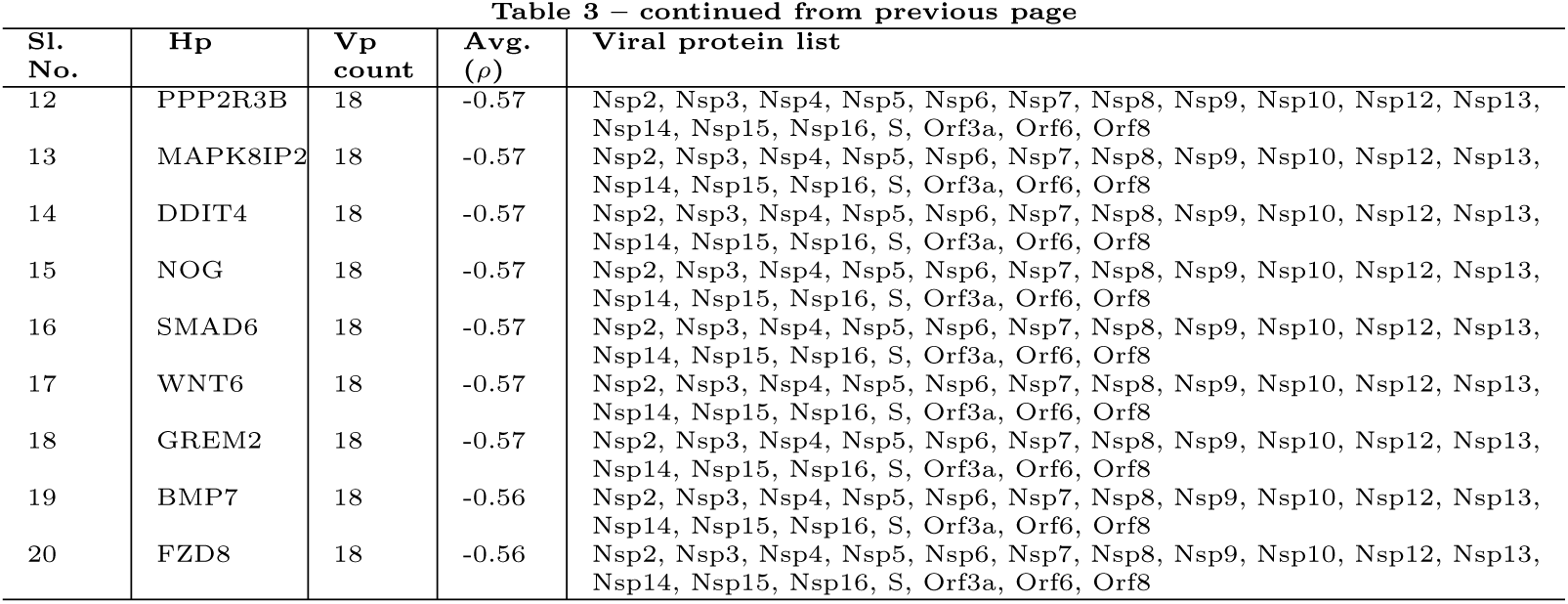
List of highly interacting host proteins (Hp) targeted by least 15 viral proteins (Vp). The host proteins are ranked by interacting viral protein count and shown along with an average correlation value. The list of the first 20 host proteins positively interact and the next 20 host proteins negatively interact with viral proteins.

### 3.1. Distribution of correlation scores

While inferring an edge we consider only the significant *p* score (*p* = .001) and use the sign derived from correlation (*ρ*) as a type of the interaction. Statistically, it is also important to study the distribution of correlation values (both positive and negative) between pairs of proteins in terms of codon usage patterns. From the distribution plot given in Figure 4 reveals that codon usage pattern between a pair of host and viral proteins (edge correlation) are non randomly associated and showing a Gaussian [43, 44] like distributions (with *p* = 1.13*e* –42 for positive and *p* = 7.819*e* –94 for negative correlation distributions based on normality test performed using SciPy.stats.normaltest ^5^). The negative correlation is varied in the range [-.73, -4.18] which covered 6325 (67%) interactions, and a positive correlation is varied within [4.18, 8.44] including 3087 (33%) interactions. Positive correlation exhibits a more wider range of values than the negative range.

**Figure 4:**
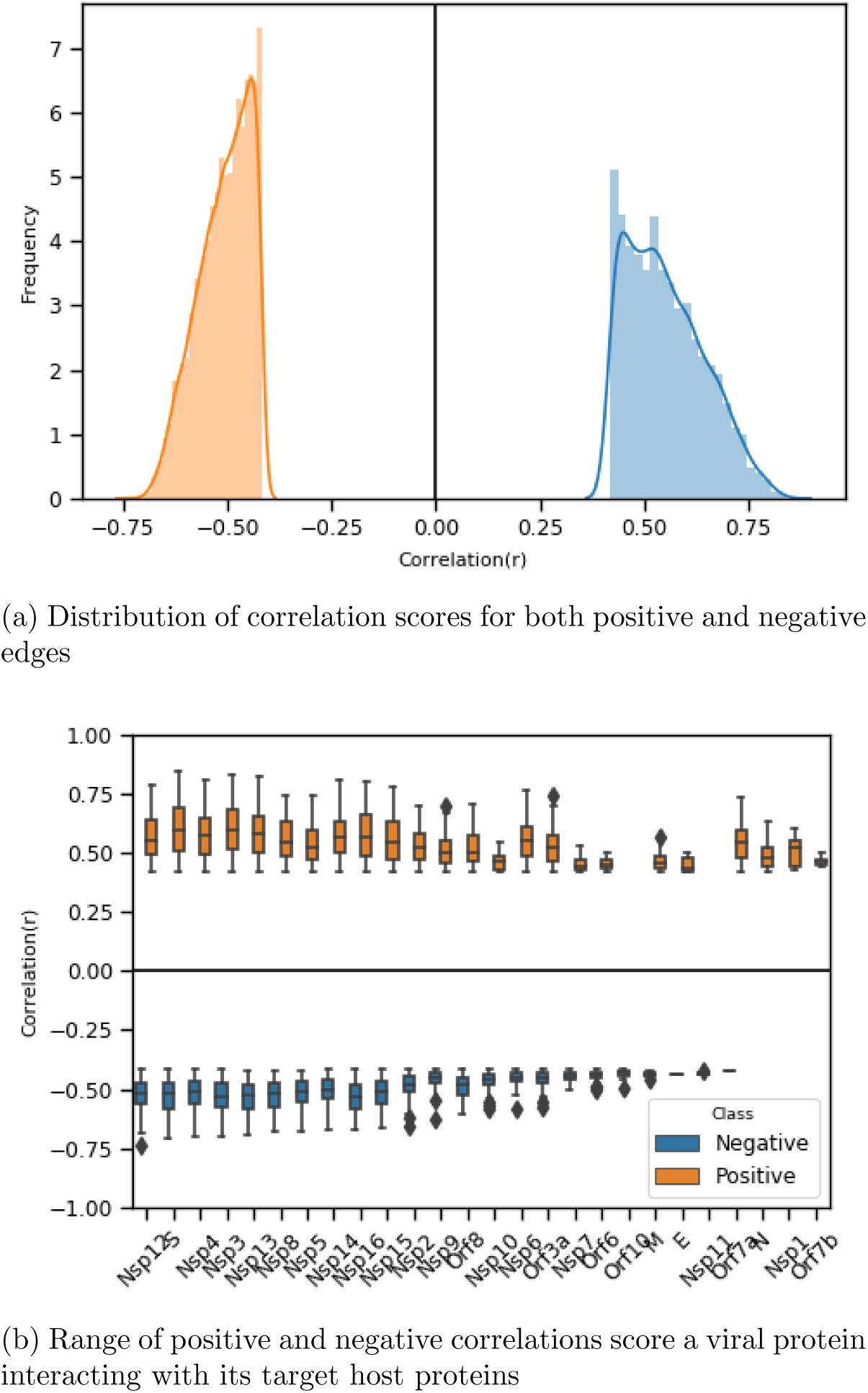
(a) Frequency distribution of positive (right) and negative (left) correlation scores for interacting proteins in terms of RSCU based codon similarity showing a normal distribution. (b) Box plot showing the range of correlation values for each viral protein while associated with its target proteins.

We further look into the correlation value distribution of a viral protein interacting with its target proteins. We report the correlation value range (both positive and negative) for 26 viral proteins in Figure 4. We observe that while fixing the *p* score at a high significance level, correlation values also appear to be significant, and it is ranging from ±0.05 and above. The majority of the viral proteins are correlated both positively and negatively with its targets except for a few in the inferred network. Orf10 and Nsp10 are viral proteins that are interacting with their target negatively. Similarly, viral proteins like N, Nsp1 and Orf7b are interacting positively only.

Based on our correlation analysis, we may confirm that while a viral protein targets its a host, it mimics similar codon usage as its target to uphold the expression of target host proteins. Similarly, viral proteins use a set of codons that are rarely used in their targets to down-regulate the expression of its target. We observe that in the case of host proteins involved in signaling pathways, the majority of SARS-CoV2 proteins aimed to break down the normal pathways by downregulating the key proteins involved in such pathways.

### 3.2. Degree distribution of host and viral proteins

In any interacting network, the node’s degree conveys essential information about the influence of the node within the network. In the case of hostviral PPI, a high degree of viral protein may be a key protein that interacts with a high number of host proteins. Pharmacologically, the identification of such proteins may help design a small molecule that may bind with it to inhibit its influence during disease pathogenesis. The same may be applicable to host proteins. If host proteins have a high degree, it indicates that the host proteins are targeted by more number of viral proteins. However, it may require further investigation about its importance in its own network i.e., host-host protein networks. If a host protein found to be significant concerning its degree, suitable repurposed drug molecules may be identified for the same.

While focusing on highly interacting viral proteins, interestingly, we observe that the maximum number of highly interacting proteins belongs to the non-structural family. In the case of structural proteins, S is highly interacting (more than 600) protein. Out of accessory proteins, Orf8 shows maximum interaction count close to protein S.

We report the degree distribution for each of the viral proteins from our network in Figure 5 (a). From the figure, we may observe that majority of the viral proteins carrying a high node degree. Out of all the SARS-CoV2 proteins, Nsp3 shows the maximum degree (≈700), which interacts with more than 80% of the candidate host proteins involved in 17 different signaling pathways. Concerning negative edges, i.e., connected, negatively, Nsp3 is still on top, followed by Nsp16, Nsp13, and so on. While considering positive edges S, Nsp6 and Orf7a are found to be highly interactive. Few viral proteins like Nsp11, Orf7b, E, Nsp1, Orf10, and Nsp11 found to be less interactive, comparatively.

**Figure 5:**
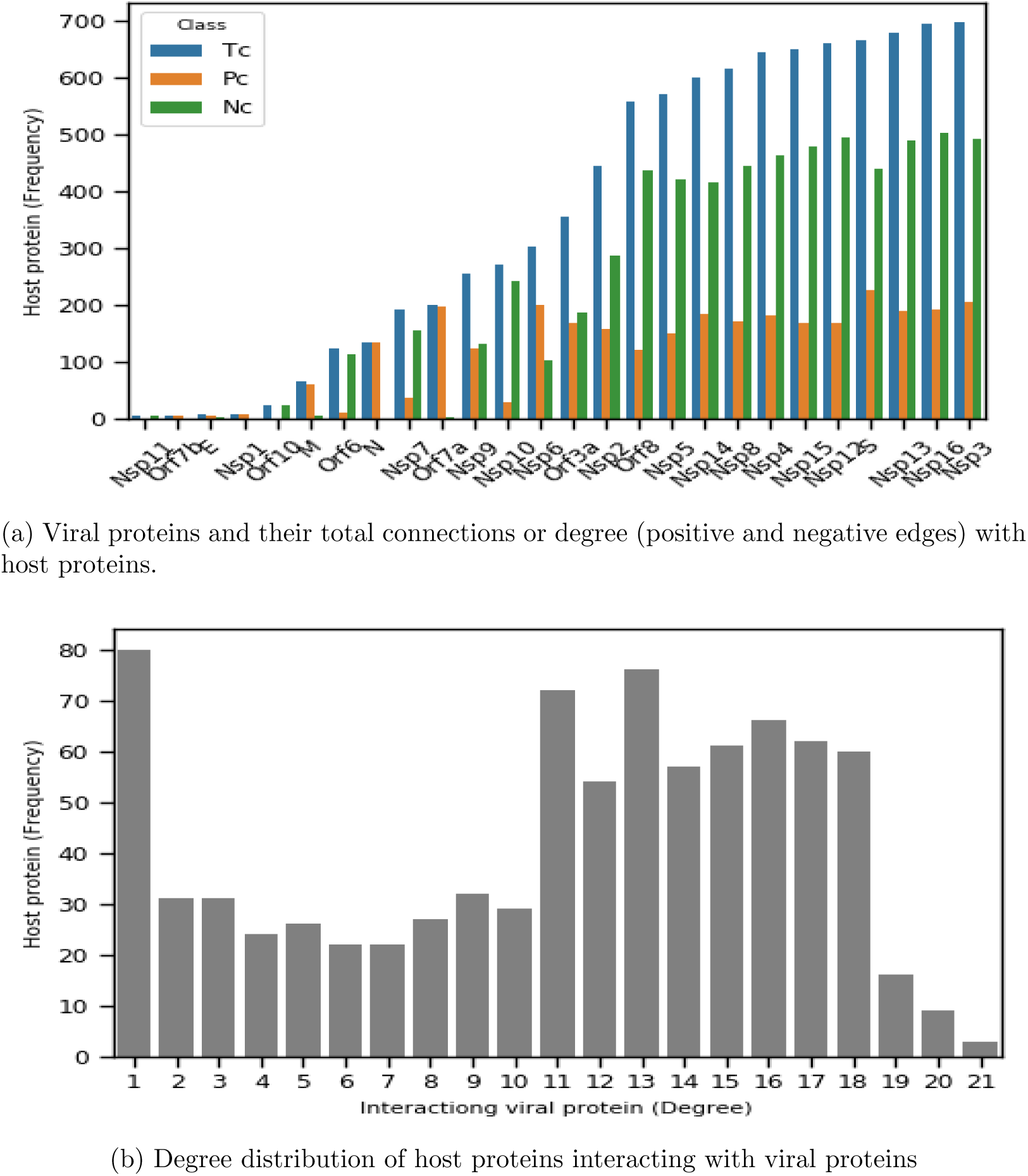
(a) The bar chart showing for number of host protein count for each viral protein based on correlation analysis (p-value*<* 0.001). Pc-positive count, Nc-negative count; positive and negative count are based on positive and negative correlation respectively. (b) Degree distribution of 859 host proteins in terms of number of associated viral proteins (degree) count (x-axis) with host protein frequency (y-axis).

We show the distribution of the degree of 859 host proteins in Figure 5 (b), interacting with 26 viral proteins. From the distribution plot, we can observe that majority (82) of the host proteins are connected with only one viral node. While considering highly targeted proteins by multiple viral proteins, we see fewer than 10 proteins with degree 21 (maximum degree), which is lowest within the distribution. Even though our network is a bipartite graph, we observe that the number of low-degree nodes is high and high degree nodes are lowest in the graph. It further indicates that hub or central nodes are relatively less, which is somehow following the scale-free properties [45] of a complex network. However, with an exception to the power-law distribution curve, we observe relatively good host nodes possessing a degree within the range 11 to 18.

### 3.3. Ranking highly targeted signaling pathways and its impact on COVID19

To study the most effected pathways in our 17 candidate set of pathways, we rank them based on the percentage of host protein targeted by any viral proteins out of total proteins involved in those pathways and report in Figure 6.

**Figure 6:**
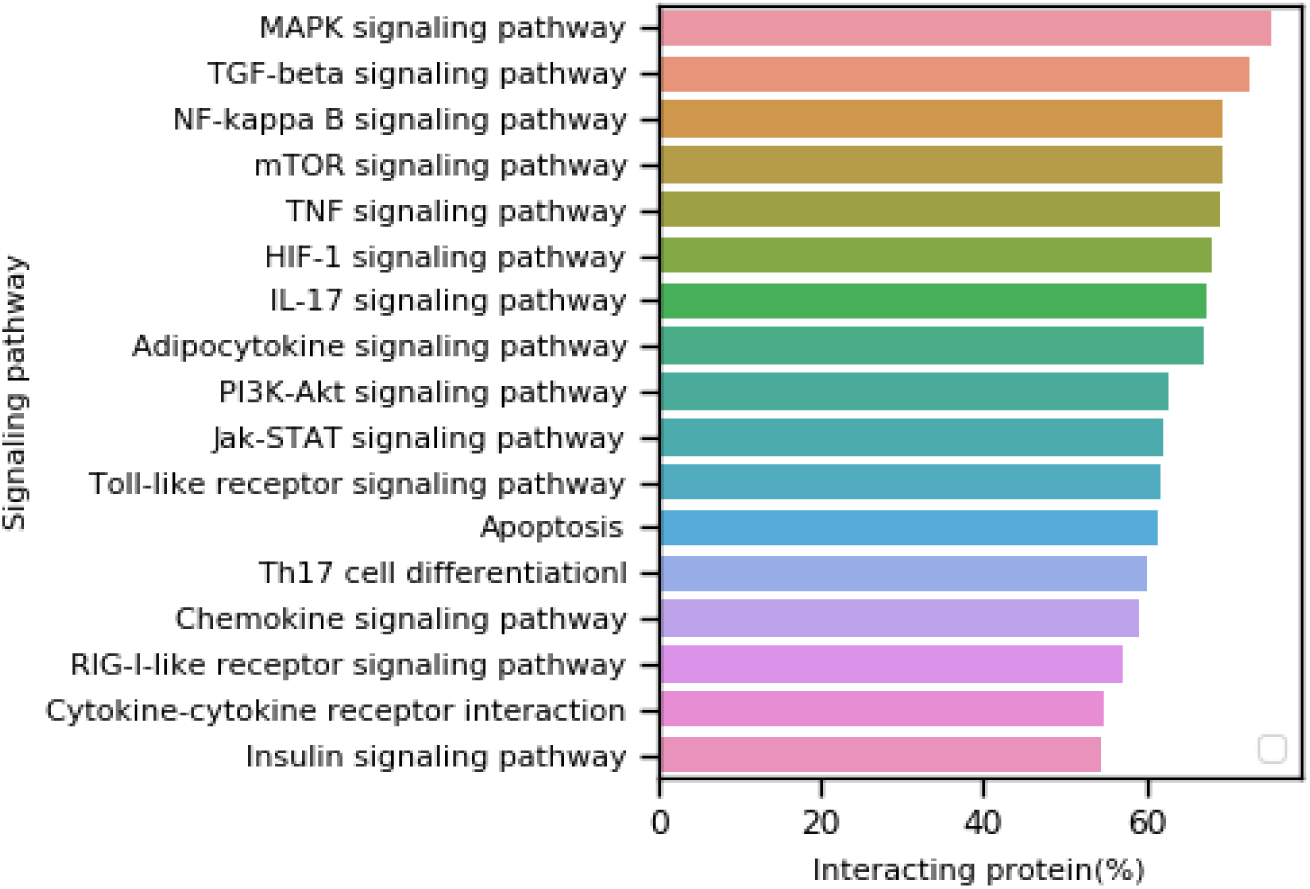
Ranking of 17 candidate signaling pathways. The pathway ranking is done by observing the host protein percentage from pathways that interact with any of the SARS-CoV2 (26) proteins.

From the figure, it is observable that Mitogen-activated protein kinase (MAPK) signaling pathways is viral proteins target profoundly affected pathways with more than 50% proteins from that pathway. MAPK proteins communicate signals from a receptor on the cell’s surface to the DNA in the nucleus of the cell, which is essential in a viral infection point of view. Besides, MAPK proteins are involved in a series of vital signal transduction pathways that regulate processes such as cell proliferation, cell differentiation, and cell death in humans.

Beside MAPK, other ranked signaling pathways are significantly affected during COVID19 infection. Under physiological conditions, adipokines act mainly in adipose tissue (paracrine or autocrine) or circulate through the blood circulation to distant target organs, regulating their growth and development, metabolism and tissue remodeling. However, under pathological conditions, the synthesis and secretion of adipokines are disordered, leading to obesity, diabetes, heart disease, and other metabolic disorders. Our results show that the adipocytokine pathway is affected during COVID19. It implicates the patients with comorbid conditions like diabetes and heart disease show worst disease aggression, which already observed in various reportings. The mTOR pathway is a central regulator of mammalian metabolism and physiology, with essential roles in the function of tissues, including liver, muscle, white and brown adipose tissue, and the brain. It dysregulated in human diseases, such as diabetes, obesity, depression, aging-related problems, and certain cancers. Our result corroborates with the same, and it’s reported that aged patients are more prone to the infection due to the dysregulation of m-TOR pathway or some other unknown reasons.

It also comes to notice that some COVID19 affected deaths are due to multiple organ failure. HIF1 and RIG1 like receptor pathways, are involved in normal immunoregulation and various organ functioning. Dysregulation may cause immune compromisation and multiple organ failure through ischaemic heart disease, acute lung injury, pulmonary hypertension, pulmonary fibrosis, chronic obstructive pulmonary disease (COPD), acute liver failure, liver fibrosis and acute kidney injury etc. Our result also supports these findings. In our ranking, the fourth most affected pathway is the TGF-*β* (Transforming growth factor-beta), which is a multi-functional cytokine belonging to the transforming growth factor superfamily that includes three different mammalian isoforms (TGF-*β* 1 to 3, HGNC symbols TGFB1, TGFB2, TGFB3) and many other signaling proteins. TGFB proteins are produced by all-white blood cell lineages. This pathway activates different downstream substrates and regulatory proteins, inducing transcription of various target genes that function in differentiation, chemotaxis, proliferation, and activation of many immune cells.

### 3.4. Centrality analysis of host proteins

Studies on human host-viral protein interactions reveal that virus tending to targeted attacks towards host proteins [46, 47, 48] by interacting with important (central) host proteins. We consider a host protein important if it interacts with many other host proteins in host-host protein networks. We use BioGRID [49] to calculate the centrality score of our candidate host proteins ^6^. We report top 100 central proteins in the Supplementary C. We observe that a good number of interacting host proteins in our network are highly central in the host PPI. We observe that a common set of viral proteins targets central genes, and such proteins are involved in multiple pathways. For instance, if we consider few top central proteins, MYC (2843), TRIM25 (2656), EGFR (2452), BRCA1 (2236), MDM2 (2219), NTRK1 (2030), KRAS (1944), ELAVL1 (1914) and HSP90AA1 (1734), they are found to be targeted commonly by the viral proteins such Nsp2, Nsp3, Nsp4, Nsp5, Nsp8, Nsp10, Nsp12, Nsp13.

If we consider the participation of most central proteins in our candidate pathways, we observe that PI3K-Akt signaling pathway (36) followed by the MAPK signaling pathway (35) contains most of the central proteins. From the disease pathogenesis perspective, a pathway may be more crucial if it contains highly central proteins targeted by viral proteins. Moving one step ahead, we may rank our 17 pathways based on the number of participating central proteins (top 100) in the above pathways and report it in the Table 4. More details about the top 100 central host proteins are listed in Supplementary C.

**Table 4:**
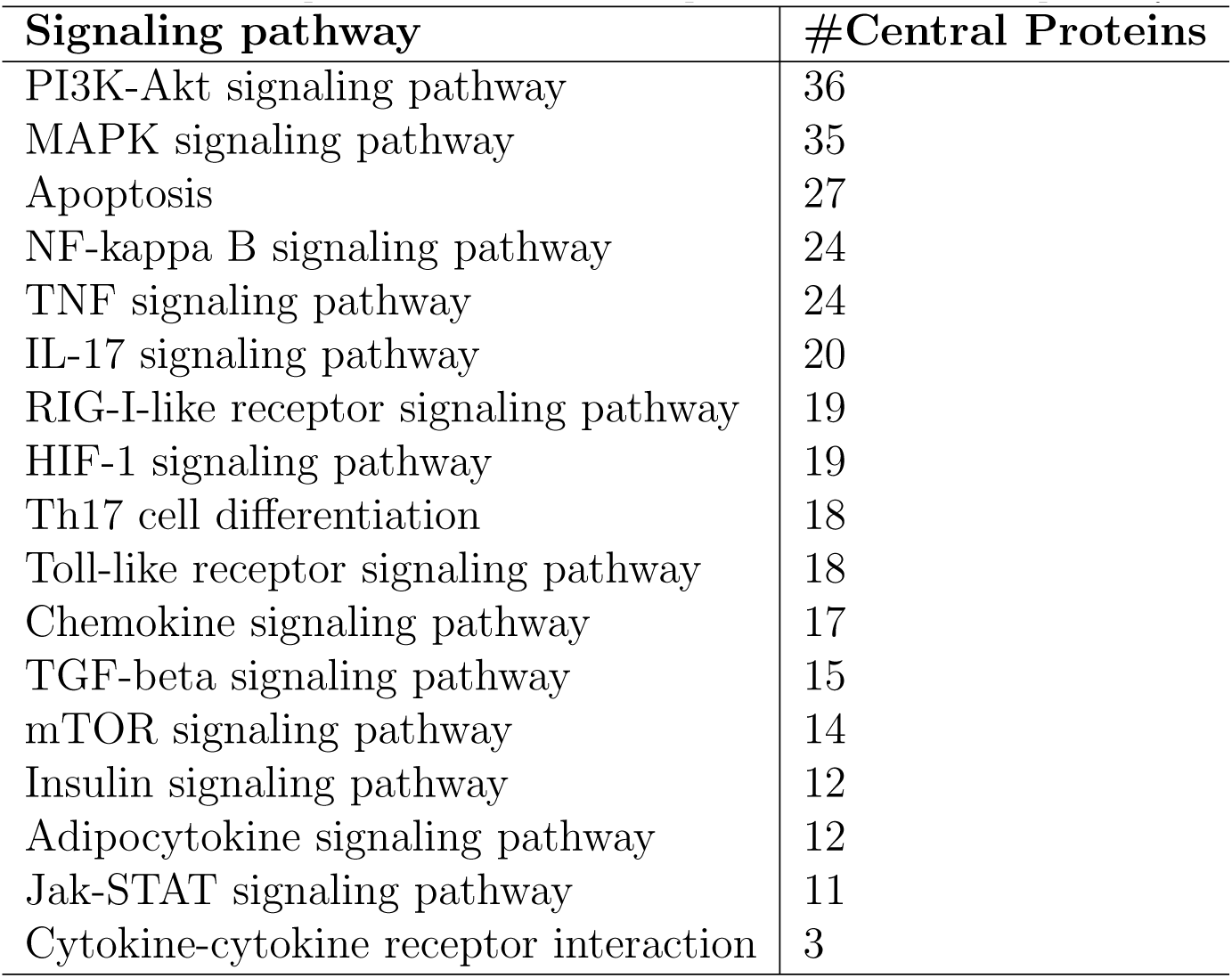
Participation count of central proteins in candidate pathways.

Interestingly, in terms of the number of targeted proteins and which are also central in host-host PPI, the signaling pathway MAPK is one of the worst affected pathways among 17 candidate pathways.

Prior researches also identified an exciting fact that viral proteins target host proteins that are pathway central, i.e., participating in multiple pathways [48]. In addition to PPI centrality, we study the pathway centrality of the host proteins regarding our 17 signaling pathways. Degree distribution of host proteins in terms of their density of participation in 17 pathways is reported in Figure 7. We observe a nice power-law [45] like distribution where the majority of proteins are participating in only one pathway and fewer numbers are having high participation in multiple pathways. We list a few top highly pathway central proteins and few interesting facts in Table 5. The table shows that the pathway central proteins are also highly connected in their own PPI and mostly targeted by multiple viral proteins.

**Table 5:** Few top pathway central proteins with the number of pathways they participating (out of 17 pathways), PPI centrality score and number of viral proteins (Vp) targeting the proteins

| Protein | #Pathway centrality | PPI centrality | #Interacting Vp count |
| --- | --- | --- | --- |
| IKBKB | 13 | 552 | 2 |
| CHUK | 12 | 462 | 11 |
| MAPK3 | 12 | 337 | 16 |
| RELA | 12 | 859 | 10 |
| AKT1 | 11 | 886 | 11 |
| AKT2 | 11 | 113 | 12 |
| AKT3 | 11 | 61 | 15 |
| IKBKG | 11 | 959 | 9 |
| TNF | 11 | 497 | 11 |
| MAPK8 | 9 | 444 | 12 |
| MAPK9 | 9 | 260 | 15 |
| NFKBIA | 9 | 501 | 8 |
| PIK3CA | 9 | 190 | 19 |
| PIK3CB | 9 | 82 | 17 |
| PIK3CD | 9 | 28 | 17 |
| PIK3R1 | 9 | 684 | 5 |
| PIK3R2 | 9 | 190 | 16 |

**Figure 7:**
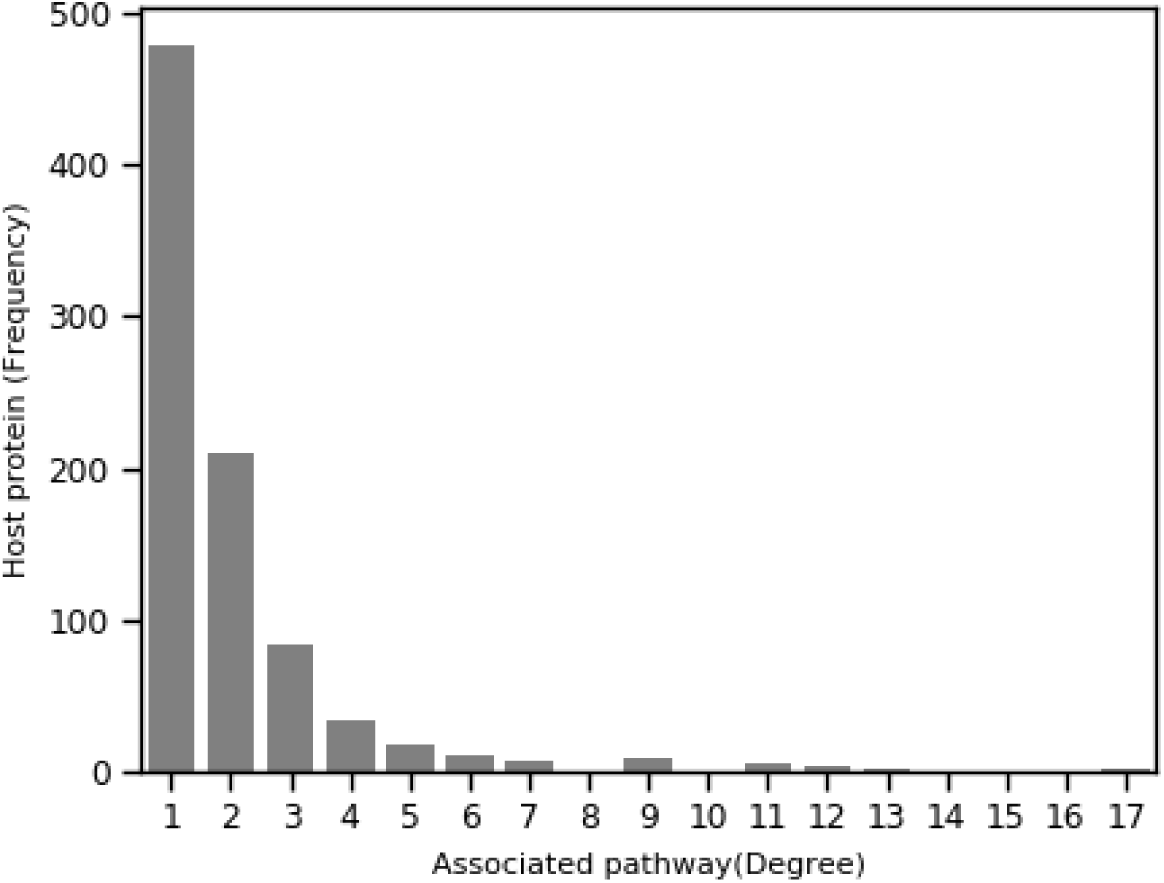
Degree distribution of 859 interacting host proteins in terms of number of associated signaling pathways (candidate).

## 4. Conclusion

In this work, we put a novel effort in recreating host-viral PPI using the codon usage pattern similarity between coding DNA sequences of host proteins that are participating in a few major signaling pathways and SARS-CoV2 viral proteins. We inferred both positive and negative edges between interacting proteins, which depict an important association between viral and host proteins. We analyze our inferred network topologically concerning degree distribution and node centrality. We observed interesting facts on how viral proteins are targeting their host proteins.

We restricted our current study on a few significant signaling pathways. Our method is generic and useful to draw a more extensive network covering all important pathways in the future.

## Supplementary Materials

**Supplementary-A:** The pathway wise list of genes involving 17 signaling pathways.

**Supplementary-B:** The host protein, each of which interacts with one or more viral interacting proteins and their list.

**Supplementary-C:** The list of top score 100 central host protein. For each protein, centrality score, correlation, interacting viral protein, and involved pathways shown in different columns.

## Author contributions

J.K.D. and S.R. conceived and designed the study. J.K.D. performed computational work, S.C. interpreted Biological validation. J.K.D. and S.R. were in charge for overall direction, planning, and supervision. All authors participate in planing, discussion and provided critical feedback. All authors participate in writing original manuscript and approved.

## Footnotes

1 www.genome.jp/kegg/pathway.html

2 https://cai.readthedocs.io/en/latest/

3 https://scipy.org

4 https://scipy.org

5 https://scipy.org/

6 https://thebiogrid.org/

