## Supplementary-A for "Impact Analysis of SARS-CoV2 on Signaling Pathways during COVID19 Pathogenesis using Codon Usage Assisted Host-Viral Protein Interactions"

**Table S1: The pathway wise list of genes involving 17 signaling pathways.**

| Pathway name | Gene count | Gene list |
|---|---|---|
| NF-kappa B signaling pathway | 105 | LCK, ZAP70, LAT, PLCG1, PRKCQ, IGH, SYK, LYN, BLNK, BTK, PLCG2, PRKCB, CARD10, CARD11, CARD14, BCL10, MALT1, IL1B, IL1R1, MYD88, IRAK1, IRAK4, TRAF6, TNF, TNFRSF1A, RIPK1, TRADD, TRAF2, TRAF5, BIRC2, BIRC3, EDA, EDAR, EDARADD, CYLD, EDA2R, DDX58, TRIM25, LBP, CD14, TLR4, LY96, TIRAP, TICAM2, TICAM1, CD40LG, CD40, TRAF3, TNFSF11, TNFRSF11A, LTA, LTB, TNFSF14, LTBR, MAP3K14, MAP3K7, TAB1, TAB2, TAB3, TNFSF13B, TNFRSF13C, IKBKG, CHUK, IKBKB, PARP1, PIAS4, UBE2I, ATM, PIDD1, ERC1, NFKBIA, NFKB1, RELA, CFLAR, XIAP, BCL2L1, BCL2, GADD45A, GADD45B, GADD45G, TRAF1, BCL2A1, NFKB2, CXCL8, TNFAIP3, PTGS2, CCL4, CCL4L2, CCL4L1, CXCL1, CXCL2, CXCL3, VCAM1, PLAU, CSNK2A1, CSNK2A2, CSNK2A3, CSNK2B, RELB, CCL13, CCL19, CCL21, CXCL12, ICAM1 |
| Cytokine-cytokine receptor interaction | 295 | CCL1, CCL25, CCL19, CCL21, CCL3, CCL3L1, CCL3L3, CCL4, CCL4L2, CCL4L1, CCL17, CCL22, CCL5, CCL8, CCL14, CCL16, CCL15, CCL23, CCL13, CCL7, CCL2, CCL11, CCL24, CCL26, CCL27, CCL28, CCL18, CCL20, CXCL1, CXCL2, CXCL3, CXCL5, CXCL6, CXCL8, PPBP, PF4, PF4V1, CXCL9, CXCL10, CXCL11, CXCL13, CXCL12, CXCL16, CXCL14, CXCL17, XCL1, XCL2, CX3CL1, IL2, IL4, IL7, IL9, IL15, IL21, TSLP, IL3, IL5, CSF2, IL13, EPO, GH1, GH2, CSH1, CSH2, PRL, THPO, CSF3, LEP, IL6, IL11, IL12A, IL12B, IL23A, IL27, IL31, CLCF1, CNTF, CTF1, LIF, OSM, IL10, IL19, IL20, IL24, IL22, IL26, IFNL2, IFNL3, IFNL1, IFNA1, IFNA2, IFNA4, IFNA5, IFNA6, IFNA7, IFNA8, IFNA10, IFNA13, IFNA14, IFNA16, IFNA17, IFNA21, IFNB1, IFNW1, IFNK, IFNE, IFNG, IL1A, IL1B, IL1RN, IL36RN, IL36A, IL36B, IL36G, IL1F10, IL37, IL18, IL33, IL17A, IL17F, IL17B, IL17C, IL17D, IL25, IL16, IL32, IL34, CSF1, TNF, LTA, LTB, TNFSF14, FASLG, TNFSF15, TNFSF10, EDA, NGF, TNFSF11, TNFSF12, CD70, TNFSF8, CD40LG, TNFSF9, TNFSF4, TNFSF18, TNFSF13, TNFSF13B, TGFB1, TGFB2, TGFB3, GDF15, GDF2, BMP10, INHA, BMP3, GDF10, GDF11, MSTN, INHBA, INHBB, GDF1, GDF3, NODAL, GDF9, INHBC, INHBE, AMH, BMP2, BMP4, GDF5, GDF6, GDF7, BMP15, BMP5, BMP6, BMP7, BMP8B, BMP8A, CCR8, CCR9, ACKR4, CCR7, CCR4, CCR5, CCR3, CCR2, CCR1, CCR10, CCR6, CXCR1, CXCR2, CXCR3, CXCR5, CXCR4, ACKR3, CXCR6, XCR1, CX3CR1, IL2RA, IL2RB, IL2RG, IL4R, IL7R, IL9R, IL15RA, IL21R, CRLF2, IL3RA, CSF2RB, IL5RA, CSF2RA, IL13RA1, IL13RA2, EPOR, GHR, PRLR, MPL, CSF3R, LEPR, IL6R, IL6ST, IL11RA, IL12RB1, IL12RB2, IL23R, IL27RA, IL31RA, CNTFR, LIFR, OSMR, IL10RA, IL10RB, IL20RA, IL20RB, IL22RA1, IFNLR1, IFNAR1, IFNAR2, IFNGR1, IFNGR2, IL1R1, IL1RAP, IL1R2, IL1RL2, IL18R1, IL18RAP, IL1RL1, IL17RA, IL17RC, IL17RB, IL17RE, CD4, CSF1R, TNFRSF1A, TNFRSF1B, LTBR, TNFRSF14, TNFRSF6B, FAS, TNFRSF25, TNFRSF10A, TNFRSF10B, TNFRSF10C, TNFRSF10D, TNFRSF21, EDAR, EDA2R, NGFR, TNFRSF11B, TNFRSF11A, TNFRSF12A, CD27, TNFRSF8, CD40, TNFRSF9, TNFRSF4, TNFRSF18, TNFRSF17, TNFRSF13B, TNFRSF13C, TNFRSF19, RELT, TGFBR1, TGFBR2, ACVRL1, ACVR2A, BMPR2, ACVR2B, ACVR1B, ACVR1C, ACVR1, AMHR2, BMPR1A, BMPR1B |
| TNF signaling pathway | 113 | TNF, TNFRSF1A, BAG4, TRADD, TRAF2, TRAF5, RIPK1, BIRC2, BIRC3, MAP3K7, TAB1, TAB2, TAB3, MAP2K4, MAP2K7, MAPK8, MAPK10, MAPK9, JUN, FOS, ITCH, CFLAR, MAP2K3, MAP2K6, MAPK11, MAPK12, MAPK13, MAPK14, CEBPB, MAP3K5, MAP3K14, IKBKG, IKBKB, CHUK, NFKBIA, RELA, NFKB1, MAP3K8, MAP2K1, MAPK1, MAPK3, RPS6KA5, RPS6KA4, CREB1, CREB3, CREB3L1, CREB3L2, CREB3L3, CREB3L4, ATF2, ATF4, CREB5, ATF6B, RIPK3, MLKL, PGAM5, DNM1L, FADD, CASP8, CASP10, CASP7, CASP3, CCL2, CCL5, CCL20, CXCL1, CXCL2, CXCL3, CXCL5, CXCL6, CXCL10, CX3CL1, CSF1, CSF2, FAS, IL18R1, JAG1, IL1B, IL6, IL15, LIF, LTA, BCL3, SOCS3, TNFAIP3, TRAF1, JUNB, MMP3, MMP9, |

| | | |
|---|---|---|
| | | MMP14, EDN1, VEGFC, VEGFD, NOD2, ICAM1, SELE, VCAM1, PTGS2, TNFRSF1B, TRAF3, PIK3CA, PIK3CD, PIK3CB, PIK3R1, PIK3R2, PIK3R3, AKT1, AKT2, AKT3, DAB2IP, IRF1, IFNB1 |
| IL-17 signaling pathway | 95 | IL25, IL17RA, IL17RB, TRADD, FADD, CASP3, CASP8, TRAF3IP2, TRAF6, NFKB1, RELA, FOS, FOSB, JUN, JUND, FOSL1, IL4, IL5, IL13, CCL17, CCL11, IL17A, IL17F, IL17RC, TRAF3, ANAPC5, TNFAIP3, HSP90AA1, HSP90AB1, HSP90B1, TBK1, TAB2, TAB3, MAP3K7, IKBKG, CHUK, IKBKB, NFKBIA, MAPK11, MAPK12, MAPK13, MAPK14, MAPK8, MAPK10, MAPK9, MAPK1, MAPK3, MAPK4, MAPK6, MAPK7, MAPK15, CEBPB, TRAF4, IKBKE, USP25, TRAF5, TRAF2, SRSF1, ELAVL1, GSK3B, CXCL1, CXCL2, CXCL3, CXCL5, CXCL6, CXCL8, CXCL10, CCL2, CCL7, CCL20, IL6, TNF, PTGS2, CSF3, CSF2, DEFB4A, DEFB4B, MUC5AC, MUC5B, S100A7, S100A7A, S100A8, S100A9, LCN2, MMP1, MMP3, MMP9, MMP13, IL17C, IL17RE, IL1B, IFNG, IL17B, IL17D |
| RIG-I-like receptor signaling pathway | 70 | DDX58, IFIH1, MAVS, DHX58, TRAF3, TANK, AZI2, TBKBP1, IKBKG, TBK1, IKBKE, IRF3, IRF7, IFNA1, IFNA2, IFNA4, IFNA5, IFNA6, IFNA7, IFNA8, IFNA10, IFNA13, IFNA14, IFNA16, IFNA17, IFNA21, IFNB1, IFNW1, IFNE, IFNK, TRADD, FADD, RIPK1, CASP8, CASP10, CHUK, IKBKB, NFKBIB, NFKBIA, NFKB1, RELA, TRAF2, MAP3K7, TRAF6, MAP3K1, MAPK8, MAPK10, MAPK9, MAPK11, MAPK12, MAPK13, MAPK14, CXCL8, TNF, IL12A, IL12B, CXCL10, TRIM25, CYLD, RNF125, ISG15, ATG5, ATG12, NLRX1, STING1, OTUD5, SIKE1, DDX3X, PIN1 |
| MAPK signaling pathway | 295 | CACNA1A, CACNA1B, CACNA1C, CACNA1D, CACNA1E, CACNA1F, CACNA1G, CACNA1H, CACNA1I, CACNA1S, CACNA2D1, CACNA2D2, CACNA2D3, CACNA2D4, CACNB1, CACNB2, CACNB3, CACNB4, CACNG1, CACNG2, CACNG3, CACNG4, CACNG5, CACNG6, CACNG7, CACNG8, PRKACA, PRKACB, PRKACG, PRKCA, PRKCB, PRKCG, GNA12, GNG12, PPP3CA, PPP3CB, PPP3CC, PPP3R1, PPP3R2, RASGRF1, RASGRF2, RASGRP1, RASGRP2, RASGRP3, RASGRP4, RAPGEF2, NF1, RASA1, RASA2, RAP1A, RAP1B, EGF, TGFA, EREG, AREG, FGF1, FGF2, FGF3, FGF4, FGF17, FGF6, FGF7, FGF8, FGF9, FGF10, FGF16, FGF5, FGF18, FGF20, FGF22, FGF19, FGF21, FGF23, NGF, BDNF, NTF3, NTF4, INS, IGF1, IGF2, PDGFA, PDGFB, PDGFC, PDGFD, CSF1, KITLG, FLT3LG, VEGFA, VEGFB, PGF, VEGFC, VEGFD, HGF, ANGPT1, ANGPT2, ANGPT4, EFNA1, EFNA2, EFNA3, EFNA4, EFNA5, EGFR, ERBB2, ERBB3, ERBB4, FGFR1, FGFR2, FGFR3, FGFR4, NGFR, NTRK1, NTRK2, INSR, IGF1R, PDGFRA, PDGFRB, CSF1R, KIT, FLT3, FLT1, FLT4, KDR, MET, TEK, EPHA2, GRB2, SOS1, SOS2, HRAS, KRAS, NRAS, RRAS, RRAS2, MRAS, ARAF, BRAF, RAF1, MAP2K1, MAP2K2, LAMTOR3, MAPK1, MAPK3, MKNK1, MKNK2, RPS6KA3, RPS6KA1, RPS6KA2, RPS6KA6, ATF4, ELK1, ELK4, MYC, SRF, FOS, MAPT, STMN1, PLA2G4E, PLA2G4A, JMJD7-PLA2G4B, PLA2G4B, PLA2G4C, PLA2G4D, PLA2G4F, TNF, IL1A, IL1B, TGFB1, TGFB2, TGFB3, TNFRSF1A, IL1R1, IL1RAP, TGFBR1, TGFBR2, FASLG, FAS, CD14, RAC1, RAC2, RAC3, CDC42, TRADD, CASP3, TRAF2, DAXX, MYD88, IRAK1, IRAK4, TRAF6, GADD45A, GADD45B, GADD45G, TAB1, TAB2, ECSIT, MAP4K3, MAP4K4, MAP4K1, PAK1, PAK2, STK4, STK3, MAP4K2, MAP3K8, MAP3K1, MAP3K11, MAP3K2, MAP3K3, MAP3K13, MAP3K12, MAP3K20, MAP3K6, MAP3K5, MAP3K7, MAP3K4, TAOK2, TAOK3, TAOK1, MAP2K4, MAP2K7, MAP2K3, MAP2K6, MAPK8IP1, MAPK8IP2, MAPK8IP3, FLNA, FLNC, FLNB, CRK, CRKL, ARRB1, ARRB2, MAPK8, MAPK10, MAPK9, MAPK11, MAPK12, MAPK13, MAPK14, MAPKAPK5, MAPKAPK2, MAPKAPK3, RPS6KA5, RPS6KA4, CDC25B, NFATC1, NFATC3, JUN, JUND, ATF2, TP53, DDIT3, MAX, MEF2C, HSPB1, AKT1, AKT2, AKT3, PPM1A, PTPRR, PTPN5, PTPN7, DUSP1, DUSP4, DUSP2, DUSP7, DUSP8, DUSP5, DUSP16, DUSP6, DUSP9, DUSP10, DUSP3, PPP5C, PPM1B, HSPA8, HSPA1A, HSPA2, HSPA1L, HSPA1B, HSPA6, MECOM, MAP2K5, MAPK7, NR4A1, MAP3K14, CHUK, IKBKB, IKBKG, NLK, NFKB1, NFKB2, RELA, RELB, |
| Chemokine signaling pathway | 190 | CXCL1, CXCL2, CXCL3, CXCL5, CXCL6, PPBP, CXCL8, CXCL9, CXCL10, CXCL11, CXCL12, CXCL13, CXCL16, PF4, PF4V1, CXCL14, XCL1, XCL2, CX3CL1, CCL1, CCL2, CCL3, CCL3L1, CCL3L3, CCL4, CCL4L2, CCL4L1, CCL5, CCL7, CCL8, CCL11, CCL13, CCL14, CCL15, CCL23, CCL16, CCL17, CCL18, CCL19, CCL20, CCL21, CCL22, CCL24, CCL25, CCL26, CCL27, CCL28, CXCR2, CXCR1, CXCR3, CXCR4, CXCR5, CXCR6, XCR1, CX3CR1, CCR8, CCR6, CCR9, CCR4, CCR7, CCR2, CCR5, CCR1, CCR3, CCR10, JAK2, JAK3, STAT1, STAT2, STAT3, STAT5B, GNAI1, GNAI3, GNAI2, ADCY1, ADCY2, ADCY3, ADCY4, ADCY5, ADCY6, ADCY7, ADCY8, ADCY9, PRKACA, PRKACB, PRKACG, LYN, HCK, FGR, SRC, SHC1, SHC2, SHC3, SHC4, GRB2, SOS1, SOS2, HRAS, KRAS, NRAS, RAF1, BRAF, MAP2K1, MAPK1, MAPK3, PIK3CA, PIK3CD, PIK3CB, PIK3R1, PIK3R2, PIK3R3, PIK3CG, PIK3R5, PIK3R6, PRKCZ, AKT1, AKT2, AKT3, FOXO3, CHUK, IKBKB, IKBKG, NFKBIA, NFKBIB, NFKB1, RELA, BAD, GSK3A, GSK3B, ITK, VAV3, VAV1, VAV2, RAC1, RAC2, RAC3, PAK1, CDC42, WAS, RHOA, ROCK1, ROCK2, GNB1, GNB2, GNB3, GNB4, GNB5, GNG2, GNG3, GNG4, GNG5, GNG7, GNG8, GNG10, GNG11, GNG12, GNG13, GNGT1, GNGT2, PREX1, ELMO1, DOCK2, PTK2, PXN, BCAR1, CRK, CRKL, PTK2B, PLCB1, PLCB2, PLCB3, PLCB4, RASGRP2, RAP1A, RAP1B, PARD3, TIAM1, PRKCB, PRKCD, NCF1, GRK7, GRK1, GRK2, GRK3, GRK4, GRK5, GRK6, ARRB1, ARRB2 |
| PI3K-Akt signaling pathway | 355 | EGF, TGFA, EREG, AREG, FGF1, FGF2, FGF3, FGF4, FGF17, FGF6, FGF7, FGF8, FGF9, FGF10, FGF16, FGF5, FGF18, FGF20, FGF22, FGF19, FGF21, FGF23, NGF, BDNF, NTF3, NTF4, INS, IGF1, IGF2, PDGFA, PDGFB, PDGFC, PDGFD, CSF1, KITLG, FLT3LG, VEGFA, VEGFB, PGF, VEGFC, VEGFD, HGF, ANGPT1, ANGPT2, ANGPT4, EFNA1, EFNA2, EFNA3, EFNA4, EFNA5, |

| | | |
|---|---|---|
| | | EGFR, ERBB2, ERBB3, ERBB4, FGFR1, FGFR2, FGFR3, FGFR4, NGFR, NTRK1, NTRK2, INSR, IGF1R, PDGFRA, PDGFRB, CSF1R, KIT, FLT3, FLT1, FLT4, KDR, MET, TEK, EPHA2, GRB2, SOS1, SOS2, HRAS, KRAS, NRAS, RAF1, MAP2K1, MAP2K2, MAPK1, MAPK3, IRS1, TLR2, TLR4, RAC1, IGH, SYK, CD19, PIK3AP1, GH1, GH2, CSH1, CSH2, PRL, OSM, IL2, IL3, IL6, IL4, IL7, IFNA1, IFNA2, IFNA4, IFNA5, IFNA6, IFNA7, IFNA8, IFNA10, IFNA13, IFNA14, IFNA16, IFNA17, IFNA21, IFNB1, EPO, CSF3, GHR, PRLR, OSMR, IL2RA, IL2RB, IL2RG, IL3RA, IL6R, IL4R, IL7R, IFNAR1, IFNAR2, EPOR, CSF3R, JAK1, JAK2, JAK3, COL1A1, COL1A2, COL2A1, COL4A2, COL4A4, COL4A6, COL4A1, COL4A5, COL4A3, COL6A1, COL6A2, COL6A3, COL6A6, COL6A5, COL9A1, COL9A2, COL9A3, LAMA1, LAMA2, LAMA3, LAMA5, LAMA4, LAMB1, LAMB2, LAMB3, LAMB4, LAMC1, LAMC2, LAMC3, CHAD, RELN, THBS1, COMP, THBS2, THBS3, THBS4, FN1, SPP1, VTN, TNC, TNN, TNR, TNXB, VWF, IBSP, ITGA1, ITGA2, ITGA2B, ITGA3, ITGA4, ITGA5, ITGA6, ITGA7, ITGA8, ITGA9, ITGA10, ITGA11, ITGAV, ITGB1, ITGB3, ITGB4, ITGB5, ITGB6, ITGB7, ITGB8, PTK2, PIK3CA, PIK3CD, PIK3CB, PIK3R1, PIK3R2, PIK3R3, F2R, CHRM1, CHRM2, LPAR1, LPAR2, LPAR3, LPAR4, LPAR5, LPAR6, GNB1, GNB2, GNB3, GNB4, GNB5, GNG2, GNG3, GNG4, GNG5, GNG7, GNG8, GNG10, GNG11, GNG12, GNG13, GNGT1, GNGT2, PIK3CG, PIK3R5, PIK3R6, PDPK1, STK11, PRKAA1, PRKAA2, DDIT4, TSC1, TSC2, RHEB, MLST8, MTOR, RPTOR, EIF4EBP1, EIF4E, EIF4E2, EIF4E1B, RPS6KB1, RPS6KB2, EIF4B, RPS6, PRKCA, PKN1, PKN2, PKN3, SGK1, SGK2, SGK3, C8orf44-SGK3, AKT1, AKT2, AKT3, MAGI1, MAGI2, PTEN, THEM4, PPP2CA, PPP2CB, PPP2R1B, PPP2R1A, PPP2R2A, PPP2R2B, PPP2R2C, PPP2R2D, PPP2R3B, PPP2R3C, PPP2R3A, PPP2R5B, PPP2R5C, PPP2R5D, PPP2R5E, PPP2R5A, HSP90AA1, HSP90AB1, HSP90B1, CDC37, CRTC2, PHLPP1, PHLPP2, TCL1A, TCL1B, MTCP1, NOS3, BRCA1, GSK3B, GYS2, GYS1, PCK1, PCK2, G6PC, G6PC2, G6PC3, MYC, CCND1, CDKN1A, CDKN1B, CDK2, CDK4, CDK6, CCND2, CCND3, CCNE1, CCNE2, FOXO3, RBL2, FASLG, BCL2L11, YWHAZ, YWHAB, YWHAQ, YWHAE, YWHAH, YWHAG, BAD, BCL2L1, BCL2, CASP9, CREB1, ATF2, ATF4, CREB3, CREB3L1, CREB3L2, CREB3L3, CREB3L4, CREB5, ATF6B, MCL1, RXRA, NR4A1, IKBKG, CHUK, IKBKB, RELA, NFKB1, MYB, MDM2, TP53 |
| Jak-STAT signaling pathway | 163 | IL2, IL3, IL4, IL5, IL6, IL7, IL9, IL10, IL11, IL12A, IL12B, IL13, IL15, IL17D, IL19, IL20, IL21, IL22, IL23A, IL24, IFNA1, IFNA2, IFNA4, IFNA5, IFNA6, IFNA7, IFNA8, IFNA10, IFNA13, IFNA14, IFNA16, IFNA17, IFNA21, IFNB1, IFNG, IFNE, IFNK, IFNL1, IFNL2, IFNL3, IFNW1, OSM, LIF, TSLP, CTF1, CSF2, CNTF, CSF3, EPO, GH1, GH2, CSH1, CSH2, LEP, THPO, PRL, EGF, PDGFA, PDGFB, IL2RA, IL2RB, IL2RG, IL3RA, IL4R, IL5RA, IL6R, IL7R, IL9R, IL10RA, IL10RB, IL11RA, IL12RB1, IL12RB2, IL13RA1, IL13RA2, IL15RA, IL20RA, IL20RB, IL21R, IL22RA1, IL22RA2, IL23R, IL27RA, IL6ST, IFNAR1, IFNAR2, IFNGR1, IFNGR2, IFNLR1, OSMR, LIFR, CRLF2, CNTFR, CSF2RA, CSF2RB, CSF3R, EPOR, GHR, LEPR, MPL, PRLR, EGFR, PDGFRA, PDGFRB, JAK1, JAK2, JAK3, TYK2, STAT1, STAT2, STAT3, STAT4, STAT5A, STAT5B, STAT6, CISH, SOCS1, SOCS2, SOCS3, SOCS4, SOCS5, SOCS7, SOCS6, BCL2, MCL1, BCL2L1, PIM1, MYC, CCND1, CCND2, CCND3, CDKN1A, AOX1, GFAP, STAM2, STAM, PTPN2, PTPN6, IRF9, CREBBP, EP300, PIAS1, PIAS2, PIAS3, PIAS4, FHL1, PTPN11, GRB2, SOS1, SOS2, HRAS, RAF1, PIK3CA, PIK3CD, PIK3CB, PIK3R1, PIK3R2, PIK3R3, AKT1, AKT2, AKT3, MTOR |
| Th17 cell differentiation | 108 | IL1B, IL1R1, IL1RAP, MAPK11, MAPK12, MAPK13, MAPK14, MTOR, IRF4, TGFB1, TGFBR1, TGFBR2, SMAD2, SMAD3, SMAD4, IL21, IL21R, IL2RG, JAK1, JAK3, IL6, IL6R, IL6ST, JAK2, IL23A, IL23R, IL12RB1, TYK2, STAT3, RORC, RORA, HIF1A, HSP90AA1, HSP90AB1, AHR, IL17A, IL17F, IL22, HLA-DMA, HLA-DMB, HLA-DOA, HLA-DOB, HLA-DPA1, HLA-DPB1, HLA-DQA1, HLA-DQA2, HLA-DQB1, HLA-DRA, HLA-DRB1, HLA-DRB3, HLA-DRB4, HLA-DRB5, CD4, LCK, CD3E, CD3G, CD247, CD3D, ZAP70, LAT, PLCG1, PPP3CA, PPP3CB, PPP3CC, PPP3R1, PPP3R2, NFATC1, NFATC2, NFATC3, PRKCQ, CHUK, IKBKB, IKBKG, NFKBIA, NFKBIB, NFKBIE, NFKB1, RELA, MAPK1, MAPK3, FOS, MAPK8, MAPK10, MAPK9, JUN, IL4, IL4R, STAT6, GATA3, RUNX1, IL17D, IL27RA, IFNG, IFNGR1, IFNGR2, STAT1, TBX21, IL2, IL2RA, IL2RB, STAT5A, STAT5B, FOXP3, RARA, RXRA, RXRB, RXRG |
| TGF-beta signaling pathway | 95 | CHRD, NOG, NBL1, MICOS10-NBL1, GREM1, GREM2, THBS1, DCN, FMOD, LEFTY1, LEFTY2, FST, BMP2, BMP4, BMP6, INHBB, BMP5, BMP7, BMP8B, BMP8A, GDF5, GDF6, GDF7, AMH, THSD4, FBN1, LTBP1, TGFB1, TGFB2, TGFB3, INHBA, INHBC, INHBE, NODAL, NEO1, HJV, BMPR1A, BMPR1B, ACVR1, BMPR2, ACVR2A, RGMA, RGMB, AMHR2, TGFBR1, TGFBR2, ACVR1B, ACVR2B, ACVR1C, BAMBI, SMAD1, SMAD5, SMAD9, SMAD2, SMAD3, SMAD4, SMAD6, SMAD7, SMURF1, SMURF2, ZFYVE9, ZFYVE16, HAMP, ID1, ID2, ID3, ID4, RBL1, E2F4, E2F5, TFDP1, CREBBP, EP300, SP1, TGIF1, TGIF2, MYC, CDKN2B, PITX2, RBX1, CUL1, SKP1, MAPK1, MAPK3, IFNG, TNF, RHOA, ROCK1, PPP2R1B, PPP2R1A, PPP2CA, PPP2CB, RPS6KB1, RPS6KB2 |
| Toll-like receptor signaling pathway | 105 | TLR1, TLR2, TLR6, LBP, CD14, LY96, TLR3, TLR4, TLR5, TLR7, TLR8, CTSK, TLR9, RAC1, PIK3CA, PIK3CD, PIK3CB, PIK3R1, PIK3R2, PIK3R3, AKT1, AKT2, AKT3, TOLLIP, MYD88, TIRAP, FADD, CASP8, IRAK4, IRAK1, TRAF6, TAB1, TAB2, MAP3K7, IKBKG, CHUK, IKBKB, NFKBIA, NFKB1, RELA, MAP3K8, MAP2K1, MAP2K2, MAPK1, MAPK3, MAP2K3, MAP2K6, MAP2K4, MAP2K7, MAPK11, MAPK12, MAPK13, MAPK14, MAPK8, MAPK10, MAPK9, JUN, FOS, TNF, IL1B, IL6, IL12A, IL12B, CXCL8, CCL5, CCL3, CCL3L1, CCL3L3, CCL4, CCL4L2, CCL4L1, TICAM2, TICAM1, RIPK1, IRF5, IRF7, SPP1, IKBKE, TBK1, TRAF3, |

| | | |
|---|---|---|
| | | IRF3, CD40, CD80, CD86, IFNA1, IFNA2, IFNA4, IFNA5, IFNA6, IFNA7, IFNA8, IFNA10, IFNA13, IFNA14, IFNA16, IFNA17, IFNA21, IFNB1, IFNAR1, IFNAR2, STAT1, CXCL10, CXCL9, CXCL11 |
| HIF-1 signaling pathway | 110 | IL6, IL6R, STAT3, TLR4, IFNG, IFNGR1, IFNGR2, RELA, NFKB1, INS, EGF, IGF1, INSR, EGFR, IGF1R, ERBB2, MAP2K1, MAP2K2, MAPK1, MAPK3, MKNK1, MKNK2, PIK3CA, PIK3CD, PIK3CB, PIK3R1, PIK3R2, PIK3R3, AKT1, AKT2, AKT3, MTOR, EIF4EBP1, EIF4E, EIF4E2, EIF4E1B, RPS6KB1, RPS6KB2, RPS6, HIF1A, VHL, RBX1, ELOC, ELOB, CUL2, EGLN1, EGLN3, EGLN2, ARNT, CREBBP, EP300, CYBB, PLCG1, PLCG2, PRKCA, PRKCB, PRKCG, CAMK2A, CAMK2D, CAMK2B, CAMK2G, TIMP1, LTBR, EPO, TF, TFRC, VEGFA, FLT1, SERPINE1, ANGPT1, ANGPT2, ANGPT4, TEK, EDN1, NOS2, NOS3, HMOX1, NPPA, SLC2A1, PDK1, HK3, HK1, HK2, HKDC1, PFKM, PFKP, PFKL, GAPDH, ALDOC, ALDOA, ALDOB, ENO3, ENO2, ENO1, ENO4, PGK2, PGK1, PFKFB3, LDHAL6A, LDHAL6B, LDHA, LDHB, LDHC, BCL2, CDKN1A, CDKN1B, PDHA2, PDHA1, PDHB |
| Apoptosis | 137 | TNFSF10, TNFRSF10A, TNFRSF10B, FASLG, FAS, FADD, TNF, TNFRSF1A, TRADD, CFLAR, CASP8, CASP10, CASP6, CASP3, CASP7, BID, BAX, BAK1, DIABLO, SEPTIN4, HTRA2, CYCS, APAF1, CASP9, PRF1, GZMB, TUBA1B, TUBA4A, TUBA3C, TUBA1A, TUBA1C, TUBA8, TUBA3E, TUBA3D, TUBAL3, MCL1, ACTG1, ACTB, SPTA1, SPTAN1, LMNA, LMNB1, LMNB2, PARP1, PARP2, PARP3, PARP4, DFFA, DFFB, ENDOG, AIFM1, ERN1, TRAF2, ITPR1, ITPR2, ITPR3, CAPN1, CAPN2, CASP12, EIF2AK3, EIF2S1, ATF4, DDIT3, CTSB, CTSC, CTSD, CTSF, CTSH, CTSK, CTSL, CTSO, CTSS, CTSV, CTSW, CTSZ, BIRC2, BIRC3, XIAP, BIRC5, BCL2L11, BCL2L1, BCL2, DAXX, RIPK1, DAB2IP, MAP3K5, MAPK8, MAPK10, MAPK9, BAD, JUN, FOS, TP53, HRK, MAP3K14, CHUK, IKBKB, IKBKG, NFKBIA, NFKB1, RELA, PTPN13, GADD45A, GADD45B, GADD45G, TRAF1, BCL2A1, ATM, PIDD1, TP53AIP1, BBC3, PMAIP1, CASP2, NGF, NTRK1, IL3, IL3RA, CSF2RB, PIK3CA, PIK3CD, PIK3CB, PIK3R1, PIK3R2, PIK3R3, PDPK1, AKT1, AKT2, AKT3, HRAS, KRAS, NRAS, RAF1, MAP2K1, MAP2K2, MAPK1, MAPK3 |
| Insulin signaling pathway | 138 | INS, INSR, IRS1, IRS2, IRS4, PIK3R1, PIK3R2, PIK3R3, PIK3CA, PIK3CD, PIK3CB, PDPK1, AKT1, AKT2, AKT3, GSK3B, GYS2, GYS1, PPP1CA, PPP1CB, PPP1CC, PPP1R3A, PPP1R3C, PPP1R3D, PPP1R3B, PPP1R3E, PPP1R3F, PHKG1, PHKG2, PHKB, PHKA2, PHKA1, CALML3, CALM2, CALM3, CALM1, CALML6, CALML5, CALML4, PYGL, PYGM, PYGB, PDE3B, PRKACA, PRKACB, PRKACG, PRKAR1A, PRKAR2A, PRKAR2B, PRKAR1B, LIPE, PRKCZ, PRKCI, SLC2A4, FLOT2, FLOT1, SH2B2, SORBS1, CBL, CBLB, CRK, CRKL, RAPGEF1, RHOQ, EXOC7, TRIP10, SREBF1, ACACA, ACACB, FASN, PKLR, HK3, HK1, HK2, HKDC1, GCK, PRKAA1, PRKAA2, PRKAB1, PRKAB2, PRKAG1, PRKAG3, PRKAG2, FOXO1, PPARGC1A, G6PC, G6PC2, G6PC3, FBP1, FBP2, PCK1, PCK2, MTOR, RPTOR, RPS6KB1, RPS6KB2, RPS6, EIF4EBP1, EIF4E, EIF4E2, EIF4E1B, TSC1, TSC2, RHEB, BAD, SHC1, SHC2, SHC3, SHC4, GRB2, SOS1, SOS2, HRAS, KRAS, NRAS, ARAF, BRAF, RAF1, MAP2K1, MAP2K2, MAPK1, MAPK3, MKNK1, MKNK2, ELK1, SOCS1, SOCS2, SOCS3, SOCS4, PTPN1, PTPRF, MAPK8, MAPK10, MAPK9, IKBKB, INPPL1, INPP5A |
| mTOR signaling pathway | 156 | SLC7A5, SLC3A2, SLC38A9, ATP6V1A, ATP6V1B1, ATP6V1B2, ATP6V1C2, ATP6V1C1, ATP6V1D, ATP6V1E2, ATP6V1E1, ATP6V1F, ATP6V1G1, ATP6V1G3, ATP6V1G2, ATP6V1H, LAMTOR1, LAMTOR2, LAMTOR3, LAMTOR4, LAMTOR5, FLCN, FNIP1, FNIP2, RRAGA, RRAGB, RRAGC, RRAGD, SESN2, CASTOR1, CASTOR2, MIOS, SEH1L, WDR24, WDR59, SEC13, DEPDC5, NPRL2, NPRL3, SKP2, RNF152, RPTOR, AKT1S1, MTOR, DEPTOR, MLST8, TELO2, TTI1, CLIP1, GRB10, LPIN1, LPIN3, LPIN2, ULK1, ULK2, EIF4EBP1, EIF4E, EIF4E2, EIF4E1B, RPS6KB1, RPS6KB2, EIF4B, RPS6, STRADA, STRADB, STK11, CAB39, CAB39L, PRKAA1, PRKAA2, TSC1, TSC2, TBC1D7, TBC1D7-LOC100130357, RHEB, DDIT4, WNT1, WNT2, WNT2B, WNT3, WNT3A, WNT4, WNT5A, WNT5B, WNT6, WNT7A, WNT7B, WNT8A, WNT8B, WNT9A, WNT9B, WNT10B, WNT10A, WNT11, WNT16, FZD1, FZD7, FZD2, FZD3, FZD4, FZD5, FZD8, FZD6, FZD10, FZD9, LRP5, LRP6, DVL3, DVL2, DVL1, GSK3B, TNF, TNFRSF1A, IKBKB, INS, IGF1, INSR, IGF1R, GRB2, SOS1, SOS2, HRAS, KRAS, NRAS, BRAF, RAF1, MAP2K1, MAP2K2, MAPK1, MAPK3, RPS6KA3, RPS6KA1, RPS6KA2, RPS6KA6, IRS1, PIK3R1, PIK3R2, PIK3R3, PIK3CA, PIK3CD, PIK3CB, PTEN, PDPK1, AKT1, AKT2, AKT3, CHUK, MAPKAP1, RICTOR, PRR5, RHOA, PRKCA, PRKCB, PRKCG, SGK1 |
| Adipocytokine signaling pathway | 70 | TNF, TNFRSF1A, TRADD, TNFRSF1B, TRAF2, MTOR, MAPK8, MAPK10, MAPK9, CHUK, IKBKB, IKBKG, NFKBIA, NFKBIB, NFKBIE, NFKB1, RELA, SOCS3, IRS1, IRS2, IRS4, AKT1, AKT2, AKT3, CD36, ACSL6, ACSL4, ACSL1, ACSL5, ACSL3, ACSBG1, ACSBG2, PRKCQ, LEP, LEPR, JAK2, STAT3, POMC, PRKAA1, PRKAA2, PRKAB1, PRKAB2, PRKAG1, PRKAG3, PRKAG2, AGRP, NPY, PPARGC1A, PCK1, PCK2, G6PC, G6PC2, G6PC3, PTPN11, PPARA, RXRA, RXRB, RXRG, ADIPOQ, ADIPOR1, ADIPOR2, STK11, CAMKK2, ACACB, CPT1A, CPT1B, CPT1C, SLC2A1, SLC2A4 |
