## Supplementary-B for "Impact Analysis of SARS-CoV2 on Signaling Pathways during COVID19 Pathogenesis using Codon Usage Assisted Host-Viral Protein Interactions"

| Viral protein | Host protein | Correlation | p-val |
|---|---|---|---|
| S | PPP1R3A | 0.844 | 4.65E-17 |
| Nsp3 | PPP1R3A | 0.828 | 6.26E-16 |
| S | IRAK4 | 0.826 | 8.54E-16 |
| Nsp13 | PPP1R3A | 0.825 | 9.13E-16 |
| S | BIRC2 | 0.822 | 1.52E-15 |
| S | PKN2 | 0.817 | 3.04E-15 |
| Nsp13 | STAM2 | 0.814 | 4.51E-15 |
| Nsp3 | IRAK4 | 0.813 | 5.42E-15 |
| Nsp4 | IRAK4 | 0.809 | 8.76E-15 |
| Nsp14 | ROCK1 | 0.808 | 1.08E-14 |
| S | LEPR | 0.807 | 1.13E-14 |
| Nsp3 | RPS6KA3 | 0.803 | 2.00E-14 |
| Nsp13 | IL13RA2 | 0.802 | 2.38E-14 |
| Nsp16 | PPP1R3A | 0.802 | 2.23E-14 |
| S | RPS6KA3 | 0.802 | 2.40E-14 |
| S | STAM2 | 0.8 | 3.01E-14 |
| Nsp4 | PPP1R3A | 0.797 | 4.23E-14 |
| Nsp13 | IRAK4 | 0.796 | 5.14E-14 |
| Nsp3 | ATM | 0.793 | 7.43E-14 |
| Nsp3 | ZFYVE16 | 0.791 | 8.54E-14 |
| Nsp13 | LIFR | 0.791 | 8.65E-14 |
| S | ATM | 0.791 | 9.26E-14 |
| Nsp13 | PKN2 | 0.79 | 1.08E-13 |
| Nsp3 | PKN2 | 0.788 | 1.31E-13 |
| Nsp3 | ROCK1 | 0.787 | 1.52E-13 |
| S | LIFR | 0.787 | 1.47E-13 |
| Nsp3 | TBK1 | 0.786 | 1.75E-13 |
| Nsp3 | STAM2 | 0.786 | 1.68E-13 |
| Nsp12 | PPP1R3A | 0.786 | 1.57E-13 |
| Nsp3 | LEPR | 0.785 | 1.94E-13 |
| S | IL6ST | 0.784 | 1.99E-13 |
| Nsp3 | BIRC2 | 0.783 | 2.29E-13 |
| Nsp16 | IRAK4 | 0.783 | 2.25E-13 |
| Nsp3 | XIAP | 0.782 | 2.60E-13 |

| | | | |
|---|---|---|---|
| S | IFNAR1 | 0.782 | 2.54E-13 |
| Nsp3 | MAP4K3 | 0.781 | 2.94E-13 |
| Nsp14 | PPP1R3A | 0.781 | 2.99E-13 |
| Nsp16 | ZFYVE16 | 0.78 | 3.22E-13 |
| S | DNM1L | 0.78 | 3.23E-13 |
| Nsp15 | LIFR | 0.779 | 3.88E-13 |
| S | TBK1 | 0.778 | 4.27E-13 |
| S | BRCA1 | 0.777 | 4.57E-13 |
| Nsp3 | LIFR | 0.776 | 5.50E-13 |
| Nsp13 | RPS6KA3 | 0.775 | 5.63E-13 |
| Nsp14 | ATM | 0.774 | 6.72E-13 |
| Nsp3 | BRCA1 | 0.773 | 7.68E-13 |
| Nsp16 | MAP4K3 | 0.773 | 7.20E-13 |
| Nsp16 | LEPR | 0.773 | 7.04E-13 |
| S | ZFYVE16 | 0.772 | 8.51E-13 |
| Nsp4 | ROCK1 | 0.771 | 9.29E-13 |
| S | MDM2 | 0.77 | 1.07E-12 |
| Nsp16 | TBK1 | 0.769 | 1.19E-12 |
| S | SGK3 | 0.767 | 1.37E-12 |
| Nsp6 | ROCK1 | 0.766 | 1.52E-12 |
| Nsp16 | PPP3CB | 0.766 | 1.63E-12 |
| Nsp3 | RPS6KA6 | 0.765 | 1.74E-12 |
| Nsp3 | BIRC3 | 0.765 | 1.68E-12 |
| Nsp3 | SOS2 | 0.764 | 1.90E-12 |
| Nsp4 | RPS6KA3 | 0.764 | 1.94E-12 |
| S | CD36 | 0.764 | 2.01E-12 |
| Nsp15 | TBK1 | 0.762 | 2.46E-12 |
| Nsp14 | IRAK4 | 0.761 | 2.60E-12 |
| Nsp15 | IRAK4 | 0.761 | 2.56E-12 |
| Nsp16 | PKN2 | 0.761 | 2.60E-12 |
| Nsp13 | MAP4K3 | 0.759 | 3.43E-12 |
| S | MAP4K3 | 0.759 | 3.43E-12 |
| S | PPM1B | 0.758 | 3.60E-12 |
| Nsp3 | USP25 | 0.756 | 4.64E-12 |
| Nsp16 | RPS6KA3 | 0.755 | 5.12E-12 |
| Nsp16 | ATM | 0.755 | 4.81E-12 |
| S | ROCK1 | 0.755 | 4.98E-12 |
| S | COL4A5 | 0.755 | 4.82E-12 |
| Nsp3 | STAM | 0.754 | 5.21E-12 |
| Nsp13 | ATM | 0.754 | 5.56E-12 |
| S | RPS6KA6 | 0.754 | 5.48E-12 |
| S | MIOS | 0.752 | 6.89E-12 |
| Nsp13 | MIOS | 0.751 | 7.54E-12 |
| S | SOS2 | 0.751 | 7.13E-12 |
| Nsp3 | MIOS | 0.749 | 8.92E-12 |
| Nsp3 | COL4A5 | 0.749 | 8.89E-12 |
| Nsp14 | TBK1 | 0.749 | 8.70E-12 |
| S | EIF2S1 | 0.748 | 9.78E-12 |
| Nsp8 | BIRC2 | 0.747 | 1.07E-11 |
| Nsp14 | ZFYVE16 | 0.747 | 1.11E-11 |

| | | | |
|---|---|---|---|
| S | BIRC3 | 0.747 | 1.12E-11 |
| Nsp4 | PKN2 | 0.746 | 1.24E-11 |
| Nsp4 | ATM | 0.746 | 1.22E-11 |
| Nsp12 | IRAK4 | 0.746 | 1.17E-11 |
| Nsp13 | LEPR | 0.746 | 1.20E-11 |
| S | APAF1 | 0.746 | 1.18E-11 |
| Nsp5 | IRAK4 | 0.745 | 1.38E-11 |
| Nsp3 | DNM1L | 0.744 | 1.46E-11 |
| Nsp8 | IRAK4 | 0.744 | 1.45E-11 |
| Nsp13 | ATF2 | 0.744 | 1.50E-11 |
| S | STAM | 0.744 | 1.40E-11 |
| Orf3a | COL4A5 | 0.744 | 1.49E-11 |
| Nsp4 | ATF2 | 0.743 | 1.65E-11 |
| Nsp16 | MIOS | 0.743 | 1.54E-11 |
| Nsp12 | PKN2 | 0.742 | 1.77E-11 |
| Nsp13 | USP25 | 0.742 | 1.76E-11 |
| S | MAPK6 | 0.742 | 1.83E-11 |
| Nsp5 | PPP1R3A | 0.741 | 2.02E-11 |
| Nsp15 | PPP1R3A | 0.741 | 1.95E-11 |
| Nsp16 | BIRC2 | 0.741 | 1.98E-11 |
| Nsp3 | IL6ST | 0.74 | 2.22E-11 |
| Nsp12 | LEPR | 0.74 | 2.05E-11 |
| Nsp13 | RPS6KA6 | 0.74 | 2.17E-11 |
| Nsp13 | ZFYVE16 | 0.74 | 2.20E-11 |
| Nsp13 | BIRC3 | 0.74 | 2.18E-11 |
| Nsp14 | BIRC3 | 0.74 | 2.18E-11 |
| Nsp15 | MIOS | 0.74 | 2.08E-11 |
| Nsp16 | LIFR | 0.739 | 2.40E-11 |
| Nsp3 | ROCK2 | 0.738 | 2.54E-11 |
| Nsp13 | CUL2 | 0.738 | 2.55E-11 |
| S | ACVR2A | 0.738 | 2.57E-11 |
| Nsp3 | HIF1A | 0.737 | 2.96E-11 |
| Nsp16 | ROCK1 | 0.737 | 2.84E-11 |
| Nsp16 | BRCA1 | 0.737 | 2.96E-11 |
| S | ITCH | 0.737 | 2.71E-11 |
| Nsp4 | BIRC2 | 0.736 | 2.97E-11 |
| Nsp13 | DNM1L | 0.736 | 3.14E-11 |
| S | XIAP | 0.736 | 3.08E-11 |
| Orf7a | MSTN | 0.736 | 3.20E-11 |
| Nsp6 | ATM | 0.735 | 3.32E-11 |
| Nsp13 | TBK1 | 0.734 | 3.85E-11 |
| S | ATF2 | 0.734 | 3.81E-11 |
| Nsp3 | CUL2 | 0.733 | 4.05E-11 |
| Nsp4 | TBK1 | 0.733 | 4.04E-11 |
| Nsp13 | SOS2 | 0.732 | 4.36E-11 |
| Nsp3 | ITCH | 0.731 | 5.02E-11 |
| Nsp12 | STAM2 | 0.731 | 5.04E-11 |
| S | PLA2G4A | 0.731 | 4.98E-11 |
| Nsp12 | ATM | 0.73 | 5.62E-11 |
| Nsp3 | ATF2 | 0.729 | 5.76E-11 |

| | | | |
|---|---|---|---|
| Nsp8 | PKN2 | 0.729 | 5.79E-11 |
| Nsp3 | PPP3CB | 0.728 | 6.58E-11 |
| Nsp3 | IFNAR1 | 0.728 | 6.20E-11 |
| Nsp8 | MIOS | 0.728 | 6.53E-11 |
| Nsp8 | RPS6KA3 | 0.728 | 6.46E-11 |
| S | USP25 | 0.728 | 6.28E-11 |
| S | HIF1A | 0.728 | 6.72E-11 |
| Nsp4 | BRCA1 | 0.727 | 7.17E-11 |
| S | CUL2 | 0.727 | 6.90E-11 |
| Orf7a | TAB3 | 0.727 | 7.16E-11 |
| Nsp4 | ZFYVE16 | 0.726 | 7.69E-11 |
| Nsp12 | RPS6KA3 | 0.726 | 7.68E-11 |
| Nsp13 | HIF1A | 0.726 | 7.46E-11 |
| Nsp14 | BRCA1 | 0.726 | 7.84E-11 |
| Nsp16 | IFNAR1 | 0.726 | 7.37E-11 |
| Nsp3 | IL13RA2 | 0.725 | 8.11E-11 |
| Nsp3 | PLA2G4A | 0.725 | 8.17E-11 |
| Nsp13 | BIRC2 | 0.725 | 8.52E-11 |
| Nsp15 | PPP3CB | 0.725 | 8.45E-11 |
| Nsp4 | SOS2 | 0.724 | 9.16E-11 |
| Nsp16 | ITCH | 0.724 | 9.09E-11 |
| S | ROCK2 | 0.724 | 8.95E-11 |
| Nsp13 | PPM1B | 0.723 | 9.96E-11 |
| Nsp16 | BIRC3 | 0.723 | 1.02E-10 |
| Nsp12 | DNM1L | 0.722 | 1.12E-10 |
| Nsp12 | XIAP | 0.722 | 1.13E-10 |
| Nsp13 | XIAP | 0.722 | 1.09E-10 |
| Nsp13 | PRKACB | 0.721 | 1.16E-10 |
| Nsp3 | PRKACB | 0.72 | 1.28E-10 |
| Nsp3 | SGK3 | 0.72 | 1.28E-10 |
| Nsp4 | ITCH | 0.72 | 1.30E-10 |
| Nsp4 | BIRC3 | 0.72 | 1.36E-10 |
| Nsp16 | SOS1 | 0.72 | 1.27E-10 |
| S | IFNGR1 | 0.72 | 1.25E-10 |
| S | SOS1 | 0.72 | 1.31E-10 |
| Nsp3 | SOS1 | 0.719 | 1.37E-10 |
| Nsp13 | IL6ST | 0.719 | 1.38E-10 |
| Nsp13 | ITCH | 0.719 | 1.47E-10 |
| Nsp14 | PKN2 | 0.719 | 1.40E-10 |
| Nsp13 | RBL1 | 0.718 | 1.56E-10 |
| Nsp15 | HIF1A | 0.718 | 1.51E-10 |
| Nsp16 | SGK3 | 0.718 | 1.55E-10 |
| Nsp16 | XIAP | 0.718 | 1.59E-10 |
| Nsp12 | TBK1 | 0.717 | 1.74E-10 |
| Nsp12 | ROCK1 | 0.717 | 1.73E-10 |
| Nsp13 | COL4A5 | 0.717 | 1.73E-10 |
| Nsp16 | RPS6KA6 | 0.717 | 1.67E-10 |
| Nsp3 | MDM2 | 0.716 | 1.87E-10 |
| Nsp3 | IFNGR1 | 0.716 | 1.80E-10 |
| Nsp8 | MDM2 | 0.716 | 1.87E-10 |

| | | | |
|---|---|---|---|
| Nsp15 | ZFYVE16 | 0.716 | 1.79E-10 |
| S | FZD6 | 0.716 | 1.90E-10 |
| S | PIAS2 | 0.716 | 1.89E-10 |
| Nsp3 | PPM1B | 0.715 | 1.97E-10 |
| Nsp4 | MAP4K3 | 0.715 | 2.07E-10 |
| Nsp5 | BIRC2 | 0.715 | 2.02E-10 |
| Nsp15 | PKN2 | 0.715 | 1.92E-10 |
| Nsp15 | SOS2 | 0.715 | 2.03E-10 |
| Nsp4 | STAM2 | 0.714 | 2.24E-10 |
| Nsp13 | TGFBR1 | 0.714 | 2.22E-10 |
| S | PTPN13 | 0.714 | 2.10E-10 |
| Orf7a | MAP4K3 | 0.714 | 2.26E-10 |
| Orf7a | HSP90B1 | 0.713 | 2.44E-10 |
| Nsp3 | MAPK6 | 0.712 | 2.61E-10 |
| Nsp4 | LIFR | 0.712 | 2.66E-10 |
| S | IL13RA2 | 0.712 | 2.61E-10 |
| Nsp8 | PPP1R3A | 0.711 | 2.72E-10 |
| Nsp5 | PPM1B | 0.71 | 3.03E-10 |
| Nsp12 | IFNAR1 | 0.71 | 3.05E-10 |
| Nsp13 | ROCK2 | 0.71 | 3.02E-10 |
| S | PRKACB | 0.71 | 3.00E-10 |
| Nsp6 | PPP1R3A | 0.709 | 3.17E-10 |
| Nsp6 | BIRC3 | 0.709 | 3.24E-10 |
| Nsp14 | PLA2G4A | 0.708 | 3.68E-10 |
| Orf8 | PPP1R3A | 0.708 | 3.60E-10 |
| Nsp3 | TAB3 | 0.707 | 4.02E-10 |
| Nsp3 | CD36 | 0.707 | 3.98E-10 |
| Nsp13 | ROCK1 | 0.707 | 3.81E-10 |
| Nsp16 | USP25 | 0.707 | 3.98E-10 |
| S | RAP1B | 0.707 | 4.05E-10 |
| Nsp4 | LEPR | 0.706 | 4.13E-10 |
| Nsp16 | ROCK2 | 0.706 | 4.21E-10 |
| S | NFATC3 | 0.706 | 4.07E-10 |
| S | COL9A1 | 0.706 | 4.27E-10 |
| Nsp12 | BIRC3 | 0.705 | 4.58E-10 |
| Nsp14 | MSTN | 0.705 | 4.45E-10 |
| Nsp14 | ITCH | 0.705 | 4.56E-10 |
| Nsp14 | AHR | 0.704 | 5.09E-10 |
| Nsp16 | RAP1B | 0.704 | 5.10E-10 |
| S | TLR6 | 0.704 | 4.97E-10 |
| Nsp6 | BRCA1 | 0.703 | 5.41E-10 |
| Orf8 | ROCK1 | 0.703 | 5.42E-10 |
| Nsp2 | RPS6KA3 | 0.702 | 5.62E-10 |
| Nsp12 | LIFR | 0.702 | 5.84E-10 |
| Nsp13 | CD36 | 0.702 | 5.67E-10 |
| Nsp14 | ROCK2 | 0.702 | 5.89E-10 |
| Nsp16 | STAM2 | 0.702 | 5.96E-10 |
| S | TAB2 | 0.702 | 5.88E-10 |
| Orf8 | TBK1 | 0.702 | 5.72E-10 |
| Nsp6 | IFNGR1 | 0.701 | 6.45E-10 |

| | | | |
|---|---|---|---|
| Nsp9 | ROCK1 | 0.701 | 6.17E-10 |
| Nsp12 | MAP4K3 | 0.701 | 6.32E-10 |
| Nsp13 | SOS1 | 0.701 | 6.10E-10 |
| Nsp15 | ATM | 0.701 | 6.22E-10 |
| Nsp16 | STAM | 0.701 | 6.19E-10 |
| Nsp16 | CUL2 | 0.701 | 6.50E-10 |
| Nsp3 | APAF1 | 0.7 | 6.62E-10 |
| Nsp4 | COL4A5 | 0.7 | 6.99E-10 |
| Nsp8 | COL4A5 | 0.7 | 7.10E-10 |
| Nsp14 | PPP3CB | 0.7 | 6.84E-10 |
| Nsp14 | PPP2R3C | 0.7 | 6.73E-10 |
| S | ZFYVE9 | 0.7 | 6.86E-10 |
| Nsp6 | ROCK2 | 0.699 | 7.18E-10 |
| Nsp12 | COL4A5 | 0.699 | 7.27E-10 |
| S | HSP90B1 | 0.699 | 7.69E-10 |
| Nsp3 | FZD6 | 0.698 | 7.83E-10 |
| Nsp8 | CUL2 | 0.698 | 8.16E-10 |
| Nsp3 | ACVR2A | 0.697 | 8.77E-10 |
| Nsp4 | PPP3CB | 0.697 | 8.37E-10 |
| Nsp5 | MIOS | 0.697 | 8.70E-10 |
| Nsp6 | BIRC2 | 0.697 | 8.61E-10 |
| S | TAB3 | 0.697 | 8.40E-10 |
| Nsp12 | MIOS | 0.696 | 9.68E-10 |
| Nsp12 | ZFYVE16 | 0.696 | 9.29E-10 |
| Nsp14 | RPS6KA3 | 0.696 | 9.37E-10 |
| Nsp15 | LEPR | 0.696 | 9.03E-10 |
| Orf3a | CD36 | 0.696 | 9.04E-10 |
| Nsp3 | ZFYVE9 | 0.695 | 1.04E-09 |
| Nsp4 | STAM | 0.695 | 1.03E-09 |
| Nsp5 | RPS6KA6 | 0.695 | 1.02E-09 |
| Nsp13 | IFNAR1 | 0.695 | 1.04E-09 |
| S | RICTOR | 0.695 | 1.02E-09 |
| Orf8 | LIFR | 0.695 | 1.05E-09 |
| Nsp2 | STAM2 | 0.694 | 1.09E-09 |
| Nsp4 | MIOS | 0.694 | 1.10E-09 |
| Nsp12 | BIRC2 | 0.694 | 1.08E-09 |
| Nsp3 | IL7 | 0.693 | 1.14E-09 |
| Nsp3 | RICTOR | 0.693 | 1.20E-09 |
| Nsp5 | LIFR | 0.693 | 1.20E-09 |
| Nsp8 | LEPR | 0.693 | 1.18E-09 |
| Nsp14 | IFNAR1 | 0.693 | 1.23E-09 |
| S | PPP3CB | 0.693 | 1.19E-09 |
| S | CYLD | 0.693 | 1.22E-09 |
| Nsp2 | BIRC2 | 0.692 | 1.25E-09 |
| Nsp3 | JAK2 | 0.692 | 1.32E-09 |
| Nsp4 | IFNGR1 | 0.692 | 1.27E-09 |
| Nsp6 | PPP3CB | 0.692 | 1.29E-09 |
| Nsp6 | KRAS | 0.692 | 1.29E-09 |
| Nsp15 | ITCH | 0.692 | 1.27E-09 |
| Nsp16 | IL6ST | 0.692 | 1.26E-09 |

| | | | |
|---|---|---|---|
| Nsp16 | FZD6 | 0.692 | 1.25E-09 |
| Nsp6 | EIF2S1 | 0.691 | 1.43E-09 |
| Nsp16 | TAB3 | 0.691 | 1.35E-09 |
| Nsp16 | APAF1 | 0.691 | 1.39E-09 |
| Nsp4 | HIF1A | 0.69 | 1.51E-09 |
| Nsp6 | LEPR | 0.69 | 1.51E-09 |
| Nsp8 | PIAS2 | 0.69 | 1.45E-09 |
| Nsp13 | STAM | 0.69 | 1.48E-09 |
| Nsp15 | ROCK1 | 0.69 | 1.49E-09 |
| Nsp16 | MDM2 | 0.69 | 1.51E-09 |
| Nsp16 | PRKACB | 0.69 | 1.54E-09 |
| Nsp6 | USP25 | 0.689 | 1.59E-09 |
| Nsp6 | ZFYVE16 | 0.689 | 1.62E-09 |
| Nsp8 | PPM1B | 0.689 | 1.57E-09 |
| Nsp14 | MAP4K3 | 0.689 | 1.66E-09 |
| S | IL23R | 0.689 | 1.64E-09 |
| Orf3a | IL18 | 0.689 | 1.56E-09 |
| Nsp4 | DNM1L | 0.688 | 1.77E-09 |
| Nsp4 | CUL2 | 0.688 | 1.79E-09 |
| Nsp5 | PKN2 | 0.688 | 1.69E-09 |
| Nsp6 | STAM | 0.688 | 1.70E-09 |
| Nsp13 | IFNGR1 | 0.688 | 1.78E-09 |
| Nsp14 | LIFR | 0.688 | 1.70E-09 |
| Nsp15 | RPS6KA3 | 0.688 | 1.77E-09 |
| Nsp15 | CUL2 | 0.688 | 1.74E-09 |
| Orf3a | RPS6KA6 | 0.688 | 1.72E-09 |
| Nsp3 | RAP1B | 0.687 | 1.85E-09 |
| Nsp3 | AHR | 0.687 | 1.86E-09 |
| Nsp3 | PTPN13 | 0.687 | 1.85E-09 |
| Nsp4 | APAF1 | 0.687 | 1.83E-09 |
| Nsp6 | RPS6KA3 | 0.687 | 1.86E-09 |
| Nsp12 | SOS2 | 0.687 | 1.88E-09 |
| Nsp16 | HIF1A | 0.687 | 1.94E-09 |
| Nsp16 | PIAS2 | 0.687 | 1.94E-09 |
| S | AHR | 0.687 | 1.91E-09 |
| S | FNIP1 | 0.687 | 1.91E-09 |
| Nsp4 | XIAP | 0.686 | 2.07E-09 |
| Nsp6 | PLA2G4A | 0.686 | 1.96E-09 |
| Nsp8 | USP25 | 0.686 | 2.10E-09 |
| Nsp8 | ITCH | 0.686 | 1.95E-09 |
| Nsp14 | HIF1A | 0.686 | 1.98E-09 |
| S | PPP2R3C | 0.686 | 1.99E-09 |
| S | ITGAV | 0.686 | 2.03E-09 |
| S | FAS | 0.686 | 1.99E-09 |
| Nsp15 | SOS1 | 0.685 | 2.18E-09 |
| Nsp16 | ACVR2A | 0.685 | 2.17E-09 |
| Nsp16 | DNM1L | 0.685 | 2.24E-09 |
| Orf7a | ITCH | 0.685 | 2.22E-09 |
| Nsp12 | BRCA1 | 0.684 | 2.36E-09 |
| Nsp13 | BRCA1 | 0.684 | 2.37E-09 |

| | | | |
|---|---|---|---|
| Orf7a | CD36 | 0.684 | 2.33E-09 |
| Nsp2 | IRAK4 | 0.683 | 2.57E-09 |
| Orf3a | RPS6KA3 | 0.683 | 2.49E-09 |
| Nsp12 | STAM | 0.682 | 2.76E-09 |
| Nsp14 | SOS1 | 0.682 | 2.63E-09 |
| Nsp16 | IFNGR1 | 0.682 | 2.69E-09 |
| S | PRKAA1 | 0.682 | 2.79E-09 |
| S | IL7 | 0.682 | 2.81E-09 |
| Nsp4 | USP25 | 0.681 | 3.01E-09 |
| Nsp5 | ACVR2A | 0.681 | 3.00E-09 |
| Nsp12 | IL13RA2 | 0.681 | 2.88E-09 |
| Nsp14 | STK3 | 0.681 | 2.91E-09 |
| S | TFRC | 0.681 | 2.91E-09 |
| Nsp4 | ROCK2 | 0.68 | 3.16E-09 |
| Nsp8 | BRCA1 | 0.68 | 3.24E-09 |
| Nsp14 | IFNGR1 | 0.68 | 3.14E-09 |
| Orf8 | MIOS | 0.68 | 3.14E-09 |
| Nsp4 | IL13RA2 | 0.679 | 3.45E-09 |
| Nsp6 | PRKACB | 0.679 | 3.35E-09 |
| Nsp8 | RPS6KA6 | 0.679 | 3.40E-09 |
| Nsp8 | SOS1 | 0.679 | 3.42E-09 |
| Nsp12 | IL7 | 0.679 | 3.42E-09 |
| Nsp14 | TLR6 | 0.679 | 3.32E-09 |
| Nsp14 | STAM | 0.679 | 3.50E-09 |
| Nsp2 | COL4A5 | 0.678 | 3.60E-09 |
| Nsp3 | PTPN2 | 0.678 | 3.56E-09 |
| Nsp6 | MAP4K3 | 0.678 | 3.63E-09 |
| Nsp6 | PIAS2 | 0.678 | 3.77E-09 |
| Nsp8 | IL6ST | 0.678 | 3.79E-09 |
| Nsp16 | TAB2 | 0.678 | 3.72E-09 |
| Nsp16 | PPP2R3C | 0.678 | 3.73E-09 |
| S | ITGB1 | 0.678 | 3.67E-09 |
| S | AKT3 | 0.678 | 3.54E-09 |
| Nsp13 | RICTOR | 0.677 | 4.03E-09 |
| Nsp13 | PLA2G4A | 0.677 | 3.87E-09 |
| Nsp15 | XIAP | 0.677 | 4.01E-09 |
| Nsp3 | TLR6 | 0.676 | 4.20E-09 |
| Nsp4 | COL9A1 | 0.676 | 4.28E-09 |
| Nsp12 | PRKACB | 0.676 | 4.32E-09 |
| Nsp14 | XIAP | 0.676 | 4.22E-09 |
| Nsp3 | IL18 | 0.675 | 4.47E-09 |
| Nsp3 | RBL1 | 0.675 | 4.39E-09 |
| Nsp4 | SOS1 | 0.675 | 4.45E-09 |
| Nsp5 | STAM2 | 0.675 | 4.38E-09 |
| Nsp12 | IL6ST | 0.675 | 4.55E-09 |
| Nsp13 | MDM2 | 0.675 | 4.40E-09 |
| Nsp16 | SOS2 | 0.675 | 4.59E-09 |
| Nsp16 | CYLD | 0.675 | 4.48E-09 |
| Orf7a | IRAK4 | 0.675 | 4.60E-09 |
| Orf7a | PPP1R3A | 0.675 | 4.48E-09 |

| | | | |
|---|---|---|---|
| Nsp5 | RPS6KA3 | 0.674 | 4.83E-09 |
| Nsp6 | IL33 | 0.674 | 4.91E-09 |
| Nsp8 | PPP3CB | 0.674 | 4.77E-09 |
| Nsp13 | TAB3 | 0.674 | 4.73E-09 |
| Nsp15 | STAM | 0.674 | 5.01E-09 |
| Nsp15 | ATF2 | 0.674 | 5.04E-09 |
| Nsp16 | PLA2G4A | 0.674 | 4.96E-09 |
| S | JAK2 | 0.674 | 4.72E-09 |
| S | TGFBR1 | 0.674 | 4.90E-09 |
| Orf3a | ATM | 0.674 | 4.94E-09 |
| Nsp2 | PPM1B | 0.673 | 5.07E-09 |
| Nsp3 | SOCS4 | 0.673 | 5.25E-09 |
| Nsp12 | HIF1A | 0.673 | 5.32E-09 |
| Nsp15 | STAM2 | 0.673 | 5.27E-09 |
| Orf3a | BIRC2 | 0.673 | 5.40E-09 |
| Orf3a | IL6ST | 0.673 | 5.16E-09 |
| Nsp2 | CUL2 | 0.672 | 5.72E-09 |
| Nsp15 | MAP3K7 | 0.672 | 5.75E-09 |
| Nsp16 | RICTOR | 0.672 | 5.48E-09 |
| Orf7a | BIRC3 | 0.672 | 5.61E-09 |
| Nsp3 | CYLD | 0.671 | 5.85E-09 |
| Nsp9 | SOS1 | 0.671 | 6.08E-09 |
| Nsp12 | CUL2 | 0.671 | 5.97E-09 |
| Nsp12 | IFNGR1 | 0.671 | 5.94E-09 |
| Nsp8 | ROCK2 | 0.67 | 6.37E-09 |
| S | ATP6V1A | 0.67 | 6.54E-09 |
| Orf7a | COL4A4 | 0.67 | 6.42E-09 |
| Nsp5 | APAF1 | 0.669 | 6.87E-09 |
| Orf7a | CACNA2D1 | 0.669 | 7.21E-09 |
| Orf7a | CYLD | 0.669 | 7.19E-09 |
| Nsp2 | LIFR | 0.668 | 7.51E-09 |
| Nsp5 | DNM1L | 0.668 | 7.49E-09 |
| Nsp5 | LEPR | 0.668 | 7.39E-09 |
| Nsp5 | SOS1 | 0.668 | 7.32E-09 |
| Nsp8 | PLA2G4A | 0.668 | 7.29E-09 |
| Nsp14 | CAB39 | 0.668 | 7.58E-09 |
| Nsp14 | IL18 | 0.668 | 7.38E-09 |
| Nsp16 | HSP90B1 | 0.668 | 7.48E-09 |
| S | PPP2R3A | 0.668 | 7.32E-09 |
| Orf3a | IL13RA2 | 0.668 | 7.57E-09 |
| Nsp3 | EIF2S1 | 0.667 | 8.02E-09 |
| Nsp8 | ZFYVE16 | 0.667 | 8.21E-09 |
| Nsp15 | RICTOR | 0.667 | 8.29E-09 |
| Nsp16 | NFATC3 | 0.667 | 7.89E-09 |
| Orf7a | RAP1B | 0.667 | 7.93E-09 |
| Nsp8 | STAM2 | 0.666 | 8.47E-09 |
| Nsp12 | USP25 | 0.666 | 8.62E-09 |
| Nsp14 | LEPR | 0.666 | 8.87E-09 |
| Nsp15 | BIRC3 | 0.666 | 8.53E-09 |
| Nsp15 | BRCA1 | 0.666 | 8.58E-09 |

| | | | |
|---|---|---|---|
| S | GNB4 | 0.666 | 8.46E-09 |
| S | PIK3CA | 0.666 | 8.56E-09 |
| Nsp3 | TAB2 | 0.665 | 9.23E-09 |
| Nsp3 | PIAS2 | 0.665 | 9.01E-09 |
| Nsp4 | PLA2G4A | 0.665 | 9.22E-09 |
| Nsp12 | ROCK2 | 0.665 | 9.21E-09 |
| S | MSTN | 0.665 | 9.48E-09 |
| Nsp4 | EIF2S1 | 0.664 | 9.68E-09 |
| Nsp12 | AHR | 0.664 | 9.69E-09 |
| S | RBL1 | 0.664 | 9.80E-09 |
| Nsp4 | ACVR2A | 0.663 | 1.08E-08 |
| Nsp4 | PRKACB | 0.663 | 1.07E-08 |
| Nsp8 | TBK1 | 0.663 | 1.06E-08 |
| Nsp8 | ATM | 0.663 | 1.03E-08 |
| Nsp15 | GNB4 | 0.663 | 1.04E-08 |
| Nsp16 | EIF2S1 | 0.663 | 1.08E-08 |
| Orf7a | ATM | 0.663 | 1.07E-08 |
| Nsp2 | DNM1L | 0.662 | 1.11E-08 |
| Nsp12 | ATF2 | 0.662 | 1.17E-08 |
| Nsp16 | MAPK6 | 0.662 | 1.17E-08 |
| S | IL18 | 0.662 | 1.17E-08 |
| S | PPM1A | 0.662 | 1.12E-08 |
| Nsp6 | IRAK4 | 0.661 | 1.19E-08 |
| Nsp6 | ITCH | 0.661 | 1.20E-08 |
| Nsp8 | ROCK1 | 0.661 | 1.24E-08 |
| Nsp8 | EIF2S1 | 0.661 | 1.21E-08 |
| Nsp12 | APAF1 | 0.661 | 1.26E-08 |
| Nsp13 | APAF1 | 0.661 | 1.18E-08 |
| Nsp14 | PIAS2 | 0.661 | 1.21E-08 |
| Orf7a | FZD6 | 0.661 | 1.22E-08 |
| Nsp3 | PRKAA2 | 0.66 | 1.27E-08 |
| Nsp12 | MDM2 | 0.66 | 1.31E-08 |
| Nsp15 | BIRC2 | 0.66 | 1.30E-08 |
| Nsp15 | AHR | 0.66 | 1.34E-08 |
| Nsp16 | CD36 | 0.66 | 1.29E-08 |
| Orf7a | SGK3 | 0.66 | 1.27E-08 |
| Nsp4 | IFNAR1 | 0.659 | 1.42E-08 |
| Nsp12 | RPS6KA6 | 0.659 | 1.41E-08 |
| S | SMURF2 | 0.659 | 1.38E-08 |
| Orf3a | PPM1B | 0.659 | 1.36E-08 |
| Nsp4 | TLR6 | 0.658 | 1.47E-08 |
| Nsp6 | IL6ST | 0.658 | 1.54E-08 |
| Nsp8 | SGK3 | 0.658 | 1.44E-08 |
| Nsp13 | AKT3 | 0.658 | 1.52E-08 |
| Nsp14 | KRAS | 0.658 | 1.44E-08 |
| Orf3a | IRAK4 | 0.658 | 1.51E-08 |
| Orf3a | STAM2 | 0.658 | 1.46E-08 |
| Nsp3 | HSP90B1 | 0.657 | 1.55E-08 |
| Nsp4 | CD36 | 0.657 | 1.55E-08 |
| Nsp8 | LIFR | 0.657 | 1.55E-08 |

| | | | |
|---|---|---|---|
| Nsp13 | PTPN13 | 0.657 | 1.62E-08 |
| Nsp13 | FZD6 | 0.657 | 1.62E-08 |
| Nsp15 | MAP4K3 | 0.657 | 1.60E-08 |
| Nsp15 | DNM1L | 0.657 | 1.62E-08 |
| Nsp16 | SOCS4 | 0.657 | 1.62E-08 |
| Orf3a | COL1A2 | 0.657 | 1.63E-08 |
| Nsp3 | NFATC3 | 0.656 | 1.72E-08 |
| Nsp4 | PPM1B | 0.656 | 1.72E-08 |
| Nsp12 | PPM1B | 0.656 | 1.68E-08 |
| Nsp14 | CCNE2 | 0.656 | 1.74E-08 |
| Nsp14 | STAM2 | 0.656 | 1.68E-08 |
| Nsp14 | SGK3 | 0.656 | 1.66E-08 |
| Nsp15 | RPS6KA6 | 0.656 | 1.71E-08 |
| Nsp16 | PTPN13 | 0.656 | 1.69E-08 |
| S | IFIH1 | 0.656 | 1.67E-08 |
| Orf7a | MAP3K1 | 0.656 | 1.74E-08 |
| Nsp4 | IL6ST | 0.655 | 1.81E-08 |
| Nsp5 | MDM2 | 0.655 | 1.84E-08 |
| Nsp5 | CD36 | 0.655 | 1.76E-08 |
| Nsp8 | IFNAR1 | 0.655 | 1.82E-08 |
| Nsp13 | MSTN | 0.655 | 1.77E-08 |
| Nsp14 | SOS2 | 0.655 | 1.77E-08 |
| Nsp16 | ZFYVE9 | 0.655 | 1.77E-08 |
| Nsp3 | PRKAA1 | 0.654 | 1.89E-08 |
| Nsp4 | RPS6KA6 | 0.654 | 1.89E-08 |
| Nsp12 | ITCH | 0.654 | 1.97E-08 |
| Nsp12 | SOS1 | 0.654 | 1.89E-08 |
| Nsp15 | IL7 | 0.654 | 1.99E-08 |
| Nsp6 | STAM2 | 0.653 | 2.05E-08 |
| Nsp8 | CD36 | 0.653 | 2.11E-08 |
| Nsp13 | CAMK2D | 0.653 | 2.13E-08 |
| Nsp13 | FNIP1 | 0.653 | 2.10E-08 |
| Nsp13 | PPP2CB | 0.653 | 2.08E-08 |
| Nsp15 | RASA1 | 0.653 | 2.09E-08 |
| Nsp16 | JAK2 | 0.653 | 2.08E-08 |
| S | SOCS4 | 0.653 | 2.03E-08 |
| S | ITGA2 | 0.653 | 2.10E-08 |
| Orf7a | PKN2 | 0.653 | 2.14E-08 |
| Nsp3 | FNIP1 | 0.652 | 2.29E-08 |
| Nsp8 | MAP4K3 | 0.652 | 2.18E-08 |
| Nsp8 | SOS2 | 0.652 | 2.18E-08 |
| Nsp14 | PRKACB | 0.652 | 2.18E-08 |
| Nsp14 | RASA2 | 0.652 | 2.28E-08 |
| Nsp16 | TLR6 | 0.652 | 2.26E-08 |
| Nsp3 | STK3 | 0.651 | 2.31E-08 |
| Nsp5 | CUL2 | 0.651 | 2.40E-08 |
| Nsp13 | RAP1B | 0.651 | 2.41E-08 |
| Nsp15 | JAK2 | 0.651 | 2.37E-08 |
| Nsp15 | RBL1 | 0.651 | 2.35E-08 |
| Orf7a | RPS6KA3 | 0.651 | 2.43E-08 |

| | | | |
|---|---|---|---|
| Nsp2 | LDHC | 0.65 | 2.47E-08 |
| Nsp4 | CCNE2 | 0.65 | 2.55E-08 |
| Nsp6 | TLR6 | 0.65 | 2.52E-08 |
| Nsp6 | DNM1L | 0.65 | 2.52E-08 |
| Nsp13 | SGK3 | 0.65 | 2.56E-08 |
| S | STAT4 | 0.65 | 2.46E-08 |
| S | MAP3K7 | 0.65 | 2.50E-08 |
| Orf7a | FNIP1 | 0.65 | 2.56E-08 |
| Nsp6 | PKN2 | 0.649 | 2.69E-08 |
| Nsp14 | CD36 | 0.649 | 2.74E-08 |
| Nsp15 | SOCS4 | 0.649 | 2.72E-08 |
| Nsp15 | PRKACB | 0.649 | 2.66E-08 |
| S | RBL2 | 0.649 | 2.76E-08 |
| Nsp5 | TBK1 | 0.648 | 2.86E-08 |
| Nsp8 | XIAP | 0.648 | 2.94E-08 |
| Nsp12 | MSTN | 0.648 | 2.87E-08 |
| Nsp15 | TAB3 | 0.648 | 2.96E-08 |
| Nsp16 | AHR | 0.648 | 2.82E-08 |
| S | PIK3CB | 0.648 | 2.86E-08 |
| Nsp6 | SOCS4 | 0.647 | 3.06E-08 |
| Nsp13 | PRKAA1 | 0.647 | 3.04E-08 |
| Nsp13 | IL7 | 0.647 | 3.06E-08 |
| Nsp14 | BIRC2 | 0.647 | 3.06E-08 |
| Nsp16 | PPM1A | 0.647 | 3.07E-08 |
| S | CCNE2 | 0.647 | 3.06E-08 |
| S | CDC42 | 0.647 | 3.05E-08 |
| Orf3a | PPP1R3A | 0.647 | 3.02E-08 |
| Orf3a | BIRC3 | 0.647 | 3.02E-08 |
| Orf7a | LIFR | 0.647 | 3.01E-08 |
| Orf7a | PPP2R5E | 0.647 | 3.15E-08 |
| Orf8 | ATM | 0.647 | 3.08E-08 |
| Nsp4 | MDM2 | 0.646 | 3.20E-08 |
| Nsp4 | PPM1A | 0.646 | 3.28E-08 |
| Nsp12 | CD36 | 0.646 | 3.21E-08 |
| Nsp15 | IFNGR1 | 0.646 | 3.29E-08 |
| Nsp3 | PPM1A | 0.645 | 3.43E-08 |
| Nsp12 | PPP3CB | 0.645 | 3.60E-08 |
| Nsp13 | PTPN2 | 0.645 | 3.47E-08 |
| Nsp15 | USP25 | 0.645 | 3.48E-08 |
| Orf7a | ITGB8 | 0.645 | 3.53E-08 |
| Orf8 | GNB4 | 0.645 | 3.62E-08 |
| Nsp2 | PPP2CA | 0.644 | 3.67E-08 |
| Nsp3 | ITGA2 | 0.644 | 3.81E-08 |
| Nsp14 | PTPN2 | 0.644 | 3.79E-08 |
| Nsp3 | CACNA2D1 | 0.643 | 3.99E-08 |
| Nsp4 | AHR | 0.643 | 3.90E-08 |
| Nsp13 | MAPK9 | 0.643 | 4.00E-08 |
| Nsp14 | COL4A5 | 0.643 | 3.91E-08 |
| S | ITGA1 | 0.643 | 3.95E-08 |
| Nsp6 | MAP3K1 | 0.642 | 4.14E-08 |

| | | | |
|---|---|---|---|
| Nsp13 | JAK2 | 0.642 | 4.16E-08 |
| Nsp2 | ATF2 | 0.641 | 4.43E-08 |
| Nsp3 | MAP3K7 | 0.641 | 4.62E-08 |
| Nsp4 | IL18 | 0.641 | 4.54E-08 |
| Nsp4 | SGK3 | 0.641 | 4.57E-08 |
| Nsp8 | STAM | 0.641 | 4.49E-08 |
| Nsp13 | IFIH1 | 0.641 | 4.49E-08 |
| S | COL1A2 | 0.641 | 4.61E-08 |
| S | TAOK1 | 0.641 | 4.47E-08 |
| Nsp8 | IL13RA2 | 0.64 | 4.86E-08 |
| Nsp8 | APAF1 | 0.64 | 4.87E-08 |
| Orf3a | XIAP | 0.64 | 4.88E-08 |
| Nsp6 | HIF1A | 0.639 | 5.06E-08 |
| Nsp9 | XIAP | 0.639 | 5.17E-08 |
| Nsp12 | JAK2 | 0.639 | 5.28E-08 |
| Nsp13 | PPP3CB | 0.639 | 5.29E-08 |
| Nsp16 | PPP2R3A | 0.639 | 5.01E-08 |
| Orf7a | COL4A3 | 0.639 | 5.26E-08 |
| Orf8 | ATF2 | 0.639 | 5.04E-08 |
| Nsp2 | RPS6KA6 | 0.638 | 5.55E-08 |
| Nsp3 | MSTN | 0.638 | 5.44E-08 |
| Nsp5 | TAB2 | 0.638 | 5.54E-08 |
| Nsp5 | BRCA1 | 0.638 | 5.54E-08 |
| Nsp6 | IFNAR1 | 0.638 | 5.55E-08 |
| Nsp8 | IL7 | 0.638 | 5.35E-08 |
| Nsp8 | ATF2 | 0.638 | 5.55E-08 |
| Nsp12 | RBL1 | 0.638 | 5.35E-08 |
| Nsp13 | AHR | 0.638 | 5.56E-08 |
| Orf3a | ROCK1 | 0.638 | 5.44E-08 |
| Nsp2 | MAPK6 | 0.637 | 5.85E-08 |
| Nsp2 | IL13RA2 | 0.637 | 5.78E-08 |
| Nsp4 | PTPN2 | 0.637 | 5.86E-08 |
| Nsp4 | HSP90B1 | 0.637 | 5.88E-08 |
| Nsp8 | HIF1A | 0.637 | 5.69E-08 |
| Nsp16 | CACNA2D1 | 0.637 | 5.69E-08 |
| Nsp16 | ATF2 | 0.637 | 5.89E-08 |
| S | SOCS5 | 0.637 | 5.91E-08 |
| Orf7a | ROCK2 | 0.637 | 5.68E-08 |
| Orf7a | IL1R1 | 0.637 | 5.71E-08 |
| Nsp5 | SOS2 | 0.636 | 6.31E-08 |
| Nsp6 | TBK1 | 0.636 | 6.12E-08 |
| Nsp6 | ITGA2 | 0.636 | 6.24E-08 |
| Nsp9 | PIAS2 | 0.636 | 6.24E-08 |
| Nsp16 | FAS | 0.636 | 6.09E-08 |
| Nsp16 | FNIP1 | 0.636 | 6.39E-08 |
| Nsp3 | RASA2 | 0.635 | 6.40E-08 |
| Nsp3 | COL9A1 | 0.635 | 6.55E-08 |
| Nsp8 | RICTOR | 0.635 | 6.42E-08 |
| Nsp14 | PTPN13 | 0.635 | 6.73E-08 |
| Nsp14 | DNM1L | 0.635 | 6.65E-08 |

| | | | |
|---|---|---|---|
| Nsp15 | PIAS2 | 0.635 | 6.61E-08 |
| Orf7a | IL23R | 0.635 | 6.79E-08 |
| Nsp4 | GNB4 | 0.634 | 7.14E-08 |
| Nsp8 | LDHC | 0.634 | 7.08E-08 |
| Nsp8 | ACVR2A | 0.634 | 6.94E-08 |
| Nsp8 | DNM1L | 0.634 | 7.17E-08 |
| Nsp13 | COL4A3 | 0.634 | 6.96E-08 |
| Nsp14 | ATF2 | 0.634 | 6.84E-08 |
| Nsp14 | RICTOR | 0.634 | 7.10E-08 |
| Nsp15 | TAB2 | 0.634 | 7.15E-08 |
| Nsp16 | IL7 | 0.634 | 7.09E-08 |
| Orf7a | CAB39 | 0.634 | 6.94E-08 |
| S | IL18R1 | 0.633 | 7.51E-08 |
| Orf7a | IL18 | 0.633 | 7.54E-08 |
| Nsp2 | TAB2 | 0.632 | 8.19E-08 |
| Nsp4 | RICTOR | 0.632 | 7.81E-08 |
| Nsp12 | SGK3 | 0.632 | 7.98E-08 |
| Nsp13 | CACNA2D1 | 0.632 | 7.85E-08 |
| Nsp13 | TFRC | 0.632 | 8.19E-08 |
| Nsp14 | EIF2S1 | 0.632 | 7.85E-08 |
| Nsp14 | TAOK1 | 0.632 | 8.00E-08 |
| Nsp16 | ITGA4 | 0.632 | 7.75E-08 |
| N | FZD6 | 0.632 | 7.96E-08 |
| Orf7a | ROCK1 | 0.632 | 8.07E-08 |
| Nsp3 | FAS | 0.631 | 8.53E-08 |
| Nsp4 | JAK2 | 0.631 | 8.50E-08 |
| Nsp5 | XIAP | 0.631 | 8.52E-08 |
| Nsp9 | SOS2 | 0.631 | 8.25E-08 |
| Nsp13 | GHR | 0.631 | 8.55E-08 |
| Orf7a | SOS1 | 0.631 | 8.65E-08 |
| Orf8 | IRAK4 | 0.631 | 8.48E-08 |
| Nsp2 | ACSL3 | 0.63 | 9.01E-08 |
| Nsp3 | CDC42 | 0.63 | 8.89E-08 |
| Nsp6 | PARP4 | 0.63 | 8.75E-08 |
| Nsp13 | IL18 | 0.63 | 9.20E-08 |
| Nsp15 | SGK3 | 0.63 | 9.22E-08 |
| Nsp16 | STK3 | 0.63 | 9.23E-08 |
| Nsp16 | COL4A5 | 0.63 | 8.73E-08 |
| Orf7a | TBK1 | 0.63 | 8.78E-08 |
| Nsp2 | COL1A2 | 0.629 | 9.37E-08 |
| Nsp3 | ITGA4 | 0.629 | 9.69E-08 |
| Nsp4 | IL7 | 0.629 | 9.42E-08 |
| Nsp8 | JAK2 | 0.629 | 9.76E-08 |
| S | PRKAA2 | 0.629 | 9.82E-08 |
| Nsp4 | ZFYVE9 | 0.628 | 9.93E-08 |
| Nsp5 | ATM | 0.628 | 1.03E-07 |
| Nsp6 | ATP6V1A | 0.628 | 9.88E-08 |
| Nsp6 | CDC42 | 0.628 | 1.04E-07 |
| Nsp12 | CCNE2 | 0.628 | 1.02E-07 |
| Nsp16 | ITGB1 | 0.628 | 1.02E-07 |

| | | | |
|---|---|---|---|
| S | ITGA4 | 0.628 | 1.01E-07 |
| Orf3a | LEPR | 0.628 | 1.04E-07 |
| Orf7a | PTPN13 | 0.628 | 9.98E-08 |
| Nsp3 | PIK3CA | 0.627 | 1.09E-07 |
| Nsp9 | JAK2 | 0.627 | 1.05E-07 |
| Nsp9 | SMAD4 | 0.627 | 1.09E-07 |
| Nsp12 | TLR6 | 0.627 | 1.07E-07 |
| Nsp13 | RASA1 | 0.627 | 1.11E-07 |
| S | FZD3 | 0.627 | 1.09E-07 |
| S | CACNA2D1 | 0.627 | 1.07E-07 |
| Nsp2 | STAM | 0.626 | 1.14E-07 |
| Nsp2 | MAP3K7 | 0.626 | 1.18E-07 |
| Nsp3 | PPP2R3C | 0.626 | 1.16E-07 |
| Nsp4 | PIAS2 | 0.626 | 1.13E-07 |
| Nsp6 | RBL2 | 0.626 | 1.15E-07 |
| Nsp6 | CD36 | 0.626 | 1.17E-07 |
| Nsp9 | ATM | 0.626 | 1.16E-07 |
| Nsp14 | USP25 | 0.626 | 1.13E-07 |
| Nsp14 | TAB3 | 0.626 | 1.13E-07 |
| S | BMPR2 | 0.626 | 1.15E-07 |
| Orf7a | STAM2 | 0.626 | 1.18E-07 |
| Nsp5 | ROCK1 | 0.625 | 1.24E-07 |
| Nsp13 | HSP90B1 | 0.625 | 1.19E-07 |
| Nsp13 | CYLD | 0.625 | 1.24E-07 |
| Nsp14 | TFRC | 0.625 | 1.24E-07 |
| Nsp15 | PRKAA2 | 0.625 | 1.20E-07 |
| Nsp16 | MSTN | 0.625 | 1.19E-07 |
| S | MAP3K1 | 0.625 | 1.24E-07 |
| Orf8 | ROCK2 | 0.625 | 1.22E-07 |
| Nsp4 | MAPK6 | 0.624 | 1.31E-07 |
| Nsp13 | ACVR2A | 0.624 | 1.28E-07 |
| Nsp15 | ROCK2 | 0.624 | 1.29E-07 |
| Orf3a | ITGAV | 0.624 | 1.25E-07 |
| Orf8 | RPS6KA3 | 0.624 | 1.27E-07 |
| Nsp2 | MDM2 | 0.623 | 1.36E-07 |
| Nsp6 | SEH1L | 0.623 | 1.33E-07 |
| Nsp6 | CACNA2D1 | 0.623 | 1.33E-07 |
| Nsp12 | ACVR2A | 0.623 | 1.40E-07 |
| Nsp13 | MAPK6 | 0.623 | 1.38E-07 |
| Nsp13 | ITGA1 | 0.623 | 1.34E-07 |
| Nsp14 | ATP6V1A | 0.623 | 1.40E-07 |
| Nsp16 | SOCS5 | 0.623 | 1.35E-07 |
| Orf8 | CUL2 | 0.623 | 1.34E-07 |
| Nsp3 | STAT4 | 0.622 | 1.45E-07 |
| Nsp6 | PTPN13 | 0.622 | 1.41E-07 |
| Nsp6 | SOS2 | 0.622 | 1.43E-07 |
| Nsp6 | ATF2 | 0.622 | 1.46E-07 |
| Nsp8 | MAPK6 | 0.622 | 1.47E-07 |
| Nsp14 | ZFYVE9 | 0.622 | 1.44E-07 |
| Nsp15 | APAF1 | 0.622 | 1.43E-07 |

| | | | |
|---|---|---|---|
| Nsp16 | RBL2 | 0.622 | 1.47E-07 |
| S | TLR8 | 0.622 | 1.48E-07 |
| S | COL4A3 | 0.622 | 1.48E-07 |
| Orf7a | CD86 | 0.622 | 1.42E-07 |
| Nsp9 | IRAK4 | 0.621 | 1.50E-07 |
| S | LDHC | 0.621 | 1.54E-07 |
| Orf8 | STAM2 | 0.621 | 1.52E-07 |
| Orf8 | SOS2 | 0.621 | 1.51E-07 |
| Nsp5 | NFATC3 | 0.62 | 1.65E-07 |
| Nsp13 | TAOK1 | 0.62 | 1.66E-07 |
| Nsp16 | IL23R | 0.62 | 1.59E-07 |
| Nsp16 | COL9A1 | 0.62 | 1.62E-07 |
| Orf7a | TGFBR1 | 0.62 | 1.63E-07 |
| Orf8 | PPM1B | 0.62 | 1.66E-07 |
| Orf8 | BIRC3 | 0.62 | 1.64E-07 |
| Nsp4 | FZD6 | 0.619 | 1.70E-07 |
| Nsp5 | MAPK6 | 0.619 | 1.73E-07 |
| Nsp9 | CCNE2 | 0.619 | 1.78E-07 |
| Nsp12 | RICTOR | 0.619 | 1.72E-07 |
| Nsp15 | IFNAR1 | 0.619 | 1.70E-07 |
| Nsp15 | FZD6 | 0.619 | 1.76E-07 |
| Nsp3 | ITGAV | 0.618 | 1.81E-07 |
| Nsp3 | AKT3 | 0.618 | 1.87E-07 |
| Nsp6 | AHR | 0.618 | 1.89E-07 |
| Nsp14 | IL6ST | 0.618 | 1.84E-07 |
| Nsp16 | ITGA1 | 0.618 | 1.86E-07 |
| S | PARP4 | 0.618 | 1.88E-07 |
| Orf3a | STAM | 0.618 | 1.83E-07 |
| Orf3a | MAPK6 | 0.618 | 1.83E-07 |
| Orf7a | COL4A6 | 0.618 | 1.86E-07 |
| Nsp3 | ITGB1 | 0.617 | 1.94E-07 |
| Nsp3 | PIK3CB | 0.617 | 1.94E-07 |
| Nsp4 | COL1A2 | 0.617 | 1.94E-07 |
| Nsp14 | FZD6 | 0.617 | 1.92E-07 |
| Orf3a | TLR6 | 0.617 | 1.98E-07 |
| Orf7a | ZFYVE16 | 0.617 | 2.00E-07 |
| Orf8 | PIAS2 | 0.617 | 1.92E-07 |
| Nsp5 | IL23R | 0.616 | 2.10E-07 |
| Nsp6 | MSTN | 0.616 | 2.02E-07 |
| Nsp12 | PTPN13 | 0.616 | 2.11E-07 |
| Nsp13 | ITGB8 | 0.616 | 2.10E-07 |
| Nsp13 | PIK3CA | 0.616 | 2.08E-07 |
| Nsp16 | IL18 | 0.616 | 2.08E-07 |
| Nsp16 | TAOK1 | 0.616 | 2.12E-07 |
| S | DDX58 | 0.616 | 2.11E-07 |
| S | IL1R1 | 0.616 | 2.05E-07 |
| Orf7a | PPP2R3C | 0.616 | 2.04E-07 |
| Nsp3 | TFRC | 0.615 | 2.13E-07 |
| Nsp6 | CAB39 | 0.615 | 2.18E-07 |
| Nsp6 | XIAP | 0.615 | 2.22E-07 |

| | | | |
|---|---|---|---|
| Nsp2 | MIOS | 0.614 | 2.27E-07 |
| Nsp2 | PKN2 | 0.614 | 2.26E-07 |
| Nsp3 | RASA1 | 0.614 | 2.30E-07 |
| Nsp6 | MAPK6 | 0.614 | 2.29E-07 |
| Nsp9 | ATF2 | 0.614 | 2.27E-07 |
| Nsp13 | ITGA4 | 0.614 | 2.26E-07 |
| Nsp14 | MIOS | 0.614 | 2.27E-07 |
| Nsp14 | CD86 | 0.614 | 2.30E-07 |
| Nsp15 | ITGB1 | 0.614 | 2.29E-07 |
| S | GHR | 0.614 | 2.29E-07 |
| S | RASA2 | 0.614 | 2.27E-07 |
| Nsp3 | TGFBR1 | 0.613 | 2.48E-07 |
| Nsp6 | TAB2 | 0.613 | 2.40E-07 |
| Nsp6 | HSP90B1 | 0.613 | 2.43E-07 |
| Nsp14 | ITGA2 | 0.613 | 2.44E-07 |
| N | IL18 | 0.613 | 2.51E-07 |
| Orf3a | MAP4K3 | 0.613 | 2.48E-07 |
| Orf3a | COL9A1 | 0.613 | 2.43E-07 |
| Nsp2 | HSP90B1 | 0.612 | 2.60E-07 |
| Nsp5 | ITGB1 | 0.612 | 2.66E-07 |
| Nsp6 | RASA2 | 0.612 | 2.56E-07 |
| Nsp6 | ZFYVE9 | 0.612 | 2.66E-07 |
| Nsp9 | TBK1 | 0.612 | 2.66E-07 |
| Nsp13 | PPP2CA | 0.612 | 2.60E-07 |
| Nsp16 | RBL1 | 0.612 | 2.56E-07 |
| S | PTPN2 | 0.612 | 2.64E-07 |
| Orf3a | SGK3 | 0.612 | 2.57E-07 |
| Orf8 | IL1R1 | 0.612 | 2.56E-07 |
| Nsp4 | PTPN13 | 0.611 | 2.72E-07 |
| Nsp6 | PPM1A | 0.611 | 2.74E-07 |
| Nsp13 | TLR6 | 0.611 | 2.79E-07 |
| Nsp13 | PRKAA2 | 0.611 | 2.74E-07 |
| Nsp13 | AZI2 | 0.611 | 2.73E-07 |
| Nsp13 | PIAS2 | 0.611 | 2.82E-07 |
| Nsp14 | CYLD | 0.611 | 2.73E-07 |
| Nsp15 | ACVR2A | 0.611 | 2.79E-07 |
| Orf3a | CCNE2 | 0.611 | 2.74E-07 |
| Orf3a | PKN2 | 0.611 | 2.71E-07 |
| Nsp3 | SOCS5 | 0.61 | 2.99E-07 |
| Nsp3 | MAP3K1 | 0.61 | 2.91E-07 |
| Nsp6 | RPS6KA6 | 0.61 | 2.98E-07 |
| Nsp6 | IL18 | 0.61 | 2.93E-07 |
| Nsp8 | NFATC3 | 0.61 | 2.95E-07 |
| Nsp12 | IL18 | 0.61 | 2.85E-07 |
| Nsp12 | PLA2G4A | 0.61 | 2.84E-07 |
| Nsp14 | RPS6KA6 | 0.61 | 2.95E-07 |
| Nsp15 | TAOK1 | 0.61 | 2.84E-07 |
| Nsp16 | CDC42 | 0.61 | 2.97E-07 |
| Nsp16 | AKT3 | 0.61 | 2.98E-07 |
| Nsp16 | IL18R1 | 0.61 | 2.99E-07 |

| | | | |
|---|---|---|---|
| Orf3a | ZFYVE16 | 0.61 | 2.96E-07 |
| Orf7a | COL4A5 | 0.61 | 2.87E-07 |
| Orf8 | TGFBR1 | 0.61 | 2.86E-07 |
| Orf8 | ZFYVE16 | 0.61 | 2.93E-07 |
| Nsp3 | GNB4 | 0.609 | 3.08E-07 |
| Nsp3 | ATP6V1A | 0.609 | 3.09E-07 |
| Nsp4 | KRAS | 0.609 | 3.13E-07 |
| Nsp8 | SOCS4 | 0.609 | 3.05E-07 |
| Nsp8 | COL9A1 | 0.609 | 3.02E-07 |
| S | RASA1 | 0.609 | 3.14E-07 |
| Orf3a | TGFBR1 | 0.609 | 3.16E-07 |
| Orf7a | PLA2G4A | 0.609 | 3.04E-07 |
| Nsp3 | IFIH1 | 0.608 | 3.28E-07 |
| Nsp3 | TAOK1 | 0.608 | 3.26E-07 |
| Nsp6 | LIFR | 0.608 | 3.21E-07 |
| Nsp14 | RBL2 | 0.608 | 3.25E-07 |
| Nsp15 | IL6ST | 0.608 | 3.31E-07 |
| Nsp15 | PIK3CA | 0.608 | 3.33E-07 |
| Nsp1 | LDHAL6A | 0.607 | 3.53E-07 |
| Nsp3 | PPP2R3A | 0.607 | 3.44E-07 |
| Nsp3 | KRAS | 0.607 | 3.47E-07 |
| Nsp4 | PPP2R3C | 0.607 | 3.46E-07 |
| Nsp4 | TFRC | 0.607 | 3.38E-07 |
| Nsp4 | TAB3 | 0.607 | 3.40E-07 |
| Nsp5 | IL6ST | 0.607 | 3.44E-07 |
| Nsp5 | ATF2 | 0.607 | 3.51E-07 |
| Nsp9 | BIRC3 | 0.607 | 3.44E-07 |
| Nsp13 | PPM1A | 0.607 | 3.45E-07 |
| Nsp14 | COL9A1 | 0.607 | 3.55E-07 |
| Nsp15 | ACSL3 | 0.607 | 3.41E-07 |
| Nsp16 | PTPN2 | 0.607 | 3.55E-07 |
| Orf7a | IL6ST | 0.607 | 3.43E-07 |
| Orf7a | CDC42 | 0.607 | 3.48E-07 |
| Orf8 | RPS6KA6 | 0.607 | 3.40E-07 |
| Nsp3 | CCNE2 | 0.606 | 3.69E-07 |
| Nsp4 | MSTN | 0.606 | 3.72E-07 |
| Nsp5 | USP25 | 0.606 | 3.59E-07 |
| Nsp5 | PIAS2 | 0.606 | 3.67E-07 |
| Nsp5 | SGK3 | 0.606 | 3.71E-07 |
| Nsp6 | COL9A1 | 0.606 | 3.73E-07 |
| Nsp14 | IL13RA2 | 0.606 | 3.56E-07 |
| Nsp16 | SMURF2 | 0.606 | 3.65E-07 |
| Nsp16 | IL13RA2 | 0.606 | 3.63E-07 |
| Orf3a | USP25 | 0.606 | 3.68E-07 |
| Orf7a | APAF1 | 0.606 | 3.62E-07 |
| Nsp3 | LDHC | 0.605 | 3.91E-07 |
| Nsp5 | ZFYVE16 | 0.605 | 3.87E-07 |
| Nsp6 | ACVR2A | 0.605 | 3.93E-07 |
| Nsp16 | PIK3CB | 0.605 | 3.86E-07 |
| Orf3a | CAMK2D | 0.605 | 3.89E-07 |

| | | | |
|---|---|---|---|
| Orf7a | STAM | 0.605 | 3.96E-07 |
| Orf7a | PHKB | 0.605 | 3.96E-07 |
| Nsp5 | IL13RA2 | 0.604 | 4.13E-07 |
| Nsp6 | COL4A5 | 0.604 | 4.01E-07 |
| Nsp12 | PRKAA1 | 0.604 | 4.08E-07 |
| Nsp12 | TAB3 | 0.604 | 4.03E-07 |
| Nsp2 | IL7 | 0.603 | 4.39E-07 |
| Nsp2 | PPP2CB | 0.603 | 4.21E-07 |
| Nsp2 | APAF1 | 0.603 | 4.25E-07 |
| Nsp4 | SOCS4 | 0.603 | 4.44E-07 |
| Nsp5 | MAP4K3 | 0.603 | 4.22E-07 |
| Nsp6 | SOS1 | 0.603 | 4.43E-07 |
| Nsp8 | HSP90B1 | 0.603 | 4.35E-07 |
| Nsp8 | CYLD | 0.603 | 4.38E-07 |
| Nsp9 | PPP1R3A | 0.603 | 4.23E-07 |
| Nsp12 | ZFYVE9 | 0.603 | 4.43E-07 |
| Nsp14 | FAS | 0.603 | 4.28E-07 |
| Nsp14 | CACNA2D1 | 0.603 | 4.23E-07 |
| Nsp14 | APAF1 | 0.603 | 4.42E-07 |
| Nsp15 | IL13RA2 | 0.603 | 4.43E-07 |
| Nsp5 | COL4A5 | 0.602 | 4.45E-07 |
| Nsp15 | PTPN2 | 0.602 | 4.46E-07 |
| Nsp16 | PRKAA1 | 0.602 | 4.50E-07 |
| S | PPP2CA | 0.602 | 4.66E-07 |
| S | RAPGEF2 | 0.602 | 4.48E-07 |
| Nsp3 | MAPK9 | 0.601 | 4.95E-07 |
| Nsp3 | IL18R1 | 0.601 | 4.79E-07 |
| Nsp8 | STAT4 | 0.601 | 4.90E-07 |
| Nsp9 | BRCA1 | 0.601 | 4.72E-07 |
| Orf8 | LEPR | 0.601 | 4.90E-07 |
| Nsp4 | STAT4 | 0.6 | 5.02E-07 |
| Nsp6 | TAB3 | 0.6 | 5.01E-07 |
| Nsp8 | IFIH1 | 0.6 | 5.05E-07 |
| Nsp12 | PPP2R3C | 0.6 | 5.04E-07 |
| Nsp13 | ZFYVE9 | 0.6 | 5.18E-07 |
| Nsp14 | PARP4 | 0.6 | 5.17E-07 |
| Nsp16 | ITGA2 | 0.6 | 5.17E-07 |
| S | KRAS | 0.6 | 5.06E-07 |
| Orf7a | DDX58 | 0.6 | 4.97E-07 |
| Orf7a | XIAP | 0.6 | 5.22E-07 |
| Orf8 | SOS1 | 0.6 | 5.16E-07 |
| Nsp2 | ATM | 0.599 | 5.47E-07 |
| Nsp3 | CAMK2D | 0.599 | 5.49E-07 |
| Nsp3 | EIF4E | 0.599 | 5.49E-07 |
| Nsp3 | ITGA1 | 0.599 | 5.45E-07 |
| Nsp3 | COL4A3 | 0.599 | 5.27E-07 |
| Nsp4 | STK3 | 0.599 | 5.35E-07 |
| Nsp6 | PRKAA2 | 0.599 | 5.28E-07 |
| Nsp8 | BIRC3 | 0.599 | 5.48E-07 |
| Nsp14 | ITGA4 | 0.599 | 5.30E-07 |

| | | | |
|---|---|---|---|
| Nsp14 | AZI2 | 0.599 | 5.45E-07 |
| Nsp14 | IL18R1 | 0.599 | 5.25E-07 |
| Nsp15 | ZFYVE9 | 0.599 | 5.29E-07 |
| S | ACSL3 | 0.599 | 5.40E-07 |
| S | AZI2 | 0.599 | 5.26E-07 |
| Orf7a | AKT3 | 0.599 | 5.28E-07 |
| Orf7a | TAOK1 | 0.599 | 5.30E-07 |
| Nsp2 | ATP6V1A | 0.598 | 5.77E-07 |
| Nsp2 | TGFBR1 | 0.598 | 5.74E-07 |
| Nsp2 | PPP1R3A | 0.598 | 5.76E-07 |
| Nsp4 | TAB2 | 0.598 | 5.61E-07 |
| Nsp8 | PPM1A | 0.598 | 5.78E-07 |
| Nsp8 | CDC42 | 0.598 | 5.81E-07 |
| Nsp12 | TAB2 | 0.598 | 5.81E-07 |
| Nsp14 | AKT3 | 0.598 | 5.74E-07 |
| S | STK3 | 0.598 | 5.65E-07 |
| S | EGF | 0.598 | 5.67E-07 |
| Nsp3 | COL1A2 | 0.597 | 5.88E-07 |
| Nsp4 | CD86 | 0.597 | 6.00E-07 |
| Nsp8 | PRKAA1 | 0.597 | 6.08E-07 |
| Nsp13 | MAP3K7 | 0.597 | 5.94E-07 |
| Nsp16 | PPM1B | 0.597 | 6.04E-07 |
| Nsp16 | PIK3CA | 0.597 | 6.12E-07 |
| S | NLK | 0.597 | 5.97E-07 |
| Orf3a | STRADB | 0.597 | 5.97E-07 |
| Nsp4 | CYLD | 0.596 | 6.32E-07 |
| Nsp6 | MDM2 | 0.596 | 6.45E-07 |
| Nsp13 | DDX58 | 0.596 | 6.24E-07 |
| S | PHKB | 0.596 | 6.46E-07 |
| Orf3a | COL4A3 | 0.596 | 6.26E-07 |
| Orf8 | PPP2R3C | 0.596 | 6.29E-07 |
| Nsp3 | AZI2 | 0.595 | 6.66E-07 |
| Nsp3 | DDX58 | 0.595 | 6.73E-07 |
| Nsp4 | PRKAA2 | 0.595 | 6.55E-07 |
| Nsp4 | RASA2 | 0.595 | 6.75E-07 |
| Nsp6 | SGK3 | 0.595 | 6.81E-07 |
| Nsp8 | SMURF2 | 0.595 | 6.70E-07 |
| Nsp8 | FNIP1 | 0.595 | 6.72E-07 |
| Nsp13 | CCNE2 | 0.595 | 6.83E-07 |
| Nsp13 | NFATC3 | 0.595 | 6.85E-07 |
| Nsp14 | IL1R1 | 0.595 | 6.59E-07 |
| Nsp15 | ITGA1 | 0.595 | 6.72E-07 |
| Nsp16 | EIF4E | 0.595 | 6.80E-07 |
| S | CAMK2D | 0.595 | 6.63E-07 |
| Orf8 | PKN2 | 0.595 | 6.64E-07 |
| Nsp2 | SOS2 | 0.594 | 6.98E-07 |
| Nsp4 | PIK3CA | 0.594 | 6.98E-07 |
| Nsp5 | ROCK2 | 0.594 | 6.96E-07 |
| Nsp12 | RAP1B | 0.594 | 7.13E-07 |
| S | TLR3 | 0.594 | 7.25E-07 |

| | | | |
|---|---|---|---|
| Orf7a | TFRC | 0.594 | 6.92E-07 |
| Orf8 | IL13RA2 | 0.594 | 7.16E-07 |
| Nsp2 | ITCH | 0.593 | 7.49E-07 |
| Nsp3 | RBL2 | 0.593 | 7.55E-07 |
| Nsp6 | STK3 | 0.593 | 7.50E-07 |
| Nsp12 | GNB4 | 0.593 | 7.44E-07 |
| Nsp12 | FZD6 | 0.593 | 7.62E-07 |
| Nsp15 | PTPN13 | 0.593 | 7.64E-07 |
| Nsp15 | RBL2 | 0.593 | 7.35E-07 |
| N | COL1A2 | 0.593 | 7.43E-07 |
| Nsp2 | COL9A1 | 0.592 | 7.75E-07 |
| Nsp4 | RBL1 | 0.592 | 7.85E-07 |
| Nsp6 | PTPN2 | 0.592 | 7.86E-07 |
| Nsp12 | HSP90B1 | 0.592 | 7.77E-07 |
| Nsp13 | IL23R | 0.592 | 7.99E-07 |
| Nsp13 | IL1R1 | 0.592 | 7.98E-07 |
| Nsp14 | JAK2 | 0.592 | 7.96E-07 |
| Nsp15 | MAPK6 | 0.592 | 8.03E-07 |
| S | PPP2R5E | 0.592 | 7.72E-07 |
| Orf3a | IFNAR1 | 0.592 | 7.98E-07 |
| Orf7a | ITGA1 | 0.592 | 7.69E-07 |
| Nsp2 | EIF2S1 | 0.591 | 8.33E-07 |
| Nsp4 | ITGA4 | 0.591 | 8.16E-07 |
| Nsp4 | ATP6V1A | 0.591 | 8.39E-07 |
| Nsp5 | COL9A1 | 0.591 | 8.42E-07 |
| Nsp6 | NLK | 0.591 | 8.25E-07 |
| Nsp6 | IL7 | 0.591 | 8.18E-07 |
| Nsp12 | MAPK6 | 0.591 | 8.40E-07 |
| Nsp13 | ITGA2 | 0.591 | 8.36E-07 |
| S | VCAM1 | 0.591 | 8.26E-07 |
| S | STRADB | 0.591 | 8.28E-07 |
| Orf3a | ITGA2 | 0.591 | 8.10E-07 |
| Orf3a | PLA2G4A | 0.591 | 8.21E-07 |
| Orf7a | PPP3CB | 0.591 | 8.50E-07 |
| Orf7a | RPS6KA6 | 0.591 | 8.23E-07 |
| Nsp8 | RAP1B | 0.59 | 8.94E-07 |
| Nsp8 | PIK3CA | 0.59 | 8.67E-07 |
| Nsp13 | MAP3K2 | 0.59 | 8.61E-07 |
| Nsp15 | PPM1B | 0.59 | 8.76E-07 |
| Nsp15 | PPM1A | 0.59 | 8.86E-07 |
| S | ITGB8 | 0.59 | 8.90E-07 |
| Orf7a | ITGA2 | 0.59 | 8.98E-07 |
| Orf7a | LEPR | 0.59 | 8.80E-07 |
| Nsp3 | IL33 | 0.589 | 9.15E-07 |
| Nsp4 | CAB39 | 0.589 | 9.29E-07 |
| Nsp4 | PPP2CA | 0.589 | 9.48E-07 |
| Nsp8 | ITGB1 | 0.589 | 9.32E-07 |
| Nsp8 | ZFYVE9 | 0.589 | 9.17E-07 |
| Nsp9 | ZFYVE16 | 0.589 | 9.27E-07 |
| Nsp14 | GNB4 | 0.589 | 9.43E-07 |

| | | | |
|---|---|---|---|
| Nsp14 | ITGA1 | 0.589 | 9.21E-07 |
| Nsp15 | FNIP1 | 0.589 | 9.37E-07 |
| Nsp16 | PRKAA2 | 0.589 | 9.15E-07 |
| Nsp3 | CD86 | 0.588 | 9.60E-07 |
| Nsp5 | ITCH | 0.588 | 9.94E-07 |
| Nsp6 | AKT3 | 0.588 | 9.83E-07 |
| Nsp9 | PPM1B | 0.588 | 9.84E-07 |
| Nsp9 | MIOS | 0.588 | 9.80E-07 |
| Nsp13 | ITGB1 | 0.588 | 9.51E-07 |
| Nsp13 | SMURF2 | 0.588 | 9.91E-07 |
| Nsp13 | PIK3CB | 0.588 | 9.89E-07 |
| N | HSP90B1 | 0.588 | 9.82E-07 |
| Orf3a | EIF4E | 0.588 | 9.56E-07 |
| Nsp2 | HSPA8 | 0.587 | 1.05E-06 |
| Nsp2 | PPP2R1B | 0.587 | 1.01E-06 |
| Nsp3 | FZD3 | 0.587 | 1.03E-06 |
| Nsp5 | IFNAR1 | 0.587 | 1.04E-06 |
| Nsp12 | PIAS2 | 0.587 | 1.02E-06 |
| Nsp14 | RBL1 | 0.587 | 1.02E-06 |
| Nsp15 | EIF2S1 | 0.587 | 1.00E-06 |
| Orf8 | XIAP | 0.587 | 1.04E-06 |
| Nsp6 | MIOS | 0.586 | 1.09E-06 |
| Nsp8 | RASA1 | 0.586 | 1.11E-06 |
| Nsp8 | MAP3K7 | 0.586 | 1.07E-06 |
| Nsp13 | EGF | 0.586 | 1.07E-06 |
| Nsp14 | COL1A2 | 0.586 | 1.06E-06 |
| Nsp15 | PLA2G4A | 0.586 | 1.08E-06 |
| Nsp2 | CD36 | 0.585 | 1.15E-06 |
| Nsp6 | CUL2 | 0.585 | 1.15E-06 |
| Nsp6 | CYLD | 0.585 | 1.17E-06 |
| Nsp8 | RBL1 | 0.585 | 1.15E-06 |
| Nsp4 | FNIP1 | 0.584 | 1.20E-06 |
| Nsp4 | PPP2R1B | 0.584 | 1.21E-06 |
| Nsp15 | COL4A5 | 0.584 | 1.21E-06 |
| Orf7a | AZI2 | 0.584 | 1.18E-06 |
| Nsp5 | RBL1 | 0.583 | 1.25E-06 |
| Nsp8 | AKT3 | 0.583 | 1.28E-06 |
| Nsp13 | ACSL3 | 0.583 | 1.29E-06 |
| Nsp14 | SOCS5 | 0.583 | 1.29E-06 |
| Nsp15 | SOCS5 | 0.583 | 1.26E-06 |
| Nsp15 | MDM2 | 0.583 | 1.26E-06 |
| Nsp16 | IL1R1 | 0.583 | 1.29E-06 |
| S | MAP3K2 | 0.583 | 1.29E-06 |
| Orf8 | COL4A5 | 0.583 | 1.25E-06 |
| Nsp2 | IL6ST | 0.582 | 1.32E-06 |
| Nsp4 | PRKAA1 | 0.582 | 1.33E-06 |
| Nsp5 | PIK3CA | 0.582 | 1.30E-06 |
| Nsp8 | GNB4 | 0.582 | 1.30E-06 |
| Nsp12 | PTPN2 | 0.582 | 1.35E-06 |
| Nsp13 | FZD3 | 0.582 | 1.31E-06 |

| | | | |
|---|---|---|---|
| Nsp13 | HGF | 0.582 | 1.32E-06 |
| Nsp14 | IL7 | 0.582 | 1.31E-06 |
| Orf3a | ROCK2 | 0.582 | 1.34E-06 |
| Orf3a | DNM1L | 0.582 | 1.32E-06 |
| Orf7a | USP25 | 0.582 | 1.35E-06 |
| Orf7a | PIAS2 | 0.582 | 1.34E-06 |
| Nsp5 | IL7 | 0.581 | 1.43E-06 |
| Nsp6 | ITGAV | 0.581 | 1.38E-06 |
| Nsp8 | ITGA2 | 0.581 | 1.43E-06 |
| Nsp12 | EIF2S1 | 0.581 | 1.37E-06 |
| Orf7a | IL18R1 | 0.581 | 1.40E-06 |
| Nsp2 | GNB4 | 0.58 | 1.44E-06 |
| Nsp2 | HIF1A | 0.58 | 1.46E-06 |
| Nsp4 | ITGA1 | 0.58 | 1.50E-06 |
| Nsp4 | RASA1 | 0.58 | 1.47E-06 |
| Nsp6 | STRADB | 0.58 | 1.50E-06 |
| Nsp8 | ATP6V1A | 0.58 | 1.52E-06 |
| Nsp9 | PPP3CB | 0.58 | 1.46E-06 |
| Nsp13 | TAB2 | 0.58 | 1.51E-06 |
| Nsp13 | SOCS4 | 0.58 | 1.49E-06 |
| Nsp13 | EIF2S1 | 0.58 | 1.45E-06 |
| Orf3a | ATF2 | 0.58 | 1.44E-06 |
| Nsp6 | SOCS5 | 0.579 | 1.58E-06 |
| Nsp6 | NFATC3 | 0.579 | 1.58E-06 |
| Nsp12 | RASA1 | 0.579 | 1.59E-06 |
| Nsp14 | MAP3K1 | 0.579 | 1.52E-06 |
| Nsp15 | LPAR6 | 0.579 | 1.54E-06 |
| Orf3a | CUL2 | 0.579 | 1.52E-06 |
| Orf7a | NFATC3 | 0.579 | 1.54E-06 |
| Orf7a | ITGB1 | 0.579 | 1.59E-06 |
| Orf8 | MAP4K3 | 0.579 | 1.54E-06 |
| Orf8 | IL6ST | 0.579 | 1.57E-06 |
| Nsp3 | MALT1 | 0.578 | 1.62E-06 |
| Nsp12 | CAMK2D | 0.578 | 1.60E-06 |
| Nsp12 | TAOK1 | 0.578 | 1.62E-06 |
| Nsp13 | RASA2 | 0.578 | 1.66E-06 |
| Nsp16 | RASA2 | 0.578 | 1.68E-06 |
| Orf3a | MDM2 | 0.578 | 1.65E-06 |
| Orf3a | HSP90B1 | 0.578 | 1.62E-06 |
| Orf7a | PTPN2 | 0.578 | 1.64E-06 |
| Orf7a | KRAS | 0.578 | 1.61E-06 |
| Nsp4 | ITGA2 | 0.577 | 1.74E-06 |
| Nsp6 | RICTOR | 0.577 | 1.74E-06 |
| Nsp12 | CACNA2D1 | 0.577 | 1.68E-06 |
| Nsp13 | GNB4 | 0.577 | 1.74E-06 |
| Nsp15 | PPP2R3C | 0.577 | 1.69E-06 |
| Orf3a | RASA2 | 0.577 | 1.68E-06 |
| Nsp2 | LEPR | 0.576 | 1.83E-06 |
| Nsp3 | MAP3K2 | 0.576 | 1.77E-06 |
| Nsp6 | MMP13 | 0.576 | 1.82E-06 |

| | | | |
|---|---|---|---|
| Nsp6 | STAT4 | 0.576 | 1.82E-06 |
| Nsp6 | COL1A2 | 0.576 | 1.83E-06 |
| Nsp9 | PKN2 | 0.576 | 1.79E-06 |
| Nsp12 | ITGA1 | 0.576 | 1.84E-06 |
| Nsp15 | TICAM2 | 0.576 | 1.84E-06 |
| Orf7a | TAB2 | 0.576 | 1.85E-06 |
| Nsp3 | IL1R1 | 0.575 | 1.89E-06 |
| Nsp4 | SOCS5 | 0.575 | 1.87E-06 |
| Nsp4 | FAS | 0.575 | 1.93E-06 |
| Nsp6 | TFRC | 0.575 | 1.89E-06 |
| Nsp14 | PIK3CA | 0.575 | 1.93E-06 |
| Nsp15 | STK3 | 0.575 | 1.91E-06 |
| S | EIF4E | 0.575 | 1.94E-06 |
| S | MAP3K4 | 0.575 | 1.88E-06 |
| Orf7a | TLR6 | 0.575 | 1.89E-06 |
| Nsp4 | AKT3 | 0.574 | 2.03E-06 |
| Nsp4 | MAP3K7 | 0.574 | 2.02E-06 |
| Nsp8 | TAB2 | 0.574 | 2.02E-06 |
| Nsp8 | PRKACB | 0.574 | 1.96E-06 |
| Nsp13 | MAP3K1 | 0.574 | 1.98E-06 |
| Nsp13 | COL9A1 | 0.574 | 1.99E-06 |
| Nsp16 | TICAM2 | 0.574 | 1.97E-06 |
| Orf3a | SOS2 | 0.574 | 2.05E-06 |
| Orf7a | TANK | 0.574 | 1.97E-06 |
| Orf7a | BIRC2 | 0.574 | 2.03E-06 |
| Nsp3 | CAB39 | 0.573 | 2.14E-06 |
| Nsp4 | RBL2 | 0.573 | 2.13E-06 |
| Nsp4 | TGFBR1 | 0.573 | 2.06E-06 |
| Nsp5 | TAB3 | 0.573 | 2.07E-06 |
| Nsp5 | COL1A2 | 0.573 | 2.08E-06 |
| Nsp8 | NF1 | 0.573 | 2.09E-06 |
| Nsp12 | IFIH1 | 0.573 | 2.13E-06 |
| Nsp12 | CDC42 | 0.573 | 2.06E-06 |
| Nsp13 | PPP2R3C | 0.573 | 2.10E-06 |
| Nsp14 | FZD3 | 0.573 | 2.15E-06 |
| S | CAB39 | 0.573 | 2.15E-06 |
| N | LAMA2 | 0.573 | 2.11E-06 |
| Nsp4 | ITGB1 | 0.572 | 2.19E-06 |
| Nsp5 | CYLD | 0.572 | 2.19E-06 |
| Nsp6 | COL4A4 | 0.572 | 2.21E-06 |
| Nsp12 | PIK3CA | 0.572 | 2.25E-06 |
| Nsp13 | ITGAV | 0.572 | 2.25E-06 |
| Nsp13 | IL18R1 | 0.572 | 2.24E-06 |
| Nsp14 | RAP1B | 0.572 | 2.26E-06 |
| Orf3a | BRCA1 | 0.572 | 2.27E-06 |
| Nsp2 | CAMK2D | 0.571 | 2.32E-06 |
| Nsp2 | ITGAV | 0.571 | 2.28E-06 |
| Nsp3 | ACSL3 | 0.571 | 2.28E-06 |
| Nsp6 | PPP2R5E | 0.571 | 2.36E-06 |
| Nsp6 | APAF1 | 0.571 | 2.29E-06 |

| | | | |
|---|---|---|---|
| Nsp13 | BMPR2 | 0.571 | 2.37E-06 |
| Nsp14 | TICAM2 | 0.571 | 2.37E-06 |
| Nsp14 | SEH1L | 0.571 | 2.32E-06 |
| Nsp15 | TLR6 | 0.571 | 2.35E-06 |
| S | NF1 | 0.571 | 2.35E-06 |
| S | MAPK9 | 0.571 | 2.29E-06 |
| Orf7a | AREG | 0.571 | 2.34E-06 |
| Nsp2 | IFNAR1 | 0.57 | 2.47E-06 |
| Nsp4 | LDHC | 0.57 | 2.44E-06 |
| Nsp6 | CCNE2 | 0.57 | 2.39E-06 |
| Nsp8 | PPP2R3C | 0.57 | 2.46E-06 |
| Nsp9 | RPS6KA3 | 0.57 | 2.44E-06 |
| Nsp12 | SOCS4 | 0.57 | 2.43E-06 |
| Nsp12 | TGFBR1 | 0.57 | 2.43E-06 |
| Nsp16 | RASA1 | 0.57 | 2.42E-06 |
| S | COL4A4 | 0.57 | 2.43E-06 |
| S | LAMA2 | 0.57 | 2.42E-06 |
| Orf3a | FNIP1 | 0.57 | 2.48E-06 |
| Orf8 | USP25 | 0.57 | 2.45E-06 |
| Nsp2 | TBK1 | 0.569 | 2.60E-06 |
| Nsp3 | STRADB | 0.569 | 2.56E-06 |
| Nsp3 | PPP2R5E | 0.569 | 2.59E-06 |
| Nsp6 | PPP2R3A | 0.569 | 2.55E-06 |
| Nsp6 | CD86 | 0.569 | 2.60E-06 |
| Nsp13 | KRAS | 0.569 | 2.62E-06 |
| Nsp15 | MALT1 | 0.569 | 2.59E-06 |
| Orf7a | MAPK9 | 0.569 | 2.52E-06 |
| Nsp1 | LDHAL6B | 0.568 | 2.66E-06 |
| Nsp3 | IL23R | 0.568 | 2.66E-06 |
| Nsp5 | SOCS5 | 0.568 | 2.69E-06 |
| Nsp5 | BIRC3 | 0.568 | 2.69E-06 |
| Nsp9 | ITGA2 | 0.568 | 2.66E-06 |
| Nsp12 | ATP6V1A | 0.568 | 2.64E-06 |
| Nsp16 | CCNE2 | 0.568 | 2.64E-06 |
| Orf3a | IL7 | 0.568 | 2.69E-06 |
| Orf3a | KRAS | 0.568 | 2.73E-06 |
| Nsp5 | STAM | 0.567 | 2.90E-06 |
| Nsp5 | ITGA1 | 0.567 | 2.85E-06 |
| Nsp5 | PIK3CB | 0.567 | 2.80E-06 |
| Nsp8 | PTPN2 | 0.567 | 2.78E-06 |
| Nsp9 | LDHC | 0.567 | 2.88E-06 |
| Nsp12 | MAP3K7 | 0.567 | 2.81E-06 |
| Nsp13 | ATG5 | 0.567 | 2.90E-06 |
| Nsp16 | TGFBR1 | 0.567 | 2.86E-06 |
| S | IL33 | 0.567 | 2.82E-06 |
| Orf3a | IFIH1 | 0.567 | 2.78E-06 |
| Orf3a | LIFR | 0.567 | 2.82E-06 |
| Orf3a | PTPN13 | 0.567 | 2.86E-06 |
| Orf3a | HIF1A | 0.567 | 2.80E-06 |
| Orf7a | FGF7 | 0.567 | 2.91E-06 |

| | | | |
|---|---|---|---|
| Nsp3 | PARP4 | 0.566 | 2.96E-06 |
| Nsp6 | AZI2 | 0.566 | 3.05E-06 |
| Nsp8 | PTPN13 | 0.566 | 2.97E-06 |
| Nsp13 | CAB39 | 0.566 | 2.92E-06 |
| Nsp14 | TAB2 | 0.566 | 2.92E-06 |
| Nsp14 | ACVR2A | 0.566 | 2.92E-06 |
| Nsp14 | CDC42 | 0.566 | 2.97E-06 |
| Nsp15 | LDHC | 0.566 | 2.94E-06 |
| S | ACSL4 | 0.566 | 2.93E-06 |
| M | APAF1 | 0.566 | 3.01E-06 |
| Orf3a | SOCS5 | 0.566 | 3.01E-06 |
| Orf7a | RICTOR | 0.566 | 2.94E-06 |
| Orf7a | RASA2 | 0.566 | 3.04E-06 |
| Orf8 | PPP3CB | 0.566 | 2.94E-06 |
| Orf8 | MAP3K7 | 0.566 | 2.93E-06 |
| Nsp3 | SMURF2 | 0.565 | 3.07E-06 |
| Nsp3 | COL4A4 | 0.565 | 3.15E-06 |
| Nsp6 | PPP2R3C | 0.565 | 3.19E-06 |
| Nsp6 | JAK2 | 0.565 | 3.14E-06 |
| Nsp6 | PPM1B | 0.565 | 3.13E-06 |
| Nsp6 | GNB4 | 0.565 | 3.15E-06 |
| Nsp13 | STK3 | 0.565 | 3.07E-06 |
| Nsp14 | PPM1B | 0.565 | 3.09E-06 |
| Nsp14 | SOCS4 | 0.565 | 3.19E-06 |
| Nsp14 | AREG | 0.565 | 3.19E-06 |
| Nsp14 | FNIP1 | 0.565 | 3.16E-06 |
| Nsp16 | LPAR6 | 0.565 | 3.21E-06 |
| Orf3a | RAP1B | 0.565 | 3.10E-06 |
| Orf3a | APAF1 | 0.565 | 3.17E-06 |
| Orf7a | FAS | 0.565 | 3.12E-06 |
| Nsp2 | FNIP1 | 0.564 | 3.21E-06 |
| Nsp6 | FZD3 | 0.564 | 3.21E-06 |
| Nsp8 | SMAD4 | 0.564 | 3.22E-06 |
| Nsp9 | PPP2R2A | 0.564 | 3.33E-06 |
| Nsp14 | PRKAA2 | 0.564 | 3.33E-06 |
| N | IL13RA2 | 0.564 | 3.31E-06 |
| Orf7a | PRKACB | 0.564 | 3.31E-06 |
| Orf8 | HIF1A | 0.564 | 3.35E-06 |
| Nsp2 | BIRC3 | 0.563 | 3.46E-06 |
| Nsp4 | CAMK2D | 0.563 | 3.45E-06 |
| Nsp4 | FZD3 | 0.563 | 3.50E-06 |
| Nsp4 | GNAI1 | 0.563 | 3.38E-06 |
| Nsp6 | MAP3K20 | 0.563 | 3.51E-06 |
| Nsp8 | CASP3 | 0.563 | 3.37E-06 |
| Nsp9 | LDHAL6A | 0.563 | 3.51E-06 |
| Nsp16 | DDX58 | 0.563 | 3.49E-06 |
| S | ITGA6 | 0.563 | 3.47E-06 |
| Orf7a | PARP4 | 0.563 | 3.42E-06 |
| Orf7a | HIF1A | 0.563 | 3.39E-06 |
| Nsp2 | FZD6 | 0.562 | 3.71E-06 |

| | | | |
|---|---|---|---|
| Nsp3 | LPAR6 | 0.562 | 3.67E-06 |
| Nsp4 | TAOK1 | 0.562 | 3.58E-06 |
| Nsp5 | GNB4 | 0.562 | 3.71E-06 |
| Nsp5 | HSP90B1 | 0.562 | 3.58E-06 |
| Nsp12 | MAP3K2 | 0.562 | 3.63E-06 |
| Nsp13 | PPP2R5E | 0.562 | 3.59E-06 |
| Nsp16 | PARP4 | 0.562 | 3.61E-06 |
| Nsp16 | AZI2 | 0.562 | 3.54E-06 |
| N | COL4A5 | 0.562 | 3.54E-06 |
| Orf8 | PPP2R1B | 0.562 | 3.57E-06 |
| Nsp2 | SGK3 | 0.561 | 3.72E-06 |
| Nsp3 | SMAD4 | 0.561 | 3.74E-06 |
| Nsp6 | ITGA4 | 0.561 | 3.87E-06 |
| Nsp6 | PIK3CB | 0.561 | 3.74E-06 |
| Nsp8 | CD86 | 0.561 | 3.89E-06 |
| Nsp8 | TAB3 | 0.561 | 3.87E-06 |
| Nsp14 | COL4A3 | 0.561 | 3.72E-06 |
| Nsp15 | PRKAA1 | 0.561 | 3.75E-06 |
| N | COL4A6 | 0.561 | 3.76E-06 |
| Orf3a | FAS | 0.561 | 3.73E-06 |
| Orf7a | IL13RA2 | 0.561 | 3.75E-06 |
| Nsp2 | USP25 | 0.56 | 4.00E-06 |
| Nsp4 | CACNA2D1 | 0.56 | 4.06E-06 |
| Nsp5 | CCNE2 | 0.56 | 3.91E-06 |
| Nsp6 | TAOK1 | 0.56 | 3.93E-06 |
| Nsp15 | TLR8 | 0.56 | 3.99E-06 |
| Nsp16 | IFIH1 | 0.56 | 3.91E-06 |
| Nsp16 | MAP3K1 | 0.56 | 4.03E-06 |
| S | SMAD5 | 0.56 | 4.09E-06 |
| S | MAPK8 | 0.56 | 4.04E-06 |
| Orf7a | AHR | 0.56 | 4.07E-06 |
| Orf7a | BRCA1 | 0.56 | 3.90E-06 |
| Nsp2 | MAPK9 | 0.559 | 4.17E-06 |
| Nsp3 | PPP2CA | 0.559 | 4.18E-06 |
| Nsp9 | ROCK2 | 0.559 | 4.24E-06 |
| Nsp9 | COL4A5 | 0.559 | 4.20E-06 |
| Nsp14 | COL4A6 | 0.559 | 4.25E-06 |
| Nsp14 | VCAM1 | 0.559 | 4.11E-06 |
| Nsp15 | PIK3CB | 0.559 | 4.28E-06 |
| Nsp16 | ITGAV | 0.559 | 4.10E-06 |
| M | COL4A5 | 0.559 | 4.21E-06 |
| Orf7a | FLT3 | 0.559 | 4.15E-06 |
| Orf8 | ITCH | 0.559 | 4.25E-06 |
| Nsp2 | PIK3CA | 0.558 | 4.44E-06 |
| Nsp12 | TFRC | 0.558 | 4.36E-06 |
| Nsp12 | COL1A2 | 0.558 | 4.46E-06 |
| Nsp15 | CCNE2 | 0.558 | 4.50E-06 |
| Nsp16 | VCAM1 | 0.558 | 4.40E-06 |
| N | ITGAV | 0.558 | 4.40E-06 |
| N | ITGA1 | 0.558 | 4.32E-06 |

| | | | |
|---|---|---|---|
| Nsp4 | AZI2 | 0.557 | 4.59E-06 |
| Nsp4 | PIK3CB | 0.557 | 4.72E-06 |
| Nsp5 | RAP1B | 0.557 | 4.58E-06 |
| Nsp6 | LDHC | 0.557 | 4.65E-06 |
| Nsp12 | PPM1A | 0.557 | 4.68E-06 |
| Nsp13 | LDHC | 0.557 | 4.65E-06 |
| N | PHKB | 0.557 | 4.70E-06 |
| N | RPS6KA3 | 0.557 | 4.56E-06 |
| Orf7a | EGF | 0.557 | 4.53E-06 |
| Orf7a | CUL2 | 0.557 | 4.72E-06 |
| Nsp4 | RAP1B | 0.556 | 4.79E-06 |
| Nsp9 | LIFR | 0.556 | 4.82E-06 |
| Nsp12 | LDHC | 0.556 | 4.81E-06 |
| Nsp15 | CACNA2D1 | 0.556 | 4.72E-06 |
| N | HIF1A | 0.556 | 4.75E-06 |
| Nsp3 | BMPR2 | 0.555 | 4.96E-06 |
| Nsp5 | IFIH1 | 0.555 | 5.03E-06 |
| Nsp8 | TLR6 | 0.555 | 5.06E-06 |
| Nsp9 | CUL2 | 0.555 | 5.18E-06 |
| Nsp13 | STAT4 | 0.555 | 5.16E-06 |
| Nsp13 | DDX3X | 0.555 | 5.03E-06 |
| Nsp14 | NLK | 0.555 | 5.16E-06 |
| Nsp15 | ITGA4 | 0.555 | 5.14E-06 |
| Nsp16 | CAB39 | 0.555 | 5.08E-06 |
| S | SMAD4 | 0.555 | 5.14E-06 |
| Orf8 | CD36 | 0.555 | 5.04E-06 |
| Nsp2 | IFIH1 | 0.554 | 5.37E-06 |
| Nsp2 | DDX3X | 0.554 | 5.40E-06 |
| Nsp3 | TLR8 | 0.554 | 5.24E-06 |
| Nsp9 | RICTOR | 0.554 | 5.29E-06 |
| Nsp12 | PRKAA2 | 0.554 | 5.41E-06 |
| Nsp12 | RASA2 | 0.554 | 5.30E-06 |
| Nsp14 | PPP2R3A | 0.554 | 5.28E-06 |
| Nsp14 | MMP13 | 0.554 | 5.31E-06 |
| Nsp14 | CUL2 | 0.554 | 5.30E-06 |
| Nsp15 | NFATC3 | 0.554 | 5.38E-06 |
| Nsp15 | MSTN | 0.554 | 5.26E-06 |
| Nsp5 | LDHC | 0.553 | 5.52E-06 |
| Nsp6 | PPP2R1B | 0.553 | 5.45E-06 |
| Nsp8 | KRAS | 0.553 | 5.45E-06 |
| Nsp13 | GNAI1 | 0.553 | 5.56E-06 |
| Nsp13 | CDC42 | 0.553 | 5.46E-06 |
| Nsp13 | COL4A4 | 0.553 | 5.58E-06 |
| Nsp14 | TNFSF18 | 0.553 | 5.45E-06 |
| S | CXCL10 | 0.553 | 5.55E-06 |
| S | EIF2AK3 | 0.553 | 5.57E-06 |
| S | LPAR6 | 0.553 | 5.68E-06 |
| N | ITGA4 | 0.553 | 5.45E-06 |
| N | ATM | 0.553 | 5.62E-06 |
| Orf3a | TAB2 | 0.553 | 5.64E-06 |

| | | | |
|---|---|---|---|
| Orf7a | MDM2 | 0.553 | 5.49E-06 |
| Orf7a | SMURF2 | 0.553 | 5.57E-06 |
| Orf7a | STK3 | 0.553 | 5.52E-06 |
| Nsp5 | JAK2 | 0.552 | 5.90E-06 |
| Nsp6 | IL18R1 | 0.552 | 5.94E-06 |
| Nsp8 | CAMK2D | 0.552 | 5.89E-06 |
| Nsp9 | SOCS4 | 0.552 | 5.86E-06 |
| Nsp12 | ITGA4 | 0.552 | 5.74E-06 |
| Nsp12 | FNIP1 | 0.552 | 5.88E-06 |
| Nsp12 | COL4A3 | 0.552 | 5.78E-06 |
| Orf3a | ZFYVE9 | 0.552 | 5.85E-06 |
| Orf7a | PPM1B | 0.552 | 5.71E-06 |
| Nsp3 | GNAI1 | 0.551 | 6.05E-06 |
| Nsp4 | IL18R1 | 0.551 | 6.16E-06 |
| Nsp5 | HIF1A | 0.551 | 6.17E-06 |
| Nsp6 | IFNAR2 | 0.551 | 6.03E-06 |
| Nsp6 | FAS | 0.551 | 6.16E-06 |
| Nsp9 | RASA2 | 0.551 | 6.02E-06 |
| Nsp13 | PHKB | 0.551 | 6.22E-06 |
| Nsp16 | STAT4 | 0.551 | 6.25E-06 |
| M | DDX3X | 0.551 | 6.16E-06 |
| N | DDX58 | 0.551 | 6.25E-06 |
| Orf7a | MAPK6 | 0.551 | 6.06E-06 |
| Orf7a | PIK3CB | 0.551 | 6.22E-06 |
| Orf8 | DNM1L | 0.551 | 5.98E-06 |
| Nsp3 | CXCL10 | 0.55 | 6.44E-06 |
| Nsp4 | SMAD4 | 0.55 | 6.27E-06 |
| Nsp4 | COL4A3 | 0.55 | 6.49E-06 |
| Nsp5 | PPP3CB | 0.55 | 6.52E-06 |
| Nsp5 | TGFBR1 | 0.55 | 6.42E-06 |
| Nsp6 | RAP1B | 0.55 | 6.40E-06 |
| Nsp8 | SMAD5 | 0.55 | 6.42E-06 |
| Nsp13 | MALT1 | 0.55 | 6.31E-06 |
| Nsp14 | MAPK6 | 0.55 | 6.45E-06 |
| Nsp14 | HSP90B1 | 0.55 | 6.55E-06 |
| Nsp16 | KRAS | 0.55 | 6.50E-06 |
| M | RPS6KA3 | 0.55 | 6.48E-06 |
| N | ATF2 | 0.55 | 6.51E-06 |
| Orf7a | MIOS | 0.55 | 6.33E-06 |
| Orf8 | TAB3 | 0.55 | 6.35E-06 |
| Nsp3 | PPP2R1B | 0.549 | 6.83E-06 |
| Nsp4 | IL33 | 0.549 | 6.58E-06 |
| Nsp5 | EIF2S1 | 0.549 | 6.63E-06 |
| Nsp8 | PIK3CB | 0.549 | 6.83E-06 |
| Nsp9 | TAB2 | 0.549 | 6.73E-06 |
| Nsp9 | ITGB1 | 0.549 | 6.69E-06 |
| Nsp10 | IRAK4 | 0.549 | 6.70E-06 |
| Nsp15 | SMURF2 | 0.549 | 6.67E-06 |
| Nsp15 | TGFBR1 | 0.549 | 6.75E-06 |
| S | MECOM | 0.549 | 6.68E-06 |

| | | | |
|---|---|---|---|
| Orf7a | MAPK8 | 0.549 | 6.79E-06 |
| Orf7a | COL9A1 | 0.549 | 6.69E-06 |
| Orf8 | RAP1B | 0.549 | 6.87E-06 |
| Nsp2 | ZFYVE16 | 0.548 | 6.98E-06 |
| Nsp3 | BCL10 | 0.548 | 7.04E-06 |
| Nsp5 | ITGAV | 0.548 | 7.15E-06 |
| Nsp6 | FZD6 | 0.548 | 6.95E-06 |
| Nsp9 | BIRC2 | 0.548 | 7.04E-06 |
| Nsp9 | EIF4E | 0.548 | 7.16E-06 |
| Nsp12 | IL18R1 | 0.548 | 7.02E-06 |
| Nsp14 | STAT4 | 0.548 | 7.08E-06 |
| Nsp16 | GNB4 | 0.548 | 7.16E-06 |
| Orf7a | IFNAR1 | 0.548 | 6.90E-06 |
| Orf7a | DNM1L | 0.548 | 7.00E-06 |
| Nsp2 | RAP1B | 0.547 | 7.28E-06 |
| Nsp5 | STAT4 | 0.547 | 7.49E-06 |
| Nsp8 | CACNA2D1 | 0.547 | 7.51E-06 |
| Nsp8 | IL23R | 0.547 | 7.50E-06 |
| Nsp9 | AHR | 0.547 | 7.49E-06 |
| Orf7a | CCNE2 | 0.547 | 7.38E-06 |
| Orf7a | ATP6V1A | 0.547 | 7.50E-06 |
| Nsp3 | ATG5 | 0.546 | 7.86E-06 |
| Nsp3 | COL4A6 | 0.546 | 7.63E-06 |
| Nsp6 | SMAD5 | 0.546 | 7.83E-06 |
| Nsp9 | RPS6KA6 | 0.546 | 7.59E-06 |
| Nsp12 | CD86 | 0.546 | 7.84E-06 |
| Nsp12 | CYLD | 0.546 | 7.67E-06 |
| Nsp13 | TLR8 | 0.546 | 7.79E-06 |
| Nsp15 | CYLD | 0.546 | 7.85E-06 |
| Nsp16 | MAP3K7 | 0.546 | 7.90E-06 |
| Orf7a | LAMA2 | 0.546 | 7.57E-06 |
| Nsp2 | LRP6 | 0.545 | 8.01E-06 |
| Nsp3 | ITGB8 | 0.545 | 8.17E-06 |
| Nsp3 | CHUK | 0.545 | 7.97E-06 |
| Nsp8 | IFNGR1 | 0.545 | 8.23E-06 |
| Nsp9 | DNM1L | 0.545 | 8.24E-06 |
| Nsp14 | LPAR4 | 0.545 | 8.00E-06 |
| Nsp16 | BMPR2 | 0.545 | 8.01E-06 |
| Nsp16 | IL33 | 0.545 | 8.18E-06 |
| S | COL4A6 | 0.545 | 8.18E-06 |
| Nsp1 | LDHA | 0.544 | 8.41E-06 |
| Nsp5 | TLR6 | 0.544 | 8.38E-06 |
| Nsp8 | FZD6 | 0.544 | 8.40E-06 |
| Nsp8 | BMPR2 | 0.544 | 8.62E-06 |
| Nsp14 | MDM2 | 0.544 | 8.63E-06 |
| Nsp14 | IL22RA2 | 0.544 | 8.28E-06 |
| Nsp15 | IL1R1 | 0.544 | 8.35E-06 |
| Nsp16 | TLR8 | 0.544 | 8.47E-06 |
| S | SEH1L | 0.544 | 8.46E-06 |
| S | MMP13 | 0.544 | 8.52E-06 |

| | | | |
|---|---|---|---|
| Nsp3 | PHKB | 0.543 | 8.71E-06 |
| Nsp4 | ATG5 | 0.543 | 8.94E-06 |
| Nsp4 | IL22RA2 | 0.543 | 8.82E-06 |
| Nsp8 | ITGAV | 0.543 | 8.66E-06 |
| Nsp8 | TAOK1 | 0.543 | 8.74E-06 |
| Nsp14 | RPS6KB1 | 0.543 | 8.93E-06 |
| Orf3a | PHKB | 0.543 | 8.97E-06 |
| Orf7a | IL21 | 0.543 | 8.81E-06 |
| Orf7a | ITGAV | 0.543 | 8.77E-06 |
| Nsp2 | RBL1 | 0.542 | 9.16E-06 |
| Nsp12 | IL1R1 | 0.542 | 9.17E-06 |
| Nsp13 | TNFSF18 | 0.542 | 9.33E-06 |
| Nsp13 | STRADB | 0.542 | 9.44E-06 |
| Nsp15 | AKT3 | 0.542 | 9.46E-06 |
| Nsp15 | CD36 | 0.542 | 9.16E-06 |
| Nsp16 | TFRC | 0.542 | 9.37E-06 |
| Orf3a | IL33 | 0.542 | 9.24E-06 |
| Nsp4 | SEH1L | 0.541 | 9.78E-06 |
| Nsp5 | IFNGR1 | 0.541 | 9.73E-06 |
| Nsp6 | EIF4E | 0.541 | 9.48E-06 |
| Nsp6 | SORBS1 | 0.541 | 9.53E-06 |
| Nsp9 | KRAS | 0.541 | 9.68E-06 |
| Nsp12 | SOCS5 | 0.541 | 9.62E-06 |
| Nsp12 | AKT3 | 0.541 | 9.79E-06 |
| Nsp16 | COL4A3 | 0.541 | 9.68E-06 |
| S | TANK | 0.541 | 9.48E-06 |
| Nsp3 | NF1 | 0.54 | 1.03E-05 |
| Nsp5 | SMURF2 | 0.54 | 1.03E-05 |
| Nsp5 | FNIP1 | 0.54 | 1.02E-05 |
| Nsp6 | IL1R1 | 0.54 | 1.03E-05 |
| S | SLC38A9 | 0.54 | 1.03E-05 |
| N | IRAK4 | 0.54 | 9.96E-06 |
| Orf3a | MIOS | 0.54 | 1.03E-05 |
| Orf3a | CACNA2D1 | 0.54 | 1.01E-05 |
| Orf3a | CD86 | 0.54 | 1.00E-05 |
| Orf7a | VCAM1 | 0.54 | 1.01E-05 |
| Orf7a | PIK3CA | 0.54 | 1.01E-05 |
| Orf8 | LDHC | 0.54 | 9.98E-06 |
| Nsp3 | MMP13 | 0.539 | 1.09E-05 |
| Nsp6 | TLR3 | 0.539 | 1.05E-05 |
| Nsp8 | COL1A2 | 0.539 | 1.06E-05 |
| Nsp13 | SMAD4 | 0.539 | 1.06E-05 |
| Nsp15 | PPP2R2A | 0.539 | 1.06E-05 |
| N | CACNA2D1 | 0.539 | 1.08E-05 |
| N | HGF | 0.539 | 1.07E-05 |
| Orf3a | AHR | 0.539 | 1.06E-05 |
| Orf3a | STK3 | 0.539 | 1.05E-05 |
| Orf3a | TAB3 | 0.539 | 1.07E-05 |
| Orf3a | SOS1 | 0.539 | 1.05E-05 |
| Nsp2 | MAP4K3 | 0.538 | 1.10E-05 |

| | | | |
|---|---|---|---|
| Nsp2 | ROCK1 | 0.538 | 1.10E-05 |
| Nsp4 | SMURF2 | 0.538 | 1.11E-05 |
| Nsp4 | PARP4 | 0.538 | 1.12E-05 |
| Nsp12 | PIK3CB | 0.538 | 1.11E-05 |
| Nsp13 | NF1 | 0.538 | 1.13E-05 |
| Nsp16 | MMP13 | 0.538 | 1.09E-05 |
| Nsp16 | TLR3 | 0.538 | 1.10E-05 |
| M | STAM | 0.538 | 1.12E-05 |
| N | PTPN13 | 0.538 | 1.13E-05 |
| Orf3a | FZD6 | 0.538 | 1.11E-05 |
| Orf3a | MAPK9 | 0.538 | 1.12E-05 |
| Orf7a | SOS2 | 0.538 | 1.12E-05 |
| Nsp2 | NFATC3 | 0.537 | 1.15E-05 |
| Nsp4 | ITGAV | 0.537 | 1.14E-05 |
| Nsp13 | SOCS5 | 0.537 | 1.18E-05 |
| Nsp13 | PPP2R3A | 0.537 | 1.16E-05 |
| S | TNFSF18 | 0.537 | 1.15E-05 |
| S | ATP6V1C1 | 0.537 | 1.18E-05 |
| Orf3a | JAK2 | 0.537 | 1.14E-05 |
| Orf7a | MET | 0.537 | 1.16E-05 |
| Orf8 | CCNE2 | 0.537 | 1.14E-05 |
| Orf8 | RICTOR | 0.537 | 1.17E-05 |
| Nsp3 | GHR | 0.536 | 1.23E-05 |
| Nsp6 | PRKAA1 | 0.536 | 1.20E-05 |
| Nsp6 | CAMK2D | 0.536 | 1.21E-05 |
| Nsp6 | TICAM2 | 0.536 | 1.24E-05 |
| Nsp6 | VCAM1 | 0.536 | 1.20E-05 |
| Nsp8 | STK3 | 0.536 | 1.19E-05 |
| Nsp9 | MAP3K7 | 0.536 | 1.19E-05 |
| Nsp9 | CD36 | 0.536 | 1.21E-05 |
| Nsp12 | PPP2CA | 0.536 | 1.19E-05 |
| Nsp14 | EIF4E | 0.536 | 1.22E-05 |
| Nsp14 | COL4A4 | 0.536 | 1.20E-05 |
| Nsp16 | MALT1 | 0.536 | 1.20E-05 |
| N | FNIP1 | 0.536 | 1.22E-05 |
| Orf3a | PPP2R3C | 0.536 | 1.21E-05 |
| Orf3a | ITCH | 0.536 | 1.22E-05 |
| Orf7a | JAK2 | 0.536 | 1.19E-05 |
| Nsp2 | TFRC | 0.535 | 1.26E-05 |
| Nsp2 | BMPR1A | 0.535 | 1.25E-05 |
| Nsp5 | RASA1 | 0.535 | 1.26E-05 |
| Nsp6 | MET | 0.535 | 1.27E-05 |
| Nsp6 | LPAR4 | 0.535 | 1.26E-05 |
| Nsp8 | EIF4E | 0.535 | 1.25E-05 |
| Nsp8 | NLK | 0.535 | 1.26E-05 |
| Nsp12 | ITGAV | 0.535 | 1.26E-05 |
| Nsp12 | MAPK9 | 0.535 | 1.25E-05 |
| Nsp15 | PPP2R3A | 0.535 | 1.29E-05 |
| M | SOS2 | 0.535 | 1.25E-05 |
| Nsp2 | RASA1 | 0.534 | 1.31E-05 |

| | | | |
|---|---|---|---|
| Nsp2 | XIAP | 0.534 | 1.32E-05 |
| Nsp3 | VCAM1 | 0.534 | 1.33E-05 |
| Nsp3 | MET | 0.534 | 1.35E-05 |
| Nsp8 | FAS | 0.534 | 1.31E-05 |
| Nsp8 | TFRC | 0.534 | 1.31E-05 |
| Nsp12 | STK3 | 0.534 | 1.34E-05 |
| Nsp12 | AZI2 | 0.534 | 1.35E-05 |
| N | EIF2S1 | 0.534 | 1.35E-05 |
| Orf7a | PRKAA1 | 0.534 | 1.34E-05 |
| Nsp3 | PDGFC | 0.533 | 1.37E-05 |
| Nsp3 | TLR3 | 0.533 | 1.36E-05 |
| Nsp6 | CHUK | 0.533 | 1.36E-05 |
| Nsp14 | PRKAA1 | 0.533 | 1.39E-05 |
| Nsp14 | LDHC | 0.533 | 1.40E-05 |
| Nsp14 | RASA1 | 0.533 | 1.41E-05 |
| Nsp14 | MAP3K7 | 0.533 | 1.38E-05 |
| Nsp15 | SMAD4 | 0.533 | 1.40E-05 |
| S | MET | 0.533 | 1.40E-05 |
| N | PPM1B | 0.533 | 1.38E-05 |
| Orf7a | PTPRR | 0.533 | 1.40E-05 |
| Orf8 | BRCA1 | 0.533 | 1.41E-05 |
| Nsp2 | PRKAA2 | 0.532 | 1.43E-05 |
| Nsp2 | PIAS2 | 0.532 | 1.44E-05 |
| Nsp2 | PIK3CB | 0.532 | 1.44E-05 |
| Nsp3 | EGF | 0.532 | 1.46E-05 |
| Nsp4 | MAP3K1 | 0.532 | 1.42E-05 |
| Nsp8 | TLR8 | 0.532 | 1.43E-05 |
| Nsp9 | CALM2 | 0.532 | 1.47E-05 |
| Nsp13 | MET | 0.532 | 1.48E-05 |
| Nsp14 | PPP2R1B | 0.532 | 1.45E-05 |
| Nsp15 | MAP3K2 | 0.532 | 1.46E-05 |
| Nsp16 | BCL10 | 0.532 | 1.46E-05 |
| Nsp16 | CXCL10 | 0.532 | 1.45E-05 |
| Nsp16 | ATP6V1A | 0.532 | 1.44E-05 |
| Nsp16 | NF1 | 0.532 | 1.48E-05 |
| Nsp16 | MAPK8 | 0.532 | 1.47E-05 |
| N | AIFM1 | 0.532 | 1.45E-05 |
| Orf3a | ATP6V1A | 0.532 | 1.43E-05 |
| Orf8 | AHR | 0.532 | 1.47E-05 |
| Nsp2 | RICTOR | 0.531 | 1.52E-05 |
| Nsp5 | GHR | 0.531 | 1.54E-05 |
| Nsp6 | FNIP1 | 0.531 | 1.52E-05 |
| Nsp6 | TGFBR1 | 0.531 | 1.50E-05 |
| Nsp6 | ITGA1 | 0.531 | 1.49E-05 |
| Nsp7 | TGFBR1 | 0.531 | 1.50E-05 |
| Nsp12 | NFATC3 | 0.531 | 1.51E-05 |
| Nsp12 | ITGA2 | 0.531 | 1.51E-05 |
| Nsp14 | ATG5 | 0.531 | 1.52E-05 |
| Nsp16 | ITGB8 | 0.531 | 1.52E-05 |
| S | MALT1 | 0.531 | 1.54E-05 |

| | | | |
|---|---|---|---|
| Orf7a | ITGA4 | 0.531 | 1.48E-05 |
| Nsp4 | DDX58 | 0.53 | 1.57E-05 |
| Nsp7 | PPP1R3A | 0.53 | 1.55E-05 |
| Nsp8 | AHR | 0.53 | 1.58E-05 |
| Nsp8 | RAPGEF2 | 0.53 | 1.56E-05 |
| Nsp9 | IL6ST | 0.53 | 1.56E-05 |
| Nsp9 | ZFYVE9 | 0.53 | 1.55E-05 |
| Nsp14 | SMURF2 | 0.53 | 1.59E-05 |
| Nsp15 | PPP2CB | 0.53 | 1.59E-05 |
| S | PPP2R1B | 0.53 | 1.56E-05 |
| N | DDX3X | 0.53 | 1.59E-05 |
| Orf3a | STAT4 | 0.53 | 1.61E-05 |
| Orf3a | PIAS2 | 0.53 | 1.58E-05 |
| Orf7a | RBL1 | 0.53 | 1.60E-05 |
| Nsp1 | LDHC | 0.529 | 1.62E-05 |
| Nsp2 | ACVR2A | 0.529 | 1.68E-05 |
| Nsp4 | PHKB | 0.529 | 1.69E-05 |
| Nsp8 | TLR3 | 0.529 | 1.65E-05 |
| Nsp9 | PPP2R3C | 0.529 | 1.68E-05 |
| Nsp9 | PARP4 | 0.529 | 1.67E-05 |
| Nsp14 | PHKB | 0.529 | 1.64E-05 |
| Nsp15 | RAP1B | 0.529 | 1.65E-05 |
| Nsp16 | PHKB | 0.529 | 1.64E-05 |
| Nsp16 | COL4A4 | 0.529 | 1.64E-05 |
| N | BMPR1A | 0.529 | 1.67E-05 |
| Orf3a | IFNGR1 | 0.529 | 1.63E-05 |
| Orf3a | LAMA2 | 0.529 | 1.62E-05 |
| Orf7a | LAMB4 | 0.529 | 1.64E-05 |
| Orf7a | BMPR2 | 0.529 | 1.66E-05 |
| Orf7a | STAT4 | 0.529 | 1.66E-05 |
| Nsp8 | IL33 | 0.528 | 1.73E-05 |
| Nsp9 | MAPK6 | 0.528 | 1.69E-05 |
| Nsp14 | GNAI1 | 0.528 | 1.73E-05 |
| Nsp14 | PPM1A | 0.528 | 1.76E-05 |
| S | LPAR4 | 0.528 | 1.76E-05 |
| N | ERBB4 | 0.528 | 1.72E-05 |
| N | FAS | 0.528 | 1.70E-05 |
| N | TFRC | 0.528 | 1.75E-05 |
| Orf3a | COL4A6 | 0.528 | 1.70E-05 |
| Orf3a | MAP3K1 | 0.528 | 1.74E-05 |
| Orf7a | COL1A2 | 0.528 | 1.74E-05 |
| Orf8 | RASA2 | 0.528 | 1.71E-05 |
| Nsp5 | PPP2R3C | 0.527 | 1.77E-05 |
| Nsp5 | EIF4E | 0.527 | 1.83E-05 |
| Nsp5 | CD86 | 0.527 | 1.84E-05 |
| Nsp5 | PTPN13 | 0.527 | 1.77E-05 |
| Nsp5 | PRKACB | 0.527 | 1.77E-05 |
| Nsp13 | MAP3K4 | 0.527 | 1.83E-05 |
| Nsp15 | ITGA2 | 0.527 | 1.81E-05 |
| Nsp16 | PPP2R2A | 0.527 | 1.81E-05 |

| | | | |
|---|---|---|---|
| Nsp16 | MET | 0.527 | 1.83E-05 |
| S | CHUK | 0.527 | 1.81E-05 |
| N | LIFR | 0.527 | 1.79E-05 |
| Orf3a | PRKAA1 | 0.527 | 1.83E-05 |
| Orf7a | IFNGR1 | 0.527 | 1.77E-05 |
| Nsp2 | SOS1 | 0.526 | 1.86E-05 |
| Nsp3 | IL22RA2 | 0.526 | 1.90E-05 |
| Nsp3 | RAPGEF2 | 0.526 | 1.85E-05 |
| Nsp4 | IFIH1 | 0.526 | 1.85E-05 |
| Nsp6 | ITGA6 | 0.526 | 1.86E-05 |
| Nsp8 | SEH1L | 0.526 | 1.87E-05 |
| Nsp12 | GHR | 0.526 | 1.86E-05 |
| Nsp12 | PPP2CB | 0.526 | 1.91E-05 |
| Nsp14 | PPP2R2A | 0.526 | 1.85E-05 |
| Nsp14 | IL23R | 0.526 | 1.87E-05 |
| Nsp15 | IFIH1 | 0.526 | 1.92E-05 |
| Nsp15 | PPP2R1B | 0.526 | 1.90E-05 |
| N | DNM1L | 0.526 | 1.86E-05 |
| N | IL18R1 | 0.526 | 1.91E-05 |
| Orf3a | FZD3 | 0.526 | 1.90E-05 |
| Orf3a | COL4A4 | 0.526 | 1.90E-05 |
| Orf7a | MECOM | 0.526 | 1.91E-05 |
| Nsp1 | COL1A2 | 0.525 | 2.00E-05 |
| Nsp3 | MECOM | 0.525 | 1.95E-05 |
| Nsp3 | SEH1L | 0.525 | 1.93E-05 |
| Nsp4 | DDX3X | 0.525 | 2.00E-05 |
| Nsp5 | PRKAA1 | 0.525 | 1.94E-05 |
| Nsp9 | MECOM | 0.525 | 1.99E-05 |
| Nsp14 | PPP2CA | 0.525 | 1.95E-05 |
| Nsp15 | FZD3 | 0.525 | 2.00E-05 |
| Nsp15 | RASA2 | 0.525 | 1.95E-05 |
| S | GYS2 | 0.525 | 1.96E-05 |
| S | CD86 | 0.525 | 1.93E-05 |
| Orf3a | LDHC | 0.525 | 1.96E-05 |
| Orf3a | ITGB8 | 0.525 | 1.98E-05 |
| Orf3a | IL18R1 | 0.525 | 1.99E-05 |
| Nsp2 | ITGB1 | 0.524 | 2.05E-05 |
| Nsp2 | PTPN13 | 0.524 | 2.07E-05 |
| Nsp2 | CASP3 | 0.524 | 2.05E-05 |
| Nsp2 | PRKACB | 0.524 | 2.06E-05 |
| Nsp2 | BMPR2 | 0.524 | 2.02E-05 |
| Nsp2 | TAB3 | 0.524 | 2.05E-05 |
| Nsp3 | SMAD2 | 0.524 | 2.02E-05 |
| Nsp4 | IL23R | 0.524 | 2.07E-05 |
| Nsp5 | MAP3K7 | 0.524 | 2.07E-05 |
| Nsp8 | ACSL3 | 0.524 | 2.05E-05 |
| N | FZD3 | 0.524 | 2.03E-05 |
| Nsp2 | HGF | 0.523 | 2.17E-05 |
| Nsp6 | E2F5 | 0.523 | 2.10E-05 |
| Nsp6 | COL4A3 | 0.523 | 2.15E-05 |

| | | | |
|---|---|---|---|
| Nsp8 | ATG5 | 0.523 | 2.16E-05 |
| Nsp8 | PPP2CA | 0.523 | 2.10E-05 |
| Nsp9 | RASA1 | 0.523 | 2.17E-05 |
| Nsp12 | IL23R | 0.523 | 2.15E-05 |
| Nsp12 | STAT4 | 0.523 | 2.13E-05 |
| Nsp13 | SMAD2 | 0.523 | 2.16E-05 |
| Nsp15 | HSP90B1 | 0.523 | 2.17E-05 |
| S | TLR1 | 0.523 | 2.14E-05 |
| S | HGF | 0.523 | 2.16E-05 |
| M | ATG5 | 0.523 | 2.18E-05 |
| M | DNM1L | 0.523 | 2.10E-05 |
| N | CAMK2D | 0.523 | 2.16E-05 |
| Orf8 | PIK3CA | 0.523 | 2.18E-05 |
| Nsp3 | ITGA6 | 0.522 | 2.25E-05 |
| Nsp5 | AHR | 0.522 | 2.22E-05 |
| Nsp9 | LEPR | 0.522 | 2.21E-05 |
| Nsp12 | ITGB1 | 0.522 | 2.20E-05 |
| Nsp12 | IL33 | 0.522 | 2.19E-05 |
| Nsp14 | MAP3K2 | 0.522 | 2.23E-05 |
| Nsp15 | DDX3X | 0.522 | 2.19E-05 |
| S | PDGFC | 0.522 | 2.22E-05 |
| N | LRP6 | 0.522 | 2.24E-05 |
| N | MET | 0.522 | 2.24E-05 |
| Nsp7 | MDM2 | 0.521 | 2.38E-05 |
| Nsp9 | GNB4 | 0.521 | 2.32E-05 |
| Nsp12 | CAB39 | 0.521 | 2.37E-05 |
| Nsp12 | PPP2R1B | 0.521 | 2.37E-05 |
| Nsp14 | TNFSF10 | 0.521 | 2.35E-05 |
| Nsp15 | DDX58 | 0.521 | 2.31E-05 |
| Nsp16 | PPP2R5E | 0.521 | 2.30E-05 |
| N | ATP6V1A | 0.521 | 2.29E-05 |
| Nsp4 | MAP3K2 | 0.52 | 2.46E-05 |
| Nsp5 | FZD6 | 0.52 | 2.40E-05 |
| Nsp5 | IL1R1 | 0.52 | 2.45E-05 |
| Nsp6 | MECOM | 0.52 | 2.42E-05 |
| Nsp10 | LDHAL6A | 0.52 | 2.46E-05 |
| Nsp13 | BCL10 | 0.52 | 2.48E-05 |
| Nsp15 | COL4A3 | 0.52 | 2.46E-05 |
| S | PLCB4 | 0.52 | 2.45E-05 |
| N | COL9A1 | 0.52 | 2.44E-05 |
| Orf7a | TLR3 | 0.52 | 2.45E-05 |
| Nsp2 | ROCK2 | 0.519 | 2.49E-05 |
| Nsp2 | IFNGR1 | 0.519 | 2.50E-05 |
| Nsp2 | PLA2G4A | 0.519 | 2.57E-05 |
| Nsp4 | EIF4E | 0.519 | 2.54E-05 |
| Nsp4 | NFATC3 | 0.519 | 2.55E-05 |
| Nsp6 | IL13RA2 | 0.519 | 2.52E-05 |
| Nsp6 | MAP3K2 | 0.519 | 2.51E-05 |
| Nsp9 | STK3 | 0.519 | 2.50E-05 |
| Nsp13 | FAS | 0.519 | 2.50E-05 |

| | | | |
|---|---|---|---|
| S | SMAD2 | 0.519 | 2.53E-05 |
| Orf7a | PPP2R3A | 0.519 | 2.58E-05 |
| Nsp3 | ACSL4 | 0.518 | 2.63E-05 |
| Nsp4 | PPP2R3A | 0.518 | 2.60E-05 |
| Nsp5 | LPAR6 | 0.518 | 2.62E-05 |
| Nsp9 | EIF2S1 | 0.518 | 2.69E-05 |
| Nsp12 | ACSL3 | 0.518 | 2.69E-05 |
| Nsp12 | MAP3K1 | 0.518 | 2.64E-05 |
| Nsp12 | RBL2 | 0.518 | 2.64E-05 |
| Nsp12 | PDGFC | 0.518 | 2.59E-05 |
| Nsp12 | TLR8 | 0.518 | 2.70E-05 |
| Nsp16 | CALM2 | 0.518 | 2.65E-05 |
| Nsp16 | CD86 | 0.518 | 2.60E-05 |
| Orf3a | TBK1 | 0.518 | 2.70E-05 |
| Orf3a | PDK1 | 0.518 | 2.66E-05 |
| Orf3a | SMAD4 | 0.518 | 2.63E-05 |
| Orf3a | CYLD | 0.518 | 2.65E-05 |
| Nsp2 | EGF | 0.517 | 2.73E-05 |
| Nsp4 | CDC42 | 0.517 | 2.79E-05 |
| Nsp5 | RICTOR | 0.517 | 2.76E-05 |
| Nsp6 | COL4A6 | 0.517 | 2.74E-05 |
| Nsp6 | SMURF2 | 0.517 | 2.77E-05 |
| Nsp6 | BRAF | 0.517 | 2.79E-05 |
| Nsp6 | HSP90AA1 | 0.517 | 2.75E-05 |
| Nsp7 | IL6ST | 0.517 | 2.82E-05 |
| Nsp8 | FZD3 | 0.517 | 2.72E-05 |
| Nsp16 | ACSL4 | 0.517 | 2.80E-05 |
| S | MAP3K20 | 0.517 | 2.76E-05 |
| N | ATP6V1C1 | 0.517 | 2.71E-05 |
| N | GRK3 | 0.517 | 2.72E-05 |
| Orf3a | ITGB1 | 0.517 | 2.71E-05 |
| Orf7a | MAP2K4 | 0.517 | 2.79E-05 |
| Orf8 | ITGB1 | 0.517 | 2.77E-05 |
| Nsp2 | PPM1A | 0.516 | 2.82E-05 |
| Nsp2 | LDHAL6A | 0.516 | 2.92E-05 |
| Nsp5 | ACSL3 | 0.516 | 2.84E-05 |
| Nsp5 | IL18 | 0.516 | 2.90E-05 |
| Nsp6 | IFIH1 | 0.516 | 2.88E-05 |
| Nsp6 | ELK4 | 0.516 | 2.86E-05 |
| Nsp7 | IL18 | 0.516 | 2.82E-05 |
| Nsp8 | SOCS5 | 0.516 | 2.82E-05 |
| Nsp9 | STAM | 0.516 | 2.85E-05 |
| Nsp10 | SOS1 | 0.516 | 2.94E-05 |
| Nsp14 | TLR8 | 0.516 | 2.90E-05 |
| S | PPP2CB | 0.516 | 2.91E-05 |
| N | BIRC2 | 0.516 | 2.84E-05 |
| N | IL6ST | 0.516 | 2.88E-05 |
| Nsp3 | SMAD5 | 0.515 | 2.95E-05 |
| Nsp3 | EIF2AK3 | 0.515 | 2.94E-05 |
| Nsp3 | DDX3X | 0.515 | 3.04E-05 |

| | | | |
|---|---|---|---|
| Nsp6 | ITGB1 | 0.515 | 2.94E-05 |
| Nsp7 | ACVR2A | 0.515 | 2.96E-05 |
| Nsp7 | STAM2 | 0.515 | 2.99E-05 |
| Nsp8 | PARP4 | 0.515 | 3.05E-05 |
| Nsp9 | ACSL4 | 0.515 | 3.06E-05 |
| Nsp13 | TANK | 0.515 | 3.03E-05 |
| Nsp14 | PPP2R5E | 0.515 | 3.06E-05 |
| Nsp15 | MAP3K1 | 0.515 | 2.95E-05 |
| Orf3a | CAB39 | 0.515 | 2.99E-05 |
| Orf3a | PPP2R5E | 0.515 | 3.05E-05 |
| Orf8 | CYLD | 0.515 | 3.06E-05 |
| Nsp3 | MAP3K4 | 0.514 | 3.17E-05 |
| Nsp4 | MMP13 | 0.514 | 3.12E-05 |
| Nsp4 | MAPK9 | 0.514 | 3.07E-05 |
| Nsp5 | PPP2R3A | 0.514 | 3.16E-05 |
| Nsp6 | MAP3K4 | 0.514 | 3.09E-05 |
| Nsp8 | MAP3K1 | 0.514 | 3.17E-05 |
| Nsp8 | PPP2R5E | 0.514 | 3.16E-05 |
| Nsp8 | RASA2 | 0.514 | 3.15E-05 |
| Nsp9 | PIK3CA | 0.514 | 3.16E-05 |
| S | IL5RA | 0.514 | 3.15E-05 |
| N | GNB4 | 0.514 | 3.07E-05 |
| Orf3a | ITGA1 | 0.514 | 3.18E-05 |
| Orf3a | PIK3CB | 0.514 | 3.13E-05 |
| Orf7a | IFIH1 | 0.514 | 3.18E-05 |
| Orf7a | NLK | 0.514 | 3.18E-05 |
| Orf8 | BIRC2 | 0.514 | 3.16E-05 |
| Orf8 | ATP6V1A | 0.514 | 3.18E-05 |
| Nsp2 | GHR | 0.513 | 3.24E-05 |
| Nsp2 | ITGA1 | 0.513 | 3.20E-05 |
| Nsp3 | NLK | 0.513 | 3.27E-05 |
| Nsp5 | SMAD4 | 0.513 | 3.29E-05 |
| Nsp8 | CCNE2 | 0.513 | 3.27E-05 |
| Nsp8 | CAB39 | 0.513 | 3.22E-05 |
| Nsp8 | IL18R1 | 0.513 | 3.26E-05 |
| Nsp9 | SOCS5 | 0.513 | 3.31E-05 |
| Nsp12 | MALT1 | 0.513 | 3.23E-05 |
| Nsp12 | PPP2R3A | 0.513 | 3.32E-05 |
| Nsp14 | IL33 | 0.513 | 3.32E-05 |
| Nsp16 | MECOM | 0.513 | 3.27E-05 |
| Nsp16 | VAV3 | 0.513 | 3.32E-05 |
| S | ATG5 | 0.513 | 3.20E-05 |
| S | VAV3 | 0.513 | 3.26E-05 |
| S | HSP90AA1 | 0.513 | 3.30E-05 |
| Orf3a | AZI2 | 0.513 | 3.26E-05 |
| Nsp3 | TANK | 0.512 | 3.40E-05 |
| Nsp6 | MAP3K7 | 0.512 | 3.36E-05 |
| Nsp14 | MAP3K20 | 0.512 | 3.41E-05 |
| Nsp15 | ATG5 | 0.512 | 3.34E-05 |
| S | LAMB4 | 0.512 | 3.45E-05 |

| | | | |
|---|---|---|---|
| S | LRP6 | 0.512 | 3.46E-05 |
| Orf3a | MMP13 | 0.512 | 3.44E-05 |
| Orf7a | CXCL10 | 0.512 | 3.43E-05 |
| Nsp2 | COL4A3 | 0.511 | 3.60E-05 |
| Nsp2 | BRCA1 | 0.511 | 3.49E-05 |
| Nsp5 | PPP2CB | 0.511 | 3.49E-05 |
| Nsp5 | PPP2R1B | 0.511 | 3.51E-05 |
| Nsp5 | ZFYVE9 | 0.511 | 3.60E-05 |
| Nsp6 | PIK3CA | 0.511 | 3.60E-05 |
| Nsp13 | AREG | 0.511 | 3.59E-05 |
| Nsp13 | ATP6V1A | 0.511 | 3.52E-05 |
| Nsp13 | PPP2R1B | 0.511 | 3.53E-05 |
| Nsp14 | STRADB | 0.511 | 3.59E-05 |
| Nsp15 | ANGPT1 | 0.511 | 3.57E-05 |
| Nsp15 | COL9A1 | 0.511 | 3.49E-05 |
| S | GNAI1 | 0.511 | 3.51E-05 |
| Nsp2 | FAS | 0.51 | 3.68E-05 |
| Nsp4 | ACSL3 | 0.51 | 3.62E-05 |
| Nsp16 | NLK | 0.51 | 3.75E-05 |
| Nsp16 | ACSL3 | 0.51 | 3.71E-05 |
| Orf7a | RPS6KB1 | 0.51 | 3.68E-05 |
| Orf7a | ZFYVE9 | 0.51 | 3.75E-05 |
| Orf8 | PPP2R5E | 0.51 | 3.66E-05 |
| Nsp5 | CREB1 | 0.509 | 3.84E-05 |
| Nsp13 | COL4A6 | 0.509 | 3.87E-05 |
| Nsp13 | RAPGEF2 | 0.509 | 3.89E-05 |
| Nsp14 | PIK3CB | 0.509 | 3.92E-05 |
| Orf7a | SOCS5 | 0.509 | 3.78E-05 |
| Orf8 | IL7 | 0.509 | 3.91E-05 |
| Orf8 | SGK3 | 0.509 | 3.77E-05 |
| Nsp5 | SOCS4 | 0.508 | 4.08E-05 |
| Nsp5 | PTPN2 | 0.508 | 4.04E-05 |
| Nsp15 | NF1 | 0.508 | 3.94E-05 |
| Nsp16 | AREG | 0.508 | 4.03E-05 |
| S | TLR7 | 0.508 | 4.02E-05 |
| S | BMPR1B | 0.508 | 4.01E-05 |
| S | DDX3X | 0.508 | 4.03E-05 |
| N | PPM1A | 0.508 | 4.00E-05 |
| Orf7a | TNFSF18 | 0.508 | 4.08E-05 |
| Orf8 | APAF1 | 0.508 | 4.01E-05 |
| Nsp2 | LAMA2 | 0.507 | 4.20E-05 |
| Nsp4 | NLK | 0.507 | 4.17E-05 |
| Nsp6 | RBL1 | 0.507 | 4.16E-05 |
| Nsp12 | COL4A6 | 0.507 | 4.25E-05 |
| Nsp14 | PDGFC | 0.507 | 4.19E-05 |
| Nsp14 | TGFBR1 | 0.507 | 4.20E-05 |
| Nsp15 | FAS | 0.507 | 4.16E-05 |
| Nsp16 | FZD3 | 0.507 | 4.09E-05 |
| M | MAPK6 | 0.507 | 4.17E-05 |
| Orf7a | CAMK2D | 0.507 | 4.22E-05 |

| | | | |
|---|---|---|---|
| Orf8 | TAB2 | 0.507 | 4.19E-05 |
| Orf8 | STAM | 0.507 | 4.16E-05 |
| Orf8 | RBL1 | 0.507 | 4.18E-05 |
| Nsp2 | IL18 | 0.506 | 4.37E-05 |
| Nsp4 | BCL10 | 0.506 | 4.32E-05 |
| Nsp12 | KRAS | 0.506 | 4.39E-05 |
| Nsp15 | STRADB | 0.506 | 4.43E-05 |
| Nsp16 | LDHC | 0.506 | 4.41E-05 |
| S | IL22RA2 | 0.506 | 4.27E-05 |
| N | MAP3K1 | 0.506 | 4.41E-05 |
| N | APAF1 | 0.506 | 4.38E-05 |
| Nsp2 | ITGB8 | 0.505 | 4.51E-05 |
| Nsp6 | TLR4 | 0.505 | 4.44E-05 |
| Nsp9 | SEH1L | 0.505 | 4.56E-05 |
| Nsp14 | BCL10 | 0.505 | 4.54E-05 |
| Nsp15 | AZI2 | 0.505 | 4.55E-05 |
| N | LAMTOR3 | 0.505 | 4.53E-05 |
| Orf6 | RPS6KA6 | 0.505 | 4.54E-05 |
| Orf8 | ACSL3 | 0.505 | 4.60E-05 |
| Orf8 | AKT3 | 0.505 | 4.47E-05 |
| Nsp8 | ITGB8 | 0.504 | 4.71E-05 |
| Nsp8 | ACSL4 | 0.504 | 4.68E-05 |
| Nsp9 | PPP2R5E | 0.504 | 4.68E-05 |
| Nsp9 | COL1A2 | 0.504 | 4.63E-05 |
| Nsp12 | DDX58 | 0.504 | 4.62E-05 |
| Nsp14 | MET | 0.504 | 4.63E-05 |
| Nsp15 | BCL10 | 0.504 | 4.67E-05 |
| N | ITGB8 | 0.504 | 4.69E-05 |
| N | RPS6 | 0.504 | 4.77E-05 |
| N | MAP3K2 | 0.504 | 4.72E-05 |
| Nsp4 | STRADB | 0.503 | 4.97E-05 |
| Nsp6 | PPP2R2A | 0.503 | 4.90E-05 |
| Nsp8 | PPP2R3A | 0.503 | 4.83E-05 |
| Nsp9 | USP25 | 0.503 | 4.92E-05 |
| Nsp13 | ITGA6 | 0.503 | 4.84E-05 |
| Nsp13 | IL22RA2 | 0.503 | 4.81E-05 |
| Nsp15 | SMAD5 | 0.503 | 4.81E-05 |
| Nsp16 | COL1A2 | 0.503 | 4.83E-05 |
| Orf3a | PPP3CB | 0.503 | 4.90E-05 |
| Nsp3 | TNFSF18 | 0.502 | 5.18E-05 |
| Nsp4 | PPP2R5E | 0.502 | 5.17E-05 |
| Nsp4 | TLR8 | 0.502 | 5.18E-05 |
| Nsp6 | MAPK9 | 0.502 | 5.12E-05 |
| Nsp9 | STAM2 | 0.502 | 5.20E-05 |
| Nsp13 | IL33 | 0.502 | 5.17E-05 |
| S | CASP3 | 0.502 | 5.20E-05 |
| N | PPP3CA | 0.502 | 5.14E-05 |
| N | PDK1 | 0.502 | 5.17E-05 |
| Orf7b | MAPK9 | 0.502 | 5.14E-05 |
| Nsp2 | PHKB | 0.501 | 5.38E-05 |

| | | | |
|---|---|---|---|
| Nsp2 | SMURF2 | 0.501 | 5.21E-05 |
| Nsp4 | MALT1 | 0.501 | 5.22E-05 |
| Nsp4 | LPAR6 | 0.501 | 5.24E-05 |
| Nsp8 | LRP6 | 0.501 | 5.28E-05 |
| Nsp9 | FZD3 | 0.501 | 5.40E-05 |
| Nsp9 | RBL1 | 0.501 | 5.28E-05 |
| Nsp10 | TBK1 | 0.501 | 5.36E-05 |
| Nsp12 | DDX3X | 0.501 | 5.31E-05 |
| Nsp14 | IFIH1 | 0.501 | 5.33E-05 |
| Nsp14 | CXCL10 | 0.501 | 5.23E-05 |
| Nsp14 | BMP5 | 0.501 | 5.35E-05 |
| Nsp16 | GNAI1 | 0.501 | 5.22E-05 |
| Nsp16 | EIF2AK3 | 0.501 | 5.24E-05 |
| S | TICAM2 | 0.501 | 5.28E-05 |
| S | ACVR1C | 0.501 | 5.29E-05 |
| N | ACSL3 | 0.501 | 5.26E-05 |
| Orf6 | NFATC3 | 0.501 | 5.41E-05 |
| Orf7a | EIF4E | 0.501 | 5.32E-05 |
| Orf7a | ACKR4 | 0.501 | 5.24E-05 |
| Orf8 | ZFYVE9 | 0.501 | 5.30E-05 |
| Orf8 | SMAD4 | 0.501 | 5.29E-05 |
| Nsp2 | PRKAA1 | 0.5 | 5.54E-05 |
| Nsp2 | PTPN2 | 0.5 | 5.48E-05 |
| Nsp13 | CHUK | 0.5 | 5.54E-05 |
| Nsp13 | LRP6 | 0.5 | 5.60E-05 |
| Nsp16 | STRADB | 0.5 | 5.62E-05 |
| Orf7a | RBL2 | 0.5 | 5.49E-05 |
| Nsp2 | SOCS4 | 0.499 | 5.67E-05 |
| Nsp5 | FZD3 | 0.499 | 5.79E-05 |
| Nsp5 | PLA2G4A | 0.499 | 5.77E-05 |
| Nsp6 | BMPR1B | 0.499 | 5.83E-05 |
| Nsp6 | TLR8 | 0.499 | 5.82E-05 |
| Nsp8 | CHUK | 0.499 | 5.66E-05 |
| Nsp10 | MIOS | 0.499 | 5.72E-05 |
| Nsp10 | COL4A5 | 0.499 | 5.66E-05 |
| Nsp12 | FZD3 | 0.499 | 5.65E-05 |
| Nsp12 | FAS | 0.499 | 5.64E-05 |
| Nsp15 | ITGAV | 0.499 | 5.73E-05 |
| E | SKP1 | 0.499 | 5.82E-05 |
| N | RELN | 0.499 | 5.68E-05 |
| Orf3a | IL23R | 0.499 | 5.70E-05 |
| Orf3a | IL22RA2 | 0.499 | 5.81E-05 |
| Orf7a | ITGA6 | 0.499 | 5.66E-05 |
| Orf7a | COL4A1 | 0.499 | 5.74E-05 |
| Nsp8 | MAP3K2 | 0.498 | 6.04E-05 |
| Nsp9 | APAF1 | 0.498 | 6.09E-05 |
| Nsp9 | COL9A1 | 0.498 | 5.99E-05 |
| Nsp13 | RBL2 | 0.498 | 5.89E-05 |
| Nsp16 | MAPK9 | 0.498 | 6.04E-05 |
| S | BCL10 | 0.498 | 6.06E-05 |

| | | | |
|---|---|---|---|
| S | TLR4 | 0.498 | 5.94E-05 |
| N | YWHAZ | 0.498 | 6.05E-05 |
| N | ITCH | 0.498 | 6.08E-05 |
| Nsp2 | FZD3 | 0.497 | 6.20E-05 |
| Nsp2 | CDC42 | 0.497 | 6.31E-05 |
| Nsp4 | LPAR4 | 0.497 | 6.34E-05 |
| Nsp5 | TFRC | 0.497 | 6.11E-05 |
| Nsp12 | GNAI1 | 0.497 | 6.13E-05 |
| Nsp12 | IL22RA2 | 0.497 | 6.14E-05 |
| Nsp13 | EIF2AK3 | 0.497 | 6.28E-05 |
| Nsp13 | BRAF | 0.497 | 6.32E-05 |
| Nsp14 | SMAD4 | 0.497 | 6.31E-05 |
| M | ATF2 | 0.497 | 6.22E-05 |
| Orf3a | SORBS1 | 0.497 | 6.24E-05 |
| Nsp2 | TLR8 | 0.496 | 6.50E-05 |
| Nsp3 | LRP6 | 0.496 | 6.50E-05 |
| Nsp5 | CXCL10 | 0.496 | 6.45E-05 |
| Nsp5 | ACSL4 | 0.496 | 6.41E-05 |
| Nsp8 | PRKAA2 | 0.496 | 6.53E-05 |
| Nsp14 | ITGB1 | 0.496 | 6.46E-05 |
| Orf3a | MMP3 | 0.496 | 6.59E-05 |
| Orf7a | IL33 | 0.496 | 6.40E-05 |
| Orf8 | COL9A1 | 0.496 | 6.45E-05 |
| Nsp4 | NF1 | 0.495 | 6.82E-05 |
| Nsp5 | ITGA2 | 0.495 | 6.74E-05 |
| Nsp5 | CDC42 | 0.495 | 6.84E-05 |
| Nsp8 | OSMR | 0.495 | 6.86E-05 |
| Nsp9 | MAP4K3 | 0.495 | 6.82E-05 |
| Nsp14 | TLR7 | 0.495 | 6.72E-05 |
| Nsp15 | CXCL10 | 0.495 | 6.80E-05 |
| Nsp15 | PARP4 | 0.495 | 6.86E-05 |
| Orf3a | HGF | 0.495 | 6.67E-05 |
| Orf8 | SOCS5 | 0.495 | 6.82E-05 |
| Orf8 | TLR6 | 0.495 | 6.71E-05 |
| Orf8 | PPM1A | 0.495 | 6.82E-05 |
| Orf8 | TAOK1 | 0.495 | 6.84E-05 |
| Nsp4 | TNFSF18 | 0.494 | 6.88E-05 |
| Nsp5 | PPM1A | 0.494 | 7.09E-05 |
| Nsp6 | KIT | 0.494 | 6.97E-05 |
| Nsp7 | PPM1B | 0.494 | 7.13E-05 |
| Nsp9 | ATP6V1A | 0.494 | 6.99E-05 |
| Nsp15 | MAPK9 | 0.494 | 6.95E-05 |
| S | PIK3R1 | 0.494 | 7.01E-05 |
| Orf7a | PDGFC | 0.494 | 7.06E-05 |
| Orf8 | TFRC | 0.494 | 7.07E-05 |
| Orf8 | PIK3CB | 0.494 | 6.96E-05 |
| Nsp2 | SMAD5 | 0.493 | 7.42E-05 |
| Nsp4 | MET | 0.493 | 7.21E-05 |
| Nsp4 | PPP2CB | 0.493 | 7.38E-05 |
| Nsp6 | LRP6 | 0.493 | 7.20E-05 |

| | | | |
|---|---|---|---|
| Nsp6 | PDGFC | 0.493 | 7.37E-05 |
| Nsp12 | SMURF2 | 0.493 | 7.32E-05 |
| Nsp14 | ITGAV | 0.493 | 7.39E-05 |
| M | PPP2CA | 0.493 | 7.19E-05 |
| N | ATP6V1H | 0.493 | 7.17E-05 |
| Orf8 | IFNGR1 | 0.493 | 7.27E-05 |
| Nsp3 | ATP6V1C1 | 0.492 | 7.69E-05 |
| Nsp4 | IL1R1 | 0.492 | 7.71E-05 |
| Nsp5 | MSTN | 0.492 | 7.43E-05 |
| Nsp6 | GDF9 | 0.492 | 7.53E-05 |
| Nsp9 | SGK3 | 0.492 | 7.57E-05 |
| Nsp9 | LPAR6 | 0.492 | 7.67E-05 |
| Nsp12 | STRADB | 0.492 | 7.54E-05 |
| Nsp14 | MECOM | 0.492 | 7.61E-05 |
| Nsp14 | NFATC3 | 0.492 | 7.61E-05 |
| M | SLC38A9 | 0.492 | 7.65E-05 |
| Orf3a | RBL1 | 0.492 | 7.50E-05 |
| Orf7a | MAP3K5 | 0.492 | 7.63E-05 |
| Orf8 | CAMK2D | 0.492 | 7.71E-05 |
| Orf8 | SOCS4 | 0.492 | 7.54E-05 |
| Nsp2 | AKT3 | 0.491 | 7.90E-05 |
| Nsp2 | RPS6 | 0.491 | 7.93E-05 |
| Nsp4 | COL4A6 | 0.491 | 7.96E-05 |
| Nsp6 | TNFSF18 | 0.491 | 7.80E-05 |
| Nsp6 | LAMA2 | 0.491 | 7.95E-05 |
| Nsp9 | TAOK1 | 0.491 | 7.96E-05 |
| Nsp10 | ROCK1 | 0.491 | 7.83E-05 |
| Nsp14 | IL13RA1 | 0.491 | 7.87E-05 |
| Nsp16 | SEH1L | 0.491 | 7.96E-05 |
| S | LDHAL6A | 0.491 | 7.77E-05 |
| N | LDHC | 0.491 | 7.74E-05 |
| N | PIK3CB | 0.491 | 7.72E-05 |
| Orf7a | PRKAA2 | 0.491 | 7.87E-05 |
| Orf7a | MMP13 | 0.491 | 7.93E-05 |
| Orf7a | PPM1A | 0.491 | 8.02E-05 |
| Orf8 | RASA1 | 0.491 | 7.74E-05 |
| Nsp3 | MAPK8 | 0.49 | 8.07E-05 |
| Nsp4 | VCAM1 | 0.49 | 8.28E-05 |
| Nsp4 | COL4A4 | 0.49 | 8.13E-05 |
| Nsp6 | MALT1 | 0.49 | 8.29E-05 |
| Nsp15 | EIF2AK3 | 0.49 | 8.05E-05 |
| S | CREB1 | 0.49 | 8.32E-05 |
| S | SORBS1 | 0.49 | 8.12E-05 |
| Nsp3 | PAK2 | 0.489 | 8.54E-05 |
| Nsp3 | HGF | 0.489 | 8.34E-05 |
| Nsp4 | BMPR2 | 0.489 | 8.49E-05 |
| Nsp9 | PPP2R1B | 0.489 | 8.59E-05 |
| Orf3a | PPP2R3A | 0.489 | 8.44E-05 |
| Orf8 | SEH1L | 0.489 | 8.44E-05 |
| Nsp3 | PPP2R2A | 0.488 | 8.74E-05 |

| | | | |
|---|---|---|---|
| Nsp4 | TLR4 | 0.488 | 8.78E-05 |
| Nsp4 | MAP3K4 | 0.488 | 8.87E-05 |
| Nsp4 | ACSL4 | 0.488 | 8.73E-05 |
| Nsp6 | SMAD4 | 0.488 | 8.84E-05 |
| Nsp8 | MSTN | 0.488 | 8.79E-05 |
| Nsp9 | HIF1A | 0.488 | 8.99E-05 |
| Nsp9 | SMAD2 | 0.488 | 8.78E-05 |
| Nsp10 | PKN2 | 0.488 | 8.77E-05 |
| Nsp15 | LDHAL6A | 0.488 | 8.70E-05 |
| S | PRKAR1A | 0.488 | 8.88E-05 |
| S | PPP2R2A | 0.488 | 8.99E-05 |
| M | PKN2 | 0.488 | 8.81E-05 |
| Orf3a | RICTOR | 0.488 | 8.92E-05 |
| Orf7a | GHR | 0.488 | 8.90E-05 |
| Orf7a | IL7 | 0.488 | 8.83E-05 |
| Orf7a | ATF2 | 0.488 | 8.86E-05 |
| Orf7a | MAP3K20 | 0.488 | 8.78E-05 |
| Orf8 | MAP3K1 | 0.488 | 8.89E-05 |
| Nsp5 | PRKAA2 | 0.487 | 9.22E-05 |
| Nsp6 | EIF2AK3 | 0.487 | 9.15E-05 |
| Nsp9 | ACVR2A | 0.487 | 9.35E-05 |
| Nsp9 | PPP3R1 | 0.487 | 9.06E-05 |
| Nsp12 | COL9A1 | 0.487 | 9.35E-05 |
| Nsp14 | DDX58 | 0.487 | 9.30E-05 |
| Nsp16 | RAPGEF2 | 0.487 | 9.16E-05 |
| Nsp16 | MAP3K2 | 0.487 | 9.25E-05 |
| M | IRAK4 | 0.487 | 9.34E-05 |
| M | LDHAL6A | 0.487 | 9.25E-05 |
| Orf3a | GDF9 | 0.487 | 9.19E-05 |
| Orf3a | NFATC3 | 0.487 | 9.24E-05 |
| Orf7a | STRADB | 0.487 | 9.19E-05 |
| Orf8 | ITGA2 | 0.487 | 9.24E-05 |
| Nsp2 | TLR3 | 0.486 | 9.41E-05 |
| Nsp3 | LAMA2 | 0.486 | 9.55E-05 |
| Nsp5 | EGF | 0.486 | 9.47E-05 |
| Nsp6 | PAK2 | 0.486 | 9.58E-05 |
| Nsp6 | DDX58 | 0.486 | 9.36E-05 |
| Nsp6 | RAPGEF2 | 0.486 | 9.71E-05 |
| Nsp8 | EGF | 0.486 | 9.57E-05 |
| Nsp8 | LDHAL6A | 0.486 | 9.41E-05 |
| Nsp9 | STAT4 | 0.486 | 9.52E-05 |
| Nsp13 | CXCL13 | 0.486 | 9.55E-05 |
| Nsp14 | ACSL4 | 0.486 | 9.58E-05 |
| Nsp16 | TANK | 0.486 | 9.44E-05 |
| Nsp16 | EGF | 0.486 | 9.58E-05 |
| S | BRAF | 0.486 | 9.61E-05 |
| M | MIOS | 0.486 | 9.55E-05 |
| N | PRKAA2 | 0.486 | 9.57E-05 |
| Orf8 | IFNAR1 | 0.486 | 9.47E-05 |
| Orf8 | PLA2G4A | 0.486 | 9.48E-05 |

| | | | |
|---|---|---|---|
| Nsp5 | CAMK2D | 0.485 | 9.91E-05 |
| Nsp5 | TAOK1 | 0.485 | 9.80E-05 |
| Nsp10 | SOS2 | 0.485 | 9.97E-05 |
| Nsp10 | RASA1 | 0.485 | 0.000100157 |
| Nsp12 | ATG5 | 0.485 | 9.98E-05 |
| Nsp14 | BMPR2 | 0.485 | 9.74E-05 |
| M | RASA1 | 0.485 | 0.000100892 |
| N | TLR6 | 0.485 | 9.87E-05 |
| N | RPS6KA6 | 0.485 | 9.78E-05 |
| Orf3a | DDX58 | 0.485 | 1.00E-04 |
| Orf3a | PRKACB | 0.485 | 0.000100243 |
| Orf7a | TLR7 | 0.485 | 9.89E-05 |
| Nsp2 | STAT4 | 0.484 | 0.000101835 |
| Nsp4 | CXCL13 | 0.484 | 0.000102737 |
| Nsp6 | PHKB | 0.484 | 0.000102367 |
| Nsp12 | EGF | 0.484 | 0.000102771 |
| Nsp14 | CHUK | 0.484 | 0.000100961 |
| Nsp15 | IL23R | 0.484 | 0.000101297 |
| M | STAM2 | 0.484 | 0.000102168 |
| M | FZD6 | 0.484 | 0.000102498 |
| N | SGK3 | 0.484 | 0.000103006 |
| Nsp3 | PPP2CB | 0.483 | 0.000107212 |
| Nsp4 | RAPGEF2 | 0.483 | 0.000105298 |
| Nsp6 | GNAI1 | 0.483 | 0.000107997 |
| Nsp8 | COL4A3 | 0.483 | 0.000105884 |
| Nsp13 | LPAR6 | 0.483 | 0.000108237 |
| Nsp15 | RPS6KB1 | 0.483 | 0.00010675 |
| Nsp15 | CHUK | 0.483 | 0.000108633 |
| E | COL1A2 | 0.483 | 0.000107769 |
| M | PPP2CB | 0.483 | 0.000107789 |
| N | PKN2 | 0.483 | 0.000105039 |
| N | LEPR | 0.483 | 0.00010756 |
| Orf3a | LRP6 | 0.483 | 0.000106249 |
| Orf7a | IL22RA2 | 0.483 | 0.000105216 |
| Nsp2 | SOCS5 | 0.482 | 0.00010967 |
| Nsp2 | JAK2 | 0.482 | 0.000111051 |
| Nsp2 | LDHA | 0.482 | 0.00011025 |
| Nsp2 | SMAD4 | 0.482 | 0.000111994 |
| Nsp4 | RPS6KB1 | 0.482 | 0.000110186 |
| Nsp8 | DDX58 | 0.482 | 0.000110641 |
| Nsp9 | PTPN2 | 0.482 | 0.000111465 |
| Nsp13 | CD86 | 0.482 | 0.000110848 |
| N | PPP3CB | 0.482 | 0.000108894 |
| N | SOS2 | 0.482 | 0.000110188 |
| N | ZFYVE16 | 0.482 | 0.000110389 |
| Orf3a | PDGFC | 0.482 | 0.000109699 |
| Nsp3 | TLR4 | 0.481 | 0.000115753 |
| Nsp12 | PHKB | 0.481 | 0.000115978 |
| Nsp12 | PPP2R5E | 0.481 | 0.000115143 |
| Nsp14 | CXCL13 | 0.481 | 0.000113271 |

| | | | |
|---|---|---|---|
| M | PIK3CA | 0.481 | 0.000115328 |
| M | MAP3K7 | 0.481 | 0.000117124 |
| N | PPP1R3A | 0.481 | 0.000114746 |
| Orf7a | RAP1A | 0.481 | 0.000113719 |
| Orf7a | FZD3 | 0.481 | 0.000114803 |
| Orf7a | RASA1 | 0.481 | 0.000116578 |
| Orf8 | STK3 | 0.481 | 0.000113448 |
| Nsp2 | MMP13 | 0.48 | 0.000118457 |
| Nsp4 | PDGFC | 0.48 | 0.000117956 |
| Nsp16 | ATP6V1C1 | 0.48 | 0.000120977 |
| M | ACSL3 | 0.48 | 0.000117968 |
| N | ZFYVE9 | 0.48 | 0.000120793 |
| Orf3a | ITGA4 | 0.48 | 0.000118076 |
| Orf7a | FLT1 | 0.48 | 0.000118058 |
| Orf8 | FZD3 | 0.48 | 0.000120947 |
| Nsp3 | BMP5 | 0.479 | 0.000122639 |
| Nsp5 | IL33 | 0.479 | 0.00012496 |
| Nsp9 | ATG5 | 0.479 | 0.000124182 |
| Nsp13 | RPS6KB1 | 0.479 | 0.000124326 |
| Nsp14 | NF1 | 0.479 | 0.000122078 |
| S | MMP3 | 0.479 | 0.000125755 |
| Orf3a | AREG | 0.479 | 0.000122027 |
| Orf3a | PLCB4 | 0.479 | 0.000123898 |
| Orf8 | TNFSF18 | 0.479 | 0.000123792 |
| Orf8 | SMURF2 | 0.479 | 0.000122154 |
| Nsp2 | BAG4 | 0.478 | 0.000128089 |
| Nsp5 | MECOM | 0.478 | 0.000129816 |
| Nsp6 | EGF | 0.478 | 0.000130807 |
| Nsp10 | LIFR | 0.478 | 0.000126859 |
| Nsp16 | TNFSF18 | 0.478 | 0.000130502 |
| Nsp16 | RAP1A | 0.478 | 0.000129641 |
| Nsp16 | E2F5 | 0.478 | 0.00013115 |
| Nsp16 | SMAD2 | 0.478 | 0.000127785 |
| S | RAP1A | 0.478 | 0.000130109 |
| S | GDF9 | 0.478 | 0.000130903 |
| S | LY96 | 0.478 | 0.000127856 |
| Orf7a | EIF2AK3 | 0.478 | 0.000129463 |
| Orf7a | BRAF | 0.478 | 0.000128867 |
| Orf8 | NFATC3 | 0.478 | 0.000129085 |
| Nsp2 | AHR | 0.477 | 0.000133841 |
| Nsp3 | TICAM2 | 0.477 | 0.000131666 |
| Nsp5 | LRP6 | 0.477 | 0.00013265 |
| Nsp8 | STRADB | 0.477 | 0.000132563 |
| Nsp15 | IL18 | 0.477 | 0.000135388 |
| Nsp15 | ATP6V1A | 0.477 | 0.000134177 |
| N | TANK | 0.477 | 0.000131907 |
| N | MMP1 | 0.477 | 0.000131542 |
| Orf3a | MAP3K20 | 0.477 | 0.000132828 |
| Orf3a | EIF2S1 | 0.477 | 0.000132104 |
| Orf7a | ACVR2A | 0.477 | 0.000134172 |

| | | | |
|---|---|---|---|
| Nsp3 | RPS6KB1 | 0.476 | 0.000141116 |
| Nsp5 | AKT3 | 0.476 | 0.000138049 |
| Nsp5 | COL4A3 | 0.476 | 0.000137358 |
| Nsp6 | CXCL13 | 0.476 | 0.000136368 |
| Nsp6 | BMP5 | 0.476 | 0.000139559 |
| Nsp6 | LPAR6 | 0.476 | 0.000139448 |
| Nsp8 | CALM2 | 0.476 | 0.000137592 |
| Nsp8 | PHKB | 0.476 | 0.000139086 |
| Nsp8 | BMPR1B | 0.476 | 0.000140687 |
| Nsp9 | CD86 | 0.476 | 0.000139356 |
| Nsp13 | LY96 | 0.476 | 0.000140953 |
| Nsp16 | PDGFC | 0.476 | 0.000136311 |
| S | TLR2 | 0.476 | 0.00013636 |
| N | PRKAR1A | 0.476 | 0.000139545 |
| Orf3a | MAP3K2 | 0.476 | 0.000140262 |
| Orf7a | HGF | 0.476 | 0.000138465 |
| Orf8 | EIF2S1 | 0.476 | 0.00013902 |
| Nsp7 | PKN2 | 0.475 | 0.000143392 |
| Nsp9 | MSTN | 0.475 | 0.000145802 |
| Orf3a | SOCS4 | 0.475 | 0.000141726 |
| Orf3a | GNAI1 | 0.475 | 0.000145473 |
| Orf3a | ACSL4 | 0.475 | 0.000145667 |
| Orf8 | JAK2 | 0.475 | 0.000141555 |
| Nsp2 | PTGS2 | 0.474 | 0.000150983 |
| Nsp3 | AREG | 0.474 | 0.000151274 |
| Nsp4 | SMAD2 | 0.474 | 0.000149036 |
| Nsp9 | RAPGEF2 | 0.474 | 0.000148545 |
| Nsp12 | ITGB8 | 0.474 | 0.000149126 |
| Nsp14 | CAMK2D | 0.474 | 0.000151741 |
| Nsp16 | ATG5 | 0.474 | 0.000148431 |
| Nsp16 | BMP5 | 0.474 | 0.000147915 |
| M | ATP6V1A | 0.474 | 0.000149384 |
| Orf3a | TLR5 | 0.474 | 0.000150678 |
| Orf7a | IL12RB2 | 0.474 | 0.000150502 |
| Orf7a | PPP2CA | 0.474 | 0.000149089 |
| Orf8 | CACNA2D1 | 0.474 | 0.000151763 |
| Nsp4 | GHR | 0.473 | 0.000155386 |
| Nsp5 | ATP6V1A | 0.473 | 0.000155059 |
| Nsp7 | BIRC2 | 0.473 | 0.000155592 |
| Nsp8 | VCAM1 | 0.473 | 0.000155132 |
| Nsp12 | SMAD4 | 0.473 | 0.000152775 |
| Nsp13 | PARP4 | 0.473 | 0.000153826 |
| Nsp13 | COL1A2 | 0.473 | 0.000152536 |
| Nsp15 | CALM2 | 0.473 | 0.000157648 |
| Nsp15 | TFRC | 0.473 | 0.000156774 |
| N | MAPK9 | 0.473 | 0.000152995 |
| N | PLA2G4A | 0.473 | 0.0001556 |
| Orf3a | LPAR4 | 0.473 | 0.000154246 |
| Orf7a | BMPR1B | 0.473 | 0.000152226 |
| Orf7a | RAPGEF2 | 0.473 | 0.000156937 |

| | | | |
|---|---|---|---|
| Orf7b | MDM2 | 0.473 | 0.000156396 |
| Nsp5 | RAPGEF2 | 0.472 | 0.000162608 |
| Nsp6 | BMPR2 | 0.472 | 0.000158197 |
| Nsp14 | PPP3R1 | 0.472 | 0.000163139 |
| Nsp15 | SMAD2 | 0.472 | 0.000157874 |
| Orf3a | PTGS2 | 0.472 | 0.0001583 |
| Orf7a | CXCL13 | 0.472 | 0.000162887 |
| Nsp2 | EIF4E | 0.471 | 0.000166577 |
| Nsp2 | CACNA2D1 | 0.471 | 0.000163836 |
| Nsp3 | BRAF | 0.471 | 0.00016693 |
| Nsp9 | IFIH1 | 0.471 | 0.000164976 |
| Nsp9 | TAB3 | 0.471 | 0.000169557 |
| Nsp9 | IFNGR1 | 0.471 | 0.000166947 |
| Nsp10 | PIAS2 | 0.471 | 0.000165656 |
| Nsp16 | ITGA6 | 0.471 | 0.000169579 |
| S | YWHAZ | 0.471 | 0.000166292 |
| N | IL36G | 0.471 | 0.000165829 |
| N | PPP2R1B | 0.471 | 0.000166172 |
| N | BIRC3 | 0.471 | 0.00016553 |
| Orf3a | SMAD2 | 0.471 | 0.000167073 |
| Orf6 | MAPK6 | 0.471 | 0.000166476 |
| Orf7a | NF1 | 0.471 | 0.000166227 |
| Nsp2 | STRADB | 0.47 | 0.000172484 |
| Nsp2 | ZFYVE9 | 0.47 | 0.000175205 |
| Nsp2 | IL33 | 0.47 | 0.000172322 |
| Nsp4 | PPP2R2A | 0.47 | 0.000174669 |
| Nsp5 | RBL2 | 0.47 | 0.000170301 |
| Nsp5 | EIF2AK3 | 0.47 | 0.000169713 |
| Nsp6 | TLR7 | 0.47 | 0.000171882 |
| Nsp8 | GHR | 0.47 | 0.000173071 |
| Nsp10 | STAM2 | 0.47 | 0.00017051 |
| Nsp13 | MAP3K20 | 0.47 | 0.000175584 |
| Nsp15 | CAB39 | 0.47 | 0.000173511 |
| Nsp16 | SMAD5 | 0.47 | 0.000172184 |
| S | KITLG | 0.47 | 0.000170753 |
| Orf8 | LPAR6 | 0.47 | 0.000175352 |
| Nsp6 | TLR1 | 0.469 | 0.000176453 |
| Nsp8 | RBL2 | 0.469 | 0.000176926 |
| Nsp8 | ITGA1 | 0.469 | 0.0001812 |
| Nsp12 | MET | 0.469 | 0.000181777 |
| Nsp12 | COL4A4 | 0.469 | 0.000176118 |
| Nsp13 | PDGFC | 0.469 | 0.000182058 |
| Nsp15 | IL18R1 | 0.469 | 0.000177527 |
| Nsp16 | PPP2R1B | 0.469 | 0.00017792 |
| S | PAK2 | 0.469 | 0.000179141 |
| S | BMP5 | 0.469 | 0.000181948 |
| N | COL4A3 | 0.469 | 0.000178176 |
| Orf8 | MSTN | 0.469 | 0.000180773 |
| Nsp3 | VAV3 | 0.468 | 0.000185071 |
| Nsp3 | MAP3K20 | 0.468 | 0.000183928 |

| | | | |
|---|---|---|---|
| Nsp5 | PPP2CA | 0.468 | 0.000185968 |
| Nsp12 | SEH1L | 0.468 | 0.000189035 |
| Nsp13 | TLR3 | 0.468 | 0.000185729 |
| Nsp16 | BRAF | 0.468 | 0.000183771 |
| Orf3a | EGF | 0.468 | 0.000187199 |
| Nsp4 | PAK2 | 0.467 | 0.000190232 |
| Nsp4 | CHUK | 0.467 | 0.000189149 |
| Nsp5 | FAS | 0.467 | 0.000189349 |
| Nsp10 | PPM1B | 0.467 | 0.000192876 |
| Nsp13 | SMAD5 | 0.467 | 0.000189379 |
| Nsp16 | LAMA2 | 0.467 | 0.000190244 |
| S | CXCL13 | 0.467 | 0.000192289 |
| S | RPS6KA5 | 0.467 | 0.000189399 |
| M | COL1A2 | 0.467 | 0.000193143 |
| Orf3a | DDX3X | 0.467 | 0.00019457 |
| Orf7a | CTSS | 0.467 | 0.000193013 |
| Orf7a | YWHAZ | 0.467 | 0.000191757 |
| Orf7b | IL36G | 0.467 | 0.000190329 |
| Orf8 | MAPK6 | 0.467 | 0.000192195 |
| Nsp2 | CCNE2 | 0.466 | 0.000196449 |
| Nsp4 | MECOM | 0.466 | 0.000196881 |
| Nsp4 | SMAD5 | 0.466 | 0.000196631 |
| Nsp4 | CXCL10 | 0.466 | 0.000202802 |
| Nsp4 | ITGB8 | 0.466 | 0.000196863 |
| Nsp6 | DIABLO | 0.466 | 0.000197182 |
| Nsp7 | IL23R | 0.466 | 0.000202523 |
| Nsp8 | MAP3K4 | 0.466 | 0.000196854 |
| Nsp16 | CHUK | 0.466 | 0.000198555 |
| S | FLT3 | 0.466 | 0.000198426 |
| N | TAB2 | 0.466 | 0.0001987 |
| N | IL22RA2 | 0.466 | 0.0001968 |
| Orf3a | ATG5 | 0.466 | 0.000202819 |
| Orf6 | RAP1B | 0.466 | 0.000200567 |
| Orf7a | GDF9 | 0.466 | 0.000199609 |
| Nsp4 | EGF | 0.465 | 0.000206099 |
| Nsp5 | MALT1 | 0.465 | 0.000203842 |
| Nsp8 | ITGA4 | 0.465 | 0.000208369 |
| Nsp10 | LDHC | 0.465 | 0.000203791 |
| Nsp13 | NLK | 0.465 | 0.000207959 |
| Nsp14 | GHR | 0.465 | 0.000203738 |
| Nsp14 | LAMA2 | 0.465 | 0.000208577 |
| M | CAMK2D | 0.465 | 0.000206487 |
| N | IL7 | 0.465 | 0.000204595 |
| Orf8 | PRKAA2 | 0.465 | 0.000206532 |
| Orf8 | KRAS | 0.465 | 0.000206014 |
| Nsp5 | ATP6V1C1 | 0.464 | 0.000212071 |
| Nsp8 | SMAD2 | 0.464 | 0.000214628 |
| Nsp9 | IL2 | 0.464 | 0.000216095 |
| Nsp12 | TNFSF18 | 0.464 | 0.000212081 |
| Nsp16 | SMAD4 | 0.464 | 0.000216975 |

| | | | |
|---|---|---|---|
| S | E2F5 | 0.464 | 0.000213439 |
| N | SOCS5 | 0.464 | 0.000211395 |
| N | CUL1 | 0.464 | 0.000216788 |
| N | PTPN2 | 0.464 | 0.00021295 |
| Orf7a | CACNB4 | 0.464 | 0.000213679 |
| Nsp2 | TAOK1 | 0.463 | 0.000222499 |
| Nsp4 | TICAM2 | 0.463 | 0.000220533 |
| Nsp5 | MAP3K4 | 0.463 | 0.000223501 |
| Nsp8 | TGFBR1 | 0.463 | 0.000221495 |
| Nsp12 | LPAR6 | 0.463 | 0.000220856 |
| Nsp13 | PPP3CA | 0.463 | 0.0002222 |
| Nsp14 | ANGPT1 | 0.463 | 0.000225273 |
| Nsp14 | LPAR6 | 0.463 | 0.000220403 |
| Nsp15 | LAMTOR3 | 0.463 | 0.000224668 |
| Nsp15 | STAT4 | 0.463 | 0.000219257 |
| N | DCN | 0.463 | 0.00022408 |
| Orf7a | EIF2S1 | 0.463 | 0.000224277 |
| Orf7b | BAG4 | 0.463 | 0.000221183 |
| Orf8 | MDM2 | 0.463 | 0.000221883 |
| Nsp3 | LDHAL6A | 0.462 | 0.000227151 |
| Nsp3 | LPAR4 | 0.462 | 0.000233586 |
| Nsp5 | RASA2 | 0.462 | 0.000230942 |
| Nsp6 | PDHA2 | 0.462 | 0.000229285 |
| Nsp7 | CD36 | 0.462 | 0.000227731 |
| Nsp12 | CXCL13 | 0.462 | 0.000227966 |
| Nsp12 | BCL10 | 0.462 | 0.000232318 |
| Nsp15 | MET | 0.462 | 0.000229205 |
| M | LIFR | 0.462 | 0.000230372 |
| Orf7a | KIT | 0.462 | 0.000228311 |
| Nsp2 | TLR6 | 0.461 | 0.000234391 |
| Nsp4 | EIF2AK3 | 0.461 | 0.00024024 |
| Nsp5 | TLR8 | 0.461 | 0.000242257 |
| Nsp8 | EIF2AK3 | 0.461 | 0.000236692 |
| Nsp9 | MDM2 | 0.461 | 0.000240079 |
| Nsp13 | LAMA2 | 0.461 | 0.000234817 |
| S | PPP3CA | 0.461 | 0.000238731 |
| S | PPARGC1A | 0.461 | 0.000236334 |
| Orf3a | MAP3K4 | 0.461 | 0.000235654 |
| Orf7a | BCL10 | 0.461 | 0.000234554 |
| Nsp4 | HGF | 0.46 | 0.000250586 |
| Nsp9 | BMPR1B | 0.46 | 0.000248853 |
| Nsp13 | PDHB | 0.46 | 0.00024808 |
| Nsp13 | ACVR1C | 0.46 | 0.000243464 |
| Nsp16 | PPP2CA | 0.46 | 0.00024881 |
| S | PTGS2 | 0.46 | 0.000247443 |
| M | PRKAA1 | 0.46 | 0.000250738 |
| M | PPM1B | 0.46 | 0.000244584 |
| N | NEO1 | 0.46 | 0.000245393 |
| Orf3a | AKT3 | 0.46 | 0.000250364 |
| Orf7a | PLCB4 | 0.46 | 0.000244219 |

| | | | |
|---|---|---|---|
| Nsp3 | TLR7 | 0.459 | 0.000259242 |
| Nsp5 | VCAM1 | 0.459 | 0.00025569 |
| Nsp8 | IL18 | 0.459 | 0.000253158 |
| Nsp12 | CXCL10 | 0.459 | 0.000255032 |
| Nsp12 | BMPR2 | 0.459 | 0.000255859 |
| Nsp13 | RAP1A | 0.459 | 0.000257203 |
| Nsp15 | PAK2 | 0.459 | 0.000256547 |
| M | SMAD4 | 0.459 | 0.000258874 |
| Orf3a | ITGA6 | 0.459 | 0.00025765 |
| Nsp2 | PPP3CB | 0.458 | 0.000260237 |
| Nsp2 | DDX58 | 0.458 | 0.000267508 |
| Nsp3 | BMPR1B | 0.458 | 0.000263229 |
| Nsp4 | LRP6 | 0.458 | 0.000269352 |
| Nsp4 | LAMA2 | 0.458 | 0.000261315 |
| Nsp7 | USP25 | 0.458 | 0.00026335 |
| Nsp9 | CYLD | 0.458 | 0.000264052 |
| Nsp10 | SMAD4 | 0.458 | 0.000267506 |
| Nsp12 | PARP4 | 0.458 | 0.000260721 |
| Nsp12 | EIF2AK3 | 0.458 | 0.00026836 |
| Nsp14 | EIF2AK3 | 0.458 | 0.000266422 |
| Nsp15 | HGF | 0.458 | 0.000266477 |
| Nsp15 | ITGA6 | 0.458 | 0.000266304 |
| Nsp16 | RPS6KB1 | 0.458 | 0.000260343 |
| M | SGK3 | 0.458 | 0.000265999 |
| Orf8 | ATG5 | 0.458 | 0.000266317 |
| Orf8 | ACVR2A | 0.458 | 0.000268292 |
| Nsp2 | ITGA6 | 0.457 | 0.000276297 |
| Nsp2 | NF1 | 0.457 | 0.000277231 |
| Nsp3 | CXCL13 | 0.457 | 0.000271519 |
| Nsp5 | TNFSF18 | 0.457 | 0.0002709 |
| Nsp8 | PPP2R1B | 0.457 | 0.00027672 |
| Nsp16 | HGF | 0.457 | 0.000276623 |
| N | PRKAA1 | 0.457 | 0.000276413 |
| N | PRKACB | 0.457 | 0.000273316 |
| Orf8 | IFIH1 | 0.457 | 0.000275414 |
| Nsp2 | MAP3K1 | 0.456 | 0.000286693 |
| Nsp2 | RBL2 | 0.456 | 0.000287922 |
| Nsp6 | ITGB8 | 0.456 | 0.000280896 |
| Nsp7 | MAP4K3 | 0.456 | 0.000285553 |
| Nsp9 | IFNAR1 | 0.456 | 0.000279111 |
| Nsp9 | PLA2G4A | 0.456 | 0.000279888 |
| Nsp14 | SMAD5 | 0.456 | 0.000283673 |
| S | MMP1 | 0.456 | 0.000281815 |
| N | ATG5 | 0.456 | 0.000287759 |
| N | MAPK6 | 0.456 | 0.000284687 |
| Nsp3 | PLCB4 | 0.455 | 0.000291441 |
| Nsp6 | IL23R | 0.455 | 0.000291663 |
| Nsp7 | SOS2 | 0.455 | 0.000292335 |
| Nsp8 | MAPK8 | 0.455 | 0.00029575 |
| Nsp9 | PTPN13 | 0.455 | 0.00029109 |

| | | | |
|---|---|---|---|
| Nsp13 | PRKAR1A | 0.455 | 0.000292759 |
| Nsp14 | MAPK8 | 0.455 | 0.000295069 |
| Nsp15 | CDC42 | 0.455 | 0.000296277 |
| Nsp16 | CXCL13 | 0.455 | 0.000288962 |
| S | IL12RB2 | 0.455 | 0.000291659 |
| S | PIAS1 | 0.455 | 0.00028897 |
| S | OSMR | 0.455 | 0.000290599 |
| N | MMP13 | 0.455 | 0.000298073 |
| N | LDHAL6A | 0.455 | 0.000290522 |
| Nsp5 | PARP4 | 0.454 | 0.000302699 |
| Nsp9 | AZI2 | 0.454 | 0.000304275 |
| Nsp12 | TICAM2 | 0.454 | 0.000300271 |
| Nsp12 | NF1 | 0.454 | 0.000299569 |
| Nsp12 | SMAD2 | 0.454 | 0.000305663 |
| Nsp16 | TLR7 | 0.454 | 0.000306811 |
| S | HSPA8 | 0.454 | 0.000302695 |
| S | AREG | 0.454 | 0.000302536 |
| N | MAP3K4 | 0.454 | 0.000306818 |
| Orf3a | PTPN2 | 0.454 | 0.000304833 |
| Orf3a | TFRC | 0.454 | 0.000302222 |
| Orf7a | PPP2R5A | 0.454 | 0.000304093 |
| Orf8 | PAK2 | 0.454 | 0.000302802 |
| Orf8 | PRKACB | 0.454 | 0.000303936 |
| Nsp2 | PRKAR1A | 0.453 | 0.000312902 |
| Nsp2 | COL4A6 | 0.453 | 0.000318709 |
| Nsp6 | PPP2CA | 0.453 | 0.000316237 |
| Nsp6 | ACSL4 | 0.453 | 0.000314353 |
| Nsp7 | IL13RA2 | 0.453 | 0.000319588 |
| Nsp9 | PAK2 | 0.453 | 0.000318458 |
| Nsp9 | TLR4 | 0.453 | 0.000318099 |
| Nsp13 | GYS2 | 0.453 | 0.000315101 |
| Nsp14 | HGF | 0.453 | 0.000311807 |
| Nsp15 | GHR | 0.453 | 0.000316536 |
| Orf3a | NLK | 0.453 | 0.000310483 |
| Orf3a | BMPR1B | 0.453 | 0.000313341 |
| Orf3a | MSTN | 0.453 | 0.00031344 |
| Orf3a | MAPK8 | 0.453 | 0.000320336 |
| Orf7a | HSP90AA1 | 0.453 | 0.000309811 |
| Orf8 | ITGA1 | 0.453 | 0.000316242 |
| Nsp5 | SMAD2 | 0.452 | 0.000322533 |
| Nsp6 | PTGS2 | 0.452 | 0.000324669 |
| Nsp8 | MECOM | 0.452 | 0.000321945 |
| Nsp8 | BLNK | 0.452 | 0.00033155 |
| Nsp10 | PPP2R1B | 0.452 | 0.000322282 |
| Nsp12 | SMAD5 | 0.452 | 0.000328579 |
| Nsp13 | PLCB4 | 0.452 | 0.000331102 |
| Nsp14 | FGF7 | 0.452 | 0.000325348 |
| Nsp15 | LRP6 | 0.452 | 0.000325518 |
| Nsp15 | KRAS | 0.452 | 0.00032153 |
| S | IL1RL1 | 0.452 | 0.000328128 |

| | | | |
|---|---|---|---|
| Orf6 | EIF4E | 0.452 | 0.000328629 |
| Orf6 | LIFR | 0.452 | 0.000325344 |
| Orf8 | PTPN13 | 0.452 | 0.000323709 |
| Nsp4 | BMPR1B | 0.451 | 0.000337027 |
| Nsp5 | MAPK8 | 0.451 | 0.000336779 |
| Nsp6 | PPP3R1 | 0.451 | 0.000338356 |
| Nsp6 | ULK2 | 0.451 | 0.000333123 |
| Nsp7 | XIAP | 0.451 | 0.000337594 |
| Nsp8 | AZI2 | 0.451 | 0.000342202 |
| Nsp9 | PDHA2 | 0.451 | 0.000335891 |
| Nsp12 | MAP3K4 | 0.451 | 0.000339763 |
| Nsp15 | CAMK2D | 0.451 | 0.000336162 |
| Nsp15 | ACSL4 | 0.451 | 0.000338701 |
| M | CCNE2 | 0.451 | 0.000337033 |
| Orf3a | ACVR2A | 0.451 | 0.00033941 |
| Orf7a | TNFSF10 | 0.451 | 0.000340939 |
| Orf8 | SMAD2 | 0.451 | 0.000335384 |
| Nsp6 | LAMB4 | 0.45 | 0.00034923 |
| Nsp8 | PPP2R2A | 0.45 | 0.000353226 |
| Nsp8 | LAMA2 | 0.45 | 0.00034533 |
| Nsp9 | RPS6KB1 | 0.45 | 0.000343438 |
| Nsp15 | BMPR2 | 0.45 | 0.000350839 |
| M | HSPA8 | 0.45 | 0.00034518 |
| N | MALT1 | 0.45 | 0.000349031 |
| Orf3a | CDC42 | 0.45 | 0.000344576 |
| Orf7a | SORBS1 | 0.45 | 0.000347968 |
| Orf8 | HSP90B1 | 0.45 | 0.00034861 |
| Nsp2 | NLK | 0.449 | 0.000366276 |
| Nsp2 | MAP3K2 | 0.449 | 0.000365695 |
| Nsp3 | GYS2 | 0.449 | 0.000355108 |
| Nsp4 | ITGA6 | 0.449 | 0.000362054 |
| Nsp6 | AREG | 0.449 | 0.000361815 |
| Nsp8 | MALT1 | 0.449 | 0.000365188 |
| Nsp9 | TLR6 | 0.449 | 0.000367089 |
| Nsp9 | NFATC3 | 0.449 | 0.0003656 |
| Nsp9 | IL1RL1 | 0.449 | 0.000357211 |
| Nsp14 | TLR4 | 0.449 | 0.000366982 |
| M | XIAP | 0.449 | 0.000363353 |
| Orf8 | STRADB | 0.449 | 0.000361674 |
| Orf8 | DDX3X | 0.449 | 0.000362455 |
| Nsp1 | HSPA8 | 0.448 | 0.000369933 |
| Nsp3 | GDF9 | 0.448 | 0.000377362 |
| Nsp5 | NF1 | 0.448 | 0.000371166 |
| Nsp7 | RAP1B | 0.448 | 0.000370187 |
| Nsp8 | BRAF | 0.448 | 0.000372958 |
| Nsp9 | PRKAA1 | 0.448 | 0.000377518 |
| Nsp13 | BMPR1B | 0.448 | 0.000373627 |
| M | LDHB | 0.448 | 0.000370736 |
| N | MDM2 | 0.448 | 0.000371996 |
| Orf3a | MAP3K7 | 0.448 | 0.000375099 |

| | | | |
|---|---|---|---|
| Orf3a | RAPGEF2 | 0.448 | 0.000368755 |
| Orf8 | PRKAA1 | 0.448 | 0.000371311 |
| Nsp2 | CYLD | 0.447 | 0.000381127 |
| Nsp3 | ACVR1C | 0.447 | 0.000389853 |
| Nsp4 | GDF9 | 0.447 | 0.000380684 |
| Nsp8 | ATP6V1C1 | 0.447 | 0.000389071 |
| Nsp9 | VCAM1 | 0.447 | 0.000385902 |
| Nsp9 | RBL2 | 0.447 | 0.000387474 |
| Nsp13 | SLC38A9 | 0.447 | 0.000380473 |
| Nsp13 | YWHAZ | 0.447 | 0.000383961 |
| Nsp15 | COL1A2 | 0.447 | 0.000390092 |
| Nsp16 | TLR1 | 0.447 | 0.000388717 |
| N | CXCL13 | 0.447 | 0.000382857 |
| N | IFIH1 | 0.447 | 0.000392714 |
| N | BRCA1 | 0.447 | 0.00038139 |
| Orf3a | MET | 0.447 | 0.000388479 |
| Orf7a | IL5RA | 0.447 | 0.000388334 |
| Nsp3 | PPP2R5A | 0.446 | 0.000400186 |
| Nsp4 | BMP5 | 0.446 | 0.000397813 |
| Nsp5 | ACVR1C | 0.446 | 0.000402965 |
| Nsp6 | CXCL10 | 0.446 | 0.000405607 |
| Nsp7 | HSP90B1 | 0.446 | 0.000404029 |
| Nsp9 | PRKAA2 | 0.446 | 0.000403959 |
| Nsp15 | TLR4 | 0.446 | 0.000404915 |
| Nsp15 | PPP2CA | 0.446 | 0.000393696 |
| Nsp15 | PDGFC | 0.446 | 0.000401539 |
| Nsp16 | IL22RA2 | 0.446 | 0.000395977 |
| Orf7a | GYS2 | 0.446 | 0.000398252 |
| Orf7a | AIFM1 | 0.446 | 0.000397158 |
| Nsp2 | HSP90AA1 | 0.445 | 0.000415475 |
| Nsp3 | PIK3R1 | 0.445 | 0.000408498 |
| Nsp6 | ATG5 | 0.445 | 0.000412471 |
| Nsp6 | GYS2 | 0.445 | 0.000413416 |
| Nsp7 | DNM1L | 0.445 | 0.000407781 |
| Nsp9 | CAMK2D | 0.445 | 0.000412781 |
| Nsp9 | IL1R1 | 0.445 | 0.000418581 |
| Nsp14 | ITGA6 | 0.445 | 0.000410267 |
| Nsp15 | TLR3 | 0.445 | 0.000406933 |
| Nsp16 | FLT3 | 0.445 | 0.000412475 |
| M | LDHC | 0.445 | 0.000409091 |
| M | BIRC2 | 0.445 | 0.000416762 |
| N | MAP4K3 | 0.445 | 0.000418174 |
| N | RICTOR | 0.445 | 0.000416939 |
| Orf3a | MECOM | 0.445 | 0.000414929 |
| Orf8 | PPP2R2A | 0.445 | 0.000411684 |
| Orf8 | HGF | 0.445 | 0.000412093 |
| Orf8 | MAP3K2 | 0.445 | 0.000410331 |
| Orf8 | IL18R1 | 0.445 | 0.000411571 |
| Nsp6 | CAB39L | 0.444 | 0.000431344 |
| Nsp6 | CTSS | 0.444 | 0.000424889 |

| | | | |
|---|---|---|---|
| Nsp7 | IRAK4 | 0.444 | 0.000424299 |
| Nsp12 | CHUK | 0.444 | 0.00043447 |
| Nsp15 | ATP6V1C1 | 0.444 | 0.000431979 |
| Nsp15 | TLR2 | 0.444 | 0.000432515 |
| S | MYB | 0.444 | 0.000426339 |
| Orf3a | PPP2R1B | 0.444 | 0.0004207 |
| Orf7a | PDK1 | 0.444 | 0.000426135 |
| Orf8 | ITGAV | 0.444 | 0.00042457 |
| Nsp3 | SORBS1 | 0.443 | 0.000442068 |
| Nsp5 | GYS2 | 0.443 | 0.000435175 |
| Nsp5 | HGF | 0.443 | 0.000439258 |
| Nsp6 | RASA1 | 0.443 | 0.000441471 |
| Nsp8 | HGF | 0.443 | 0.000435767 |
| Nsp9 | OSMR | 0.443 | 0.000440737 |
| Nsp10 | PPP1R3A | 0.443 | 0.000437673 |
| Nsp14 | MALT1 | 0.443 | 0.000449089 |
| Nsp16 | PLCB4 | 0.443 | 0.000448961 |
| M | TFRC | 0.443 | 0.000448846 |
| M | RBL1 | 0.443 | 0.000445216 |
| N | MIOS | 0.443 | 0.000442364 |
| N | CXCL11 | 0.443 | 0.000448302 |
| Orf3a | PIK3R1 | 0.443 | 0.000435239 |
| Orf3a | SMURF2 | 0.443 | 0.000447652 |
| Nsp3 | SLC38A9 | 0.442 | 0.000454441 |
| Nsp3 | PPP3CA | 0.442 | 0.000454131 |
| Nsp5 | PIAS1 | 0.442 | 0.000455028 |
| Nsp6 | GHR | 0.442 | 0.000460906 |
| Nsp9 | ITCH | 0.442 | 0.000450055 |
| Nsp14 | RAPGEF2 | 0.442 | 0.000458962 |
| Nsp15 | SEH1L | 0.442 | 0.000459667 |
| Nsp16 | CASP3 | 0.442 | 0.00045429 |
| Nsp16 | MAP3K4 | 0.442 | 0.000452182 |
| N | STAM | 0.442 | 0.000450263 |
| Orf3a | LPAR6 | 0.442 | 0.000457174 |
| Nsp2 | RAPGEF2 | 0.441 | 0.000468938 |
| Nsp3 | ANGPT1 | 0.441 | 0.000473458 |
| Nsp3 | CASP3 | 0.441 | 0.000478224 |
| Nsp5 | BMPR2 | 0.441 | 0.000470641 |
| Nsp9 | CXCL10 | 0.441 | 0.000466707 |
| Nsp13 | ACSL4 | 0.441 | 0.00047444 |
| Nsp14 | PTPRR | 0.441 | 0.000468512 |
| Nsp14 | BMPR1B | 0.441 | 0.000477601 |
| Nsp16 | GYS2 | 0.441 | 0.000477243 |
| M | GNAI1 | 0.441 | 0.000468745 |
| M | ATM | 0.441 | 0.000471654 |
| N | PIK3R1 | 0.441 | 0.000473427 |
| N | IFNAR1 | 0.441 | 0.000476412 |
| Orf3a | SKP1 | 0.441 | 0.000477725 |
| Orf3a | OSMR | 0.441 | 0.000479083 |
| Orf7a | PDHB | 0.441 | 0.000473124 |

| | | | |
|---|---|---|---|
| Orf7a | ELK4 | 0.441 | 0.000470345 |
| Orf7a | LPAR6 | 0.441 | 0.000474424 |
| Orf7b | TAB2 | 0.441 | 0.00047789 |
| Orf8 | PPP3R1 | 0.441 | 0.000476312 |
| Nsp5 | SEH1L | 0.44 | 0.000488546 |
| Nsp5 | PPP3CA | 0.44 | 0.00048928 |
| Nsp6 | TLR5 | 0.44 | 0.00048212 |
| Nsp8 | COL4A4 | 0.44 | 0.000494647 |
| Nsp10 | PPP3CB | 0.44 | 0.000494621 |
| Nsp13 | SEH1L | 0.44 | 0.000480649 |
| Nsp13 | ELOC | 0.44 | 0.000485371 |
| Nsp13 | FLT3 | 0.44 | 0.000481659 |
| Nsp14 | MMP1 | 0.44 | 0.000488998 |
| M | MDM2 | 0.44 | 0.000494054 |
| N | LDHA | 0.44 | 0.000490454 |
| Orf7a | CFLAR | 0.44 | 0.000492914 |
| Orf7b | TGFBR1 | 0.44 | 0.000493399 |
| Orf8 | RBL2 | 0.44 | 0.000490587 |
| Orf8 | AZI2 | 0.44 | 0.000494306 |
| Orf8 | PPP2CB | 0.44 | 0.000484417 |
| Nsp2 | IL23R | 0.439 | 0.000511399 |
| Nsp5 | PAK2 | 0.439 | 0.00050742 |
| Nsp5 | DDX58 | 0.439 | 0.000511487 |
| Nsp6 | TNFSF10 | 0.439 | 0.000497086 |
| Nsp7 | SOS1 | 0.439 | 0.000506712 |
| Nsp8 | RPS6KA5 | 0.439 | 0.000502048 |
| Nsp9 | IL13RA2 | 0.439 | 0.000497772 |
| Nsp12 | TANK | 0.439 | 0.000498585 |
| Nsp12 | VCAM1 | 0.439 | 0.000506579 |
| Nsp15 | LY96 | 0.439 | 0.000507534 |
| Nsp15 | MAPK8 | 0.439 | 0.000497305 |
| S | CASP12 | 0.439 | 0.000501634 |
| S | ATP6V1D | 0.439 | 0.000501314 |
| E | ATP6V1A | 0.439 | 0.000503167 |
| M | IFNAR1 | 0.439 | 0.000497751 |
| N | JAK2 | 0.439 | 0.000499648 |
| Orf3a | PARP4 | 0.439 | 0.000499282 |
| Orf7a | IL13RA1 | 0.439 | 0.000509183 |
| Nsp4 | LDHAL6A | 0.438 | 0.00052445 |
| Nsp5 | MMP3 | 0.438 | 0.000528158 |
| Nsp7 | RPS6KA6 | 0.438 | 0.000522623 |
| Nsp7 | CD3G | 0.438 | 0.000527438 |
| Nsp14 | GDF9 | 0.438 | 0.000523834 |
| Nsp14 | ITGB8 | 0.438 | 0.000515366 |
| Nsp15 | GNAI1 | 0.438 | 0.000514613 |
| S | IL31RA | 0.438 | 0.000526977 |
| N | AHR | 0.438 | 0.000522361 |
| Orf6 | CXCL10 | 0.438 | 0.000520562 |
| Nsp2 | RASA2 | 0.437 | 0.000545346 |
| Nsp3 | LY96 | 0.437 | 0.000544942 |

| | | | |
|---|---|---|---|
| Nsp5 | TLR1 | 0.437 | 0.000546764 |
| Nsp8 | MAPK9 | 0.437 | 0.0005381 |
| Nsp9 | TICAM2 | 0.437 | 0.000538155 |
| Nsp13 | ATP6V1C1 | 0.437 | 0.000539543 |
| S | KIT | 0.437 | 0.000530549 |
| M | BCL10 | 0.437 | 0.000544031 |
| M | GNB4 | 0.437 | 0.000533303 |
| M | IL26 | 0.437 | 0.000539933 |
| N | HSPA8 | 0.437 | 0.000546611 |
| N | BMPR1B | 0.437 | 0.000538253 |
| N | STAT4 | 0.437 | 0.000536626 |
| Orf3a | GHR | 0.437 | 0.000536574 |
| Orf3a | CCL28 | 0.437 | 0.000546535 |
| Orf3a | MALT1 | 0.437 | 0.000531596 |
| Orf3a | PRKAA2 | 0.437 | 0.000534521 |
| Orf7a | MYB | 0.437 | 0.000536824 |
| Orf7a | PTPN11 | 0.437 | 0.000530798 |
| Orf8 | GHR | 0.437 | 0.000535714 |
| Orf8 | PPP2R3A | 0.437 | 0.000532055 |
| Nsp2 | CHUK | 0.436 | 0.000555937 |
| Nsp4 | TLR3 | 0.436 | 0.00055389 |
| Nsp6 | IL22RA2 | 0.436 | 0.00055787 |
| Nsp12 | ACSL4 | 0.436 | 0.000557958 |
| Nsp12 | RAPGEF2 | 0.436 | 0.000555778 |
| Nsp15 | CD86 | 0.436 | 0.00054807 |
| S | RPS6KB1 | 0.436 | 0.00055133 |
| M | LDHA | 0.436 | 0.000551533 |
| N | CASP12 | 0.436 | 0.00056335 |
| N | RBL2 | 0.436 | 0.000558587 |
| Orf7a | BLNK | 0.436 | 0.000551046 |
| Nsp7 | NFATC3 | 0.435 | 0.000570081 |
| Nsp7 | IFNAR1 | 0.435 | 0.00057544 |
| Nsp7 | IL33 | 0.435 | 0.000583572 |
| Nsp8 | HSP90AA1 | 0.435 | 0.000568413 |
| Nsp9 | ACSL3 | 0.435 | 0.000577741 |
| Nsp12 | PPP2R2A | 0.435 | 0.000580946 |
| Nsp14 | MAP3K4 | 0.435 | 0.000580088 |
| Nsp15 | VCAM1 | 0.435 | 0.00057657 |
| Nsp16 | COL4A6 | 0.435 | 0.000584692 |
| Nsp16 | MMP1 | 0.435 | 0.000579266 |
| M | PPP2R1B | 0.435 | 0.000574266 |
| Nsp2 | CD86 | 0.434 | 0.000586006 |
| Nsp5 | BMPR1B | 0.434 | 0.000591421 |
| Nsp8 | IL1R1 | 0.434 | 0.000597763 |
| Nsp8 | DDX3X | 0.434 | 0.000588462 |
| Nsp9 | MAP3K1 | 0.434 | 0.000588637 |
| Nsp10 | RPS6KA3 | 0.434 | 0.000593318 |
| Nsp14 | SMAD2 | 0.434 | 0.000603378 |
| Nsp15 | VAV3 | 0.434 | 0.000602312 |
| S | PDHB | 0.434 | 0.000600558 |

| | | | |
|---|---|---|---|
| S | MAP2K4 | 0.434 | 0.000598028 |
| M | CREB1 | 0.434 | 0.000602495 |
| M | COL4A3 | 0.434 | 0.000599717 |
| N | IL23R | 0.434 | 0.000598028 |
| Orf6 | STAM2 | 0.434 | 0.000601776 |
| Nsp2 | IL36G | 0.433 | 0.000613808 |
| Nsp2 | MAP3K4 | 0.433 | 0.000613394 |
| Nsp2 | GNG11 | 0.433 | 0.000618443 |
| Nsp4 | SLC38A9 | 0.433 | 0.000604801 |
| Nsp8 | TRAF6 | 0.433 | 0.000618899 |
| Nsp13 | TLR7 | 0.433 | 0.000609827 |
| Nsp16 | LAMB4 | 0.433 | 0.00060929 |
| Nsp16 | MAP2K4 | 0.433 | 0.000609131 |
| Nsp2 | ITGA2 | 0.432 | 0.000631165 |
| Nsp2 | MSTN | 0.432 | 0.000636936 |
| Nsp3 | E2F5 | 0.432 | 0.000638563 |
| Nsp3 | TLR1 | 0.432 | 0.000636894 |
| Nsp4 | MAP3K20 | 0.432 | 0.000629178 |
| Nsp5 | ITGA6 | 0.432 | 0.000627619 |
| Nsp9 | IL7R | 0.432 | 0.000627039 |
| Nsp10 | ZFYVE16 | 0.432 | 0.000629745 |
| Nsp12 | HGF | 0.432 | 0.000629616 |
| Nsp13 | CASP3 | 0.432 | 0.00063259 |
| Nsp15 | MECOM | 0.432 | 0.000626526 |
| Nsp15 | EIF4E | 0.432 | 0.000625708 |
| Nsp15 | EGF | 0.432 | 0.000624422 |
| Nsp15 | RAPGEF2 | 0.432 | 0.000628333 |
| Nsp16 | KIT | 0.432 | 0.000632936 |
| S | CFLAR | 0.432 | 0.000635848 |
| Orf3a | PGK2 | 0.432 | 0.000633683 |
| Orf7a | RELN | 0.432 | 0.000635034 |
| Orf8 | CAB39 | 0.432 | 0.000626586 |
| Nsp2 | ACVR1C | 0.431 | 0.000653512 |
| Nsp2 | LPAR4 | 0.431 | 0.000649256 |
| Nsp6 | NF1 | 0.431 | 0.000664675 |
| Nsp6 | MAPK8 | 0.431 | 0.000655671 |
| Nsp8 | RAP1A | 0.431 | 0.000657729 |
| Nsp8 | TLR1 | 0.431 | 0.000652989 |
| Nsp9 | IL5 | 0.431 | 0.000655089 |
| Nsp10 | PIK3CA | 0.431 | 0.000664403 |
| Nsp13 | MECOM | 0.431 | 0.000664277 |
| Nsp14 | ELK4 | 0.431 | 0.000659101 |
| Nsp15 | GNAI3 | 0.431 | 0.000651232 |
| M | GHR | 0.431 | 0.000661008 |
| M | HIF1A | 0.431 | 0.000648504 |
| N | CCNE2 | 0.431 | 0.000646568 |
| Orf3a | BAG4 | 0.431 | 0.000650018 |
| Orf8 | MAPK9 | 0.431 | 0.000654992 |
| Nsp1 | COL4A5 | 0.43 | 0.000669444 |
| Nsp2 | BRAF | 0.43 | 0.000681813 |

| | | | |
|---|---|---|---|
| Nsp5 | ITGB8 | 0.43 | 0.000686478 |
| Nsp5 | ATP6V1D | 0.43 | 0.000680303 |
| Nsp6 | OSMR | 0.43 | 0.00067174 |
| Nsp6 | MYB | 0.43 | 0.000669991 |
| Nsp7 | LY96 | 0.43 | 0.000672495 |
| Nsp10 | RICTOR | 0.43 | 0.000686808 |
| Nsp15 | BRAF | 0.43 | 0.000673086 |
| Nsp16 | PIK3R1 | 0.43 | 0.00067582 |
| Nsp16 | ANGPT1 | 0.43 | 0.000675403 |
| Nsp16 | LY96 | 0.43 | 0.000666215 |
| S | PTK2 | 0.43 | 0.000668256 |
| M | EIF2S1 | 0.43 | 0.00067142 |
| N | CTSS | 0.43 | 0.000675113 |
| N | CTSC | 0.43 | 0.000667308 |
| N | CSNK2A3 | 0.43 | 0.000679078 |
| N | MAP3K7 | 0.43 | 0.000687193 |
| Orf7a | MAP3K4 | 0.43 | 0.000669983 |
| Nsp3 | RAP1A | 0.429 | 0.000705223 |
| Nsp4 | AREG | 0.429 | 0.000703364 |
| Nsp5 | BCL10 | 0.429 | 0.000707101 |
| Nsp5 | ITGA4 | 0.429 | 0.000703041 |
| Nsp6 | HGF | 0.429 | 0.000697648 |
| Nsp6 | AIFM1 | 0.429 | 0.000700126 |
| Nsp6 | PTPN11 | 0.429 | 0.00070054 |
| Nsp7 | CUL2 | 0.429 | 0.000690108 |
| Nsp12 | MECOM | 0.429 | 0.000703766 |
| Nsp13 | KITLG | 0.429 | 0.000688654 |
| Nsp16 | PIAS1 | 0.429 | 0.000708398 |
| S | IL21 | 0.429 | 0.000696187 |
| S | RELN | 0.429 | 0.000688391 |
| N | KRAS | 0.429 | 0.000697603 |
| Orf3a | ATP6V1C1 | 0.429 | 0.000701275 |
| Nsp3 | KITLG | 0.428 | 0.000711395 |
| Nsp3 | PDHB | 0.428 | 0.000726047 |
| Nsp5 | IL18R1 | 0.428 | 0.000713883 |
| Nsp8 | LPAR4 | 0.428 | 0.00071691 |
| Nsp9 | STRADB | 0.428 | 0.000718216 |
| Nsp9 | MAP3K20 | 0.428 | 0.000713835 |
| Nsp12 | MMP13 | 0.428 | 0.000727931 |
| Nsp13 | PPP2R5A | 0.428 | 0.000717682 |
| Nsp14 | SLC38A9 | 0.428 | 0.000731166 |
| Nsp15 | PPP2R5A | 0.428 | 0.000726055 |
| Nsp16 | GHR | 0.428 | 0.000711092 |
| Nsp16 | LPAR4 | 0.428 | 0.000724222 |
| S | TRAF6 | 0.428 | 0.000711396 |
| S | ULK2 | 0.428 | 0.000712814 |
| M | HSP90B1 | 0.428 | 0.000728562 |
| Orf3a | BLNK | 0.428 | 0.000718598 |
| Orf7a | SLC38A9 | 0.428 | 0.00071633 |
| Orf7a | ITGB6 | 0.428 | 0.000709621 |

| | | | |
|---|---|---|---|
| Orf7a | ACSL4 | 0.428 | 0.000728058 |
| Orf7a | TLR8 | 0.428 | 0.000725172 |
| Nsp2 | CREB1 | 0.427 | 0.000743001 |
| Nsp2 | YWHAZ | 0.427 | 0.000738536 |
| Nsp3 | LAMTOR3 | 0.427 | 0.00074036 |
| Nsp3 | HSP90AA1 | 0.427 | 0.000736143 |
| Nsp3 | FLT3 | 0.427 | 0.000749421 |
| Nsp4 | PPP2R5A | 0.427 | 0.000742735 |
| Nsp6 | IL1RL2 | 0.427 | 0.000749053 |
| Nsp6 | IL1RL1 | 0.427 | 0.000740669 |
| Nsp7 | ITGA2 | 0.427 | 0.000750994 |
| Nsp8 | MMP3 | 0.427 | 0.000735309 |
| Nsp9 | PIK3CB | 0.427 | 0.000755331 |
| Nsp9 | MAP3K2 | 0.427 | 0.000734198 |
| Nsp10 | MSTN | 0.427 | 0.000751059 |
| Nsp12 | EIF4E | 0.427 | 0.000744233 |
| Nsp13 | PTK2 | 0.427 | 0.000745974 |
| M | PIAS1 | 0.427 | 0.000751267 |
| N | ROCK1 | 0.427 | 0.000736741 |
| N | SORBS1 | 0.427 | 0.00075468 |
| Orf3a | PPM1A | 0.427 | 0.000754717 |
| Orf7a | PPP3CA | 0.427 | 0.00073818 |
| Orf7a | IL1RL1 | 0.427 | 0.000741183 |
| Orf8 | FNIP1 | 0.427 | 0.000741958 |
| Nsp1 | DDX3X | 0.426 | 0.000760239 |
| Nsp2 | ATG5 | 0.426 | 0.000772028 |
| Nsp2 | BCL10 | 0.426 | 0.000759132 |
| Nsp6 | RPS6KB1 | 0.426 | 0.0007644 |
| Nsp7 | MSTN | 0.426 | 0.00076956 |
| Nsp12 | ITGA6 | 0.426 | 0.000775155 |
| Nsp12 | LY96 | 0.426 | 0.000771796 |
| Nsp14 | PTPN11 | 0.426 | 0.000757389 |
| Nsp15 | MAP3K4 | 0.426 | 0.000773723 |
| S | MAP3K5 | 0.426 | 0.000759321 |
| Nsp5 | AZI2 | 0.425 | 0.000790637 |
| Nsp6 | BCL10 | 0.425 | 0.000797702 |
| Nsp6 | ATP6V1D | 0.425 | 0.000782633 |
| Nsp7 | TLR6 | 0.425 | 0.000796332 |
| Nsp7 | LIFR | 0.425 | 0.00079658 |
| Nsp8 | GNAI1 | 0.425 | 0.000791805 |
| Nsp10 | IL7 | 0.425 | 0.000783706 |
| Nsp12 | ANGPT1 | 0.425 | 0.000782451 |
| Nsp12 | PPP3CA | 0.425 | 0.000792457 |
| Nsp12 | LDHAL6A | 0.425 | 0.000791045 |
| Nsp12 | MAPK8 | 0.425 | 0.000783615 |
| Nsp15 | PPP3CA | 0.425 | 0.00078688 |
| Nsp16 | PTPN11 | 0.425 | 0.000804014 |
| N | PARP4 | 0.425 | 0.000802725 |
| Orf3a | ACSL3 | 0.425 | 0.000787602 |
| Orf7a | BCL2A1 | 0.425 | 0.000781347 |

| | | | |
|---|---|---|---|
| Nsp2 | ATP6V1C1 | 0.424 | 0.000819329 |
| Nsp4 | HSP90AA1 | 0.424 | 0.000819393 |
| Nsp6 | TANK | 0.424 | 0.00082231 |
| Nsp8 | GDF9 | 0.424 | 0.000827934 |
| Nsp9 | NF1 | 0.424 | 0.000813566 |
| Nsp13 | VCAM1 | 0.424 | 0.000813774 |
| Nsp14 | IL21 | 0.424 | 0.000827962 |
| Nsp14 | E2F5 | 0.424 | 0.000815693 |
| Nsp16 | TLR4 | 0.424 | 0.000830235 |
| Nsp16 | PTPRR | 0.424 | 0.000823427 |
| Nsp16 | TLR2 | 0.424 | 0.000818697 |
| M | IL13RA2 | 0.424 | 0.000826029 |
| N | STAM2 | 0.424 | 0.000810092 |
| N | TGFBR1 | 0.424 | 0.000807536 |
| Orf3a | LAMB4 | 0.424 | 0.000819747 |
| Orf3a | LY96 | 0.424 | 0.000822167 |
| Orf7a | CASP12 | 0.424 | 0.000826737 |
| Orf8 | PTPN2 | 0.424 | 0.000819474 |
| Nsp2 | PPP2R3C | 0.423 | 0.000836723 |
| Nsp2 | PPP2R5E | 0.423 | 0.000850524 |
| Nsp3 | ATP6V1D | 0.423 | 0.000852834 |
| Nsp8 | LAMB4 | 0.423 | 0.000853353 |
| Nsp8 | MAP2K4 | 0.423 | 0.00084039 |
| Nsp8 | ITGA6 | 0.423 | 0.000851512 |
| Nsp12 | TLR4 | 0.423 | 0.000852586 |
| Nsp14 | CD40LG | 0.423 | 0.00084196 |
| Nsp16 | CAMK2D | 0.423 | 0.000850108 |
| S | IL7R | 0.423 | 0.000833766 |
| M | COL9A1 | 0.423 | 0.000848198 |
| N | STRADB | 0.423 | 0.000844514 |
| N | BMPR2 | 0.423 | 0.000840321 |
| Orf3a | TAOK1 | 0.423 | 0.000851623 |
| Orf7a | PPP2CB | 0.423 | 0.000836859 |
| Nsp3 | CASP12 | 0.422 | 0.000875628 |
| Nsp3 | IL1RL1 | 0.422 | 0.000867037 |
| Nsp4 | GYS2 | 0.422 | 0.000867173 |
| Nsp4 | ATP6V1D | 0.422 | 0.000863896 |
| Nsp5 | YWHAZ | 0.422 | 0.000860689 |
| Nsp7 | TAB2 | 0.422 | 0.000872157 |
| Nsp12 | TLR3 | 0.422 | 0.000863182 |
| Nsp13 | CXCL10 | 0.422 | 0.000879759 |
| Nsp13 | BLNK | 0.422 | 0.000879431 |
| Nsp14 | EGF | 0.422 | 0.000868954 |
| Nsp2 | ACSL4 | 0.421 | 0.000907599 |
| Nsp2 | SORBS1 | 0.421 | 0.000909354 |
| Nsp7 | CDC42 | 0.421 | 0.00090312 |
| Nsp7 | COL4A5 | 0.421 | 0.000897684 |
| Nsp10 | RPS6KA6 | 0.421 | 0.000894458 |
| Nsp13 | BMPR1A | 0.421 | 0.000910309 |
| Nsp14 | ACSL3 | 0.421 | 0.000895704 |

| | | | |
|---|---|---|---|
| Nsp14 | TLR3 | 0.421 | 0.000912482 |
| Nsp14 | DDX3X | 0.421 | 0.000908493 |
| Nsp15 | CREB1 | 0.421 | 0.000905958 |
| Nsp16 | ACVR1C | 0.421 | 0.000891133 |
| S | RPS6 | 0.421 | 0.000888936 |
| M | PPP2R5A | 0.421 | 0.000898559 |
| N | PIK3CA | 0.421 | 0.000912451 |
| N | LPAR4 | 0.421 | 0.000887951 |
| N | PLCB4 | 0.421 | 0.000904428 |
| Orf3a | TANK | 0.421 | 0.00090871 |
| Orf3a | BMPR2 | 0.421 | 0.000897507 |
| Orf3a | EIF2AK3 | 0.421 | 0.000886321 |
| Orf8 | COL1A2 | 0.421 | 0.000896022 |
| Nsp2 | ATP6V1H | 0.42 | 0.000918757 |
| Nsp2 | GNAI1 | 0.42 | 0.000927593 |
| Nsp4 | MAPK8 | 0.42 | 0.000936805 |
| Nsp9 | MAP3K4 | 0.42 | 0.000941624 |
| Nsp9 | PPM1A | 0.42 | 0.000941615 |
| Nsp10 | IFNAR1 | 0.42 | 0.00093651 |
| Nsp13 | PAK2 | 0.42 | 0.000935255 |
| Nsp13 | MMP13 | 0.42 | 0.000936961 |
| M | CUL2 | 0.42 | 0.000918879 |
| Orf3a | CUL1 | 0.42 | 0.000922459 |
| Orf6 | IL1R1 | 0.42 | 0.000928119 |
| Orf7a | SOCS4 | 0.42 | 0.000939814 |
| Nsp3 | TRAF6 | 0.419 | 0.000949041 |
| Nsp3 | PTK2 | 0.419 | 0.000971225 |
| Nsp4 | RPS6 | 0.419 | 0.000959509 |
| Nsp6 | PRKAR1A | 0.419 | 0.000952033 |
| Nsp7 | APAF1 | 0.419 | 0.000945255 |
| Nsp12 | RPS6KB1 | 0.419 | 0.000962653 |
| M | TBK1 | 0.419 | 0.000952228 |
| N | PPP2R3C | 0.419 | 0.00096801 |
| N | CUL2 | 0.419 | 0.000966538 |
| Orf3a | PIK3CA | 0.419 | 0.000948402 |
| Orf6 | RBL1 | 0.419 | 0.000950329 |
| Nsp3 | MMP1 | 0.418 | 0.000992693 |
| Nsp5 | RAP1A | 0.418 | 0.00099554 |
| Nsp6 | ATP6V1B2 | 0.418 | 0.000993664 |
| Nsp6 | CASP3 | 0.418 | 0.00099466 |
| Nsp9 | PPP2R3A | 0.418 | 0.000978952 |
| Nsp10 | GNB4 | 0.418 | 0.00098499 |
| Nsp13 | CASP12 | 0.418 | 0.000973061 |
| Nsp13 | VAV3 | 0.418 | 0.00099001 |
| Nsp15 | MMP13 | 0.418 | 0.000976636 |
| Nsp16 | IL12RB2 | 0.418 | 0.000981893 |
| E | TAB2 | 0.418 | 0.000984686 |
| E | HSP90B1 | 0.418 | 0.000987241 |
| M | EIF2AK3 | 0.418 | 0.000999101 |
| Orf7a | PRKAR1A | 0.418 | 0.000997196 |

| | | | |
|---|---|---|---|
| Orf7a | IL18RAP | 0.418 | 0.000992703 |
| Orf7a | ACVR1C | 0.418 | 0.000982399 |
| Orf7a | LY96 | 0.418 | 0.00097251 |
| Nsp2 | ITGA3 | -0.418 | 0.000974623 |
| Nsp2 | ELAVL1 | -0.418 | 0.000998798 |
| Nsp2 | ID2 | -0.418 | 0.000984066 |
| Nsp3 | ELK1 | -0.418 | 0.000981448 |
| Nsp3 | ARRB1 | -0.418 | 0.000982051 |
| Nsp3 | HAMP | -0.418 | 0.000997436 |
| Nsp4 | TUBA3E | -0.418 | 0.000973263 |
| Nsp4 | NFKBIA | -0.418 | 0.000996597 |
| Nsp6 | MAPK4 | -0.418 | 0.000999596 |
| Nsp7 | FZD7 | -0.418 | 0.000997819 |
| Nsp7 | IKBKE | -0.418 | 0.000980034 |
| Nsp7 | GDF5 | -0.418 | 0.000982136 |
| Nsp9 | IRF7 | -0.418 | 0.000992643 |
| Nsp9 | ACKR3 | -0.418 | 0.000984377 |
| Nsp9 | LCK | -0.418 | 0.000995278 |
| Nsp9 | ADCY3 | -0.418 | 0.000983624 |
| Nsp10 | PHLPP1 | -0.418 | 0.000996203 |
| Nsp10 | CACNA2D2 | -0.418 | 0.000994018 |
| Nsp10 | PIN1 | -0.418 | 0.000987645 |
| Nsp12 | IL25 | -0.418 | 0.000993423 |
| Nsp12 | ARRB1 | -0.418 | 0.000992651 |
| Nsp12 | ATP6V1F | -0.418 | 0.000976359 |
| Nsp14 | IRF5 | -0.418 | 0.000999911 |
| Nsp14 | IKBKG | -0.418 | 0.000994948 |
| Nsp15 | RPS6KA1 | -0.418 | 0.000983484 |
| Nsp15 | CCL24 | -0.418 | 0.000993264 |
| Nsp16 | ADCY2 | -0.418 | 0.000989218 |
| S | TIMP1 | -0.418 | 0.000977673 |
| S | ID2 | -0.418 | 0.000994067 |
| S | PTPN5 | -0.418 | 0.000976259 |
| S | CISH | -0.418 | 0.000980652 |
| Orf3a | MAPK15 | -0.418 | 0.000991543 |
| Orf3a | FZD2 | -0.418 | 0.00099162 |
| Orf3a | ITGA2B | -0.418 | 0.000972885 |
| Orf6 | PIDD1 | -0.418 | 0.000977005 |
| Orf6 | JUNB | -0.418 | 0.000992476 |
| Orf6 | RXRA | -0.418 | 0.000998013 |
| Orf6 | DUSP6 | -0.418 | 0.000999103 |
| Orf6 | STK11 | -0.418 | 0.000977939 |
| Orf6 | ZAP70 | -0.418 | 0.00097514 |
| Orf8 | CCL25 | -0.418 | 0.000985145 |
| Orf8 | PRKACA | -0.418 | 0.000990533 |
| Orf10 | TNFSF4 | -0.418 | 0.000989031 |
| Orf10 | HSPA1A | -0.418 | 0.00097765 |
| Orf10 | HSPA1B | -0.418 | 0.000987927 |
| Nsp2 | NGFR | -0.419 | 0.000954579 |
| Nsp3 | RRAGA | -0.419 | 0.000961972 |

| | | | |
|---|---|---|---|
| Nsp4 | EGFR | -0.419 | 0.000945394 |
| Nsp4 | ERN1 | -0.419 | 0.000956748 |
| Nsp5 | TGFB3 | -0.419 | 0.000946871 |
| Nsp6 | CAPN2 | -0.419 | 0.000970479 |
| Nsp6 | JUNB | -0.419 | 0.000966656 |
| Nsp6 | WNT9A | -0.419 | 0.000960631 |
| Nsp6 | JUND | -0.419 | 0.000948516 |
| Nsp7 | TNFRSF6B | -0.419 | 0.000942867 |
| Nsp7 | TYK2 | -0.419 | 0.000949449 |
| Nsp8 | PGF | -0.419 | 0.000943408 |
| Nsp8 | GH2 | -0.419 | 0.000946193 |
| Nsp8 | IKBKG | -0.419 | 0.000944533 |
| Nsp9 | TNFRSF18 | -0.419 | 0.000947847 |
| Nsp9 | PIK3CD | -0.419 | 0.000966815 |
| Nsp10 | CDKN2B | -0.419 | 0.000954712 |
| Nsp10 | FGF19 | -0.419 | 0.000945239 |
| Nsp10 | TNFRSF13C | -0.419 | 0.000943951 |
| Nsp12 | BMP4 | -0.419 | 0.000948352 |
| Nsp12 | FLNB | -0.419 | 0.000956902 |
| Nsp13 | HKDC1 | -0.419 | 0.000957126 |
| Nsp14 | ADCY8 | -0.419 | 0.000961349 |
| Nsp14 | SESN2 | -0.419 | 0.000960885 |
| Nsp14 | TNFRSF12A | -0.419 | 0.000950479 |
| Nsp15 | SHC3 | -0.419 | 0.000944991 |
| Nsp15 | IL15RA | -0.419 | 0.000944781 |
| Nsp16 | IL2RB | -0.419 | 0.000956182 |
| Nsp16 | FLNB | -0.419 | 0.000951765 |
| S | CSF2 | -0.419 | 0.000942309 |
| S | HK3 | -0.419 | 0.000963558 |
| S | IL12RB1 | -0.419 | 0.00096641 |
| Orf6 | LIF | -0.419 | 0.000943454 |
| Orf7a | CCL14 | -0.419 | 0.000960104 |
| Nsp2 | HCK | -0.42 | 0.00094101 |
| Nsp2 | SMAD3 | -0.42 | 0.000924642 |
| Nsp2 | IL17B | -0.42 | 0.000918461 |
| Nsp2 | FOXO3 | -0.42 | 0.000939873 |
| Nsp2 | CACNG5 | -0.42 | 0.000941034 |
| Nsp3 | IL2RB | -0.42 | 0.000941787 |
| Nsp3 | CXCR2 | -0.42 | 0.000936675 |
| Nsp3 | TGFA | -0.42 | 0.000930333 |
| Nsp3 | ERN1 | -0.42 | 0.00092054 |
| Nsp3 | CSF2 | -0.42 | 0.000928534 |
| Nsp3 | SERPINE1 | -0.42 | 0.000938833 |
| Nsp4 | PRKAB1 | -0.42 | 0.00093066 |
| Nsp4 | CPT1A | -0.42 | 0.000916206 |
| Nsp4 | RASGRP4 | -0.42 | 0.00092326 |
| Nsp4 | ITGA7 | -0.42 | 0.000933559 |
| Nsp5 | IL17RE | -0.42 | 0.000936581 |
| Nsp5 | CLCF1 | -0.42 | 0.000930721 |
| Nsp6 | PIAS4 | -0.42 | 0.000918759 |

| | | | |
|---|---|---|---|
| Nsp7 | HSPA6 | -0.42 | 0.00091639 |
| Nsp7 | GNA12 | -0.42 | 0.000940627 |
| Nsp7 | NCF1 | -0.42 | 0.000935088 |
| Nsp7 | GDF1 | -0.42 | 0.000916218 |
| Nsp7 | PLCB3 | -0.42 | 0.000921357 |
| Nsp7 | FGF8 | -0.42 | 0.000931465 |
| Nsp8 | IL11 | -0.42 | 0.000918645 |
| Nsp8 | ELAVL1 | -0.42 | 0.000936287 |
| Nsp9 | TNFRSF25 | -0.42 | 0.000941541 |
| Nsp9 | CAMKK2 | -0.42 | 0.000915853 |
| Nsp10 | PDGFA | -0.42 | 0.000914351 |
| Nsp10 | TNFRSF14 | -0.42 | 0.000930584 |
| Nsp10 | HRK | -0.42 | 0.000934727 |
| Nsp12 | ANGPT4 | -0.42 | 0.000924762 |
| Nsp13 | MMP14 | -0.42 | 0.000914748 |
| Nsp14 | PIDD1 | -0.42 | 0.000918679 |
| Nsp16 | FOS | -0.42 | 0.000928201 |
| S | TUBA8 | -0.42 | 0.000915024 |
| Orf3a | IRF7 | -0.42 | 0.000922434 |
| Orf3a | ADCY9 | -0.42 | 0.000921336 |
| Orf3a | EIF4E1B | -0.42 | 0.000932019 |
| Orf3a | PHLPP1 | -0.42 | 0.000914136 |
| Orf6 | TNFRSF4 | -0.42 | 0.000936469 |
| Orf6 | CACNA1I | -0.42 | 0.000933097 |
| Orf6 | RARA | -0.42 | 0.000913566 |
| Orf6 | FADD | -0.42 | 0.000920706 |
| Orf8 | IRF5 | -0.42 | 0.000917869 |
| Orf10 | DVL2 | -0.42 | 0.000940437 |
| Nsp2 | GNG13 | -0.421 | 0.000895979 |
| Nsp2 | SESN2 | -0.421 | 0.000902015 |
| Nsp2 | TNF | -0.421 | 0.000903241 |
| Nsp2 | CPT1C | -0.421 | 0.000912795 |
| Nsp3 | VEGFB | -0.421 | 0.000907984 |
| Nsp4 | PPP2R5B | -0.421 | 0.000889419 |
| Nsp4 | CSF3 | -0.421 | 0.000900385 |
| Nsp4 | FGFR1 | -0.421 | 0.000895166 |
| Nsp5 | PDGFA | -0.421 | 0.000890393 |
| Nsp5 | HMOX1 | -0.421 | 0.000886188 |
| Nsp5 | BCL2L1 | -0.421 | 0.000900081 |
| Nsp6 | WNT3 | -0.421 | 0.000898137 |
| Nsp6 | ID4 | -0.421 | 0.00090454 |
| Nsp6 | COL6A1 | -0.421 | 0.000908705 |
| Nsp7 | FOXO1 | -0.421 | 0.000912837 |
| Nsp7 | PFKFB3 | -0.421 | 0.000903224 |
| Nsp8 | MAP4K1 | -0.421 | 0.00091251 |
| Nsp8 | EGLN2 | -0.421 | 0.000912033 |
| Nsp9 | CARD14 | -0.421 | 0.000905693 |
| Nsp9 | TRADD | -0.421 | 0.000911265 |
| Nsp9 | CACNG1 | -0.421 | 0.000904382 |
| Nsp9 | TGFB1 | -0.421 | 0.000887973 |

| | | | |
|---|---|---|---|
| Nsp10 | CXCR5 | -0.421 | 0.000907805 |
| Nsp10 | IRF7 | -0.421 | 0.000891552 |
| Nsp10 | ICAM1 | -0.421 | 0.000899864 |
| Nsp10 | ACVRL1 | -0.421 | 0.000897203 |
| Nsp10 | TNFRSF1A | -0.421 | 0.00091304 |
| Nsp12 | IRF4 | -0.421 | 0.000901542 |
| Nsp13 | FLNB | -0.421 | 0.00090729 |
| Nsp15 | FGF2 | -0.421 | 0.000892434 |
| Nsp15 | GADD45A | -0.421 | 0.00090117 |
| Nsp16 | IL4 | -0.421 | 0.000889702 |
| Nsp16 | EGFR | -0.421 | 0.000899284 |
| Nsp16 | ACACB | -0.421 | 0.000885784 |
| S | ARRB2 | -0.421 | 0.00090792 |
| S | TRAF3 | -0.421 | 0.000895059 |
| Orf3a | CACNA1C | -0.421 | 0.000904978 |
| Orf3a | EPOR | -0.421 | 0.000901401 |
| Orf3a | IGF2 | -0.421 | 0.000895122 |
| Orf6 | CAMKK2 | -0.421 | 0.0009016 |
| Orf6 | WNT10B | -0.421 | 0.000909435 |
| Orf6 | RELA | -0.421 | 0.000898069 |
| Orf6 | SRF | -0.421 | 0.000892058 |
| Orf7a | PTPN6 | -0.421 | 0.000890056 |
| Orf8 | IRS1 | -0.421 | 0.000886562 |
| Orf8 | CCL11 | -0.421 | 0.000905226 |
| Orf8 | ADCY6 | -0.421 | 0.000908139 |
| Orf10 | MAP4K2 | -0.421 | 0.000905436 |
| Nsp2 | CACNG7 | -0.422 | 0.000883967 |
| Nsp2 | YWHAG | -0.422 | 0.00086617 |
| Nsp2 | VHL | -0.422 | 0.000858316 |
| Nsp3 | TNFRSF14 | -0.422 | 0.000881895 |
| Nsp3 | TAOK2 | -0.422 | 0.000879847 |
| Nsp3 | EIF4E1B | -0.422 | 0.000860249 |
| Nsp4 | CTSF | -0.422 | 0.000858635 |
| Nsp5 | AKT1S1 | -0.422 | 0.000869464 |
| Nsp5 | GH2 | -0.422 | 0.000876495 |
| Nsp6 | CXCR3 | -0.422 | 0.000858899 |
| Nsp6 | DUSP4 | -0.422 | 0.000882123 |
| Nsp6 | CAPN1 | -0.422 | 0.000878405 |
| Nsp8 | CREBBP | -0.422 | 0.000878116 |
| Nsp9 | LEFTY1 | -0.422 | 0.000880761 |
| Nsp10 | ACVR2B | -0.422 | 0.000883794 |
| Nsp10 | DUSP4 | -0.422 | 0.000868431 |
| Nsp11 | RXRG | -0.422 | 0.000875432 |
| Nsp12 | SLC2A4 | -0.422 | 0.00087105 |
| Nsp13 | CTSF | -0.422 | 0.00087774 |
| Nsp13 | CACNG3 | -0.422 | 0.000858094 |
| Nsp14 | EGLN2 | -0.422 | 0.000878308 |
| Nsp15 | NOD2 | -0.422 | 0.000861605 |
| Nsp16 | TUBA3E | -0.422 | 0.000870957 |
| S | FGF2 | -0.422 | 0.000880683 |

| | | | |
|---|---|---|---|
| S | CACNA1S | -0.422 | 0.000884776 |
| S | TAOK2 | -0.422 | 0.000884957 |
| S | TNF | -0.422 | 0.000879293 |
| S | LAMTOR2 | -0.422 | 0.000882704 |
| S | IKBKG | -0.422 | 0.000874703 |
| Orf3a | ITGB4 | -0.422 | 0.000862296 |
| Orf3a | CTSD | -0.422 | 0.000864565 |
| Orf3a | TYK2 | -0.422 | 0.000865466 |
| Orf3a | ULK1 | -0.422 | 0.000861643 |
| Orf3a | FADD | -0.422 | 0.000866216 |
| Orf6 | DAB2IP | -0.422 | 0.00088365 |
| Orf8 | TGFB3 | -0.422 | 0.000881513 |
| Orf8 | IL4R | -0.422 | 0.000878833 |
| Orf8 | HRAS | -0.422 | 0.000862354 |
| Orf8 | INS | -0.422 | 0.000883762 |
| Orf10 | DHX58 | -0.422 | 0.000862284 |
| Nsp2 | ITPR3 | -0.423 | 0.000851217 |
| Nsp2 | EDAR | -0.423 | 0.000841027 |
| Nsp2 | WDR24 | -0.423 | 0.000833182 |
| Nsp2 | RAPGEF1 | -0.423 | 0.000850505 |
| Nsp3 | CALML6 | -0.423 | 0.00084793 |
| Nsp4 | NBL1 | -0.423 | 0.000838256 |
| Nsp5 | CSF1R | -0.423 | 0.000853034 |
| Nsp5 | RELA | -0.423 | 0.000831795 |
| Nsp5 | MAPKAPK2 | -0.423 | 0.000855074 |
| Nsp6 | WNT5B | -0.423 | 0.000838097 |
| Nsp7 | LRP5 | -0.423 | 0.00085742 |
| Nsp7 | RASGRP2 | -0.423 | 0.000847168 |
| Nsp7 | GNB3 | -0.423 | 0.000847827 |
| Nsp7 | STK11 | -0.423 | 0.000853671 |
| Nsp7 | ZAP70 | -0.423 | 0.000834209 |
| Nsp8 | CPT1A | -0.423 | 0.000844679 |
| Nsp9 | LIF | -0.423 | 0.000835893 |
| Nsp9 | ID4 | -0.423 | 0.000847402 |
| Nsp10 | BCL2 | -0.423 | 0.000849714 |
| Nsp12 | TNR | -0.423 | 0.000853119 |
| Nsp12 | LAMB3 | -0.423 | 0.000834901 |
| Nsp12 | EFNA4 | -0.423 | 0.000854797 |
| Nsp13 | MAPT | -0.423 | 0.000851759 |
| Nsp13 | ELOB | -0.423 | 0.000848646 |
| Nsp14 | GH1 | -0.423 | 0.000855085 |
| Nsp14 | TNFRSF13B | -0.423 | 0.000842108 |
| Nsp15 | EGFR | -0.423 | 0.00084045 |
| Nsp15 | CD19 | -0.423 | 0.000843982 |
| Orf3a | FGF22 | -0.423 | 0.000856924 |
| Orf6 | GDF1 | -0.423 | 0.000839519 |
| Orf6 | SREBF1 | -0.423 | 0.000849822 |
| Orf8 | PDGFA | -0.423 | 0.000845975 |
| Orf8 | EFNA4 | -0.423 | 0.000854322 |
| Orf8 | HK3 | -0.423 | 0.000835449 |

| | | | |
|---|---|---|---|
| Orf8 | PLA2G4D | -0.423 | 0.000837764 |
| Orf8 | IL17C | -0.423 | 0.000832287 |
| Orf8 | MRAS | -0.423 | 0.000842359 |
| Orf8 | CSF3R | -0.423 | 0.000840684 |
| Nsp2 | CSF2RB | -0.424 | 0.000809041 |
| Nsp2 | FLT3LG | -0.424 | 0.000810288 |
| Nsp3 | S100A9 | -0.424 | 0.000828153 |
| Nsp4 | TNFSF12 | -0.424 | 0.000829724 |
| Nsp4 | CTSZ | -0.424 | 0.000830156 |
| Nsp5 | PPARA | -0.424 | 0.000820397 |
| Nsp5 | PTPN5 | -0.424 | 0.000824323 |
| Nsp6 | CACNG6 | -0.424 | 0.000813053 |
| Nsp6 | NOS3 | -0.424 | 0.000830896 |
| Nsp6 | CSH2 | -0.424 | 0.000825947 |
| Nsp8 | PTPN5 | -0.424 | 0.000815496 |
| Nsp9 | BCL3 | -0.424 | 0.000831099 |
| Nsp10 | PLCG1 | -0.424 | 0.000805851 |
| Nsp10 | MAPK8IP3 | -0.424 | 0.000817077 |
| Nsp12 | IKBKB | -0.424 | 0.00080648 |
| Nsp13 | GNG8 | -0.424 | 0.000822518 |
| Nsp13 | CREB3L3 | -0.424 | 0.00081846 |
| Nsp13 | YWHAH | -0.424 | 0.000830631 |
| Nsp13 | CRK | -0.424 | 0.000810019 |
| Nsp13 | IL36RN | -0.424 | 0.000827377 |
| Nsp14 | PARP3 | -0.424 | 0.000816554 |
| Nsp14 | IL17RE | -0.424 | 0.000824948 |
| Nsp14 | ENO3 | -0.424 | 0.00080717 |
| Nsp15 | CPT1A | -0.424 | 0.000818918 |
| Nsp15 | GH1 | -0.424 | 0.000805616 |
| Nsp15 | ERBB2 | -0.424 | 0.000826784 |
| Nsp16 | SGK2 | -0.424 | 0.000821267 |
| Nsp16 | GDF2 | -0.424 | 0.000808671 |
| Nsp16 | STAT6 | -0.424 | 0.00082239 |
| Orf3a | SRC | -0.424 | 0.000826288 |
| Orf3a | CCND3 | -0.424 | 0.000806059 |
| Orf3a | MMP9 | -0.424 | 0.000828366 |
| Orf3a | INSR | -0.424 | 0.00081087 |
| Orf3a | COL6A1 | -0.424 | 0.000813657 |
| Orf6 | PPP2R2C | -0.424 | 0.000824432 |
| Orf6 | WNT11 | -0.424 | 0.000809589 |
| Orf8 | TUBA3C | -0.424 | 0.000809037 |
| Orf8 | EGFR | -0.424 | 0.000821066 |
| Orf8 | GDF2 | -0.424 | 0.000826121 |
| Orf8 | NFKBIE | -0.424 | 0.000811892 |
| Orf8 | IKBKG | -0.424 | 0.000827764 |
| Orf8 | CACNG2 | -0.424 | 0.000809652 |
| Nsp2 | TAB1 | -0.425 | 0.000783167 |
| Nsp2 | GNB2 | -0.425 | 0.000802002 |
| Nsp2 | ITGB5 | -0.425 | 0.000784285 |
| Nsp2 | ACKR3 | -0.425 | 0.000790884 |

| | | | |
|---|---|---|---|
| Nsp2 | BCAR1 | -0.425 | 0.000786097 |
| Nsp2 | ADCY4 | -0.425 | 0.000786824 |
| Nsp2 | ELOB | -0.425 | 0.000792756 |
| Nsp3 | DVL2 | -0.425 | 0.000786684 |
| Nsp4 | CCL14 | -0.425 | 0.000791339 |
| Nsp4 | WNT8B | -0.425 | 0.000805529 |
| Nsp5 | LTB | -0.425 | 0.00078775 |
| Nsp6 | FZD7 | -0.425 | 0.000801673 |
| Nsp6 | PPP2R2C | -0.425 | 0.00079493 |
| Nsp6 | PIK3CD | -0.425 | 0.000799568 |
| Nsp6 | TNFRSF13C | -0.425 | 0.000792517 |
| Nsp7 | TLR9 | -0.425 | 0.000786314 |
| Nsp7 | WNT10B | -0.425 | 0.000787659 |
| Nsp7 | PPP2R3B | -0.425 | 0.000785599 |
| Nsp7 | NFATC1 | -0.425 | 0.000795756 |
| Nsp8 | NFKBIE | -0.425 | 0.00078186 |
| Nsp9 | PRKAR1B | -0.425 | 0.000801387 |
| Nsp9 | XCR1 | -0.425 | 0.000784224 |
| Nsp10 | IGF1R | -0.425 | 0.00080337 |
| Nsp12 | CALML6 | -0.425 | 0.000800814 |
| Nsp12 | TRAF3 | -0.425 | 0.000804407 |
| Nsp13 | GADD45G | -0.425 | 0.0007949 |
| Nsp13 | NFKBIA | -0.425 | 0.000789073 |
| Nsp14 | CTSF | -0.425 | 0.000802206 |
| Nsp14 | NGF | -0.425 | 0.000786431 |
| Nsp15 | NFKB2 | -0.425 | 0.000794494 |
| Nsp15 | ITGA5 | -0.425 | 0.000785219 |
| Nsp16 | CTSW | -0.425 | 0.000786424 |
| Nsp16 | TUBA3C | -0.425 | 0.000795588 |
| Nsp16 | TUBA3D | -0.425 | 0.000783462 |
| Nsp16 | VAV1 | -0.425 | 0.000791177 |
| S | ELAVL1 | -0.425 | 0.000790683 |
| S | MMP14 | -0.425 | 0.000786252 |
| M | FGF4 | -0.425 | 0.000784371 |
| Orf3a | DAB2IP | -0.425 | 0.000802224 |
| Orf3a | GDF5 | -0.425 | 0.000803144 |
| Orf6 | TNFRSF25 | -0.425 | 0.000781587 |
| Orf6 | RASGRP4 | -0.425 | 0.000800561 |
| Orf6 | FLCN | -0.425 | 0.000785987 |
| Orf6 | PIAS4 | -0.425 | 0.000799588 |
| Orf8 | FGR | -0.425 | 0.000796413 |
| Orf8 | CD14 | -0.425 | 0.000797703 |
| Orf8 | CCL4L2 | -0.425 | 0.000787638 |
| Nsp2 | CACNA2D4 | -0.426 | 0.000762323 |
| Nsp2 | PDGFB | -0.426 | 0.000757404 |
| Nsp2 | THBS2 | -0.426 | 0.000774715 |
| Nsp3 | ACVR1B | -0.426 | 0.000768441 |
| Nsp4 | CREB3L3 | -0.426 | 0.000764604 |
| Nsp4 | PIDD1 | -0.426 | 0.000771262 |
| Nsp5 | PLA2G4D | -0.426 | 0.000774766 |

| | | | |
|---|---|---|---|
| Nsp7 | IRF7 | -0.426 | 0.000759815 |
| Nsp7 | PHLPP1 | -0.426 | 0.000758964 |
| Nsp8 | DVL2 | -0.426 | 0.00077868 |
| Nsp8 | RELA | -0.426 | 0.00077392 |
| Nsp10 | VAV2 | -0.426 | 0.000763894 |
| Nsp10 | ERN1 | -0.426 | 0.000776912 |
| Nsp10 | IRF3 | -0.426 | 0.000774939 |
| Nsp10 | POMC | -0.426 | 0.000760515 |
| Nsp12 | NBL1 | -0.426 | 0.000764283 |
| Nsp12 | IL17RE | -0.426 | 0.000759512 |
| Nsp13 | IRF4 | -0.426 | 0.000778053 |
| Nsp13 | PRKCA | -0.426 | 0.000764477 |
| Nsp13 | ITGA5 | -0.426 | 0.000756204 |
| Nsp14 | TIMP1 | -0.426 | 0.000770541 |
| Nsp14 | CAMK2A | -0.426 | 0.000765598 |
| Nsp15 | ELMO1 | -0.426 | 0.000775991 |
| Nsp16 | CXCR2 | -0.426 | 0.000778954 |
| Nsp16 | LPIN3 | -0.426 | 0.000758248 |
| Nsp16 | CACNG2 | -0.426 | 0.000763248 |
| S | FGFR1 | -0.426 | 0.00077072 |
| Orf3a | WNT3A | -0.426 | 0.000760313 |
| Orf3a | TELO2 | -0.426 | 0.000779418 |
| Orf3a | LIF | -0.426 | 0.00076728 |
| Orf6 | PFKFB3 | -0.426 | 0.00078034 |
| Orf6 | BCL3 | -0.426 | 0.000777202 |
| Orf8 | FGF6 | -0.426 | 0.000776758 |
| Orf8 | IL17RE | -0.426 | 0.000757221 |
| Orf8 | MMP14 | -0.426 | 0.000771767 |
| Orf8 | SOCS7 | -0.426 | 0.000770833 |
| Nsp2 | RAC3 | -0.427 | 0.000748207 |
| Nsp2 | SREBF1 | -0.427 | 0.00075013 |
| Nsp3 | IFNL1 | -0.427 | 0.000743404 |
| Nsp3 | HRAS | -0.427 | 0.000755968 |
| Nsp3 | IKBKG | -0.427 | 0.000739791 |
| Nsp4 | TNFRSF25 | -0.427 | 0.00074948 |
| Nsp4 | MAP2K3 | -0.427 | 0.000744834 |
| Nsp4 | CTSW | -0.427 | 0.0007556 |
| Nsp4 | S100A9 | -0.427 | 0.000745545 |
| Nsp4 | CSF3R | -0.427 | 0.000753725 |
| Nsp5 | GFAP | -0.427 | 0.000732931 |
| Nsp5 | FGF16 | -0.427 | 0.000738837 |
| Nsp6 | TNFRSF4 | -0.427 | 0.00074802 |
| Nsp7 | FGF18 | -0.427 | 0.000742392 |
| Nsp7 | NFKBIB | -0.427 | 0.000736493 |
| Nsp7 | DUSP7 | -0.427 | 0.000737483 |
| Nsp7 | WNT11 | -0.427 | 0.000755734 |
| Nsp7 | CARD11 | -0.427 | 0.000748895 |
| Nsp8 | CSF1R | -0.427 | 0.000742848 |
| Nsp9 | CACNA1H | -0.427 | 0.000745043 |
| Nsp9 | EPOR | -0.427 | 0.000734917 |

| | | | |
|---|---|---|---|
| Nsp9 | EFNA3 | -0.427 | 0.000733463 |
| Nsp9 | TICAM1 | -0.427 | 0.000743683 |
| Nsp10 | GNB2 | -0.427 | 0.000748494 |
| Nsp10 | MMP14 | -0.427 | 0.000753983 |
| Nsp10 | ID4 | -0.427 | 0.000747449 |
| Nsp12 | AKT1S1 | -0.427 | 0.000747363 |
| Nsp12 | TGFA | -0.427 | 0.000733547 |
| Nsp12 | TCL1B | -0.427 | 0.000750533 |
| Nsp12 | LPIN3 | -0.427 | 0.000753204 |
| Nsp12 | IKBKG | -0.427 | 0.000745704 |
| Nsp13 | IKBKB | -0.427 | 0.000742697 |
| Nsp13 | PRL | -0.427 | 0.000754979 |
| Nsp14 | IRS1 | -0.427 | 0.000735828 |
| Nsp14 | NPRL2 | -0.427 | 0.000754152 |
| Nsp15 | LEP | -0.427 | 0.000753544 |
| Nsp15 | LAMTOR1 | -0.427 | 0.00074645 |
| Nsp16 | TUBA8 | -0.427 | 0.000749224 |
| Nsp16 | GAPDH | -0.427 | 0.000741409 |
| S | GNB5 | -0.427 | 0.000750958 |
| Orf3a | ITPR3 | -0.427 | 0.000749295 |
| Orf3a | CXCR3 | -0.427 | 0.000732703 |
| Orf3a | CCR10 | -0.427 | 0.000742621 |
| Orf3a | FGF17 | -0.427 | 0.000748261 |
| Orf3a | RPS6KB2 | -0.427 | 0.000733086 |
| Orf3a | PPP1R3E | -0.427 | 0.000737451 |
| Orf3a | BCAR1 | -0.427 | 0.000747506 |
| Orf6 | IRS1 | -0.427 | 0.000754234 |
| Orf6 | MMP9 | -0.427 | 0.00074705 |
| Orf6 | LRP5 | -0.427 | 0.000735702 |
| Orf6 | EFNA2 | -0.427 | 0.000739303 |
| Orf6 | GDF11 | -0.427 | 0.000754402 |
| Orf6 | FGF23 | -0.427 | 0.000745995 |
| Orf6 | FOXO3 | -0.427 | 0.000753849 |
| Orf8 | BMP4 | -0.427 | 0.00075155 |
| Orf10 | ITGA2B | -0.427 | 0.000740045 |
| Nsp2 | DVL3 | -0.428 | 0.000723526 |
| Nsp2 | MLST8 | -0.428 | 0.00072773 |
| Nsp2 | FGF23 | -0.428 | 0.000719337 |
| Nsp3 | ELOB | -0.428 | 0.000721362 |
| Nsp4 | IRF4 | -0.428 | 0.000719833 |
| Nsp5 | IL27 | -0.428 | 0.000724758 |
| Nsp5 | TUBA3C | -0.428 | 0.0007119 |
| Nsp6 | GRK2 | -0.428 | 0.000719074 |
| Nsp7 | CACNG6 | -0.428 | 0.000718202 |
| Nsp7 | FGFR4 | -0.428 | 0.000728288 |
| Nsp7 | MAPK8IP1 | -0.428 | 0.000725601 |
| Nsp8 | ELMO1 | -0.428 | 0.000732394 |
| Nsp8 | ERN1 | -0.428 | 0.00071561 |
| Nsp8 | EDA | -0.428 | 0.000730457 |
| Nsp8 | ERBB2 | -0.428 | 0.000726424 |

| | | | |
|---|---|---|---|
| Nsp9 | CCR10 | -0.428 | 0.000725023 |
| Nsp10 | DUSP5 | -0.428 | 0.000714896 |
| Nsp10 | JUND | -0.428 | 0.000728522 |
| Nsp12 | ELOB | -0.428 | 0.00071675 |
| Nsp13 | MAP2K3 | -0.428 | 0.000714194 |
| Nsp13 | IRF5 | -0.428 | 0.000725368 |
| Nsp13 | TNFRSF11A | -0.428 | 0.000731321 |
| Nsp14 | RNF152 | -0.428 | 0.000715376 |
| Nsp14 | SLC2A1 | -0.428 | 0.000713299 |
| Nsp15 | CSF3 | -0.428 | 0.000727838 |
| Nsp16 | ADCY8 | -0.428 | 0.000730067 |
| Nsp16 | IL10RA | -0.428 | 0.00071413 |
| S | IRF5 | -0.428 | 0.000722009 |
| Orf3a | FGFR4 | -0.428 | 0.000731455 |
| Orf3a | TNFRSF4 | -0.428 | 0.000721106 |
| Orf3a | CACNA1B | -0.428 | 0.000723324 |
| Orf3a | RARA | -0.428 | 0.000711877 |
| Orf3a | ADCY6 | -0.428 | 0.000732332 |
| Orf6 | PTPN6 | -0.428 | 0.000726252 |
| Orf6 | LPAR5 | -0.428 | 0.000731038 |
| Orf6 | WNT10A | -0.428 | 0.000713924 |
| Orf6 | PPP2R3B | -0.428 | 0.000709754 |
| Orf8 | IRF4 | -0.428 | 0.000717297 |
| Orf8 | ERN1 | -0.428 | 0.000712743 |
| Orf10 | TGFB3 | -0.428 | 0.000711563 |
| Nsp2 | TLR9 | -0.429 | 0.000691961 |
| Nsp2 | FOXP3 | -0.429 | 0.000709045 |
| Nsp2 | RUNX1 | -0.429 | 0.000696179 |
| Nsp3 | CXCL1 | -0.429 | 0.000703197 |
| Nsp4 | CRK | -0.429 | 0.000698319 |
| Nsp5 | IRAK1 | -0.429 | 0.000687286 |
| Nsp5 | IL27RA | -0.429 | 0.000700174 |
| Nsp6 | GDF6 | -0.429 | 0.000708093 |
| Nsp6 | FZD2 | -0.429 | 0.00070268 |
| Nsp8 | NFKB2 | -0.429 | 0.000704857 |
| Nsp9 | SLC7A5 | -0.429 | 0.000687625 |
| Nsp10 | TGFB3 | -0.429 | 0.000700466 |
| Nsp10 | HCK | -0.429 | 0.00070551 |
| Nsp10 | GYS1 | -0.429 | 0.000693396 |
| Nsp10 | JUNB | -0.429 | 0.000704115 |
| Nsp10 | MAPK11 | -0.429 | 0.000690752 |
| Nsp10 | IL9R | -0.429 | 0.000698872 |
| Nsp14 | IL11 | -0.429 | 0.00069229 |
| Nsp14 | HK3 | -0.429 | 0.00070492 |
| Nsp14 | ITGA7 | -0.429 | 0.00069241 |
| Nsp15 | CDC37 | -0.429 | 0.000704478 |
| Nsp15 | VTN | -0.429 | 0.000704397 |
| Nsp15 | FGF23 | -0.429 | 0.000695392 |
| Nsp15 | EGLN2 | -0.429 | 0.000689653 |
| Nsp15 | HK2 | -0.429 | 0.000698794 |

| | | | |
|---|---|---|---|
| Nsp15 | SLC2A1 | -0.429 | 0.000701274 |
| Nsp16 | COL9A3 | -0.429 | 0.000706942 |
| Nsp16 | MAX | -0.429 | 0.000702302 |
| S | ELK1 | -0.429 | 0.00069748 |
| S | PARP3 | -0.429 | 0.000689993 |
| S | ITGA11 | -0.429 | 0.000692768 |
| Orf3a | MAP3K6 | -0.429 | 0.000699118 |
| Orf3a | LCK | -0.429 | 0.000698185 |
| Orf6 | CACNG7 | -0.429 | 0.000701113 |
| Orf6 | CEBPB | -0.429 | 0.000702414 |
| Orf6 | IRF7 | -0.429 | 0.000702709 |
| Orf6 | DUSP2 | -0.429 | 0.000689524 |
| Orf6 | FZD2 | -0.429 | 0.000701759 |
| Orf8 | ACVRL1 | -0.429 | 0.000709257 |
| Orf10 | TGIF2 | -0.429 | 0.000694063 |
| Orf10 | CRTC2 | -0.429 | 0.000693399 |
| Nsp2 | CACNA1C | -0.43 | 0.000665957 |
| Nsp3 | TNFRSF13B | -0.43 | 0.000673003 |
| Nsp4 | DFFB | -0.43 | 0.000673705 |
| Nsp4 | IFNL1 | -0.43 | 0.000670726 |
| Nsp4 | TAOK2 | -0.43 | 0.000676182 |
| Nsp4 | GADD45G | -0.43 | 0.000681491 |
| Nsp4 | ELOB | -0.43 | 0.000669682 |
| Nsp5 | EDAR | -0.43 | 0.000675284 |
| Nsp6 | SMAD7 | -0.43 | 0.000684266 |
| Nsp6 | GDF10 | -0.43 | 0.000672545 |
| Nsp6 | HRK | -0.43 | 0.000680092 |
| Nsp6 | FGFR3 | -0.43 | 0.000671523 |
| Nsp7 | ISG15 | -0.43 | 0.000674039 |
| Nsp7 | WNT6 | -0.43 | 0.00068024 |
| Nsp7 | SLC7A5 | -0.43 | 0.000669254 |
| Nsp7 | SH2B2 | -0.43 | 0.000683902 |
| Nsp8 | CSF2 | -0.43 | 0.000677402 |
| Nsp8 | FGF16 | -0.43 | 0.000677514 |
| Nsp9 | IL17D | -0.43 | 0.000682541 |
| Nsp9 | WNT1 | -0.43 | 0.000680181 |
| Nsp10 | COL9A3 | -0.43 | 0.000666604 |
| Nsp10 | CACNG1 | -0.43 | 0.000672584 |
| Nsp10 | CXCR3 | -0.43 | 0.000666007 |
| Nsp10 | PRKCD | -0.43 | 0.000666965 |
| Nsp10 | DUSP9 | -0.43 | 0.000672056 |
| Nsp10 | RPS6KB2 | -0.43 | 0.000674072 |
| Nsp12 | HKDC1 | -0.43 | 0.000682302 |
| Nsp12 | SOCS7 | -0.43 | 0.000677657 |
| Nsp14 | PDGFA | -0.43 | 0.000667054 |
| Nsp14 | PF4V1 | -0.43 | 0.000671521 |
| Nsp14 | SMAD3 | -0.43 | 0.000685177 |
| Nsp14 | FADD | -0.43 | 0.000668705 |
| Nsp15 | IL25 | -0.43 | 0.000683436 |
| Nsp15 | COL9A3 | -0.43 | 0.000671408 |

| | | | |
|---|---|---|---|
| Nsp15 | IRF3 | -0.43 | 0.000677103 |
| Nsp16 | MAVS | -0.43 | 0.000669603 |
| Orf3a | FLT3LG | -0.43 | 0.000682729 |
| Orf3a | WNT9B | -0.43 | 0.0006767 |
| Orf3a | MAPK4 | -0.43 | 0.000682083 |
| Orf6 | FGF18 | -0.43 | 0.000678211 |
| Orf6 | SH2B2 | -0.43 | 0.000685566 |
| Orf6 | NOG | -0.43 | 0.000672525 |
| Orf8 | PGF | -0.43 | 0.000680925 |
| Orf8 | MAPK12 | -0.43 | 0.000683791 |
| Orf8 | GH2 | -0.43 | 0.000681031 |
| Orf8 | PKN3 | -0.43 | 0.000685334 |
| Orf8 | PTPN5 | -0.43 | 0.000686896 |
| Orf8 | GNB3 | -0.43 | 0.000667896 |
| Orf8 | LPAR2 | -0.43 | 0.000680494 |
| Orf10 | HSPA6 | -0.43 | 0.00068417 |
| Nsp2 | ATP6V1B1 | -0.431 | 0.000659449 |
| Nsp2 | TOLLIP | -0.431 | 0.00064516 |
| Nsp2 | MAPK13 | -0.431 | 0.000662629 |
| Nsp2 | ID1 | -0.431 | 0.000651065 |
| Nsp2 | TYK2 | -0.431 | 0.000650905 |
| Nsp4 | CACNA1F | -0.431 | 0.000662073 |
| Nsp5 | NFKBIA | -0.431 | 0.000659733 |
| Nsp6 | CACNA1H | -0.431 | 0.000664922 |
| Nsp6 | CCND1 | -0.431 | 0.000653617 |
| Nsp7 | EPHA2 | -0.431 | 0.000664052 |
| Nsp7 | MAPK12 | -0.431 | 0.000655384 |
| Nsp7 | CCR10 | -0.431 | 0.000656534 |
| Nsp7 | WDR24 | -0.431 | 0.000656118 |
| Nsp7 | POMC | -0.431 | 0.000651332 |
| Nsp8 | PPP1R3F | -0.431 | 0.000655218 |
| Nsp9 | MMP9 | -0.431 | 0.000661021 |
| Nsp9 | WNT9A | -0.431 | 0.000644695 |
| Nsp9 | FGF19 | -0.431 | 0.000656646 |
| Nsp9 | RASGRP2 | -0.431 | 0.00065088 |
| Nsp9 | SH2B2 | -0.431 | 0.000648296 |
| Nsp10 | ARAF | -0.431 | 0.000663908 |
| Nsp10 | GDF6 | -0.431 | 0.000661498 |
| Nsp10 | PFKFB3 | -0.431 | 0.000653606 |
| Nsp10 | TNFRSF1B | -0.431 | 0.000651332 |
| Nsp12 | CSF1R | -0.431 | 0.000653436 |
| Nsp13 | ACVR1B | -0.431 | 0.00064635 |
| Nsp14 | TAOK2 | -0.431 | 0.000659354 |
| Nsp14 | GFAP | -0.431 | 0.000661739 |
| Nsp14 | HRAS | -0.431 | 0.000655695 |
| Nsp15 | ELK1 | -0.431 | 0.000662044 |
| Nsp15 | FGF6 | -0.431 | 0.000658834 |
| Nsp16 | ARRB1 | -0.431 | 0.000662574 |
| Nsp16 | ID2 | -0.431 | 0.000648452 |
| S | CTSF | -0.431 | 0.000653433 |

| | | | |
|---|---|---|---|
| S | G6PC3 | -0.431 | 0.000650056 |
| Orf3a | CACNG4 | -0.431 | 0.000663544 |
| Orf3a | DUSP5 | -0.431 | 0.000661026 |
| Orf3a | GDF6 | -0.431 | 0.000661614 |
| Orf3a | S100A9 | -0.431 | 0.00065044 |
| Orf6 | GDF7 | -0.431 | 0.000654102 |
| Orf6 | BCAR1 | -0.431 | 0.000646124 |
| Orf8 | CXCR2 | -0.431 | 0.000650373 |
| Orf8 | RORC | -0.431 | 0.000657023 |
| Orf8 | NODAL | -0.431 | 0.000655856 |
| Orf8 | PDGFRB | -0.431 | 0.000661145 |
| Orf10 | TNFRSF10D | -0.431 | 0.000645527 |
| Orf10 | TNFRSF8 | -0.431 | 0.000662042 |
| Nsp2 | NTRK1 | -0.432 | 0.000631369 |
| Nsp2 | PDGFA | -0.432 | 0.000641508 |
| Nsp2 | FMOD | -0.432 | 0.000637345 |
| Nsp2 | PPP5C | -0.432 | 0.000631059 |
| Nsp2 | ADCY3 | -0.432 | 0.000644082 |
| Nsp3 | MAP4K1 | -0.432 | 0.000629216 |
| Nsp4 | PARP3 | -0.432 | 0.000638584 |
| Nsp4 | GDF2 | -0.432 | 0.000626171 |
| Nsp4 | MRAS | -0.432 | 0.000629701 |
| Nsp5 | PARP3 | -0.432 | 0.000641681 |
| Nsp5 | VAV2 | -0.432 | 0.000631228 |
| Nsp5 | SHC3 | -0.432 | 0.000637583 |
| Nsp6 | NCF1 | -0.432 | 0.000631092 |
| Nsp7 | CACNA1B | -0.432 | 0.000628436 |
| Nsp7 | BCAR1 | -0.432 | 0.000629295 |
| Nsp9 | BMP8B | -0.432 | 0.000640796 |
| Nsp12 | CREBBP | -0.432 | 0.000632667 |
| Nsp12 | TNFRSF13B | -0.432 | 0.000630606 |
| Nsp13 | PARP3 | -0.432 | 0.000637916 |
| Nsp13 | RELA | -0.432 | 0.00064357 |
| Nsp13 | PF4 | -0.432 | 0.000634097 |
| Nsp14 | TGFB3 | -0.432 | 0.000625886 |
| Nsp14 | INS | -0.432 | 0.000627189 |
| Nsp14 | HK2 | -0.432 | 0.000640552 |
| Nsp15 | NBL1 | -0.432 | 0.000638614 |
| Nsp15 | IL4R | -0.432 | 0.000636486 |
| Nsp15 | FGF16 | -0.432 | 0.000626119 |
| Nsp15 | IKBKG | -0.432 | 0.000641164 |
| Nsp16 | IFNL1 | -0.432 | 0.000642146 |
| Nsp16 | IL13 | -0.432 | 0.000633774 |
| S | ACTB | -0.432 | 0.000640934 |
| Orf3a | RGMA | -0.432 | 0.000627078 |
| Orf3a | WNT9A | -0.432 | 0.000624687 |
| Orf3a | CTF1 | -0.432 | 0.000626791 |
| Orf3a | SOCS3 | -0.432 | 0.000630477 |
| Orf3a | FGF8 | -0.432 | 0.000639511 |
| Orf8 | GREM1 | -0.432 | 0.000636801 |

| | | | |
|---|---|---|---|
| Orf8 | PLA2G4F | -0.432 | 0.000639961 |
| Nsp2 | TNFRSF8 | -0.433 | 0.000614738 |
| Nsp4 | HRAS | -0.433 | 0.000609646 |
| Nsp4 | VEGFB | -0.433 | 0.000617937 |
| Nsp4 | PIK3R6 | -0.433 | 0.000622481 |
| Nsp6 | CSH1 | -0.433 | 0.000617487 |
| Nsp7 | DUSP4 | -0.433 | 0.000621453 |
| Nsp7 | BMP8A | -0.433 | 0.000609183 |
| Nsp7 | VHL | -0.433 | 0.000609318 |
| Nsp8 | BMP4 | -0.433 | 0.000609759 |
| Nsp9 | DVL1 | -0.433 | 0.000617044 |
| Nsp9 | PRKCG | -0.433 | 0.000611452 |
| Nsp10 | TNXB | -0.433 | 0.000623726 |
| Nsp10 | DVL3 | -0.433 | 0.000609359 |
| Nsp10 | TNFRSF12A | -0.433 | 0.000606649 |
| Nsp10 | THBS2 | -0.433 | 0.000609906 |
| Nsp12 | FGF2 | -0.433 | 0.000610181 |
| Nsp12 | RRAGA | -0.433 | 0.000610911 |
| Nsp13 | EGFR | -0.433 | 0.00061432 |
| Nsp13 | CXCL1 | -0.433 | 0.000613662 |
| Nsp13 | LPIN3 | -0.433 | 0.000611301 |
| Nsp14 | NTRK1 | -0.433 | 0.000606483 |
| Nsp14 | IL17B | -0.433 | 0.000612731 |
| Nsp14 | PRKACA | -0.433 | 0.000622421 |
| Nsp14 | VWF | -0.433 | 0.000622389 |
| Nsp15 | IL11 | -0.433 | 0.000615573 |
| Nsp15 | CACNA1F | -0.433 | 0.000610227 |
| S | RNF152 | -0.433 | 0.000621539 |
| S | VEGFB | -0.433 | 0.000607439 |
| Orf3a | PIK3R2 | -0.433 | 0.00061631 |
| Orf3a | WNT10A | -0.433 | 0.000614794 |
| Orf6 | MAP3K11 | -0.433 | 0.000618944 |
| Orf6 | FGF4 | -0.433 | 0.00061739 |
| Orf6 | GNAI2 | -0.433 | 0.000608581 |
| Orf6 | PPP1R3E | -0.433 | 0.000620672 |
| Orf8 | NR4A1 | -0.433 | 0.000618923 |
| Orf8 | SHC3 | -0.433 | 0.000616605 |
| Nsp3 | SHC3 | -0.434 | 0.00059628 |
| Nsp4 | BMP4 | -0.434 | 0.000596627 |
| Nsp4 | VAV1 | -0.434 | 0.000602889 |
| Nsp5 | CACNA1S | -0.434 | 0.000593921 |
| Nsp5 | CACNB3 | -0.434 | 0.000590587 |
| Nsp5 | IL17B | -0.434 | 0.00058885 |
| Nsp5 | CXCL3 | -0.434 | 0.000594752 |
| Nsp5 | FOSL1 | -0.434 | 0.000589448 |
| Nsp5 | IL21R | -0.434 | 0.000596322 |
| Nsp5 | INS | -0.434 | 0.000586429 |
| Nsp5 | VWF | -0.434 | 0.000594883 |
| Nsp6 | BMP7 | -0.434 | 0.000592645 |
| Nsp6 | BCL3 | -0.434 | 0.000588029 |

| | | | |
|---|---|---|---|
| Nsp6 | EGLN1 | -0.434 | 0.000592466 |
| Nsp7 | TELO2 | -0.434 | 0.000588304 |
| Nsp7 | CD70 | -0.434 | 0.000591901 |
| Nsp7 | WNT5A | -0.434 | 0.000587324 |
| Nsp10 | ENDOG | -0.434 | 0.000599516 |
| Nsp10 | CAMK2B | -0.434 | 0.00060019 |
| Nsp10 | LIF | -0.434 | 0.000590095 |
| Nsp10 | AMH | -0.434 | 0.000587592 |
| Nsp11 | CCL3L1 | -0.434 | 0.00058817 |
| Nsp11 | CCL3L3 | -0.434 | 0.00058817 |
| Nsp12 | ITGA5 | -0.434 | 0.000594367 |
| Nsp12 | NFKBIE | -0.434 | 0.000586004 |
| Nsp14 | NFKB2 | -0.434 | 0.000602867 |
| Nsp14 | GH2 | -0.434 | 0.000588252 |
| Nsp15 | CAMK2B | -0.434 | 0.000594745 |
| S | IL17RE | -0.434 | 0.000590973 |
| S | ITGA7 | -0.434 | 0.0005977 |
| S | FOXP3 | -0.434 | 0.000585729 |
| E | NTF3 | -0.434 | 0.000597722 |
| Orf6 | WNT6 | -0.434 | 0.000603213 |
| Orf6 | WNT9A | -0.434 | 0.000589605 |
| Orf8 | ID3 | -0.434 | 0.000588126 |
| Nsp2 | RAC2 | -0.435 | 0.000584359 |
| Nsp2 | TSC2 | -0.435 | 0.000582748 |
| Nsp3 | DFFB | -0.435 | 0.00058002 |
| Nsp4 | CREBBP | -0.435 | 0.000567862 |
| Nsp4 | LBP | -0.435 | 0.000575095 |
| Nsp5 | RELT | -0.435 | 0.000573007 |
| Nsp6 | PGAM5 | -0.435 | 0.00057032 |
| Nsp6 | YWHAG | -0.435 | 0.000582122 |
| Nsp6 | MCL1 | -0.435 | 0.000570913 |
| Nsp8 | FOS | -0.435 | 0.000579383 |
| Nsp8 | PRKACA | -0.435 | 0.000577844 |
| Nsp9 | GRK2 | -0.435 | 0.000570379 |
| Nsp9 | RAC3 | -0.435 | 0.000568317 |
| Nsp9 | VEGFA | -0.435 | 0.000568886 |
| Nsp10 | PTPN6 | -0.435 | 0.000581997 |
| Nsp10 | PIK3R2 | -0.435 | 0.000571315 |
| Nsp10 | COL6A2 | -0.435 | 0.00058054 |
| Nsp10 | TNFRSF13B | -0.435 | 0.000576277 |
| Nsp10 | CDKN1B | -0.435 | 0.000580577 |
| Nsp12 | PF4 | -0.435 | 0.000579484 |
| Nsp13 | NLRX1 | -0.435 | 0.000566766 |
| Nsp14 | ELAVL1 | -0.435 | 0.000571457 |
| Nsp14 | RPS6KA2 | -0.435 | 0.000571097 |
| Nsp14 | CPT1B | -0.435 | 0.000566903 |
| Nsp14 | GNB5 | -0.435 | 0.000572635 |
| Nsp14 | IFNL2 | -0.435 | 0.000575713 |
| Nsp15 | TNFRSF1A | -0.435 | 0.000584054 |
| Nsp16 | TGFA | -0.435 | 0.000568558 |

| | | | |
|---|---|---|---|
| Nsp16 | FGF21 | -0.435 | 0.000578002 |
| S | BBC3 | -0.435 | 0.000581606 |
| Orf6 | GREM2 | -0.435 | 0.000574838 |
| Orf8 | NOS2 | -0.435 | 0.000568912 |
| Orf8 | STAT5A | -0.435 | 0.000583468 |
| Orf8 | PTK2B | -0.435 | 0.000566823 |
| Orf8 | CSF2 | -0.435 | 0.000566453 |
| Orf8 | ADCY2 | -0.435 | 0.00058387 |
| Orf10 | RNF152 | -0.435 | 0.00057442 |
| Nsp2 | PGF | -0.436 | 0.00055289 |
| Nsp2 | S100A9 | -0.436 | 0.000550505 |
| Nsp2 | WNT7A | -0.436 | 0.000548026 |
| Nsp3 | CTSZ | -0.436 | 0.000560527 |
| Nsp3 | ITGA5 | -0.436 | 0.000560682 |
| Nsp4 | NTF3 | -0.436 | 0.000560056 |
| Nsp4 | ACVR2B | -0.436 | 0.000561693 |
| Nsp5 | ERN1 | -0.436 | 0.000557482 |
| Nsp5 | CACNA1E | -0.436 | 0.000563003 |
| Nsp5 | TCL1A | -0.436 | 0.00055744 |
| Nsp6 | LRP5 | -0.436 | 0.000556026 |
| Nsp6 | BMP8B | -0.436 | 0.000551724 |
| Nsp7 | IL34 | -0.436 | 0.000550259 |
| Nsp7 | WNT7B | -0.436 | 0.000560218 |
| Nsp7 | BMP6 | -0.436 | 0.000565312 |
| Nsp7 | LAMA5 | -0.436 | 0.000562905 |
| Nsp8 | CAMK2B | -0.436 | 0.000551634 |
| Nsp8 | SOCS7 | -0.436 | 0.000554598 |
| Nsp9 | IL27 | -0.436 | 0.000562836 |
| Nsp9 | CHAD | -0.436 | 0.000554599 |
| Nsp9 | DUSP4 | -0.436 | 0.000550957 |
| Nsp9 | RELB | -0.436 | 0.000557039 |
| Nsp10 | GDF5 | -0.436 | 0.000558237 |
| Nsp12 | HMOX1 | -0.436 | 0.000556044 |
| Nsp12 | LPAR2 | -0.436 | 0.000562716 |
| Nsp13 | CAMK2B | -0.436 | 0.000566004 |
| Nsp14 | PKN3 | -0.436 | 0.000563646 |
| Nsp14 | CXCL3 | -0.436 | 0.000556887 |
| Nsp14 | GNB3 | -0.436 | 0.000552187 |
| Nsp15 | WNT8B | -0.436 | 0.00054912 |
| Nsp16 | IL11 | -0.436 | 0.000554026 |
| S | NLRX1 | -0.436 | 0.000556618 |
| S | FGR | -0.436 | 0.000564503 |
| S | EIF4E1B | -0.436 | 0.000556276 |
| M | SHC2 | -0.436 | 0.000554861 |
| M | FGF3 | -0.436 | 0.00055228 |
| Orf3a | FGF18 | -0.436 | 0.000565882 |
| Orf3a | MAPK11 | -0.436 | 0.000555399 |
| Orf3a | IL17RA | -0.436 | 0.00055013 |
| Orf6 | MAPK8IP2 | -0.436 | 0.000558131 |
| Orf8 | NFKBIB | -0.436 | 0.00056314 |

| | | | |
|---|---|---|---|
| Orf8 | CTSZ | -0.436 | 0.000557086 |
| Orf8 | DHX58 | -0.436 | 0.000561075 |
| Orf8 | IL11RA | -0.436 | 0.000549084 |
| Nsp2 | TNFSF12 | -0.437 | 0.000547345 |
| Nsp2 | PLA2G4D | -0.437 | 0.000538187 |
| Nsp3 | PGF | -0.437 | 0.000540696 |
| Nsp3 | PARP3 | -0.437 | 0.000544174 |
| Nsp3 | PPP2R5B | -0.437 | 0.000531534 |
| Nsp4 | NFKB2 | -0.437 | 0.000531275 |
| Nsp5 | CTSF | -0.437 | 0.000532326 |
| Nsp5 | ACACB | -0.437 | 0.000543356 |
| Nsp6 | FLT4 | -0.437 | 0.000546382 |
| Nsp7 | PRKCG | -0.437 | 0.000532668 |
| Nsp7 | PDGFRB | -0.437 | 0.000541836 |
| Nsp8 | TGFB3 | -0.437 | 0.000543269 |
| Nsp8 | ITGA5 | -0.437 | 0.000546055 |
| Nsp9 | DUSP5 | -0.437 | 0.000547504 |
| Nsp9 | BMP6 | -0.437 | 0.000539014 |
| Nsp9 | COMP | -0.437 | 0.000531545 |
| Nsp9 | WNT10A | -0.437 | 0.000532012 |
| Nsp9 | STK11 | -0.437 | 0.000542544 |
| Nsp9 | SOCS3 | -0.437 | 0.000534628 |
| Nsp10 | IRF4 | -0.437 | 0.000537998 |
| Nsp10 | FGR | -0.437 | 0.000541016 |
| Nsp10 | WNT9A | -0.437 | 0.000542419 |
| Nsp10 | PPP1R3E | -0.437 | 0.000533465 |
| Nsp10 | FGFR3 | -0.437 | 0.000540983 |
| Nsp12 | CCR7 | -0.437 | 0.000539315 |
| Nsp12 | CDC37 | -0.437 | 0.0005393 |
| Nsp13 | ATP6V1F | -0.437 | 0.000539985 |
| Nsp13 | EIF4EBP1 | -0.437 | 0.000540244 |
| Nsp14 | DUSP3 | -0.437 | 0.000545264 |
| Nsp14 | MAPK12 | -0.437 | 0.000535527 |
| Nsp14 | IKBKE | -0.437 | 0.000543772 |
| Nsp14 | ERBB2 | -0.437 | 0.000544505 |
| Nsp15 | IRF5 | -0.437 | 0.000545463 |
| Nsp15 | NLRX1 | -0.437 | 0.000536201 |
| Nsp15 | ARRB2 | -0.437 | 0.000537354 |
| Nsp15 | CACNB3 | -0.437 | 0.000539118 |
| Nsp15 | EDA | -0.437 | 0.000538994 |
| Nsp15 | MAX | -0.437 | 0.000539242 |
| Nsp15 | CNTFR | -0.437 | 0.000547753 |
| Nsp16 | IL25 | -0.437 | 0.000531778 |
| Nsp16 | ERN1 | -0.437 | 0.000542288 |
| Nsp16 | LAMTOR1 | -0.437 | 0.000532932 |
| S | PRKAB1 | -0.437 | 0.000544008 |
| S | CACNB3 | -0.437 | 0.000539841 |
| Orf3a | SLC7A5 | -0.437 | 0.000544696 |
| Orf6 | TNFRSF6B | -0.437 | 0.000541883 |
| Orf8 | ELK1 | -0.437 | 0.000537797 |

| | | | |
|---|---|---|---|
| Orf8 | S100A7A | -0.437 | 0.000545224 |
| Orf8 | CSF1 | -0.437 | 0.00053885 |
| Orf8 | IL12RB1 | -0.437 | 0.000547625 |
| Orf8 | LAMTOR2 | -0.437 | 0.000534322 |
| Orf8 | ERBB2 | -0.437 | 0.000539017 |
| Nsp2 | PREX1 | -0.438 | 0.000519039 |
| Nsp2 | DHX58 | -0.438 | 0.000519993 |
| Nsp2 | TNFRSF1B | -0.438 | 0.000528306 |
| Nsp2 | PXN | -0.438 | 0.000528742 |
| Nsp3 | ELMO1 | -0.438 | 0.000529595 |
| Nsp4 | TUBA3D | -0.438 | 0.000515716 |
| Nsp4 | GH1 | -0.438 | 0.000526321 |
| Nsp4 | ACTB | -0.438 | 0.000524293 |
| Nsp7 | CACNA1H | -0.438 | 0.000522229 |
| Nsp7 | NOS3 | -0.438 | 0.000524391 |
| Nsp7 | RPTOR | -0.438 | 0.000528804 |
| Nsp7 | PDGFB | -0.438 | 0.000525432 |
| Nsp7 | ADCY7 | -0.438 | 0.000519254 |
| Nsp9 | EGLN1 | -0.438 | 0.000515837 |
| Nsp9 | PTPN7 | -0.438 | 0.0005258 |
| Nsp11 | GNG5 | -0.438 | 0.000521373 |
| Nsp12 | CCND2 | -0.438 | 0.000521392 |
| Nsp12 | DVL2 | -0.438 | 0.000525283 |
| Nsp12 | CSF3 | -0.438 | 0.000523868 |
| Nsp13 | DVL2 | -0.438 | 0.000518957 |
| Nsp13 | RORC | -0.438 | 0.000523792 |
| Nsp14 | MAP2K3 | -0.438 | 0.000515994 |
| Nsp14 | IFNL3 | -0.438 | 0.000516463 |
| Nsp14 | ITGB5 | -0.438 | 0.000515721 |
| Nsp14 | ID3 | -0.438 | 0.000514628 |
| Nsp15 | HMOX1 | -0.438 | 0.000522377 |
| Nsp15 | CTSF | -0.438 | 0.000514709 |
| Nsp15 | MMP14 | -0.438 | 0.000515839 |
| Nsp16 | NOD2 | -0.438 | 0.000516093 |
| Nsp16 | LAMTOR2 | -0.438 | 0.00052356 |
| S | S100A9 | -0.438 | 0.00052425 |
| Orf3a | YWHAG | -0.438 | 0.00052985 |
| Orf3a | NOS3 | -0.438 | 0.000527662 |
| Orf3a | WNT3 | -0.438 | 0.000519239 |
| Orf6 | TNFRSF1B | -0.438 | 0.000524344 |
| Orf8 | PPP2R1A | -0.438 | 0.000523066 |
| Orf8 | PIK3R6 | -0.438 | 0.000521566 |
| Nsp2 | NFATC2 | -0.439 | 0.000504652 |
| Nsp2 | WNT11 | -0.439 | 0.000507904 |
| Nsp2 | CXCL14 | -0.439 | 0.000508911 |
| Nsp3 | MAP2K3 | -0.439 | 0.000506048 |
| Nsp3 | RPS6KA1 | -0.439 | 0.000507605 |
| Nsp4 | TGFB3 | -0.439 | 0.000511463 |
| Nsp4 | ADCY8 | -0.439 | 0.000505125 |
| Nsp4 | ARRB2 | -0.439 | 0.00050421 |

| | | | |
|---|---|---|---|
| Nsp4 | HK2 | -0.439 | 0.000508146 |
| Nsp5 | DFFB | -0.439 | 0.000502238 |
| Nsp5 | PRF1 | -0.439 | 0.000510399 |
| Nsp5 | ACVR2B | -0.439 | 0.000510864 |
| Nsp5 | RPS6KA1 | -0.439 | 0.000499137 |
| Nsp5 | CISH | -0.439 | 0.000497966 |
| Nsp7 | FZD10 | -0.439 | 0.000504487 |
| Nsp7 | BMP7 | -0.439 | 0.00050181 |
| Nsp7 | DHX58 | -0.439 | 0.000497047 |
| Nsp8 | CHRD | -0.439 | 0.000512055 |
| Nsp8 | FOXO1 | -0.439 | 0.000509901 |
| Nsp8 | CXCL3 | -0.439 | 0.000504011 |
| Nsp8 | LAMTOR2 | -0.439 | 0.000509348 |
| Nsp10 | CACNA1C | -0.439 | 0.000507768 |
| Nsp10 | FOXO1 | -0.439 | 0.000501504 |
| Nsp10 | NOG | -0.439 | 0.000497025 |
| Nsp12 | IRS1 | -0.439 | 0.000509273 |
| Nsp12 | DFFB | -0.439 | 0.000505674 |
| Nsp12 | WNT8B | -0.439 | 0.00050665 |
| Nsp12 | FGF23 | -0.439 | 0.000505389 |
| Nsp13 | BMP4 | -0.439 | 0.000505331 |
| Nsp14 | TUBA3E | -0.439 | 0.000500477 |
| Nsp14 | PPP1R3F | -0.439 | 0.000510178 |
| Nsp15 | TNFRSF14 | -0.439 | 0.000508633 |
| Nsp15 | PCK1 | -0.439 | 0.000496764 |
| Nsp15 | TAOK2 | -0.439 | 0.000506888 |
| Nsp15 | GDF2 | -0.439 | 0.000512267 |
| Nsp16 | TNFRSF14 | -0.439 | 0.000497853 |
| Nsp16 | TRAF4 | -0.439 | 0.000501333 |
| Nsp16 | VHL | -0.439 | 0.000506344 |
| S | FGF23 | -0.439 | 0.000513043 |
| S | RAPGEF1 | -0.439 | 0.000501931 |
| Orf3a | CAMKK2 | -0.439 | 0.000497132 |
| Orf3a | NTRK1 | -0.439 | 0.000512318 |
| Orf3a | ISG15 | -0.439 | 0.000498042 |
| Orf3a | GREM2 | -0.439 | 0.000497544 |
| Orf6 | MAPK11 | -0.439 | 0.000508918 |
| Orf8 | DUSP2 | -0.439 | 0.000509122 |
| Orf8 | GRK6 | -0.439 | 0.000499659 |
| Orf8 | CX3CL1 | -0.439 | 0.00049774 |
| Orf8 | CPT1C | -0.439 | 0.000500451 |
| Orf10 | ELMO1 | -0.439 | 0.000512216 |
| Nsp2 | IL21R | -0.44 | 0.000488224 |
| Nsp2 | WNT5A | -0.44 | 0.000495675 |
| Nsp3 | AKT1S1 | -0.44 | 0.000490312 |
| Nsp3 | HK3 | -0.44 | 0.000485331 |
| Nsp3 | MAPKAPK2 | -0.44 | 0.00049097 |
| Nsp4 | TUBA8 | -0.44 | 0.000489918 |
| Nsp5 | MAP2K3 | -0.44 | 0.000488424 |
| Nsp5 | EGFR | -0.44 | 0.000488427 |

| | | | |
|---|---|---|---|
| Nsp5 | ERBB2 | -0.44 | 0.00048147 |
| Nsp8 | PLA2G4B | -0.44 | 0.000488745 |
| Nsp10 | PITX2 | -0.44 | 0.00049586 |
| Nsp10 | LAMA5 | -0.44 | 0.000485652 |
| Nsp10 | SOCS3 | -0.44 | 0.000486235 |
| Nsp10 | PTPN7 | -0.44 | 0.000488933 |
| Nsp12 | GADD45G | -0.44 | 0.000490813 |
| Nsp12 | ID3 | -0.44 | 0.000486538 |
| Nsp13 | FGF16 | -0.44 | 0.000490911 |
| Nsp14 | TNFRSF25 | -0.44 | 0.000491862 |
| Nsp14 | FGF2 | -0.44 | 0.000490585 |
| Nsp14 | TUBA3D | -0.44 | 0.000481361 |
| Nsp14 | IGF2 | -0.44 | 0.000493129 |
| Nsp14 | LPAR2 | -0.44 | 0.000493643 |
| Nsp15 | CTSZ | -0.44 | 0.000480329 |
| Nsp15 | GNB3 | -0.44 | 0.000488342 |
| Nsp16 | LAMB3 | -0.44 | 0.000481352 |
| Nsp16 | MAPKAPK2 | -0.44 | 0.000494973 |
| Nsp16 | TNFRSF1A | -0.44 | 0.000480619 |
| S | GRK5 | -0.44 | 0.00048686 |
| S | FGF16 | -0.44 | 0.000490487 |
| E | BAD | -0.44 | 0.000485587 |
| Orf3a | BMP6 | -0.44 | 0.00049099 |
| Orf3a | GATA3 | -0.44 | 0.000480427 |
| Orf3a | LAMA5 | -0.44 | 0.000493653 |
| Orf6 | JUND | -0.44 | 0.000493077 |
| Orf6 | FOSL1 | -0.44 | 0.000481284 |
| Orf6 | MAP2K2 | -0.44 | 0.000482306 |
| Orf8 | CACNB1 | -0.44 | 0.000481924 |
| Orf8 | ITGB3 | -0.44 | 0.000482638 |
| Nsp2 | PRF1 | -0.441 | 0.000473774 |
| Nsp2 | DUSP3 | -0.441 | 0.000479228 |
| Nsp2 | CACNA2D2 | -0.441 | 0.000477777 |
| Nsp3 | TUBA3C | -0.441 | 0.000472642 |
| Nsp3 | EGLN2 | -0.441 | 0.000470709 |
| Nsp4 | ANGPT4 | -0.441 | 0.000472224 |
| Nsp4 | GADD45A | -0.441 | 0.000469028 |
| Nsp4 | ID2 | -0.441 | 0.000467964 |
| Nsp5 | CAPN2 | -0.441 | 0.0004725 |
| Nsp5 | CXCL2 | -0.441 | 0.000479837 |
| Nsp5 | NPRL3 | -0.441 | 0.000477612 |
| Nsp5 | ENO3 | -0.441 | 0.00047653 |
| Nsp5 | ITGB3 | -0.441 | 0.000474904 |
| Nsp7 | HSPA2 | -0.441 | 0.000469423 |
| Nsp7 | CNTFR | -0.441 | 0.000470202 |
| Nsp8 | HRAS | -0.441 | 0.000466161 |
| Nsp8 | VEGFB | -0.441 | 0.000473236 |
| Nsp10 | DAB2IP | -0.441 | 0.000471518 |
| Nsp10 | PREX1 | -0.441 | 0.00046694 |
| Nsp10 | PRKACG | -0.441 | 0.000475293 |

| | | | |
|---|---|---|---|
| Nsp10 | PLCB3 | -0.441 | 0.000479881 |
| Nsp12 | TNFRSF25 | -0.441 | 0.000467865 |
| Nsp14 | RELT | -0.441 | 0.000472561 |
| Nsp14 | PLCG2 | -0.441 | 0.000468707 |
| Nsp14 | PRKAB1 | -0.441 | 0.000470921 |
| Nsp14 | GADD45A | -0.441 | 0.000468588 |
| Nsp15 | IFNL2 | -0.441 | 0.000474015 |
| Nsp16 | IFNL3 | -0.441 | 0.00046642 |
| Nsp16 | IRF4 | -0.441 | 0.000467188 |
| Nsp16 | ELAVL1 | -0.441 | 0.000465509 |
| Nsp16 | SLC3A2 | -0.441 | 0.000473375 |
| Nsp16 | PIK3R6 | -0.441 | 0.000477857 |
| S | CRK | -0.441 | 0.000476764 |
| Orf3a | FZD7 | -0.441 | 0.000473559 |
| Orf3a | MAP2K7 | -0.441 | 0.000470067 |
| Orf6 | HSPA2 | -0.441 | 0.00047372 |
| Orf6 | DUSP4 | -0.441 | 0.000473948 |
| Orf6 | ECSIT | -0.441 | 0.00048016 |
| Orf6 | NCF1 | -0.441 | 0.000475703 |
| Orf8 | PRKCD | -0.441 | 0.000475919 |
| Orf10 | RXRB | -0.441 | 0.00047124 |
| Orf10 | CDKN1A | -0.441 | 0.000478252 |
| Orf10 | PLAU | -0.441 | 0.000469357 |
| Nsp2 | TNFSF14 | -0.442 | 0.000460676 |
| Nsp2 | GNAI2 | -0.442 | 0.000455797 |
| Nsp3 | WNT8B | -0.442 | 0.000452081 |
| Nsp3 | CSF3R | -0.442 | 0.000460586 |
| Nsp5 | ELK1 | -0.442 | 0.00046451 |
| Nsp5 | VEGFB | -0.442 | 0.000454173 |
| Nsp6 | LMNB2 | -0.442 | 0.000457913 |
| Nsp6 | MAPK8IP3 | -0.442 | 0.000463261 |
| Nsp7 | PKN1 | -0.442 | 0.000456845 |
| Nsp7 | VEGFA | -0.442 | 0.00045189 |
| Nsp8 | ELK1 | -0.442 | 0.000456738 |
| Nsp9 | SOCS1 | -0.442 | 0.000454388 |
| Nsp9 | ELAVL1 | -0.442 | 0.000461048 |
| Nsp9 | BMP8A | -0.442 | 0.000463555 |
| Nsp10 | DVL1 | -0.442 | 0.000458509 |
| Nsp10 | HSPA1B | -0.442 | 0.000450192 |
| Nsp12 | RASGRP4 | -0.442 | 0.00046355 |
| Nsp12 | CX3CL1 | -0.442 | 0.000453084 |
| Nsp13 | FGF23 | -0.442 | 0.000459324 |
| Nsp14 | CACNA1S | -0.442 | 0.000463064 |
| Nsp14 | ACTG1 | -0.442 | 0.000453514 |
| Nsp14 | ANGPT4 | -0.442 | 0.000462752 |
| Nsp14 | CHRM1 | -0.442 | 0.00044949 |
| Nsp14 | ITGA11 | -0.442 | 0.000462036 |
| Nsp14 | OSM | -0.442 | 0.000459433 |
| Nsp14 | CXCL2 | -0.442 | 0.000459816 |
| Nsp14 | TNFRSF1A | -0.442 | 0.000462274 |

| | | | |
|---|---|---|---|
| Nsp14 | CACNG2 | -0.442 | 0.000461788 |
| Nsp15 | MAP2K3 | -0.442 | 0.000453998 |
| Nsp15 | PLA2G4F | -0.442 | 0.000459443 |
| Nsp16 | GH1 | -0.442 | 0.000463167 |
| S | RELT | -0.442 | 0.000459241 |
| S | PRKCZ | -0.442 | 0.000463962 |
| S | ANGPT4 | -0.442 | 0.000457256 |
| S | PRKCD | -0.442 | 0.00046277 |
| S | IL21R | -0.442 | 0.000451343 |
| S | MRAS | -0.442 | 0.000458147 |
| S | CPT1B | -0.442 | 0.000453991 |
| Orf3a | TNFRSF8 | -0.442 | 0.000463478 |
| Orf3a | MAP2K2 | -0.442 | 0.000464344 |
| Orf6 | CHAD | -0.442 | 0.000457403 |
| Orf6 | SMAD6 | -0.442 | 0.000463397 |
| Orf8 | NFKB2 | -0.442 | 0.000457735 |
| Orf8 | EPOR | -0.442 | 0.000459994 |
| Orf8 | CACNA1E | -0.442 | 0.000458903 |
| Nsp2 | AKT2 | -0.443 | 0.000441355 |
| Nsp2 | SHC3 | -0.443 | 0.00044477 |
| Nsp2 | ECSIT | -0.443 | 0.000442881 |
| Nsp2 | BCL3 | -0.443 | 0.000436056 |
| Nsp2 | WNT4 | -0.443 | 0.000441592 |
| Nsp3 | ITGB3 | -0.443 | 0.000441206 |
| Nsp5 | TRAF1 | -0.443 | 0.000438991 |
| Nsp5 | PRKCD | -0.443 | 0.0004423 |
| Nsp5 | MAP4K2 | -0.443 | 0.00043594 |
| Nsp5 | TNFRSF12A | -0.443 | 0.000436073 |
| Nsp6 | DUSP7 | -0.443 | 0.00043535 |
| Nsp6 | ZAP70 | -0.443 | 0.000436655 |
| Nsp8 | MAP2K3 | -0.443 | 0.000439138 |
| Nsp8 | ARRB2 | -0.443 | 0.000435601 |
| Nsp8 | ID2 | -0.443 | 0.000449064 |
| Nsp8 | CSF3 | -0.443 | 0.000446302 |
| Nsp8 | G6PC3 | -0.443 | 0.00044325 |
| Nsp10 | SRC | -0.443 | 0.000436242 |
| Nsp10 | CARD10 | -0.443 | 0.000438971 |
| Nsp10 | DUSP6 | -0.443 | 0.000436406 |
| Nsp10 | LTB | -0.443 | 0.000448079 |
| Nsp10 | ADCY5 | -0.443 | 0.000448534 |
| Nsp12 | TUBA3C | -0.443 | 0.000435246 |
| Nsp12 | GH1 | -0.443 | 0.000436569 |
| Nsp12 | RPS6KA1 | -0.443 | 0.000439502 |
| Nsp12 | RELA | -0.443 | 0.000445815 |
| Nsp12 | CACNB3 | -0.443 | 0.000436361 |
| Nsp12 | EGLN2 | -0.443 | 0.000444256 |
| Nsp12 | CACNA1F | -0.443 | 0.000435696 |
| Nsp13 | CDC37 | -0.443 | 0.000442659 |
| Nsp13 | COL9A2 | -0.443 | 0.000436059 |
| Nsp14 | GADD45G | -0.443 | 0.00044478 |

| | | | |
|---|---|---|---|
| Nsp14 | CACNA2D2 | -0.443 | 0.00043874 |
| Nsp14 | MAPKAPK2 | -0.443 | 0.000435573 |
| Nsp15 | CREB3L3 | -0.443 | 0.000436059 |
| Nsp15 | ACSBG1 | -0.443 | 0.000437863 |
| Nsp16 | CDKN1A | -0.443 | 0.0004404 |
| S | NOS2 | -0.443 | 0.000436824 |
| S | TUBA3C | -0.443 | 0.000444469 |
| Orf3a | CACNA1G | -0.443 | 0.00044555 |
| Orf3a | PRR5 | -0.443 | 0.000441467 |
| Orf3a | CARD11 | -0.443 | 0.000448838 |
| Orf6 | FZD8 | -0.443 | 0.000448372 |
| Orf6 | WDR24 | -0.443 | 0.000436697 |
| Orf6 | JUN | -0.443 | 0.000445731 |
| Orf6 | FZD9 | -0.443 | 0.00043913 |
| Orf6 | AMH | -0.443 | 0.000438344 |
| Orf6 | PXN | -0.443 | 0.000439483 |
| Orf8 | CTSW | -0.443 | 0.000446646 |
| Orf8 | G6PC3 | -0.443 | 0.000442756 |
| Nsp2 | MAPK3 | -0.444 | 0.000428259 |
| Nsp2 | CD14 | -0.444 | 0.000428937 |
| Nsp2 | INSR | -0.444 | 0.000425502 |
| Nsp2 | FOSL1 | -0.444 | 0.000432008 |
| Nsp2 | SOCS3 | -0.444 | 0.000422352 |
| Nsp3 | IRF5 | -0.444 | 0.000425869 |
| Nsp3 | IL17RE | -0.444 | 0.000427211 |
| Nsp3 | MAX | -0.444 | 0.000434161 |
| Nsp5 | G6PC3 | -0.444 | 0.000429824 |
| Nsp7 | COMP | -0.444 | 0.000427477 |
| Nsp7 | TNFRSF13C | -0.444 | 0.000422776 |
| Nsp8 | PPP2R5D | -0.444 | 0.000427324 |
| Nsp8 | HMOX1 | -0.444 | 0.000432096 |
| Nsp8 | PRL | -0.444 | 0.000430738 |
| Nsp8 | ENO3 | -0.444 | 0.00042814 |
| Nsp9 | IGF2 | -0.444 | 0.00042852 |
| Nsp9 | RARA | -0.444 | 0.000427562 |
| Nsp10 | RPTOR | -0.444 | 0.000421152 |
| Nsp10 | LAMC3 | -0.444 | 0.000433137 |
| Nsp10 | CACNG5 | -0.444 | 0.000421975 |
| Nsp13 | GH2 | -0.444 | 0.000433051 |
| Nsp13 | RRAGA | -0.444 | 0.000427964 |
| Nsp14 | ECSIT | -0.444 | 0.000429507 |
| Nsp14 | FOSL1 | -0.444 | 0.000423481 |
| Nsp14 | PDGFRB | -0.444 | 0.000422318 |
| Nsp15 | IL17A | -0.444 | 0.000432266 |
| Nsp15 | PRKACA | -0.444 | 0.000434606 |
| Nsp15 | CDKN1A | -0.444 | 0.000426584 |
| Nsp16 | DVL2 | -0.444 | 0.000430341 |
| Nsp16 | TNFRSF13B | -0.444 | 0.000429291 |
| S | NPRL2 | -0.444 | 0.000424706 |
| Orf3a | PRKCG | -0.444 | 0.000426597 |

| | | | |
|---|---|---|---|
| Orf6 | ISG15 | -0.444 | 0.000425638 |
| Orf6 | FLT4 | -0.444 | 0.000420578 |
| Orf6 | WNT5B | -0.444 | 0.000426469 |
| Orf8 | ISG15 | -0.444 | 0.000434411 |
| Orf8 | TAOK2 | -0.444 | 0.000422696 |
| Orf8 | NOD2 | -0.444 | 0.000429801 |
| Orf10 | DDIT3 | -0.444 | 0.000425943 |
| Nsp2 | DUSP1 | -0.445 | 0.000413233 |
| Nsp2 | PLCB2 | -0.445 | 0.000416923 |
| Nsp2 | PTPN6 | -0.445 | 0.000413253 |
| Nsp2 | EGLN2 | -0.445 | 0.000410421 |
| Nsp2 | PRKACG | -0.445 | 0.000411969 |
| Nsp3 | NBL1 | -0.445 | 0.000417212 |
| Nsp3 | IFNL2 | -0.445 | 0.000408175 |
| Nsp4 | CX3CL1 | -0.445 | 0.00041499 |
| Nsp5 | JMJD7-PLA2G | -0.445 | 0.000418675 |
| Nsp5 | TNFSF9 | -0.445 | 0.000413568 |
| Nsp5 | CSF2 | -0.445 | 0.000407281 |
| Nsp5 | PKN3 | -0.445 | 0.000416079 |
| Nsp5 | HRAS | -0.445 | 0.000415353 |
| Nsp6 | DDIT4 | -0.445 | 0.000414328 |
| Nsp6 | INHBB | -0.445 | 0.00041516 |
| Nsp7 | CARD14 | -0.445 | 0.000419994 |
| Nsp7 | TRAF3 | -0.445 | 0.000418446 |
| Nsp8 | GNG8 | -0.445 | 0.000407613 |
| Nsp8 | SHC3 | -0.445 | 0.000418813 |
| Nsp8 | ELOB | -0.445 | 0.000409449 |
| Nsp9 | PIK3R2 | -0.445 | 0.000407935 |
| Nsp12 | PF4V1 | -0.445 | 0.000417313 |
| Nsp13 | PPP2R5D | -0.445 | 0.00041452 |
| Nsp13 | CPT1A | -0.445 | 0.000416864 |
| Nsp13 | NPRL2 | -0.445 | 0.000410619 |
| Nsp14 | ID2 | -0.445 | 0.000407692 |
| Nsp14 | PTPN5 | -0.445 | 0.000414896 |
| Nsp15 | CCND3 | -0.445 | 0.000408972 |
| Nsp15 | CCND2 | -0.445 | 0.000413331 |
| Nsp15 | RAC3 | -0.445 | 0.000409792 |
| Nsp15 | TNFSF12 | -0.445 | 0.00041074 |
| Nsp15 | CAMK2A | -0.445 | 0.000408042 |
| Nsp15 | VHL | -0.445 | 0.000409664 |
| Orf3a | PIAS4 | -0.445 | 0.000415673 |
| Orf3a | RASGRP2 | -0.445 | 0.000407456 |
| Orf6 | RPS6KA2 | -0.445 | 0.000414357 |
| Orf6 | COL6A1 | -0.445 | 0.000418226 |
| Orf8 | FOS | -0.445 | 0.000416896 |
| Orf8 | CAMK2B | -0.445 | 0.000420185 |
| Orf8 | BCAR1 | -0.445 | 0.000408518 |
| Orf8 | FOXO3 | -0.445 | 0.000410185 |
| Nsp2 | CREB3L1 | -0.446 | 0.00039588 |
| Nsp3 | BMP4 | -0.446 | 0.000403406 |

| | | | |
|---|---|---|---|
| Nsp3 | MRAS | -0.446 | 0.00040582 |
| Nsp3 | CACNA1F | -0.446 | 0.000404835 |
| Nsp3 | G6PC3 | -0.446 | 0.00040031 |
| Nsp4 | MMP14 | -0.446 | 0.000394186 |
| Nsp4 | PF4 | -0.446 | 0.000402318 |
| Nsp4 | PRL | -0.446 | 0.000395346 |
| Nsp4 | INS | -0.446 | 0.000395198 |
| Nsp5 | TIMP1 | -0.446 | 0.000405389 |
| Nsp7 | TOLLIP | -0.446 | 0.000400762 |
| Nsp7 | PTPN6 | -0.446 | 0.000393714 |
| Nsp7 | COL9A2 | -0.446 | 0.000400906 |
| Nsp7 | ADCY5 | -0.446 | 0.000404385 |
| Nsp10 | IL25 | -0.446 | 0.000399614 |
| Nsp10 | NOS3 | -0.446 | 0.000404351 |
| Nsp12 | IGF2 | -0.446 | 0.00040345 |
| Nsp13 | RASGRP4 | -0.446 | 0.000401343 |
| Nsp15 | CSF3R | -0.446 | 0.000398189 |
| Nsp16 | NBL1 | -0.446 | 0.000393379 |
| Nsp16 | PLA2G4F | -0.446 | 0.000397235 |
| Nsp16 | MRAS | -0.446 | 0.000393924 |
| S | PDGFA | -0.446 | 0.000396168 |
| S | CSF2RB | -0.446 | 0.000399727 |
| Orf3a | RXRA | -0.446 | 0.000406453 |
| Orf3a | PPP5C | -0.446 | 0.000394577 |
| Orf6 | GRK6 | -0.446 | 0.000402388 |
| Orf6 | MAPK4 | -0.446 | 0.000393975 |
| Orf6 | CDKN1A | -0.446 | 0.00039757 |
| Orf8 | CHRM1 | -0.446 | 0.000401284 |
| Orf8 | CARD10 | -0.446 | 0.000396905 |
| Orf8 | HSPA6 | -0.446 | 0.000394568 |
| Orf8 | TNFRSF1A | -0.446 | 0.000394384 |
| Nsp2 | RRAS | -0.447 | 0.000381351 |
| Nsp2 | CACNG1 | -0.447 | 0.000380364 |
| Nsp2 | MAPK15 | -0.447 | 0.00038226 |
| Nsp2 | LAMC3 | -0.447 | 0.000384783 |
| Nsp2 | PHLPP1 | -0.447 | 0.00039166 |
| Nsp2 | ACVRL1 | -0.447 | 0.000385502 |
| Nsp3 | CREBBP | -0.447 | 0.000392456 |
| Nsp3 | NLRX1 | -0.447 | 0.000380648 |
| Nsp3 | VAV1 | -0.447 | 0.000390234 |
| Nsp4 | FOS | -0.447 | 0.000390576 |
| Nsp4 | CACNA1E | -0.447 | 0.000392917 |
| Nsp5 | NTF3 | -0.447 | 0.00038892 |
| Nsp7 | MAPK3 | -0.447 | 0.000392882 |
| Nsp7 | SMAD6 | -0.447 | 0.000381991 |
| Nsp7 | MMP9 | -0.447 | 0.000385946 |
| Nsp7 | INHBB | -0.447 | 0.000389464 |
| Nsp7 | JUND | -0.447 | 0.000381782 |
| Nsp7 | BCL2 | -0.447 | 0.000385179 |
| Nsp8 | HK3 | -0.447 | 0.000391876 |

| | | | |
|---|---|---|---|
| Nsp8 | FGF23 | -0.447 | 0.000381256 |
| Nsp8 | VWF | -0.447 | 0.000387208 |
| Nsp9 | MAPK8IP2 | -0.447 | 0.000382939 |
| Nsp10 | FOSB | -0.447 | 0.000384743 |
| Nsp10 | YWHAG | -0.447 | 0.000389046 |
| Nsp10 | WNT9B | -0.447 | 0.000391572 |
| Nsp10 | PPP1R3D | -0.447 | 0.00038711 |
| Nsp10 | HSPA1A | -0.447 | 0.000392516 |
| Nsp10 | LCK | -0.447 | 0.000380572 |
| Nsp12 | CPT1A | -0.447 | 0.000381858 |
| Nsp14 | FGF6 | -0.447 | 0.000391185 |
| Nsp15 | ADCY8 | -0.447 | 0.000391608 |
| Nsp16 | CCND3 | -0.447 | 0.000386772 |
| Nsp16 | CRK | -0.447 | 0.000385337 |
| Nsp16 | FGFR1 | -0.447 | 0.000384781 |
| S | MAPK15 | -0.447 | 0.000384204 |
| S | GFAP | -0.447 | 0.000391741 |
| S | ENO3 | -0.447 | 0.000392033 |
| Orf3a | CACNA1H | -0.447 | 0.000388591 |
| Orf3a | RAC3 | -0.447 | 0.000387849 |
| Orf3a | ZAP70 | -0.447 | 0.000383039 |
| Orf8 | TNFRSF14 | -0.447 | 0.000389511 |
| Orf8 | BBC3 | -0.447 | 0.00038681 |
| Nsp2 | BAD | -0.448 | 0.000369362 |
| Nsp2 | RASGRF1 | -0.448 | 0.000379006 |
| Nsp3 | LBP | -0.448 | 0.000371618 |
| Nsp3 | FGF21 | -0.448 | 0.000378914 |
| Nsp4 | PDGFA | -0.448 | 0.000377567 |
| Nsp4 | AKT1S1 | -0.448 | 0.000374256 |
| Nsp4 | FOSL1 | -0.448 | 0.000371143 |
| Nsp5 | AKT1 | -0.448 | 0.000377522 |
| Nsp5 | ELOB | -0.448 | 0.000372773 |
| Nsp6 | HSPA2 | -0.448 | 0.000370918 |
| Nsp7 | CARD10 | -0.448 | 0.000379593 |
| Nsp7 | MAPK4 | -0.448 | 0.000377022 |
| Nsp7 | BMP8B | -0.448 | 0.000377382 |
| Nsp7 | BAD | -0.448 | 0.000373942 |
| Nsp8 | RNF152 | -0.448 | 0.000379403 |
| Nsp8 | HK2 | -0.448 | 0.000376147 |
| Nsp9 | ENDOG | -0.448 | 0.000370037 |
| Nsp9 | MAPK3 | -0.448 | 0.000369352 |
| Nsp10 | HSPA2 | -0.448 | 0.00037446 |
| Nsp10 | ITGA11 | -0.448 | 0.000373565 |
| Nsp10 | PPP2R2C | -0.448 | 0.0003685 |
| Nsp10 | PIK3CD | -0.448 | 0.000377342 |
| Nsp10 | CSF3R | -0.448 | 0.000372253 |
| Nsp12 | IL2RB | -0.448 | 0.00037296 |
| Nsp12 | MAP4K1 | -0.448 | 0.000377591 |
| Nsp12 | TAOK2 | -0.448 | 0.000379772 |
| Nsp12 | S100A9 | -0.448 | 0.000370036 |

| | | | |
|---|---|---|---|
| Nsp12 | TIMP1 | -0.448 | 0.000371461 |
| Nsp12 | EIF4EBP1 | -0.448 | 0.000375785 |
| Nsp12 | MRAS | -0.448 | 0.000379656 |
| Nsp13 | ELK1 | -0.448 | 0.000372523 |
| Nsp13 | DFFB | -0.448 | 0.000374775 |
| Nsp14 | MMP14 | -0.448 | 0.000378812 |
| Nsp15 | ERN1 | -0.448 | 0.000367704 |
| Nsp16 | CREBBP | -0.448 | 0.000374808 |
| S | FOXO1 | -0.448 | 0.00037302 |
| S | PPP2R5D | -0.448 | 0.000373309 |
| S | RASGRP4 | -0.448 | 0.00037104 |
| S | IGF1R | -0.448 | 0.000370918 |
| S | RRAGA | -0.448 | 0.00037099 |
| Orf3a | CAPN1 | -0.448 | 0.000376833 |
| Orf8 | IL2RB | -0.448 | 0.000376456 |
| Orf8 | LMNA | -0.448 | 0.000373899 |
| Nsp2 | SOCS7 | -0.449 | 0.000357076 |
| Nsp2 | PPP1R3E | -0.449 | 0.000360196 |
| Nsp3 | IRF4 | -0.449 | 0.000357423 |
| Nsp3 | FGR | -0.449 | 0.000365947 |
| Nsp3 | GDF2 | -0.449 | 0.0003667 |
| Nsp3 | FGF16 | -0.449 | 0.000359045 |
| Nsp4 | CSF1R | -0.449 | 0.000361184 |
| Nsp4 | GNG13 | -0.449 | 0.000362651 |
| Nsp4 | IL17RE | -0.449 | 0.000357659 |
| Nsp4 | FGF21 | -0.449 | 0.000360037 |
| Nsp4 | FGF23 | -0.449 | 0.000365668 |
| Nsp5 | PIDD1 | -0.449 | 0.000362525 |
| Nsp5 | HSPA6 | -0.449 | 0.000361965 |
| Nsp5 | NPRL2 | -0.449 | 0.000362031 |
| Nsp6 | SOCS1 | -0.449 | 0.0003574 |
| Nsp6 | CARD14 | -0.449 | 0.000361945 |
| Nsp6 | WNT7B | -0.449 | 0.000360017 |
| Nsp6 | LEFTY1 | -0.449 | 0.000363313 |
| Nsp7 | RELT | -0.449 | 0.000361792 |
| Nsp8 | IL17RE | -0.449 | 0.000355953 |
| Nsp9 | GDF11 | -0.449 | 0.000361795 |
| Nsp9 | PPP2R3B | -0.449 | 0.000366746 |
| Nsp9 | TNFRSF12A | -0.449 | 0.000356357 |
| Nsp9 | ADCY5 | -0.449 | 0.000359253 |
| Nsp10 | CSF1 | -0.449 | 0.000359523 |
| Nsp10 | ULK1 | -0.449 | 0.000359359 |
| Nsp12 | NPRL2 | -0.449 | 0.000359164 |
| Nsp12 | CACNG2 | -0.449 | 0.000357007 |
| Nsp13 | FGR | -0.449 | 0.000363625 |
| Nsp13 | MAP4K1 | -0.449 | 0.000363623 |
| Nsp13 | IL17B | -0.449 | 0.000356149 |
| Nsp13 | CACNA1F | -0.449 | 0.000363499 |
| Nsp14 | MAPK15 | -0.449 | 0.000355961 |
| Nsp14 | WNT10B | -0.449 | 0.00036213 |

| | | | |
|---|---|---|---|
| Nsp14 | GNAI2 | -0.449 | 0.000366145 |
| Nsp14 | CX3CL1 | -0.449 | 0.000355595 |
| Nsp14 | IL12RB1 | -0.449 | 0.000359077 |
| Nsp14 | ADCY6 | -0.449 | 0.000363885 |
| Nsp15 | IL17RE | -0.449 | 0.000357719 |
| Nsp15 | IL13 | -0.449 | 0.000357036 |
| Nsp15 | FGFR1 | -0.449 | 0.000362839 |
| Nsp15 | TNFRSF13B | -0.449 | 0.000356558 |
| Nsp15 | NFKBIA | -0.449 | 0.00035737 |
| Orf3a | PLCB3 | -0.449 | 0.000361305 |
| Orf6 | CACNA1H | -0.449 | 0.000361711 |
| Orf6 | FZD5 | -0.449 | 0.000365308 |
| Orf6 | FZD1 | -0.449 | 0.00036339 |
| Orf8 | PLA2G4B | -0.449 | 0.00035526 |
| Orf8 | CDC37 | -0.449 | 0.000360475 |
| Orf8 | PDGFB | -0.449 | 0.000366044 |
| Nsp3 | PF4V1 | -0.45 | 0.000349789 |
| Nsp3 | CACNB3 | -0.45 | 0.000349117 |
| Nsp4 | ACSBG1 | -0.45 | 0.000349732 |
| Nsp4 | RORC | -0.45 | 0.000343615 |
| Nsp4 | GFAP | -0.45 | 0.00035188 |
| Nsp6 | BMP8A | -0.45 | 0.000352817 |
| Nsp7 | EXOC7 | -0.45 | 0.000346853 |
| Nsp7 | COL6A2 | -0.45 | 0.0003458 |
| Nsp8 | STAT5A | -0.45 | 0.000351916 |
| Nsp8 | ITGA11 | -0.45 | 0.000352849 |
| Nsp9 | PGAM5 | -0.45 | 0.000347642 |
| Nsp9 | WNT3 | -0.45 | 0.000352396 |
| Nsp10 | DUSP3 | -0.45 | 0.000353921 |
| Nsp10 | PFKP | -0.45 | 0.000354094 |
| Nsp10 | TBX21 | -0.45 | 0.000347659 |
| Nsp10 | SMAD3 | -0.45 | 0.000353151 |
| Nsp12 | PRKCD | -0.45 | 0.000349396 |
| Nsp13 | HK2 | -0.45 | 0.000353765 |
| Nsp14 | RORC | -0.45 | 0.000343721 |
| Nsp14 | RUNX1 | -0.45 | 0.00035241 |
| Nsp15 | IRF4 | -0.45 | 0.000349621 |
| Nsp15 | FOXO1 | -0.45 | 0.000345757 |
| Nsp15 | LAMB3 | -0.45 | 0.000352493 |
| Nsp16 | AKT1S1 | -0.45 | 0.000347356 |
| Nsp16 | RNF152 | -0.45 | 0.000354079 |
| Nsp16 | RRAGA | -0.45 | 0.000352815 |
| Nsp16 | CSF3 | -0.45 | 0.00034858 |
| S | CCND3 | -0.45 | 0.000350882 |
| S | CLCF1 | -0.45 | 0.000344144 |
| Orf3a | HSPA6 | -0.45 | 0.000347366 |
| Orf3a | PPP1R3D | -0.45 | 0.000343223 |
| Orf3a | PKN1 | -0.45 | 0.000351733 |
| Orf6 | RPS6KA4 | -0.45 | 0.000354031 |
| Orf8 | ARRB2 | -0.45 | 0.000351528 |

| | | | |
|---|---|---|---|
| Orf8 | SESN2 | -0.45 | 0.000347107 |
| Orf8 | RAC2 | -0.45 | 0.000346806 |
| Orf8 | TNN | -0.45 | 0.000350995 |
| Orf8 | FLNC | -0.45 | 0.000346066 |
| Nsp2 | EPOR | -0.451 | 0.000334744 |
| Nsp2 | TRAF2 | -0.451 | 0.000339307 |
| Nsp2 | WNT3 | -0.451 | 0.000335462 |
| Nsp3 | TGFB3 | -0.451 | 0.000332142 |
| Nsp3 | CREB3L3 | -0.451 | 0.000338549 |
| Nsp3 | EGFR | -0.451 | 0.000336341 |
| Nsp3 | FGFR1 | -0.451 | 0.000336733 |
| Nsp5 | CTSW | -0.451 | 0.000336917 |
| Nsp5 | MAPK15 | -0.451 | 0.000336462 |
| Nsp5 | CDC37 | -0.451 | 0.000337898 |
| Nsp6 | HSPB1 | -0.451 | 0.000342095 |
| Nsp6 | GREM2 | -0.451 | 0.000341688 |
| Nsp7 | ELK1 | -0.451 | 0.000338617 |
| Nsp7 | TRAF4 | -0.451 | 0.000341949 |
| Nsp7 | LPAR5 | -0.451 | 0.000338707 |
| Nsp8 | CAMK2A | -0.451 | 0.000333495 |
| Nsp8 | CACNG2 | -0.451 | 0.00033683 |
| Nsp9 | RPS6KA4 | -0.451 | 0.000335815 |
| Nsp9 | YWHAG | -0.451 | 0.000336565 |
| Nsp9 | RAC2 | -0.451 | 0.0003366 |
| Nsp9 | PPP1R3E | -0.451 | 0.000341666 |
| Nsp9 | NOG | -0.451 | 0.000341925 |
| Nsp10 | TRIM25 | -0.451 | 0.000343033 |
| Nsp10 | ID2 | -0.451 | 0.000340799 |
| Nsp12 | SHC3 | -0.451 | 0.000336413 |
| Nsp12 | PLA2G4F | -0.451 | 0.000333333 |
| Nsp14 | TRAF1 | -0.451 | 0.000341698 |
| Nsp14 | EXOC7 | -0.451 | 0.00033351 |
| Nsp15 | NOS2 | -0.451 | 0.000334623 |
| Nsp15 | CTSW | -0.451 | 0.000339001 |
| Nsp15 | ELAVL1 | -0.451 | 0.000341387 |
| Nsp16 | MAP4K1 | -0.451 | 0.000337389 |
| Nsp16 | S100A9 | -0.451 | 0.000334931 |
| S | IRAK1 | -0.451 | 0.000336067 |
| S | RELA | -0.451 | 0.000335784 |
| S | PPP2R5B | -0.451 | 0.000342403 |
| S | IL17B | -0.451 | 0.000338615 |
| Orf3a | CDC25B | -0.451 | 0.000332994 |
| Orf3a | MAPK3 | -0.451 | 0.000337496 |
| Orf3a | WNT6 | -0.451 | 0.000337109 |
| Orf3a | CCND1 | -0.451 | 0.000339703 |
| Orf6 | DDIT4 | -0.451 | 0.000337328 |
| Orf6 | GRK2 | -0.451 | 0.000340038 |
| Orf6 | MAPK8IP3 | -0.451 | 0.00033808 |
| Orf6 | IL36RN | -0.451 | 0.00033284 |
| Orf8 | CSF1R | -0.451 | 0.000335945 |

| | | | |
|---|---|---|---|
| Orf8 | IL17RC | -0.451 | 0.000340991 |
| Orf8 | FGF18 | -0.451 | 0.000340586 |
| Orf8 | IFNLR1 | -0.451 | 0.000335096 |
| Orf8 | S100A9 | -0.451 | 0.000335683 |
| Nsp2 | CCR10 | -0.452 | 0.000320574 |
| Nsp4 | FGR | -0.452 | 0.000327275 |
| Nsp4 | CAMK2A | -0.452 | 0.000322005 |
| Nsp4 | PPP1R3F | -0.452 | 0.000327346 |
| Nsp4 | FGF16 | -0.452 | 0.000321687 |
| Nsp5 | GSK3A | -0.452 | 0.000329563 |
| Nsp5 | GNG13 | -0.452 | 0.000329958 |
| Nsp5 | DUSP3 | -0.452 | 0.000330965 |
| Nsp5 | ITGB5 | -0.452 | 0.000325059 |
| Nsp5 | WNT8B | -0.452 | 0.000329737 |
| Nsp6 | PPP1R3D | -0.452 | 0.00032377 |
| Nsp7 | FLT4 | -0.452 | 0.000329962 |
| Nsp7 | CACNA1I | -0.452 | 0.000330539 |
| Nsp8 | PDGFA | -0.452 | 0.000327327 |
| Nsp8 | GADD45G | -0.452 | 0.000320774 |
| Nsp9 | LMNB2 | -0.452 | 0.000328528 |
| Nsp9 | TNFRSF6B | -0.452 | 0.000322517 |
| Nsp9 | CACNA1A | -0.452 | 0.000321307 |
| Nsp9 | BAD | -0.452 | 0.000330664 |
| Nsp10 | FZD8 | -0.452 | 0.000327894 |
| Nsp10 | CACNA2D4 | -0.452 | 0.000322492 |
| Nsp10 | PIAS4 | -0.452 | 0.000328935 |
| Nsp12 | CCL14 | -0.452 | 0.000320621 |
| Nsp12 | ARRB2 | -0.452 | 0.00032954 |
| Nsp12 | ACACB | -0.452 | 0.000331325 |
| Nsp12 | MAPKAPK2 | -0.452 | 0.000325725 |
| Nsp13 | ARRB1 | -0.452 | 0.000320604 |
| Nsp13 | IGF2 | -0.452 | 0.000325654 |
| Nsp14 | CDC37 | -0.452 | 0.000326914 |
| Nsp15 | PGF | -0.452 | 0.000323284 |
| Nsp15 | GADD45G | -0.452 | 0.000321849 |
| Nsp16 | PDGFA | -0.452 | 0.000329489 |
| Nsp16 | TAOK2 | -0.452 | 0.000328976 |
| Nsp16 | CSF3R | -0.452 | 0.000329101 |
| S | IL11 | -0.452 | 0.000324552 |
| S | CSH1 | -0.452 | 0.000329582 |
| S | TUBA3E | -0.452 | 0.000322103 |
| Orf3a | HRK | -0.452 | 0.000331102 |
| Orf3a | CACNA1A | -0.452 | 0.000327384 |
| Orf3a | BAD | -0.452 | 0.000324981 |
| Orf6 | PKN1 | -0.452 | 0.000321094 |
| Orf8 | JMJD7-PLA2G | -0.452 | 0.000321181 |
| Orf8 | ECSIT | -0.452 | 0.000320412 |
| Orf8 | GFAP | -0.452 | 0.000330584 |
| Orf8 | CREB3L1 | -0.452 | 0.000330318 |
| Nsp2 | MAPK12 | -0.453 | 0.000309592 |

| | | | |
|---|---|---|---|
| Nsp2 | PIN1 | -0.453 | 0.000309815 |
| Nsp3 | IL11 | -0.453 | 0.000311654 |
| Nsp3 | ANGPT4 | -0.453 | 0.000309825 |
| Nsp3 | GADD45A | -0.453 | 0.000310302 |
| Nsp3 | PF4 | -0.453 | 0.000312164 |
| Nsp4 | FGF2 | -0.453 | 0.000309812 |
| Nsp4 | EGLN2 | -0.453 | 0.000315451 |
| Nsp4 | SLC2A1 | -0.453 | 0.000320229 |
| Nsp4 | CACNG2 | -0.453 | 0.000313194 |
| Nsp5 | MCL1 | -0.453 | 0.000318125 |
| Nsp5 | PPP1R3F | -0.453 | 0.000314812 |
| Nsp5 | LCK | -0.453 | 0.000317839 |
| Nsp5 | CACNA1F | -0.453 | 0.000311118 |
| Nsp6 | FGF22 | -0.453 | 0.000314668 |
| Nsp7 | NTRK1 | -0.453 | 0.000318022 |
| Nsp8 | GDF5 | -0.453 | 0.00031356 |
| Nsp10 | FZD7 | -0.453 | 0.000314465 |
| Nsp10 | RRAS | -0.453 | 0.000312253 |
| Nsp10 | CACNA1G | -0.453 | 0.000319455 |
| Nsp10 | TSC2 | -0.453 | 0.000318892 |
| Nsp10 | ADCY4 | -0.453 | 0.000313897 |
| Nsp10 | MKNK2 | -0.453 | 0.000316659 |
| Nsp13 | TAOK2 | -0.453 | 0.000319506 |
| Nsp13 | ACACB | -0.453 | 0.000315303 |
| Nsp14 | CCND3 | -0.453 | 0.000313642 |
| Nsp14 | CACNA2D4 | -0.453 | 0.000314258 |
| Nsp15 | GNG13 | -0.453 | 0.000317122 |
| Nsp15 | LPAR2 | -0.453 | 0.000311051 |
| Nsp16 | DFFB | -0.453 | 0.00031093 |
| Nsp16 | PPP2R5B | -0.453 | 0.000320167 |
| S | FGF6 | -0.453 | 0.000318299 |
| S | TUBA3D | -0.453 | 0.000313395 |
| Orf3a | IL17D | -0.453 | 0.00031391 |
| Orf3a | SH2B2 | -0.453 | 0.000319764 |
| Orf6 | NFATC1 | -0.453 | 0.000312264 |
| Orf8 | IRF3 | -0.453 | 0.000312703 |
| Nsp2 | FLCN | -0.454 | 0.000303592 |
| Nsp3 | ELAVL1 | -0.454 | 0.000307563 |
| Nsp4 | CACNB3 | -0.454 | 0.000308641 |
| Nsp4 | LPAR2 | -0.454 | 0.000301402 |
| Nsp5 | TNFRSF8 | -0.454 | 0.000308847 |
| Nsp5 | GRK5 | -0.454 | 0.000300625 |
| Nsp5 | CSF3R | -0.454 | 0.000301805 |
| Nsp6 | CACNA1I | -0.454 | 0.000300888 |
| Nsp7 | FZD8 | -0.454 | 0.000305234 |
| Nsp7 | FZD5 | -0.454 | 0.000304775 |
| Nsp7 | CACNG8 | -0.454 | 0.0003028 |
| Nsp8 | PTK2B | -0.454 | 0.000307611 |
| Nsp8 | OSM | -0.454 | 0.000305745 |
| Nsp8 | GRK5 | -0.454 | 0.000307709 |

| | | | |
|---|---|---|---|
| Nsp10 | MAP2K3 | -0.454 | 0.00030776 |
| Nsp10 | FGF6 | -0.454 | 0.000302205 |
| Nsp10 | WNT3 | -0.454 | 0.000304684 |
| Nsp12 | TUBA3E | -0.454 | 0.000308657 |
| Nsp12 | CXCL1 | -0.454 | 0.000303451 |
| Nsp13 | ELMO1 | -0.454 | 0.000301677 |
| Nsp13 | PLA2G4F | -0.454 | 0.000299736 |
| Nsp13 | PTPN5 | -0.454 | 0.000299103 |
| Nsp14 | PRKCZ | -0.454 | 0.000304824 |
| Nsp14 | ACVR2B | -0.454 | 0.000308021 |
| Nsp14 | ACVRL1 | -0.454 | 0.000306114 |
| Nsp16 | STAT5B | -0.454 | 0.000302335 |
| Nsp16 | IL11RA | -0.454 | 0.00030087 |
| Orf3a | CACNG1 | -0.454 | 0.000307622 |
| Orf3a | ID4 | -0.454 | 0.000302854 |
| Orf3a | HSPA1B | -0.454 | 0.000303704 |
| Orf3a | PRKACG | -0.454 | 0.000302725 |
| Orf8 | NTRK1 | -0.454 | 0.000305663 |
| Orf8 | VWF | -0.454 | 0.00030876 |
| Nsp2 | LIF | -0.455 | 0.000294464 |
| Nsp2 | ULK1 | -0.455 | 0.000289262 |
| Nsp3 | CLCF1 | -0.455 | 0.000289185 |
| Nsp3 | GNB5 | -0.455 | 0.000289503 |
| Nsp4 | RELA | -0.455 | 0.000288891 |
| Nsp4 | PKN3 | -0.455 | 0.000295024 |
| Nsp4 | IGF2 | -0.455 | 0.000289553 |
| Nsp4 | MAPKAPK2 | -0.455 | 0.000296096 |
| Nsp6 | PIM1 | -0.455 | 0.000296558 |
| Nsp7 | SHC2 | -0.455 | 0.000292947 |
| Nsp7 | DUSP9 | -0.455 | 0.000292323 |
| Nsp7 | FGFR3 | -0.455 | 0.000289966 |
| Nsp8 | FOXP3 | -0.455 | 0.000292307 |
| Nsp9 | PPP1R3D | -0.455 | 0.000291319 |
| Nsp10 | RAC2 | -0.455 | 0.000297315 |
| Nsp10 | FZD5 | -0.455 | 0.00029146 |
| Nsp10 | MAP2K2 | -0.455 | 0.000294537 |
| Nsp12 | ADCY8 | -0.455 | 0.000294826 |
| Nsp12 | MMP14 | -0.455 | 0.000289632 |
| Nsp13 | TRAF1 | -0.455 | 0.00029394 |
| Nsp13 | ACSBG1 | -0.455 | 0.000294693 |
| Nsp13 | FGFR1 | -0.455 | 0.000294033 |
| Nsp14 | AKT1S1 | -0.455 | 0.000296971 |
| Nsp14 | NFATC2 | -0.455 | 0.000298063 |
| Nsp15 | DFFB | -0.455 | 0.000294639 |
| Nsp15 | ITGA7 | -0.455 | 0.000298342 |
| Nsp16 | INHBA | -0.455 | 0.000292286 |
| Nsp16 | IL17RE | -0.455 | 0.00029166 |
| Nsp16 | PRL | -0.455 | 0.000296284 |
| Nsp16 | IKBKG | -0.455 | 0.000290327 |
| S | PLA2G4B | -0.455 | 0.000292723 |

| | | | |
|---|---|---|---|
| S | IRF3 | -0.455 | 0.000294268 |
| S | HK2 | -0.455 | 0.000295928 |
| Orf3a | GNG13 | -0.455 | 0.000293761 |
| Orf3a | TOLLIP | -0.455 | 0.000291348 |
| Orf3a | INHBB | -0.455 | 0.00029177 |
| Orf8 | CTSF | -0.455 | 0.000294967 |
| Orf8 | ITGB5 | -0.455 | 0.000294952 |
| Orf8 | FOSL1 | -0.455 | 0.000297566 |
| Orf8 | IGF2 | -0.455 | 0.000289022 |
| Orf8 | SREBF1 | -0.455 | 0.000292281 |
| Orf8 | CACNA2D2 | -0.455 | 0.000290417 |
| Nsp2 | WNT3A | -0.456 | 0.000279993 |
| Nsp2 | ITGB4 | -0.456 | 0.000279233 |
| Nsp3 | FOXO1 | -0.456 | 0.000280408 |
| Nsp3 | ARRB2 | -0.456 | 0.000283308 |
| Nsp3 | ID2 | -0.456 | 0.000281455 |
| Nsp3 | TNFRSF1A | -0.456 | 0.000288385 |
| Nsp5 | FGR | -0.456 | 0.000284544 |
| Nsp6 | PRKAR1B | -0.456 | 0.000286138 |
| Nsp6 | LEFTY2 | -0.456 | 0.000286253 |
| Nsp6 | COMP | -0.456 | 0.000281496 |
| Nsp6 | GDF11 | -0.456 | 0.000281181 |
| Nsp7 | CD14 | -0.456 | 0.000283401 |
| Nsp7 | PIK3CD | -0.456 | 0.000280107 |
| Nsp7 | IL36RN | -0.456 | 0.000287436 |
| Nsp8 | PLA2G4D | -0.456 | 0.000279564 |
| Nsp8 | CPT1B | -0.456 | 0.000284631 |
| Nsp8 | RAPGEF1 | -0.456 | 0.000288373 |
| Nsp9 | DDIT4 | -0.456 | 0.000287 |
| Nsp9 | WNT7A | -0.456 | 0.000281064 |
| Nsp10 | RGMA | -0.456 | 0.000283783 |
| Nsp10 | GRK2 | -0.456 | 0.000284585 |
| Nsp10 | CREB3L1 | -0.456 | 0.000285158 |
| Nsp10 | COL6A1 | -0.456 | 0.000283511 |
| Nsp10 | ZAP70 | -0.456 | 0.000284936 |
| Nsp12 | ACVR1B | -0.456 | 0.000286185 |
| Nsp12 | G6PC3 | -0.456 | 0.000288645 |
| Nsp14 | PPP2R5D | -0.456 | 0.000282221 |
| Nsp14 | IL27RA | -0.456 | 0.000285437 |
| Nsp14 | INHBA | -0.456 | 0.000287876 |
| Nsp14 | GRK5 | -0.456 | 0.000284387 |
| Nsp16 | TIRAP | -0.456 | 0.000285672 |
| Nsp16 | CSF2 | -0.456 | 0.000279817 |
| S | GNB3 | -0.456 | 0.000283921 |
| S | ERBB2 | -0.456 | 0.000281319 |
| Orf3a | FLT4 | -0.456 | 0.000282014 |
| Orf3a | ICAM1 | -0.456 | 0.000285847 |
| Orf3a | WNT5B | -0.456 | 0.000288722 |
| Orf3a | FZD1 | -0.456 | 0.000284419 |
| Orf6 | SHC2 | -0.456 | 0.000283537 |

| | | | |
|---|---|---|---|
| Orf6 | LMNB2 | -0.456 | 0.000282972 |
| Orf6 | WNT1 | -0.456 | 0.000288539 |
| Orf8 | PARP3 | -0.456 | 0.000287527 |
| Orf8 | GYS1 | -0.456 | 0.000285236 |
| Orf8 | EIF4E1B | -0.456 | 0.000287982 |
| Orf8 | INHBA | -0.456 | 0.000279161 |
| Orf8 | JAK3 | -0.456 | 0.000279459 |
| Orf10 | PTPN7 | -0.456 | 0.000287276 |
| Nsp3 | FOS | -0.457 | 0.000270441 |
| Nsp3 | HK2 | -0.457 | 0.000270796 |
| Nsp4 | NPRL2 | -0.457 | 0.000278564 |
| Nsp4 | GNB5 | -0.457 | 0.00027196 |
| Nsp4 | G6PC3 | -0.457 | 0.000271382 |
| Nsp5 | CHRM1 | -0.457 | 0.000271088 |
| Nsp5 | ITGA11 | -0.457 | 0.000276139 |
| Nsp5 | CAMK2A | -0.457 | 0.000270614 |
| Nsp6 | ENDOG | -0.457 | 0.000277534 |
| Nsp6 | IRF7 | -0.457 | 0.000274477 |
| Nsp6 | SMAD6 | -0.457 | 0.000273852 |
| Nsp7 | TBKBP1 | -0.457 | 0.000276992 |
| Nsp7 | YWHAG | -0.457 | 0.000270916 |
| Nsp8 | RORC | -0.457 | 0.000277781 |
| Nsp8 | NFKBIA | -0.457 | 0.000277414 |
| Nsp9 | RGMA | -0.457 | 0.00027714 |
| Nsp9 | JUND | -0.457 | 0.000273233 |
| Nsp9 | NFATC1 | -0.457 | 0.000274814 |
| Nsp10 | CACNG6 | -0.457 | 0.000278737 |
| Nsp10 | DUSP7 | -0.457 | 0.000278831 |
| Nsp10 | MAPK8IP2 | -0.457 | 0.000270607 |
| Nsp10 | GRK1 | -0.457 | 0.000272732 |
| Nsp12 | GNB5 | -0.457 | 0.000272511 |
| Nsp13 | GH1 | -0.457 | 0.000269857 |
| Nsp13 | ITGA7 | -0.457 | 0.000275859 |
| Nsp14 | CSF2RB | -0.457 | 0.000277649 |
| Nsp14 | VHL | -0.457 | 0.000271897 |
| Nsp15 | MAPKAPK2 | -0.457 | 0.000269776 |
| Nsp16 | NFKB2 | -0.457 | 0.000274435 |
| Nsp16 | GH2 | -0.457 | 0.000272869 |
| Nsp16 | LTB | -0.457 | 0.00027802 |
| S | THSD4 | -0.457 | 0.000270402 |
| S | EDAR | -0.457 | 0.000273098 |
| S | TNFRSF12A | -0.457 | 0.000276916 |
| Orf3a | CACNG6 | -0.457 | 0.000273286 |
| Orf3a | DVL1 | -0.457 | 0.000276542 |
| Orf3a | WNT11 | -0.457 | 0.000276458 |
| Orf3a | FGF3 | -0.457 | 0.000271118 |
| Orf8 | FLCN | -0.457 | 0.000271161 |
| Nsp2 | TCL1A | -0.458 | 0.000264972 |
| Nsp3 | GNG8 | -0.458 | 0.000264613 |
| Nsp3 | NFKBIA | -0.458 | 0.000261548 |

| | | | |
|---|---|---|---|
| Nsp4 | THSD4 | -0.458 | 0.000266675 |
| Nsp4 | TNFRSF12A | -0.458 | 0.000261073 |
| Nsp4 | TNFRSF1A | -0.458 | 0.000268156 |
| Nsp5 | PPP2R5B | -0.458 | 0.000266116 |
| Nsp6 | NFATC1 | -0.458 | 0.00026589 |
| Nsp7 | ADCY6 | -0.458 | 0.000264462 |
| Nsp8 | NPRL3 | -0.458 | 0.000269211 |
| Nsp8 | NPRL2 | -0.458 | 0.000265002 |
| Nsp8 | ITGB3 | -0.458 | 0.000268897 |
| Nsp8 | SLC2A1 | -0.458 | 0.000260417 |
| Nsp9 | GDF10 | -0.458 | 0.000260834 |
| Nsp9 | DUSP9 | -0.458 | 0.000267462 |
| Nsp9 | NCF1 | -0.458 | 0.000266422 |
| Nsp10 | ELAVL1 | -0.458 | 0.000263639 |
| Nsp10 | WNT5B | -0.458 | 0.000268595 |
| Nsp10 | CACNA1A | -0.458 | 0.000267084 |
| Nsp10 | BAD | -0.458 | 0.000268372 |
| Nsp12 | CHRM1 | -0.458 | 0.00026739 |
| Nsp12 | TUBA8 | -0.458 | 0.000261451 |
| Nsp12 | GNB3 | -0.458 | 0.000261942 |
| Nsp12 | HK2 | -0.458 | 0.000268295 |
| Nsp12 | SLC2A1 | -0.458 | 0.000264052 |
| Nsp13 | CCL14 | -0.458 | 0.000269154 |
| Nsp13 | TUBA8 | -0.458 | 0.000267029 |
| Nsp13 | IKBKG | -0.458 | 0.000266744 |
| Nsp13 | G6PC3 | -0.458 | 0.000268754 |
| Nsp14 | GREM1 | -0.458 | 0.00026808 |
| Nsp14 | SREBF1 | -0.458 | 0.00026164 |
| Nsp14 | WNT5A | -0.458 | 0.000269178 |
| Nsp15 | IL4 | -0.458 | 0.000264042 |
| Orf3a | GRK1 | -0.458 | 0.000263762 |
| Orf8 | MLST8 | -0.458 | 0.000261861 |
| Orf8 | IFNL2 | -0.458 | 0.000267607 |
| Orf8 | PIN1 | -0.458 | 0.000261149 |
| Orf10 | DUSP16 | -0.458 | 0.000268143 |
| Nsp2 | POMC | -0.459 | 0.00025834 |
| Nsp3 | ADCY2 | -0.459 | 0.000251417 |
| Nsp4 | RELT | -0.459 | 0.000258088 |
| Nsp4 | TNF | -0.459 | 0.000253187 |
| Nsp4 | ADCY6 | -0.459 | 0.000257774 |
| Nsp6 | RPS6KA4 | -0.459 | 0.00025354 |
| Nsp7 | TAB1 | -0.459 | 0.000253952 |
| Nsp8 | IRS1 | -0.459 | 0.000255102 |
| Nsp8 | PARP3 | -0.459 | 0.000255204 |
| Nsp8 | MAPK15 | -0.459 | 0.000258147 |
| Nsp8 | S100A9 | -0.459 | 0.000254152 |
| Nsp9 | WNT3A | -0.459 | 0.00025447 |
| Nsp9 | JUN | -0.459 | 0.000257933 |
| Nsp9 | FGF22 | -0.459 | 0.000257023 |
| Nsp12 | ERN1 | -0.459 | 0.000259786 |

| | | | |
|---|---|---|---|
| Nsp12 | HK3 | -0.459 | 0.000255891 |
| Nsp13 | IL21R | -0.459 | 0.00025827 |
| Nsp14 | DUSP2 | -0.459 | 0.000257661 |
| Nsp14 | FOXP3 | -0.459 | 0.000256397 |
| Nsp15 | LBP | -0.459 | 0.000252243 |
| Nsp15 | MAPK13 | -0.459 | 0.000258509 |
| S | CCND2 | -0.459 | 0.000252139 |
| Orf3a | GRK2 | -0.459 | 0.000256767 |
| Orf3a | TNFRSF18 | -0.459 | 0.000251481 |
| Orf3a | FZD5 | -0.459 | 0.000256708 |
| Orf6 | TRIP10 | -0.459 | 0.000256324 |
| Orf8 | TIRAP | -0.459 | 0.000257448 |
| Nsp3 | CSF1R | -0.46 | 0.000248008 |
| Nsp3 | CTSF | -0.46 | 0.000243167 |
| Nsp3 | RNF152 | -0.46 | 0.000244617 |
| Nsp3 | TUBA3E | -0.46 | 0.000246677 |
| Nsp3 | CAMK2A | -0.46 | 0.000242763 |
| Nsp3 | FGF23 | -0.46 | 0.000244059 |
| Nsp4 | HMOX1 | -0.46 | 0.000247589 |
| Nsp5 | RASGRP4 | -0.46 | 0.000243047 |
| Nsp5 | GADD45G | -0.46 | 0.000244067 |
| Nsp5 | CNTFR | -0.46 | 0.000251137 |
| Nsp6 | DVL1 | -0.46 | 0.000244345 |
| Nsp10 | PLCG2 | -0.46 | 0.000246701 |
| Nsp10 | RASGRP2 | -0.46 | 0.000243036 |
| Nsp12 | PDGFA | -0.46 | 0.000243234 |
| Nsp12 | PIDD1 | -0.46 | 0.000249065 |
| Nsp12 | IL15RA | -0.46 | 0.000247178 |
| Nsp13 | IL2RB | -0.46 | 0.000247417 |
| Nsp13 | LTBR | -0.46 | 0.000250058 |
| Nsp13 | PPP2R5B | -0.46 | 0.000248045 |
| Nsp14 | HMOX1 | -0.46 | 0.000243236 |
| Nsp14 | GNB2 | -0.46 | 0.000249811 |
| Nsp15 | RASGRP4 | -0.46 | 0.000245765 |
| Nsp15 | S100A9 | -0.46 | 0.000245544 |
| Nsp15 | VAV1 | -0.46 | 0.000248689 |
| Nsp15 | PIK3R6 | -0.46 | 0.000244263 |
| Nsp16 | SEC13 | -0.46 | 0.000242625 |
| Nsp16 | ARRB2 | -0.46 | 0.000247823 |
| S | CSH2 | -0.46 | 0.00024696 |
| Orf3a | GDF7 | -0.46 | 0.000249474 |
| Orf6 | CARD11 | -0.46 | 0.000250697 |
| Orf8 | PIK3R5 | -0.46 | 0.000245451 |
| Orf8 | TNFSF9 | -0.46 | 0.000246513 |
| Orf8 | IKBKE | -0.46 | 0.000243449 |
| Orf8 | ADCY4 | -0.46 | 0.000244725 |
| Orf10 | PHLPP1 | -0.46 | 0.00024951 |
| Nsp3 | TUBA8 | -0.461 | 0.000238019 |
| Nsp3 | CACNA1E | -0.461 | 0.000242147 |
| Nsp4 | CCR7 | -0.461 | 0.000239686 |

| | | | |
|---|---|---|---|
| Nsp4 | CDC37 | -0.461 | 0.000241725 |
| Nsp4 | ADCY2 | -0.461 | 0.000234756 |
| Nsp5 | ACTB | -0.461 | 0.000238366 |
| Nsp5 | CALML5 | -0.461 | 0.00023662 |
| Nsp5 | SLC2A1 | -0.461 | 0.000238362 |
| Nsp5 | WNT5A | -0.461 | 0.000235403 |
| Nsp7 | SRF | -0.461 | 0.000234501 |
| Nsp7 | PIK3R2 | -0.461 | 0.000242006 |
| Nsp7 | GADD45B | -0.461 | 0.000239272 |
| Nsp7 | FZD1 | -0.461 | 0.00023646 |
| Nsp9 | FZD2 | -0.461 | 0.000238735 |
| Nsp10 | IL27RA | -0.461 | 0.000241018 |
| Nsp12 | LBP | -0.461 | 0.000242308 |
| Nsp13 | HMOX1 | -0.461 | 0.000240821 |
| Nsp14 | GNG13 | -0.461 | 0.000236143 |
| Nsp15 | CALML6 | -0.461 | 0.000234181 |
| S | NFKB2 | -0.461 | 0.000238695 |
| M | GDF15 | -0.461 | 0.000241275 |
| Orf8 | EPHA2 | -0.461 | 0.00023852 |
| Orf8 | TAB1 | -0.461 | 0.000239268 |
| Nsp2 | RELB | -0.462 | 0.000226168 |
| Nsp3 | PRKAB1 | -0.462 | 0.000233435 |
| Nsp4 | SREBF1 | -0.462 | 0.000227322 |
| Nsp5 | PHLPP1 | -0.462 | 0.00023223 |
| Nsp5 | THBS2 | -0.462 | 0.000234103 |
| Nsp6 | BMP6 | -0.462 | 0.000232765 |
| Nsp6 | MAPK8IP2 | -0.462 | 0.000227678 |
| Nsp7 | TNFSF12 | -0.462 | 0.000233147 |
| Nsp7 | NGFR | -0.462 | 0.000227886 |
| Nsp7 | EFNA2 | -0.462 | 0.000226411 |
| Nsp7 | JUN | -0.462 | 0.000226201 |
| Nsp7 | EGLN1 | -0.462 | 0.000226605 |
| Nsp8 | TUBA3C | -0.462 | 0.000230685 |
| Nsp8 | ITGA7 | -0.462 | 0.000229976 |
| Nsp10 | IL21R | -0.462 | 0.000232802 |
| Nsp10 | IL17RA | -0.462 | 0.000228452 |
| Nsp10 | TICAM1 | -0.462 | 0.000230592 |
| Nsp12 | PPP2R5D | -0.462 | 0.00022692 |
| Nsp12 | GH2 | -0.462 | 0.000233013 |
| Nsp13 | FGF21 | -0.462 | 0.000231655 |
| Nsp14 | THSD4 | -0.462 | 0.0002302 |
| Nsp14 | ACSBG1 | -0.462 | 0.000231018 |
| Nsp15 | TGFB3 | -0.462 | 0.000229531 |
| Nsp15 | PPP5C | -0.462 | 0.000227639 |
| Nsp15 | GDF5 | -0.462 | 0.000226259 |
| S | PLCG2 | -0.462 | 0.000227287 |
| Orf3a | TNFRSF6B | -0.462 | 0.00022774 |
| Orf3a | FGFR3 | -0.462 | 0.000230194 |
| Orf6 | IRS2 | -0.462 | 0.000230408 |
| Orf6 | VEGFA | -0.462 | 0.000227059 |

| | | | |
|---|---|---|---|
| Orf8 | TRAF1 | -0.462 | 0.000228948 |
| Orf8 | ITGA3 | -0.462 | 0.000226455 |
| Orf8 | DVL2 | -0.462 | 0.000232173 |
| Orf8 | IGF1R | -0.462 | 0.000232861 |
| Orf8 | WNT9A | -0.462 | 0.000231496 |
| Orf8 | WNT8B | -0.462 | 0.000228883 |
| Nsp2 | FGF17 | -0.463 | 0.000224033 |
| Nsp2 | CACNA1A | -0.463 | 0.0002223 |
| Nsp3 | RASGRP4 | -0.463 | 0.000225791 |
| Nsp4 | WNT10B | -0.463 | 0.000223817 |
| Nsp4 | IFNL2 | -0.463 | 0.000221305 |
| Nsp5 | OSM | -0.463 | 0.000221833 |
| Nsp5 | PRKACA | -0.463 | 0.000222896 |
| Nsp5 | PFKL | -0.463 | 0.00022593 |
| Nsp6 | MAP2K7 | -0.463 | 0.000220645 |
| Nsp7 | GDF7 | -0.463 | 0.000223071 |
| Nsp7 | PPP1R3E | -0.463 | 0.000223237 |
| Nsp8 | IRF5 | -0.463 | 0.000224933 |
| Nsp8 | CREB3L3 | -0.463 | 0.000223991 |
| Nsp8 | TNFSF12 | -0.463 | 0.000220825 |
| Nsp8 | PRKCD | -0.463 | 0.000224616 |
| Nsp8 | ACVRL1 | -0.463 | 0.000219711 |
| Nsp10 | ATP6V1B1 | -0.463 | 0.000218649 |
| Nsp12 | ACTB | -0.463 | 0.000223363 |
| Nsp12 | IL21R | -0.463 | 0.000219894 |
| Nsp12 | PIK3R6 | -0.463 | 0.000223924 |
| Nsp13 | RAC2 | -0.463 | 0.000220744 |
| Nsp14 | PIK3R5 | -0.463 | 0.000221653 |
| Nsp14 | IL21R | -0.463 | 0.000223623 |
| Nsp15 | ID3 | -0.463 | 0.000220492 |
| Nsp15 | PRL | -0.463 | 0.000221289 |
| Nsp15 | ADCY6 | -0.463 | 0.000220724 |
| Nsp16 | HMOX1 | -0.463 | 0.000223121 |
| S | HMOX1 | -0.463 | 0.000220839 |
| S | NODAL | -0.463 | 0.000225884 |
| S | WNT8B | -0.463 | 0.000221862 |
| S | ELOB | -0.463 | 0.000223451 |
| Orf3a | SMAD6 | -0.463 | 0.000219331 |
| Orf3a | ADCY7 | -0.463 | 0.000221951 |
| Orf8 | PXN | -0.463 | 0.000219077 |
| Nsp2 | CARD10 | -0.464 | 0.000215926 |
| Nsp2 | PLCB3 | -0.464 | 0.000216927 |
| Nsp3 | IL27RA | -0.464 | 0.000213579 |
| Nsp3 | RORC | -0.464 | 0.000213787 |
| Nsp3 | ITGB5 | -0.464 | 0.000213934 |
| Nsp4 | CCND3 | -0.464 | 0.000210686 |
| Nsp4 | DUSP3 | -0.464 | 0.000212389 |
| Nsp4 | PPP2R5D | -0.464 | 0.000215362 |
| Nsp4 | CHRM1 | -0.464 | 0.000212321 |
| Nsp4 | PRKCD | -0.464 | 0.000211291 |

| | | | |
|---|---|---|---|
| Nsp4 | ELAVL1 | -0.464 | 0.000213411 |
| Nsp4 | INHBA | -0.464 | 0.000211676 |
| Nsp4 | TIMP1 | -0.464 | 0.000216974 |
| Nsp5 | IGF1R | -0.464 | 0.000215304 |
| Nsp5 | ECSIT | -0.464 | 0.000217674 |
| Nsp5 | CACNB1 | -0.464 | 0.000213031 |
| Nsp5 | IKBKE | -0.464 | 0.000216014 |
| Nsp5 | ITGA2B | -0.464 | 0.000213607 |
| Nsp5 | BBC3 | -0.464 | 0.000216157 |
| Nsp6 | LPAR5 | -0.464 | 0.000212697 |
| Nsp6 | IL17RA | -0.464 | 0.000210736 |
| Nsp7 | CEBPB | -0.464 | 0.00021257 |
| Nsp7 | DDIT4 | -0.464 | 0.000217752 |
| Nsp7 | PRR5 | -0.464 | 0.000214684 |
| Nsp9 | CACNG7 | -0.464 | 0.000215051 |
| Nsp10 | ADCY1 | -0.464 | 0.000211156 |
| Nsp10 | FGF3 | -0.464 | 0.000213542 |
| Nsp10 | CXCL12 | -0.464 | 0.000215029 |
| Nsp12 | RELT | -0.464 | 0.000213877 |
| Nsp12 | TGFB3 | -0.464 | 0.000213339 |
| Nsp13 | SLC2A1 | -0.464 | 0.000213678 |
| Nsp14 | THBS2 | -0.464 | 0.000216321 |
| Nsp15 | CSF1R | -0.464 | 0.000217583 |
| Nsp15 | CACNA2D4 | -0.464 | 0.000215972 |
| Nsp15 | G6PC3 | -0.464 | 0.000216509 |
| Nsp16 | ITGB5 | -0.464 | 0.000214947 |
| S | EIF4EBP1 | -0.464 | 0.000211731 |
| Orf3a | SOCS1 | -0.464 | 0.000213997 |
| Orf3a | AKT1S1 | -0.464 | 0.000215845 |
| Orf3a | HSPA1A | -0.464 | 0.000216589 |
| Orf6 | ENDOG | -0.464 | 0.000214508 |
| Orf6 | GDF15 | -0.464 | 0.000211202 |
| Orf8 | FGFR4 | -0.464 | 0.000211193 |
| Orf8 | TBX21 | -0.464 | 0.000217905 |
| Nsp2 | IRF7 | -0.465 | 0.000208356 |
| Nsp2 | FGF19 | -0.465 | 0.000209801 |
| Nsp2 | MYC | -0.465 | 0.000207293 |
| Nsp2 | PRKCG | -0.465 | 0.000206739 |
| Nsp2 | SH2B2 | -0.465 | 0.00020862 |
| Nsp2 | LAMA5 | -0.465 | 0.000205086 |
| Nsp3 | TNFSF12 | -0.465 | 0.000205901 |
| Nsp3 | EIF4EBP1 | -0.465 | 0.000207955 |
| Nsp3 | CACNG2 | -0.465 | 0.000210384 |
| Nsp4 | SHC3 | -0.465 | 0.000208807 |
| Nsp4 | MAX | -0.465 | 0.000206774 |
| Nsp4 | PTPN5 | -0.465 | 0.000209557 |
| Nsp4 | VWF | -0.465 | 0.00020974 |
| Nsp5 | PTK2B | -0.465 | 0.000209105 |
| Nsp5 | PPP3R2 | -0.465 | 0.000209645 |
| Nsp5 | FGFR1 | -0.465 | 0.000208491 |

| | | | |
|---|---|---|---|
| Nsp5 | TNFRSF1A | -0.465 | 0.000204051 |
| Nsp6 | PPP2R3B | -0.465 | 0.000207943 |
| Nsp7 | GRK2 | -0.465 | 0.000208583 |
| Nsp7 | DUSP8 | -0.465 | 0.000209849 |
| Nsp7 | RPS6KA2 | -0.465 | 0.000208592 |
| Nsp7 | SMAD3 | -0.465 | 0.000203234 |
| Nsp8 | JMJD7-PLA2G | -0.465 | 0.000207275 |
| Nsp8 | IKBKE | -0.465 | 0.000204664 |
| Nsp9 | LRP5 | -0.465 | 0.000210349 |
| Nsp9 | WNT5B | -0.465 | 0.000206248 |
| Nsp10 | TRAF2 | -0.465 | 0.000208172 |
| Nsp10 | CACNA1I | -0.465 | 0.000208683 |
| Nsp12 | RNF152 | -0.465 | 0.000207471 |
| Nsp13 | CSF1R | -0.465 | 0.000205465 |
| Nsp13 | CACNB3 | -0.465 | 0.000209037 |
| Nsp14 | FGR | -0.465 | 0.000209543 |
| Nsp14 | EIF4EBP1 | -0.465 | 0.000207857 |
| Nsp14 | NODAL | -0.465 | 0.00020451 |
| Nsp14 | CSH2 | -0.465 | 0.000209726 |
| Nsp14 | RAPGEF1 | -0.465 | 0.000203346 |
| Nsp15 | IL2RB | -0.465 | 0.000206636 |
| Nsp15 | IGF2 | -0.465 | 0.000208013 |
| Nsp15 | SREBF1 | -0.465 | 0.000203146 |
| Nsp16 | LBP | -0.465 | 0.00020496 |
| S | IRS1 | -0.465 | 0.000209182 |
| S | DUSP3 | -0.465 | 0.000209484 |
| S | TNFRSF1A | -0.465 | 0.000204175 |
| Orf3a | HSPA2 | -0.465 | 0.000210382 |
| Orf3a | DUSP4 | -0.465 | 0.000208453 |
| Orf3a | MAPK13 | -0.465 | 0.000209059 |
| Orf8 | CAMKK2 | -0.465 | 0.00020606 |
| Orf8 | DVL3 | -0.465 | 0.00020425 |
| Orf8 | IRAK1 | -0.465 | 0.0002103 |
| Orf8 | WDR24 | -0.465 | 0.000206868 |
| Orf8 | CXCL14 | -0.465 | 0.000204993 |
| Orf8 | FOXP3 | -0.465 | 0.000203292 |
| Nsp2 | CHRD | -0.466 | 0.000196267 |
| Nsp2 | VAV2 | -0.466 | 0.000197667 |
| Nsp2 | CAPN1 | -0.466 | 0.000196246 |
| Nsp3 | NFKB2 | -0.466 | 0.000201863 |
| Nsp3 | FGF6 | -0.466 | 0.000197206 |
| Nsp3 | PRKCD | -0.466 | 0.0002008 |
| Nsp4 | IL11 | -0.466 | 0.000202951 |
| Nsp4 | SESN2 | -0.466 | 0.00020197 |
| Nsp5 | IRS1 | -0.466 | 0.000203063 |
| Nsp7 | EFNA3 | -0.466 | 0.000199113 |
| Nsp7 | PPP1R3D | -0.466 | 0.000200708 |
| Nsp8 | CCND3 | -0.466 | 0.000197578 |
| Nsp8 | ECSIT | -0.466 | 0.000196533 |
| Nsp8 | CTSF | -0.466 | 0.000197926 |

| | | | |
|---|---|---|---|
| Nsp8 | CXCL2 | -0.466 | 0.000196899 |
| Nsp12 | ACSBG1 | -0.466 | 0.000202379 |
| Nsp12 | CAMK2A | -0.466 | 0.000196729 |
| Nsp13 | TUBA3C | -0.466 | 0.000198386 |
| Nsp14 | STAT5A | -0.466 | 0.000202248 |
| Nsp14 | CHRD | -0.466 | 0.000201519 |
| Nsp14 | TRIP10 | -0.466 | 0.000201506 |
| Nsp16 | CREB3L3 | -0.466 | 0.000201472 |
| S | JMJD7-PLA2G | -0.466 | 0.000200736 |
| S | PLA2G4D | -0.466 | 0.000202744 |
| Orf3a | PPP2R3B | -0.466 | 0.000202201 |
| Orf6 | INHBA | -0.466 | 0.000202257 |
| Orf8 | TUBA8 | -0.466 | 0.000202799 |
| Orf8 | TRIP10 | -0.466 | 0.000199603 |
| Orf8 | TYK2 | -0.466 | 0.000198056 |
| Orf8 | CNTFR | -0.466 | 0.00020303 |
| Orf8 | TNFRSF13B | -0.466 | 0.000201742 |
| Nsp3 | CX3CL1 | -0.467 | 0.000191608 |
| Nsp3 | IL17B | -0.467 | 0.000194188 |
| Nsp3 | PTPN5 | -0.467 | 0.000189434 |
| Nsp3 | PIK3R6 | -0.467 | 0.000195316 |
| Nsp4 | TRAF1 | -0.467 | 0.000194448 |
| Nsp4 | NPRL3 | -0.467 | 0.000194236 |
| Nsp5 | RAC3 | -0.467 | 0.000194106 |
| Nsp5 | CACNG4 | -0.467 | 0.000192381 |
| Nsp5 | TBX21 | -0.467 | 0.000193419 |
| Nsp6 | WNT6 | -0.467 | 0.000190058 |
| Nsp8 | CACNA1S | -0.467 | 0.00018993 |
| Nsp8 | IRAK1 | -0.467 | 0.000195275 |
| Nsp8 | DHX58 | -0.467 | 0.000190801 |
| Nsp8 | ADCY2 | -0.467 | 0.000190987 |
| Nsp9 | CACNA1I | -0.467 | 0.000195335 |
| Nsp10 | RPS6KA4 | -0.467 | 0.000192005 |
| Nsp10 | GNG13 | -0.467 | 0.000190333 |
| Nsp10 | EDAR | -0.467 | 0.000189445 |
| Nsp10 | TNFRSF6B | -0.467 | 0.000195697 |
| Nsp12 | TNFSF9 | -0.467 | 0.000194077 |
| Nsp12 | CTSF | -0.467 | 0.000189652 |
| Nsp13 | TNFRSF25 | -0.467 | 0.000190989 |
| Nsp13 | CCND2 | -0.467 | 0.00019579 |
| Nsp13 | GNG13 | -0.467 | 0.000194377 |
| Nsp14 | PRKACG | -0.467 | 0.000194138 |
| Nsp15 | IFNL3 | -0.467 | 0.000190933 |
| Nsp15 | TNFSF9 | -0.467 | 0.000191942 |
| Nsp15 | PIDD1 | -0.467 | 0.000194347 |
| Nsp15 | RELA | -0.467 | 0.000191074 |
| Nsp15 | HK3 | -0.467 | 0.000190132 |
| Nsp16 | ID3 | -0.467 | 0.000190775 |
| Nsp16 | FGF23 | -0.467 | 0.000190117 |
| S | GDF5 | -0.467 | 0.000191333 |

| | | | |
|---|---|---|---|
| Orf8 | AKT1 | -0.467 | 0.000189101 |
| Orf8 | TLR9 | -0.467 | 0.000191101 |
| Orf8 | ACTG1 | -0.467 | 0.000192072 |
| Orf8 | ACSBG1 | -0.467 | 0.00019549 |
| Orf8 | GH1 | -0.467 | 0.000193965 |
| Orf8 | MYC | -0.467 | 0.000189809 |
| Orf8 | PFKL | -0.467 | 0.00019501 |
| Orf8 | ITGA7 | -0.467 | 0.00019166 |
| Orf8 | PLAU | -0.467 | 0.000194229 |
| Nsp4 | CACNA1S | -0.468 | 0.000187189 |
| Nsp4 | MAPK12 | -0.468 | 0.000183063 |
| Nsp4 | IRF3 | -0.468 | 0.000187761 |
| Nsp4 | FOXP3 | -0.468 | 0.000188107 |
| Nsp5 | FGF6 | -0.468 | 0.000183221 |
| Nsp5 | FOXO1 | -0.468 | 0.000183744 |
| Nsp5 | CACNA2D4 | -0.468 | 0.000189014 |
| Nsp7 | RPS6KA4 | -0.468 | 0.000187589 |
| Nsp7 | HSPB1 | -0.468 | 0.000187494 |
| Nsp7 | GDF15 | -0.468 | 0.000188538 |
| Nsp7 | MAPK11 | -0.468 | 0.00018275 |
| Nsp8 | TAB1 | -0.468 | 0.000186501 |
| Nsp8 | CLCF1 | -0.468 | 0.000183845 |
| Nsp8 | IL17C | -0.468 | 0.000187655 |
| Nsp8 | ADCY6 | -0.468 | 0.000182928 |
| Nsp9 | LPAR5 | -0.468 | 0.000186311 |
| Nsp9 | INHBB | -0.468 | 0.000184435 |
| Nsp10 | IL34 | -0.468 | 0.000182918 |
| Nsp10 | PGAM5 | -0.468 | 0.000184898 |
| Nsp10 | HSPB1 | -0.468 | 0.000188059 |
| Nsp10 | ADCY9 | -0.468 | 0.000187349 |
| Nsp10 | MMP9 | -0.468 | 0.000186196 |
| Nsp10 | ITGB5 | -0.468 | 0.000186063 |
| Nsp10 | BMP8B | -0.468 | 0.000186607 |
| Nsp12 | CREB3L3 | -0.468 | 0.000188164 |
| Nsp12 | CTSZ | -0.468 | 0.000187918 |
| Nsp12 | EGFR | -0.468 | 0.00018815 |
| Nsp12 | FOSL1 | -0.468 | 0.000186893 |
| Nsp12 | PTPN5 | -0.468 | 0.000182535 |
| Nsp13 | CHRM1 | -0.468 | 0.000184271 |
| Nsp13 | PF4V1 | -0.468 | 0.000188254 |
| Nsp13 | CNTFR | -0.468 | 0.0001889 |
| Nsp14 | CXCR5 | -0.468 | 0.000183929 |
| Nsp14 | PLCB2 | -0.468 | 0.000183299 |
| Nsp14 | CSH1 | -0.468 | 0.000183147 |
| Nsp14 | SOCS3 | -0.468 | 0.000182537 |
| Nsp15 | THSD4 | -0.468 | 0.000184448 |
| Nsp15 | MAP4K1 | -0.468 | 0.000183046 |
| Nsp15 | ARAF | -0.468 | 0.000185391 |
| Nsp16 | TNFRSF12A | -0.468 | 0.000184961 |
| S | AKT1S1 | -0.468 | 0.000186603 |

| | | | |
|---|---|---|---|
| Orf3a | FZD8 | -0.468 | 0.000188665 |
| Orf3a | JUNB | -0.468 | 0.000182942 |
| Orf3a | PTPN6 | -0.468 | 0.000187613 |
| Orf3a | LTB | -0.468 | 0.00018839 |
| Orf3a | EGLN1 | -0.468 | 0.000187026 |
| Orf8 | DFFB | -0.468 | 0.000182561 |
| Orf8 | PFKFB3 | -0.468 | 0.000185749 |
| Nsp2 | MAPK8IP3 | -0.469 | 0.000179275 |
| Nsp2 | TNFRSF12A | -0.469 | 0.000180889 |
| Nsp3 | RELT | -0.469 | 0.000180983 |
| Nsp3 | GNG13 | -0.469 | 0.000182362 |
| Nsp3 | TUBA3D | -0.469 | 0.000176149 |
| Nsp4 | IRAK1 | -0.469 | 0.000177417 |
| Nsp5 | STAT5A | -0.469 | 0.000180476 |
| Nsp5 | EIF4EBP1 | -0.469 | 0.000181238 |
| Nsp5 | GNB3 | -0.469 | 0.000178081 |
| Nsp6 | TNFRSF6B | -0.469 | 0.000180783 |
| Nsp6 | FZD9 | -0.469 | 0.000178298 |
| Nsp8 | GREM1 | -0.469 | 0.000182244 |
| Nsp8 | GFAP | -0.469 | 0.000181421 |
| Nsp8 | SREBF1 | -0.469 | 0.000177569 |
| Nsp10 | PPP3R2 | -0.469 | 0.000176005 |
| Nsp12 | MAP2K3 | -0.469 | 0.000176954 |
| Nsp12 | TNFSF14 | -0.469 | 0.000179926 |
| Nsp12 | CACNA1S | -0.469 | 0.000180189 |
| Nsp12 | TUBA3D | -0.469 | 0.000176373 |
| Nsp12 | VAV1 | -0.469 | 0.00017606 |
| Nsp13 | NFKB2 | -0.469 | 0.000178933 |
| Nsp13 | TGFB3 | -0.469 | 0.000178315 |
| Nsp13 | ANGPT4 | -0.469 | 0.000181329 |
| Nsp13 | ITGB5 | -0.469 | 0.00017822 |
| Nsp13 | HK3 | -0.469 | 0.0001797 |
| Nsp13 | CAMK2A | -0.469 | 0.000177236 |
| Nsp13 | TNFRSF12A | -0.469 | 0.000179907 |
| Nsp14 | SRF | -0.469 | 0.00017797 |
| Nsp15 | IRS1 | -0.469 | 0.000177403 |
| Nsp15 | SEC13 | -0.469 | 0.000179693 |
| Nsp15 | ITGA2B | -0.469 | 0.000181484 |
| S | GNG8 | -0.469 | 0.000179247 |
| S | ACVR2B | -0.469 | 0.000176677 |
| S | HRAS | -0.469 | 0.000181237 |
| S | INS | -0.469 | 0.000175951 |
| Orf3a | PIK3CD | -0.469 | 0.000182084 |
| Orf8 | ITGA11 | -0.469 | 0.000181106 |
| Orf8 | TUBA3D | -0.469 | 0.000176559 |
| Nsp2 | IL34 | -0.47 | 0.000175655 |
| Nsp2 | CACNG4 | -0.47 | 0.000174265 |
| Nsp3 | ACSBG1 | -0.47 | 0.000173147 |
| Nsp3 | ITGA7 | -0.47 | 0.000172477 |
| Nsp3 | ERBB2 | -0.47 | 0.000170643 |

| | | | |
|---|---|---|---|
| Nsp4 | NOS2 | -0.47 | 0.000170177 |
| Nsp4 | ACTG1 | -0.47 | 0.000171561 |
| Nsp4 | CD14 | -0.47 | 0.000175857 |
| Nsp4 | ERBB2 | -0.47 | 0.000174688 |
| Nsp5 | TNFSF12 | -0.47 | 0.00017519 |
| Nsp5 | CPT1C | -0.47 | 0.000170141 |
| Nsp6 | FZD5 | -0.47 | 0.000172901 |
| Nsp6 | GDF1 | -0.47 | 0.000172836 |
| Nsp7 | LMNB2 | -0.47 | 0.000174414 |
| Nsp7 | IRS2 | -0.47 | 0.000174398 |
| Nsp7 | GDF11 | -0.47 | 0.000169653 |
| Nsp8 | PPP3R2 | -0.47 | 0.000171588 |
| Nsp8 | ACTB | -0.47 | 0.000170343 |
| Nsp10 | FLT4 | -0.47 | 0.000175471 |
| Nsp12 | FGF21 | -0.47 | 0.000170273 |
| Nsp13 | SHC3 | -0.47 | 0.000171584 |
| Nsp14 | FLT3LG | -0.47 | 0.000174918 |
| Nsp14 | TCL1A | -0.47 | 0.000173312 |
| Nsp15 | DUSP2 | -0.47 | 0.000173919 |
| Nsp15 | CACNA1S | -0.47 | 0.000175286 |
| Nsp15 | ACTB | -0.47 | 0.000175775 |
| Nsp16 | ELK1 | -0.47 | 0.000170781 |
| Nsp16 | PLCG2 | -0.47 | 0.000171901 |
| Nsp16 | CDC37 | -0.47 | 0.000173577 |
| Nsp16 | CX3CL1 | -0.47 | 0.000174827 |
| S | ACSBG1 | -0.47 | 0.000169754 |
| S | NPRL3 | -0.47 | 0.000170315 |
| S | SLC2A1 | -0.47 | 0.00017295 |
| Orf3a | COMP | -0.47 | 0.000175273 |
| Orf3a | TNFRSF13C | -0.47 | 0.000175667 |
| Orf8 | IFNL3 | -0.47 | 0.000173679 |
| Orf8 | TRADD | -0.47 | 0.000175146 |
| Orf8 | GNAI2 | -0.47 | 0.000174702 |
| Nsp3 | CSF3 | -0.471 | 0.000165065 |
| Nsp4 | TNFSF9 | -0.471 | 0.000165622 |
| Nsp4 | IGF1R | -0.471 | 0.00016481 |
| Nsp4 | IL17B | -0.471 | 0.000166712 |
| Nsp6 | CXCL14 | -0.471 | 0.000169041 |
| Nsp7 | TRADD | -0.471 | 0.000168808 |
| Nsp7 | WNT9A | -0.471 | 0.000164651 |
| Nsp8 | NOS2 | -0.471 | 0.00016703 |
| Nsp8 | RASGRP4 | -0.471 | 0.000168167 |
| Nsp8 | SESN2 | -0.471 | 0.000163872 |
| Nsp8 | CACNG5 | -0.471 | 0.000167459 |
| Nsp10 | TBKBP1 | -0.471 | 0.000167938 |
| Nsp10 | PPARA | -0.471 | 0.000168412 |
| Nsp10 | SLC7A5 | -0.471 | 0.00016516 |
| Nsp12 | FOS | -0.471 | 0.000165845 |
| Nsp12 | SESN2 | -0.471 | 0.000166693 |
| Nsp12 | WNT7A | -0.471 | 0.000165803 |

| | | | |
|---|---|---|---|
| Nsp13 | NBL1 | -0.471 | 0.000166665 |
| Nsp13 | PGF | -0.471 | 0.000164124 |
| Nsp13 | EGLN2 | -0.471 | 0.000168287 |
| Nsp13 | LPAR2 | -0.471 | 0.000164303 |
| Nsp14 | FGFR4 | -0.471 | 0.000168087 |
| Nsp14 | ATP6V1B1 | -0.471 | 0.000166551 |
| Nsp15 | PARP3 | -0.471 | 0.000168782 |
| Nsp15 | GH2 | -0.471 | 0.00016624 |
| Nsp16 | CAMK2G | -0.471 | 0.000163708 |
| Nsp16 | MMP14 | -0.471 | 0.000169074 |
| S | TAB1 | -0.471 | 0.00016503 |
| S | SESN2 | -0.471 | 0.000167251 |
| S | IKBKE | -0.471 | 0.000165929 |
| S | CAMK2A | -0.471 | 0.000165177 |
| Orf3a | PGAM5 | -0.471 | 0.000169342 |
| Orf3a | GDF10 | -0.471 | 0.000165953 |
| Orf8 | TNFSF14 | -0.471 | 0.000164868 |
| Orf8 | HCK | -0.471 | 0.000166846 |
| Orf8 | MAP3K6 | -0.471 | 0.000164447 |
| Orf8 | TRIM25 | -0.471 | 0.000163887 |
| Orf8 | RPS6KB2 | -0.471 | 0.00016621 |
| Nsp2 | FZD7 | -0.472 | 0.000160127 |
| Nsp3 | CCL14 | -0.472 | 0.000161704 |
| Nsp3 | PIDD1 | -0.472 | 0.000161968 |
| Nsp3 | GADD45G | -0.472 | 0.000161623 |
| Nsp3 | MMP14 | -0.472 | 0.000163391 |
| Nsp3 | INS | -0.472 | 0.000162614 |
| Nsp3 | VWF | -0.472 | 0.000158507 |
| Nsp4 | EXOC7 | -0.472 | 0.000162055 |
| Nsp4 | SMAD3 | -0.472 | 0.00016042 |
| Nsp4 | RAPGEF1 | -0.472 | 0.000162251 |
| Nsp5 | TNFRSF25 | -0.472 | 0.000157814 |
| Nsp5 | GADD45A | -0.472 | 0.000159408 |
| Nsp5 | NODAL | -0.472 | 0.000158483 |
| Nsp6 | FZD10 | -0.472 | 0.000163313 |
| Nsp6 | EFNA3 | -0.472 | 0.00016348 |
| Nsp7 | DUSP2 | -0.472 | 0.000161688 |
| Nsp7 | TNFRSF18 | -0.472 | 0.000158269 |
| Nsp8 | IL27 | -0.472 | 0.000160425 |
| Nsp8 | TRAF1 | -0.472 | 0.000160307 |
| Nsp8 | CSF2RB | -0.472 | 0.000161544 |
| Nsp8 | MAP3K11 | -0.472 | 0.000159533 |
| Nsp9 | DUSP8 | -0.472 | 0.000160985 |
| Nsp12 | GDF2 | -0.472 | 0.000160389 |
| Nsp13 | PIDD1 | -0.472 | 0.000158282 |
| Nsp13 | ARRB2 | -0.472 | 0.000161841 |
| Nsp13 | CX3CL1 | -0.472 | 0.00016352 |
| Nsp13 | SOCS7 | -0.472 | 0.000159498 |
| Nsp14 | DHX58 | -0.472 | 0.000161491 |
| Nsp14 | PLA2G4D | -0.472 | 0.000157908 |

| | | | |
|---|---|---|---|
| Nsp15 | PDGFRB | -0.472 | 0.000160511 |
| Nsp15 | TCL1A | -0.472 | 0.000157941 |
| Nsp16 | FOXO1 | -0.472 | 0.00016341 |
| Nsp16 | TNFSF12 | -0.472 | 0.000162971 |
| S | CXCL3 | -0.472 | 0.000162599 |
| S | CACNG2 | -0.472 | 0.000161472 |
| Orf3a | CACNA1I | -0.472 | 0.000163376 |
| Nsp2 | TRIM25 | -0.473 | 0.000153471 |
| Nsp2 | FADD | -0.473 | 0.000154953 |
| Nsp4 | CACNA2D4 | -0.473 | 0.000156586 |
| Nsp4 | EIF4EBP1 | -0.473 | 0.000157416 |
| Nsp4 | PDGFRB | -0.473 | 0.000157347 |
| Nsp4 | ENO3 | -0.473 | 0.000156871 |
| Nsp4 | GNB3 | -0.473 | 0.000153853 |
| Nsp5 | CXCR5 | -0.473 | 0.000155094 |
| Nsp5 | CCND3 | -0.473 | 0.000153966 |
| Nsp5 | CACNA1C | -0.473 | 0.00015378 |
| Nsp5 | PPP2R1A | -0.473 | 0.000155084 |
| Nsp6 | SHC2 | -0.473 | 0.000157706 |
| Nsp6 | AMH | -0.473 | 0.000156689 |
| Nsp7 | FZD9 | -0.473 | 0.000153723 |
| Nsp8 | IRF4 | -0.473 | 0.000156371 |
| Nsp8 | TUBA8 | -0.473 | 0.000155108 |
| Nsp8 | TNF | -0.473 | 0.000153677 |
| Nsp9 | FOXO3 | -0.473 | 0.000155894 |
| Nsp10 | PRKCZ | -0.473 | 0.000157707 |
| Nsp12 | CNTFR | -0.473 | 0.000156378 |
| Nsp12 | SREBF1 | -0.473 | 0.000155508 |
| Nsp12 | ITGA7 | -0.473 | 0.000156983 |
| Nsp13 | CACNA1S | -0.473 | 0.000153376 |
| Nsp13 | ACTB | -0.473 | 0.000153214 |
| Nsp13 | CACNA1E | -0.473 | 0.000154468 |
| Nsp14 | TNFSF14 | -0.473 | 0.000154815 |
| Nsp14 | TSC2 | -0.473 | 0.000156446 |
| Nsp14 | PF4 | -0.473 | 0.00015558 |
| Nsp15 | CRK | -0.473 | 0.000155948 |
| Nsp15 | INS | -0.473 | 0.000152217 |
| Nsp15 | VWF | -0.473 | 0.000154826 |
| Nsp16 | ITGA5 | -0.473 | 0.000157469 |
| S | PTK2B | -0.473 | 0.000152201 |
| S | CACNA2D4 | -0.473 | 0.000153687 |
| S | GADD45G | -0.473 | 0.000153778 |
| S | NFKBIA | -0.473 | 0.000152951 |
| Orf6 | DUSP5 | -0.473 | 0.000153127 |
| Orf6 | CD14 | -0.473 | 0.000156765 |
| Orf6 | DVL1 | -0.473 | 0.000156495 |
| Orf8 | PITX2 | -0.473 | 0.000152615 |
| Orf8 | RRAS | -0.473 | 0.00015628 |
| Orf8 | MAPK15 | -0.473 | 0.000153023 |
| Orf8 | PLCG2 | -0.473 | 0.000156449 |

| | | | |
|---|---|---|---|
| Orf8 | WNT11 | -0.473 | 0.000153031 |
| Nsp2 | GADD45A | -0.474 | 0.00014669 |
| Nsp2 | WNT9A | -0.474 | 0.000149565 |
| Nsp2 | ICAM1 | -0.474 | 0.000148616 |
| Nsp3 | IRS1 | -0.474 | 0.000151249 |
| Nsp3 | TRAF1 | -0.474 | 0.000151438 |
| Nsp3 | ACTB | -0.474 | 0.000149158 |
| Nsp3 | ID3 | -0.474 | 0.000147221 |
| Nsp3 | TNFRSF12A | -0.474 | 0.000150159 |
| Nsp4 | ITGB5 | -0.474 | 0.000148671 |
| Nsp4 | GH2 | -0.474 | 0.000150103 |
| Nsp5 | NOS2 | -0.474 | 0.00015049 |
| Nsp5 | ITGA3 | -0.474 | 0.00015206 |
| Nsp5 | TICAM1 | -0.474 | 0.000147454 |
| Nsp6 | STK11 | -0.474 | 0.000148953 |
| Nsp7 | GREM2 | -0.474 | 0.00014667 |
| Nsp8 | CSH2 | -0.474 | 0.000147779 |
| Nsp10 | PRKAR1B | -0.474 | 0.000148526 |
| Nsp10 | CDC25B | -0.474 | 0.000148734 |
| Nsp10 | RAC3 | -0.474 | 0.000148681 |
| Nsp10 | FZD9 | -0.474 | 0.000147366 |
| Nsp13 | FOS | -0.474 | 0.000151191 |
| Nsp13 | TNFSF9 | -0.474 | 0.000148739 |
| Nsp13 | WNT7A | -0.474 | 0.000151518 |
| Nsp13 | IL17C | -0.474 | 0.000150903 |
| Nsp13 | MAPKAPK2 | -0.474 | 0.000151484 |
| Nsp13 | GNB3 | -0.474 | 0.00014926 |
| Nsp14 | FLCN | -0.474 | 0.000147334 |
| Nsp14 | WDR24 | -0.474 | 0.000147543 |
| Nsp15 | RELT | -0.474 | 0.000146959 |
| Nsp16 | VWF | -0.474 | 0.000149637 |
| Nsp16 | ERBB2 | -0.474 | 0.000147992 |
| S | GREM1 | -0.474 | 0.00014926 |
| S | ITGA2B | -0.474 | 0.000151461 |
| S | ACVRL1 | -0.474 | 0.000147108 |
| S | THBS2 | -0.474 | 0.000147029 |
| Orf3a | LMNB2 | -0.474 | 0.000148655 |
| Orf3a | BMP8A | -0.474 | 0.000149419 |
| Orf3a | BMP7 | -0.474 | 0.000147074 |
| Orf3a | FGF19 | -0.474 | 0.000146627 |
| Orf3a | ADCY5 | -0.474 | 0.000151189 |
| Orf6 | DVL3 | -0.474 | 0.000148935 |
| Orf6 | TNFRSF18 | -0.474 | 0.000150252 |
| Orf8 | CHAD | -0.474 | 0.000147653 |
| Nsp2 | PFKFB3 | -0.475 | 0.000143832 |
| Nsp4 | IL21R | -0.475 | 0.000143022 |
| Nsp5 | MAPK12 | -0.475 | 0.00014434 |
| Nsp5 | WNT10B | -0.475 | 0.000146395 |
| Nsp5 | SOCS7 | -0.475 | 0.000144257 |
| Nsp6 | JUN | -0.475 | 0.000144067 |

| | | | |
|---|---|---|---|
| Nsp8 | FGR | -0.475 | 0.000143896 |
| Nsp8 | PLCG2 | -0.475 | 0.000144063 |
| Nsp10 | COMP | -0.475 | 0.000144966 |
| Nsp12 | GADD45A | -0.475 | 0.000145641 |
| Nsp12 | IRF3 | -0.475 | 0.000144511 |
| Nsp13 | ACVR2B | -0.475 | 0.000141424 |
| Nsp13 | GADD45A | -0.475 | 0.000143943 |
| Nsp14 | IRAK1 | -0.475 | 0.000146451 |
| Nsp15 | ENO3 | -0.475 | 0.000143872 |
| Nsp16 | TGFB3 | -0.475 | 0.000146323 |
| Nsp16 | CTSF | -0.475 | 0.000141485 |
| Nsp16 | GADD45G | -0.475 | 0.000142584 |
| S | PIDD1 | -0.475 | 0.000142527 |
| S | DHX58 | -0.475 | 0.000142404 |
| Orf3a | DUSP6 | -0.475 | 0.000144092 |
| Orf6 | GADD45B | -0.475 | 0.000144368 |
| Orf8 | SRC | -0.475 | 0.000142348 |
| Orf8 | ATP6V1B1 | -0.475 | 0.000144686 |
| Orf8 | CACNA1S | -0.475 | 0.000145466 |
| Orf8 | TNF | -0.475 | 0.000142877 |
| Orf8 | SRF | -0.475 | 0.000142453 |
| Nsp2 | CXCR3 | -0.476 | 0.000137932 |
| Nsp2 | CTF1 | -0.476 | 0.000140202 |
| Nsp3 | CCND2 | -0.476 | 0.000137845 |
| Nsp4 | GREM1 | -0.476 | 0.000139424 |
| Nsp5 | ISG15 | -0.476 | 0.000138121 |
| Nsp5 | EGLN2 | -0.476 | 0.000139097 |
| Nsp6 | NOG | -0.476 | 0.000137052 |
| Nsp8 | GNAI2 | -0.476 | 0.000138878 |
| Nsp8 | IL21R | -0.476 | 0.000136681 |
| Nsp9 | FZD7 | -0.476 | 0.000139859 |
| Nsp9 | GREM2 | -0.476 | 0.0001412 |
| Nsp9 | SMAD6 | -0.476 | 0.0001387 |
| Nsp10 | DDIT4 | -0.476 | 0.000140235 |
| Nsp12 | CSF2RB | -0.476 | 0.000136838 |
| Nsp12 | PKN3 | -0.476 | 0.000140375 |
| Nsp12 | PRKACA | -0.476 | 0.000137593 |
| Nsp13 | RELT | -0.476 | 0.00014017 |
| Nsp13 | IRS1 | -0.476 | 0.000138858 |
| Nsp13 | ICAM1 | -0.476 | 0.000138106 |
| Nsp14 | ID1 | -0.476 | 0.000136898 |
| Nsp14 | MLST8 | -0.476 | 0.000140684 |
| Nsp14 | ACKR3 | -0.476 | 0.000137254 |
| Nsp14 | WNT11 | -0.476 | 0.000137892 |
| Nsp14 | JAK3 | -0.476 | 0.000140446 |
| Nsp15 | CCR7 | -0.476 | 0.000136193 |
| Nsp15 | CACNA1E | -0.476 | 0.000137845 |
| Nsp16 | CACNA1F | -0.476 | 0.000140175 |
| Nsp16 | ELOB | -0.476 | 0.000140305 |
| S | VAV2 | -0.476 | 0.000139692 |

| | | | |
|---|---|---|---|
| Orf3a | CARD14 | -0.476 | 0.000138367 |
| Orf3a | TGFB1 | -0.476 | 0.000141039 |
| Orf8 | ACVR2B | -0.476 | 0.000138198 |
| Orf8 | MUC5B | -0.476 | 0.000137784 |
| Orf8 | WNT5A | -0.476 | 0.000137386 |
| Nsp2 | GADD45B | -0.477 | 0.000132081 |
| Nsp3 | IL12RB1 | -0.477 | 0.000133806 |
| Nsp3 | PRL | -0.477 | 0.000133522 |
| Nsp4 | IL12RB1 | -0.477 | 0.00013425 |
| Nsp6 | IL17D | -0.477 | 0.000135203 |
| Nsp6 | FZD8 | -0.477 | 0.000134856 |
| Nsp6 | DUSP8 | -0.477 | 0.000132681 |
| Nsp6 | EFNA2 | -0.477 | 0.00013484 |
| Nsp8 | TNFRSF1A | -0.477 | 0.000134038 |
| Nsp9 | CXCR3 | -0.477 | 0.000132449 |
| Nsp9 | RXRA | -0.477 | 0.000133554 |
| Nsp9 | DUSP6 | -0.477 | 0.000132462 |
| Nsp10 | CACNA1H | -0.477 | 0.000135784 |
| Nsp10 | TAB1 | -0.477 | 0.000135452 |
| Nsp10 | OSM | -0.477 | 0.000134041 |
| Nsp10 | EFNA2 | -0.477 | 0.00013388 |
| Nsp12 | ELK1 | -0.477 | 0.000133318 |
| Nsp12 | IRF7 | -0.477 | 0.00013151 |
| Nsp12 | CACNA1E | -0.477 | 0.000134883 |
| Nsp12 | DHX58 | -0.477 | 0.000136016 |
| Nsp12 | PIN1 | -0.477 | 0.000132314 |
| Nsp13 | TUBA3E | -0.477 | 0.000131546 |
| Nsp14 | FLNC | -0.477 | 0.000136028 |
| Nsp14 | LTB | -0.477 | 0.000135511 |
| Nsp15 | FOS | -0.477 | 0.000131215 |
| Nsp15 | PPP2R5D | -0.477 | 0.000134852 |
| Nsp15 | ACTG1 | -0.477 | 0.00013193 |
| Nsp15 | ACVR2B | -0.477 | 0.000134378 |
| Nsp16 | IRF5 | -0.477 | 0.000132113 |
| Nsp16 | IKBKE | -0.477 | 0.00013195 |
| Nsp16 | PPP1R3F | -0.477 | 0.00013579 |
| Nsp16 | VEGFB | -0.477 | 0.00013552 |
| S | TSC2 | -0.477 | 0.000135053 |
| S | EGLN2 | -0.477 | 0.000133459 |
| S | CSF3 | -0.477 | 0.000136038 |
| Orf3a | DUSP7 | -0.477 | 0.000132989 |
| Orf3a | BCL2 | -0.477 | 0.000132094 |
| Orf8 | FLT3LG | -0.477 | 0.000131535 |
| Orf8 | GDF5 | -0.477 | 0.000135191 |
| Orf8 | PLCG1 | -0.477 | 0.000133254 |
| Nsp2 | CACNA1G | -0.478 | 0.000128896 |
| Nsp4 | ID3 | -0.478 | 0.000129032 |
| Nsp4 | CSH2 | -0.478 | 0.000127616 |
| Nsp4 | THBS2 | -0.478 | 0.00012693 |
| Nsp5 | ACSBG1 | -0.478 | 0.000128685 |

| | | | |
|---|---|---|---|
| Nsp8 | TUBA3D | -0.478 | 0.000127451 |
| Nsp8 | WNT8B | -0.478 | 0.000129767 |
| Nsp9 | HSPA2 | -0.478 | 0.00012688 |
| Nsp10 | RAPGEF1 | -0.478 | 0.000126937 |
| Nsp12 | TRAF1 | -0.478 | 0.00013084 |
| Nsp12 | PGF | -0.478 | 0.000129321 |
| Nsp12 | RAC2 | -0.478 | 0.000127059 |
| Nsp13 | PRKCD | -0.478 | 0.000130398 |
| Nsp13 | WNT8B | -0.478 | 0.000126464 |
| Nsp14 | LIF | -0.478 | 0.000127682 |
| Nsp15 | TRAF1 | -0.478 | 0.000126648 |
| Nsp16 | CSF1R | -0.478 | 0.000126616 |
| Nsp16 | ANGPT4 | -0.478 | 0.000127218 |
| Nsp16 | CACNB3 | -0.478 | 0.000129194 |
| Nsp16 | HRAS | -0.478 | 0.000129135 |
| Nsp16 | IL17B | -0.478 | 0.000126424 |
| Nsp16 | CAMK2A | -0.478 | 0.000129858 |
| Nsp16 | FGF16 | -0.478 | 0.000127706 |
| Nsp16 | SLC2A1 | -0.478 | 0.000130869 |
| S | CXCL2 | -0.478 | 0.000131005 |
| S | ID3 | -0.478 | 0.000128755 |
| Orf8 | TNXB | -0.478 | 0.000128996 |
| Orf8 | PPP5C | -0.478 | 0.000129659 |
| Orf8 | NPRL3 | -0.478 | 0.000127277 |
| Nsp3 | THSD4 | -0.479 | 0.000122824 |
| Nsp3 | IFNL3 | -0.479 | 0.000122402 |
| Nsp3 | SLC2A1 | -0.479 | 0.000123878 |
| Nsp4 | FGF6 | -0.479 | 0.000126129 |
| Nsp4 | ECSIT | -0.479 | 0.000126166 |
| Nsp7 | FGF4 | -0.479 | 0.000123373 |
| Nsp8 | BBC3 | -0.479 | 0.000124118 |
| Nsp8 | LPAR2 | -0.479 | 0.000125573 |
| Nsp10 | MAPK3 | -0.479 | 0.000126096 |
| Nsp10 | EFNA3 | -0.479 | 0.000124768 |
| Nsp10 | BMP7 | -0.479 | 0.000123614 |
| Nsp10 | ADCY7 | -0.479 | 0.000123082 |
| Nsp12 | NFKB2 | -0.479 | 0.000125272 |
| Nsp12 | CHRD | -0.479 | 0.000125153 |
| Nsp12 | HRAS | -0.479 | 0.000124787 |
| Nsp12 | ADCY2 | -0.479 | 0.000124619 |
| Nsp13 | LBP | -0.479 | 0.000125181 |
| Nsp13 | RPS6KB2 | -0.479 | 0.000122663 |
| Nsp13 | ADCY2 | -0.479 | 0.00012291 |
| Nsp14 | CDC25B | -0.479 | 0.000122098 |
| Nsp14 | GRK6 | -0.479 | 0.000123959 |
| Nsp14 | BCAR1 | -0.479 | 0.000121935 |
| Nsp15 | NTRK1 | -0.479 | 0.000123999 |
| Nsp15 | CCL14 | -0.479 | 0.000125727 |
| Nsp15 | CHRM1 | -0.479 | 0.000126239 |
| Nsp15 | WNT10B | -0.479 | 0.000122129 |

| | | | |
|---|---|---|---|
| Nsp15 | IKBKE | -0.479 | 0.000122614 |
| Nsp15 | PKN3 | -0.479 | 0.000125467 |
| Nsp15 | LTB | -0.479 | 0.000122067 |
| S | PFKP | -0.479 | 0.000123713 |
| Orf8 | DUSP3 | -0.479 | 0.000123937 |
| Orf8 | MAPK13 | -0.479 | 0.000124606 |
| Orf8 | PTPN7 | -0.479 | 0.000123663 |
| Nsp3 | HMOX1 | -0.48 | 0.000118374 |
| Nsp3 | NPRL2 | -0.48 | 0.000121697 |
| Nsp3 | LTB | -0.48 | 0.000118236 |
| Nsp4 | LMNA | -0.48 | 0.000120248 |
| Nsp5 | TAB1 | -0.48 | 0.00011917 |
| Nsp5 | LMNA | -0.48 | 0.000118386 |
| Nsp6 | WNT1 | -0.48 | 0.000121016 |
| Nsp8 | COL9A3 | -0.48 | 0.000121513 |
| Nsp8 | BAD | -0.48 | 0.000117774 |
| Nsp8 | FADD | -0.48 | 0.000118814 |
| Nsp10 | SMAD7 | -0.48 | 0.000118967 |
| Nsp10 | LEFTY2 | -0.48 | 0.000119061 |
| Nsp10 | FGF23 | -0.48 | 0.000121451 |
| Nsp10 | CARD11 | -0.48 | 0.000119193 |
| Nsp12 | ADCY6 | -0.48 | 0.000117587 |
| Nsp13 | GNB5 | -0.48 | 0.000121704 |
| Nsp15 | CXCR5 | -0.48 | 0.000117748 |
| Nsp15 | RRAS | -0.48 | 0.000117589 |
| Nsp15 | CACNB1 | -0.48 | 0.000119445 |
| Nsp16 | NLRX1 | -0.48 | 0.000120994 |
| Nsp16 | ENO3 | -0.48 | 0.000118537 |
| S | PPP3R2 | -0.48 | 0.000118204 |
| S | GNG13 | -0.48 | 0.000120131 |
| S | CHRM1 | -0.48 | 0.00011816 |
| S | PRKACA | -0.48 | 0.000118812 |
| Orf3a | MAPK8IP1 | -0.48 | 0.000121786 |
| Orf6 | COMP | -0.48 | 0.000117743 |
| Orf8 | MAPK8IP1 | -0.48 | 0.000119533 |
| Orf8 | TNFRSF12A | -0.48 | 0.000119374 |
| Orf8 | VEGFA | -0.48 | 0.000121314 |
| Orf8 | RAPGEF1 | -0.48 | 0.000118611 |
| Nsp2 | WNT1 | -0.481 | 0.000116418 |
| Nsp3 | TNFRSF25 | -0.481 | 0.000113575 |
| Nsp4 | IFNL3 | -0.481 | 0.000115585 |
| Nsp4 | PLCG2 | -0.481 | 0.000115739 |
| Nsp4 | ITGA11 | -0.481 | 0.000116557 |
| Nsp4 | GNAI2 | -0.481 | 0.000113809 |
| Nsp4 | PRKACA | -0.481 | 0.00011715 |
| Nsp5 | PIK3R5 | -0.481 | 0.000114714 |
| Nsp5 | MAP3K11 | -0.481 | 0.000113772 |
| Nsp7 | SOCS1 | -0.481 | 0.000116769 |
| Nsp8 | CDC37 | -0.481 | 0.000113065 |
| Nsp8 | TUBA3E | -0.481 | 0.00011541 |

| | | | |
|---|---|---|---|
| Nsp9 | JUNB | -0.481 | 0.000113677 |
| Nsp10 | TRAF1 | -0.481 | 0.000116291 |
| Nsp10 | WNT6 | -0.481 | 0.000113359 |
| Nsp10 | GADD45B | -0.481 | 0.000113311 |
| Nsp12 | LMNA | -0.481 | 0.000115993 |
| Nsp12 | IFNL2 | -0.481 | 0.000113572 |
| Nsp12 | ITGB3 | -0.481 | 0.000115908 |
| Nsp13 | MAP3K11 | -0.481 | 0.000113145 |
| Nsp13 | EIF4E1B | -0.481 | 0.000115505 |
| Nsp13 | LCK | -0.481 | 0.000114997 |
| Nsp13 | ADCY6 | -0.481 | 0.000114714 |
| Nsp13 | BBC3 | -0.481 | 0.000114562 |
| Nsp14 | HSPA6 | -0.481 | 0.000115335 |
| Nsp14 | PFKP | -0.481 | 0.000113023 |
| Nsp14 | NPRL3 | -0.481 | 0.000114364 |
| Nsp15 | STAT5A | -0.481 | 0.000116348 |
| Nsp16 | FGF6 | -0.481 | 0.000114801 |
| S | IL27 | -0.481 | 0.000116028 |
| S | CACNA2D2 | -0.481 | 0.000116654 |
| Orf3a | CEBPB | -0.481 | 0.000113301 |
| Orf3a | DDIT4 | -0.481 | 0.00011494 |
| Orf3a | DUSP8 | -0.481 | 0.000113357 |
| Orf3a | MAPK8IP3 | -0.481 | 0.000113626 |
| Orf8 | GSK3A | -0.481 | 0.000116339 |
| Orf8 | MAP4K2 | -0.481 | 0.000115152 |
| Orf8 | TSC2 | -0.481 | 0.000113199 |
| Nsp2 | FOSB | -0.482 | 0.000110802 |
| Nsp2 | GRK1 | -0.482 | 0.000109723 |
| Nsp2 | PIK3CD | -0.482 | 0.000110204 |
| Nsp2 | CRK | -0.482 | 0.000111134 |
| Nsp3 | CNTFR | -0.482 | 0.000109793 |
| Nsp4 | STAT5A | -0.482 | 0.000110142 |
| Nsp4 | CSH1 | -0.482 | 0.000111437 |
| Nsp4 | ITGA2B | -0.482 | 0.000109023 |
| Nsp5 | FGF23 | -0.482 | 0.000112394 |
| Nsp5 | GDF5 | -0.482 | 0.000112955 |
| Nsp5 | CPT1B | -0.482 | 0.00011267 |
| Nsp6 | IRS2 | -0.482 | 0.000108918 |
| Nsp8 | IL34 | -0.482 | 0.000110128 |
| Nsp10 | LMNB2 | -0.482 | 0.000109926 |
| Nsp10 | BMP8A | -0.482 | 0.000112482 |
| Nsp10 | RARA | -0.482 | 0.000112661 |
| Nsp12 | FGFR1 | -0.482 | 0.000112147 |
| Nsp12 | ERBB2 | -0.482 | 0.000110622 |
| Nsp13 | TRIP10 | -0.482 | 0.000110789 |
| Nsp13 | VAV1 | -0.482 | 0.000110293 |
| Nsp13 | MRAS | -0.482 | 0.000112639 |
| Nsp14 | RAC2 | -0.482 | 0.000110983 |
| Nsp15 | MAPK12 | -0.482 | 0.000110322 |
| Nsp16 | LPAR2 | -0.482 | 0.000110961 |

| | | | |
|---|---|---|---|
| S | OSM | -0.482 | 0.000109981 |
| Orf3a | PRKAR1B | -0.482 | 0.000112111 |
| Orf3a | DUSP9 | -0.482 | 0.000110619 |
| Orf8 | ID4 | -0.482 | 0.000112178 |
| Orf8 | FGF23 | -0.482 | 0.000110638 |
| Nsp2 | DUSP2 | -0.483 | 0.000108412 |
| Nsp2 | WNT7B | -0.483 | 0.00010594 |
| Nsp2 | WNT9B | -0.483 | 0.000106834 |
| Nsp2 | PIM1 | -0.483 | 0.000106353 |
| Nsp3 | RAC3 | -0.483 | 0.000107246 |
| Nsp3 | TIMP1 | -0.483 | 0.000104937 |
| Nsp3 | GFAP | -0.483 | 0.000104964 |
| Nsp3 | ADCY6 | -0.483 | 0.000104859 |
| Nsp4 | NTRK1 | -0.483 | 0.00010611 |
| Nsp4 | PLA2G4B | -0.483 | 0.000108088 |
| Nsp5 | PLCG1 | -0.483 | 0.000106261 |
| Nsp6 | TNFRSF18 | -0.483 | 0.000104859 |
| Nsp6 | RXRA | -0.483 | 0.000106795 |
| Nsp6 | FGF4 | -0.483 | 0.000107459 |
| Nsp6 | TGFB1 | -0.483 | 0.000105818 |
| Nsp8 | PRKAB1 | -0.483 | 0.000107341 |
| Nsp8 | PKN3 | -0.483 | 0.000108719 |
| Nsp10 | TNFRSF4 | -0.483 | 0.000105952 |
| Nsp12 | IL11 | -0.483 | 0.000106412 |
| Nsp12 | TRIP10 | -0.483 | 0.000106485 |
| Nsp12 | PFKL | -0.483 | 0.000106054 |
| Nsp12 | CACNA2D2 | -0.483 | 0.000106291 |
| Nsp12 | ENO3 | -0.483 | 0.00010504 |
| Nsp12 | FOXP3 | -0.483 | 0.000106816 |
| Nsp13 | CHRD | -0.483 | 0.000105038 |
| Nsp14 | NGFR | -0.483 | 0.000106554 |
| Nsp14 | ITGA2B | -0.483 | 0.000106724 |
| Nsp14 | XCR1 | -0.483 | 0.000106596 |
| Nsp15 | PRKCD | -0.483 | 0.000108004 |
| Nsp16 | RASGRP4 | -0.483 | 0.000108331 |
| Nsp16 | MCL1 | -0.483 | 0.000108758 |
| Nsp16 | OSM | -0.483 | 0.000106855 |
| S | ADCY6 | -0.483 | 0.0001079 |
| Orf3a | CDKN2B | -0.483 | 0.000108604 |
| Orf8 | NFATC2 | -0.483 | 0.000107971 |
| Orf8 | PRKCZ | -0.483 | 0.000108597 |
| Orf8 | TUBA3E | -0.483 | 0.000107689 |
| Orf8 | ITGA2B | -0.483 | 0.000107235 |
| Orf8 | FGF22 | -0.483 | 0.000106603 |
| Nsp2 | PRR5 | -0.484 | 0.000102259 |
| Nsp4 | CPT1B | -0.484 | 0.000103849 |
| Nsp4 | BBC3 | -0.484 | 0.000102082 |
| Nsp5 | CCR7 | -0.484 | 0.000104081 |
| Nsp5 | MKNK2 | -0.484 | 0.000101708 |
| Nsp5 | PRKACG | -0.484 | 0.000101673 |

| | | | |
|---|---|---|---|
| Nsp6 | CACNG8 | -0.484 | 0.000102869 |
| Nsp8 | PIDD1 | -0.484 | 0.000103031 |
| Nsp8 | HSPA6 | -0.484 | 0.000102595 |
| Nsp8 | TYK2 | -0.484 | 0.000103832 |
| Nsp8 | CACNA2D2 | -0.484 | 0.000103662 |
| Nsp9 | HRK | -0.484 | 0.000104727 |
| Nsp9 | CAPN1 | -0.484 | 0.000101223 |
| Nsp10 | IL27 | -0.484 | 0.000104692 |
| Nsp12 | CHAD | -0.484 | 0.000101506 |
| Nsp12 | RORC | -0.484 | 0.000104689 |
| Nsp12 | TNFRSF1A | -0.484 | 0.000101179 |
| Nsp12 | VWF | -0.484 | 0.000104459 |
| Nsp13 | GDF2 | -0.484 | 0.00010171 |
| Nsp14 | FOSB | -0.484 | 0.000101546 |
| Nsp14 | RAC3 | -0.484 | 0.000104473 |
| Nsp14 | LMNA | -0.484 | 0.000104649 |
| Nsp15 | FLT3LG | -0.484 | 0.00010368 |
| Nsp15 | HSPA6 | -0.484 | 0.00010352 |
| Nsp16 | TNF | -0.484 | 0.00010459 |
| Nsp16 | CISH | -0.484 | 0.000101659 |
| Nsp16 | ITGA7 | -0.484 | 0.00010443 |
| S | MAP4K2 | -0.484 | 0.000102237 |
| S | CDC37 | -0.484 | 0.000101811 |
| S | IGF2 | -0.484 | 0.000104201 |
| Orf3a | FZD10 | -0.484 | 0.000103121 |
| Orf8 | RAC3 | -0.484 | 0.000104024 |
| Orf8 | GNB2 | -0.484 | 0.000103611 |
| Orf8 | PFKP | -0.484 | 0.00010442 |
| Orf8 | WNT10A | -0.484 | 0.000103039 |
| Orf8 | CALML5 | -0.484 | 0.000104651 |
| Orf8 | PPP1R3E | -0.484 | 0.00010165 |
| Nsp2 | CAPN2 | -0.485 | 9.91E-05 |
| Nsp2 | PIK3R2 | -0.485 | 9.92E-05 |
| Nsp2 | CCND1 | -0.485 | 9.84E-05 |
| Nsp4 | IKBKE | -0.485 | 9.76E-05 |
| Nsp4 | DHX58 | -0.485 | 9.76E-05 |
| Nsp4 | CXCL14 | -0.485 | 9.81E-05 |
| Nsp5 | ATP6V1B1 | -0.485 | 0.000100439 |
| Nsp5 | PLCG2 | -0.485 | 9.81E-05 |
| Nsp5 | EXOC7 | -0.485 | 9.77E-05 |
| Nsp7 | DUSP6 | -0.485 | 9.80E-05 |
| Nsp8 | EIF4EBP1 | -0.485 | 9.98E-05 |
| Nsp8 | ID3 | -0.485 | 9.90E-05 |
| Nsp9 | WNT7B | -0.485 | 9.97E-05 |
| Nsp9 | MAP2K7 | -0.485 | 9.95E-05 |
| Nsp10 | IL17D | -0.485 | 0.000100657 |
| Nsp10 | FZD1 | -0.485 | 0.000100372 |
| Nsp13 | RNF152 | -0.485 | 9.92E-05 |
| Nsp13 | HRAS | -0.485 | 0.000100336 |
| Nsp13 | ACVRL1 | -0.485 | 0.000100444 |

| | | | |
|---|---|---|---|
| Nsp14 | GSK3A | -0.485 | 0.000100851 |
| Nsp14 | VEGFA | -0.485 | 9.90E-05 |
| Nsp15 | PPP2R1A | -0.485 | 0.00010025 |
| Nsp15 | MAP4K2 | -0.485 | 9.73E-05 |
| Nsp15 | CLCF1 | -0.485 | 9.98E-05 |
| Nsp15 | ID2 | -0.485 | 0.000100096 |
| Nsp16 | TNFSF9 | -0.485 | 0.000100799 |
| Nsp16 | CACNA2D4 | -0.485 | 0.000100284 |
| Orf3a | EFNA3 | -0.485 | 9.91E-05 |
| Orf3a | BMP8B | -0.485 | 9.81E-05 |
| Orf8 | GADD45A | -0.485 | 9.86E-05 |
| Orf8 | GRK5 | -0.485 | 0.00010054 |
| Orf8 | XCR1 | -0.485 | 9.96E-05 |
| Nsp2 | ADCY9 | -0.486 | 9.53E-05 |
| Nsp2 | PPP2R2C | -0.486 | 9.49E-05 |
| Nsp2 | ADCY7 | -0.486 | 9.45E-05 |
| Nsp2 | FGF8 | -0.486 | 9.54E-05 |
| Nsp3 | SREBF1 | -0.486 | 9.38E-05 |
| Nsp4 | MAPK15 | -0.486 | 9.49E-05 |
| Nsp4 | HSPA6 | -0.486 | 9.44E-05 |
| Nsp4 | TNFRSF1B | -0.486 | 9.54E-05 |
| Nsp7 | AMH | -0.486 | 9.62E-05 |
| Nsp8 | INSR | -0.486 | 9.39E-05 |
| Nsp8 | TSC2 | -0.486 | 9.42E-05 |
| Nsp8 | TNFRSF1B | -0.486 | 9.59E-05 |
| Nsp10 | LEFTY1 | -0.486 | 9.39E-05 |
| Nsp12 | NTF3 | -0.486 | 9.65E-05 |
| Nsp12 | CD14 | -0.486 | 9.54E-05 |
| Nsp12 | CRK | -0.486 | 9.68E-05 |
| Nsp13 | IRAK1 | -0.486 | 9.42E-05 |
| Nsp13 | IRF3 | -0.486 | 9.53E-05 |
| Nsp13 | ITGA2B | -0.486 | 9.66E-05 |
| Nsp14 | TLR9 | -0.486 | 9.42E-05 |
| Nsp14 | CARD10 | -0.486 | 9.46E-05 |
| Nsp14 | PDGFB | -0.486 | 9.67E-05 |
| Nsp15 | PLA2G4B | -0.486 | 9.44E-05 |
| Nsp15 | IL27RA | -0.486 | 9.59E-05 |
| Nsp15 | CSH1 | -0.486 | 9.58E-05 |
| Nsp16 | PARP3 | -0.486 | 9.68E-05 |
| Nsp16 | MAPK15 | -0.486 | 9.53E-05 |
| S | CACNA1C | -0.486 | 9.45E-05 |
| S | GSK3A | -0.486 | 9.62E-05 |
| S | TNFSF12 | -0.486 | 9.69E-05 |
| S | WNT10B | -0.486 | 9.38E-05 |
| Orf3a | TRADD | -0.486 | 9.49E-05 |
| Orf8 | IL10 | -0.486 | 9.45E-05 |
| Orf8 | PPP3R2 | -0.486 | 9.62E-05 |
| Orf8 | INSR | -0.486 | 9.58E-05 |
| Orf8 | CCND1 | -0.486 | 9.38E-05 |
| Orf8 | CSH2 | -0.486 | 9.36E-05 |

| | | | |
|---|---|---|---|
| Nsp2 | PITX2 | -0.487 | 9.18E-05 |
| Nsp2 | ZAP70 | -0.487 | 9.32E-05 |
| Nsp3 | ACVRL1 | -0.487 | 9.03E-05 |
| Nsp4 | PRKCZ | -0.487 | 9.11E-05 |
| Nsp5 | PLCB2 | -0.487 | 9.02E-05 |
| Nsp5 | TRAF2 | -0.487 | 9.16E-05 |
| Nsp5 | ID1 | -0.487 | 9.15E-05 |
| Nsp5 | GNB2 | -0.487 | 9.03E-05 |
| Nsp5 | TYK2 | -0.487 | 9.34E-05 |
| Nsp5 | SREBF1 | -0.487 | 9.16E-05 |
| Nsp7 | CDKN2B | -0.487 | 9.09E-05 |
| Nsp8 | FLT3LG | -0.487 | 9.12E-05 |
| Nsp9 | GDF1 | -0.487 | 9.16E-05 |
| Nsp9 | PIM1 | -0.487 | 9.31E-05 |
| Nsp9 | FGF8 | -0.487 | 9.34E-05 |
| Nsp10 | SMAD6 | -0.487 | 9.09E-05 |
| Nsp10 | LPAR5 | -0.487 | 9.23E-05 |
| Nsp10 | TELO2 | -0.487 | 9.19E-05 |
| Nsp10 | GDF11 | -0.487 | 9.31E-05 |
| Nsp12 | STAT5A | -0.487 | 9.16E-05 |
| Nsp12 | IL27RA | -0.487 | 9.17E-05 |
| Nsp13 | CSF3 | -0.487 | 9.19E-05 |
| Nsp14 | EPHA2 | -0.487 | 9.03E-05 |
| Nsp14 | PPP5C | -0.487 | 9.17E-05 |
| Nsp14 | IL17C | -0.487 | 9.04E-05 |
| Nsp14 | PLCG1 | -0.487 | 9.03E-05 |
| Nsp15 | FGF18 | -0.487 | 9.03E-05 |
| Nsp15 | TNF | -0.487 | 9.16E-05 |
| Nsp15 | TNFRSF1B | -0.487 | 9.08E-05 |
| Nsp15 | FOSL1 | -0.487 | 9.23E-05 |
| Nsp16 | IGF1R | -0.487 | 9.28E-05 |
| Nsp16 | CACNA1E | -0.487 | 9.00E-05 |
| S | CCR7 | -0.487 | 9.33E-05 |
| S | PKN3 | -0.487 | 9.35E-05 |
| Orf3a | STK11 | -0.487 | 9.13E-05 |
| Orf8 | WNT4 | -0.487 | 9.22E-05 |
| Orf8 | PPP1R3F | -0.487 | 9.12E-05 |
| Nsp2 | GRK6 | -0.488 | 8.81E-05 |
| Nsp2 | IKBKE | -0.488 | 8.86E-05 |
| Nsp3 | CDC37 | -0.488 | 8.66E-05 |
| Nsp3 | SESN2 | -0.488 | 8.80E-05 |
| Nsp4 | TNFSF14 | -0.488 | 8.86E-05 |
| Nsp4 | WNT5A | -0.488 | 8.75E-05 |
| Nsp5 | GREM1 | -0.488 | 8.84E-05 |
| Nsp5 | TNFRSF13C | -0.488 | 8.90E-05 |
| Nsp5 | LPAR2 | -0.488 | 8.99E-05 |
| Nsp6 | CEBPB | -0.488 | 8.79E-05 |
| Nsp7 | DVL1 | -0.488 | 8.76E-05 |
| Nsp7 | MAPK8IP3 | -0.488 | 8.90E-05 |
| Nsp8 | NR4A1 | -0.488 | 8.90E-05 |

| | | | |
|---|---|---|---|
| Nsp8 | CSH1 | -0.488 | 8.81E-05 |
| Nsp8 | IRF3 | -0.488 | 8.80E-05 |
| Nsp8 | TCL1A | -0.488 | 8.84E-05 |
| Nsp10 | CCL14 | -0.488 | 8.90E-05 |
| Nsp10 | HAMP | -0.488 | 8.76E-05 |
| Nsp12 | PLA2G4B | -0.488 | 8.96E-05 |
| Nsp12 | JAK3 | -0.488 | 8.94E-05 |
| Nsp13 | TUBA3D | -0.488 | 8.94E-05 |
| Nsp14 | ITGA3 | -0.488 | 8.92E-05 |
| Nsp14 | LAMC3 | -0.488 | 8.82E-05 |
| Nsp15 | RNF152 | -0.488 | 8.67E-05 |
| Nsp16 | NOS2 | -0.488 | 8.94E-05 |
| Nsp16 | RPS6KA1 | -0.488 | 8.90E-05 |
| S | STAT5A | -0.488 | 8.74E-05 |
| S | NFATC2 | -0.488 | 8.80E-05 |
| S | LTB | -0.488 | 8.82E-05 |
| Orf3a | HSPB1 | -0.488 | 8.89E-05 |
| Orf6 | COL6A2 | -0.488 | 8.85E-05 |
| Orf8 | IRF7 | -0.488 | 8.76E-05 |
| Orf8 | COL6A2 | -0.488 | 8.77E-05 |
| Nsp2 | RGMA | -0.489 | 8.35E-05 |
| Nsp2 | SRC | -0.489 | 8.65E-05 |
| Nsp2 | CDC25B | -0.489 | 8.63E-05 |
| Nsp2 | MAP3K6 | -0.489 | 8.37E-05 |
| Nsp2 | FZD2 | -0.489 | 8.60E-05 |
| Nsp2 | CACNA1B | -0.489 | 8.36E-05 |
| Nsp2 | COL6A2 | -0.489 | 8.43E-05 |
| Nsp2 | FGF22 | -0.489 | 8.66E-05 |
| Nsp3 | NOS2 | -0.489 | 8.66E-05 |
| Nsp3 | IKBKE | -0.489 | 8.43E-05 |
| Nsp4 | ACVRL1 | -0.489 | 8.58E-05 |
| Nsp5 | ACTG1 | -0.489 | 8.65E-05 |
| Nsp5 | CARD10 | -0.489 | 8.37E-05 |
| Nsp5 | SESN2 | -0.489 | 8.59E-05 |
| Nsp5 | TRIM25 | -0.489 | 8.47E-05 |
| Nsp6 | FZD1 | -0.489 | 8.61E-05 |
| Nsp7 | TGFB1 | -0.489 | 8.45E-05 |
| Nsp8 | GNG13 | -0.489 | 8.55E-05 |
| Nsp8 | NODAL | -0.489 | 8.36E-05 |
| Nsp8 | INS | -0.489 | 8.49E-05 |
| Nsp9 | FZD8 | -0.489 | 8.64E-05 |
| Nsp9 | FZD5 | -0.489 | 8.37E-05 |
| Nsp12 | IFNL3 | -0.489 | 8.49E-05 |
| Nsp12 | ID1 | -0.489 | 8.37E-05 |
| Nsp12 | OSM | -0.489 | 8.43E-05 |
| Nsp12 | ID2 | -0.489 | 8.41E-05 |
| Nsp13 | FGF6 | -0.489 | 8.42E-05 |
| Nsp13 | WNT10B | -0.489 | 8.65E-05 |
| Nsp13 | CTSZ | -0.489 | 8.43E-05 |
| Nsp13 | ITGA11 | -0.489 | 8.57E-05 |

| | | | |
|---|---|---|---|
| Nsp13 | TCL1A | -0.489 | 8.50E-05 |
| Nsp13 | TNFRSF1A | -0.489 | 8.50E-05 |
| Nsp13 | FADD | -0.489 | 8.47E-05 |
| Nsp13 | VWF | -0.489 | 8.50E-05 |
| Nsp14 | TNFRSF1B | -0.489 | 8.44E-05 |
| Nsp15 | PLCG2 | -0.489 | 8.42E-05 |
| Nsp15 | ECSIT | -0.489 | 8.52E-05 |
| Nsp15 | RAC2 | -0.489 | 8.59E-05 |
| Nsp16 | MAP2K3 | -0.489 | 8.40E-05 |
| Nsp16 | CCND2 | -0.489 | 8.63E-05 |
| Nsp16 | TNFRSF13C | -0.489 | 8.39E-05 |
| S | VHL | -0.489 | 8.40E-05 |
| Orf3a | CACNG8 | -0.489 | 8.54E-05 |
| Orf6 | BMP7 | -0.489 | 8.50E-05 |
| Orf8 | PRKAB1 | -0.489 | 8.42E-05 |
| Nsp2 | WNT10A | -0.49 | 8.31E-05 |
| Nsp3 | IRAK1 | -0.49 | 8.32E-05 |
| Nsp3 | MAPK15 | -0.49 | 8.19E-05 |
| Nsp3 | MAPK12 | -0.49 | 8.14E-05 |
| Nsp3 | INHBA | -0.49 | 8.30E-05 |
| Nsp3 | RELA | -0.49 | 8.10E-05 |
| Nsp3 | PKN3 | -0.49 | 8.19E-05 |
| Nsp3 | PLA2G4D | -0.49 | 8.03E-05 |
| Nsp4 | ARAF | -0.49 | 8.29E-05 |
| Nsp4 | FLNC | -0.49 | 8.15E-05 |
| Nsp5 | WNT9B | -0.49 | 8.20E-05 |
| Nsp5 | CACNA2D2 | -0.49 | 8.21E-05 |
| Nsp8 | VAV2 | -0.49 | 8.10E-05 |
| Nsp13 | THSD4 | -0.49 | 8.29E-05 |
| Nsp13 | FGF2 | -0.49 | 8.13E-05 |
| Nsp14 | PLA2G4B | -0.49 | 8.32E-05 |
| Nsp14 | PTK2B | -0.49 | 8.30E-05 |
| Nsp14 | GNA12 | -0.49 | 8.11E-05 |
| Nsp14 | RASGRF1 | -0.49 | 8.09E-05 |
| Nsp15 | IFNL1 | -0.49 | 8.14E-05 |
| Nsp15 | ITGA3 | -0.49 | 8.21E-05 |
| Nsp15 | RORC | -0.49 | 8.18E-05 |
| Nsp15 | INHBA | -0.49 | 8.15E-05 |
| Nsp15 | PHLPP1 | -0.49 | 8.02E-05 |
| Nsp15 | PRKACG | -0.49 | 8.12E-05 |
| Nsp16 | IL27RA | -0.49 | 8.23E-05 |
| Nsp16 | TIMP1 | -0.49 | 8.17E-05 |
| Nsp16 | BAD | -0.49 | 8.06E-05 |
| Nsp16 | NFKBIA | -0.49 | 8.20E-05 |
| S | SMAD3 | -0.49 | 8.06E-05 |
| Orf3a | AMH | -0.49 | 8.21E-05 |
| Orf8 | WNT7A | -0.49 | 8.28E-05 |
| Nsp2 | FGF18 | -0.491 | 7.77E-05 |
| Nsp3 | WNT10B | -0.491 | 7.97E-05 |
| Nsp3 | PFKP | -0.491 | 7.97E-05 |

| | | | |
|---|---|---|---|
| Nsp3 | ITGA2B | -0.491 | 8.02E-05 |
| Nsp3 | IL21R | -0.491 | 7.72E-05 |
| Nsp3 | GNB3 | -0.491 | 7.93E-05 |
| Nsp4 | AKT1 | -0.491 | 7.99E-05 |
| Nsp4 | JMJD7-PLA2G | -0.491 | 8.00E-05 |
| Nsp4 | PIK3R5 | -0.491 | 7.90E-05 |
| Nsp5 | PIN1 | -0.491 | 7.72E-05 |
| Nsp8 | THSD4 | -0.491 | 7.87E-05 |
| Nsp8 | CCR7 | -0.491 | 7.93E-05 |
| Nsp9 | MCL1 | -0.491 | 7.80E-05 |
| Nsp9 | GADD45B | -0.491 | 8.00E-05 |
| Nsp12 | ITGA11 | -0.491 | 7.85E-05 |
| Nsp12 | PPP1R3F | -0.491 | 7.89E-05 |
| Nsp12 | PDGFRB | -0.491 | 7.83E-05 |
| Nsp13 | WNT9A | -0.491 | 7.90E-05 |
| Nsp14 | JMJD7-PLA2G | -0.491 | 7.98E-05 |
| Nsp14 | PPP3R2 | -0.491 | 7.94E-05 |
| Nsp15 | CHRD | -0.491 | 7.85E-05 |
| Nsp15 | PPP1R3F | -0.491 | 7.86E-05 |
| Nsp15 | FOXP3 | -0.491 | 7.83E-05 |
| Nsp16 | GADD45A | -0.491 | 7.78E-05 |
| Nsp16 | EIF4EBP1 | -0.491 | 7.87E-05 |
| S | SREBF1 | -0.491 | 7.98E-05 |
| S | LPAR2 | -0.491 | 7.84E-05 |
| Orf8 | RPTOR | -0.491 | 7.78E-05 |
| Orf8 | EXOC7 | -0.491 | 7.96E-05 |
| Orf8 | RPS6KA2 | -0.491 | 7.83E-05 |
| Nsp2 | CACNA1H | -0.492 | 7.69E-05 |
| Nsp2 | HSPB1 | -0.492 | 7.71E-05 |
| Nsp2 | DUSP5 | -0.492 | 7.52E-05 |
| Nsp2 | TNFRSF6B | -0.492 | 7.48E-05 |
| Nsp3 | PLA2G4B | -0.492 | 7.47E-05 |
| Nsp3 | STAT5A | -0.492 | 7.69E-05 |
| Nsp3 | ITGA11 | -0.492 | 7.43E-05 |
| Nsp3 | GH1 | -0.492 | 7.45E-05 |
| Nsp3 | CACNA2D4 | -0.492 | 7.65E-05 |
| Nsp3 | PRKACA | -0.492 | 7.52E-05 |
| Nsp3 | IGF2 | -0.492 | 7.48E-05 |
| Nsp4 | CSF2RB | -0.492 | 7.52E-05 |
| Nsp4 | DUSP2 | -0.492 | 7.44E-05 |
| Nsp4 | PLA2G4D | -0.492 | 7.55E-05 |
| Nsp5 | TSC2 | -0.492 | 7.45E-05 |
| Nsp8 | PPP2R1A | -0.492 | 7.49E-05 |
| Nsp8 | RPS6KB2 | -0.492 | 7.49E-05 |
| Nsp8 | TNFRSF12A | -0.492 | 7.67E-05 |
| Nsp8 | GNB3 | -0.492 | 7.63E-05 |
| Nsp9 | FZD1 | -0.492 | 7.52E-05 |
| Nsp10 | RELB | -0.492 | 7.68E-05 |
| Nsp12 | FOXO1 | -0.492 | 7.68E-05 |
| Nsp12 | GNG13 | -0.492 | 7.69E-05 |

| | | | |
|---|---|---|---|
| Nsp12 | ACVR2B | -0.492 | 7.68E-05 |
| Nsp12 | FLNC | -0.492 | 7.57E-05 |
| Nsp13 | NOS2 | -0.492 | 7.59E-05 |
| Nsp13 | PIK3R6 | -0.492 | 7.50E-05 |
| Nsp14 | EPOR | -0.492 | 7.53E-05 |
| Nsp14 | PPP2R1A | -0.492 | 7.48E-05 |
| Nsp14 | FMOD | -0.492 | 7.62E-05 |
| Nsp14 | TRIM25 | -0.492 | 7.45E-05 |
| Nsp15 | JMJD7-PLA2G | -0.492 | 7.58E-05 |
| Nsp15 | EXOC7 | -0.492 | 7.58E-05 |
| Nsp16 | EDA | -0.492 | 7.67E-05 |
| Nsp16 | GDF5 | -0.492 | 7.48E-05 |
| S | PIK3R5 | -0.492 | 7.56E-05 |
| Orf3a | LRP5 | -0.492 | 7.58E-05 |
| Orf3a | IRS2 | -0.492 | 7.70E-05 |
| Orf8 | THSD4 | -0.492 | 7.53E-05 |
| Orf8 | ANGPT4 | -0.492 | 7.61E-05 |
| Orf10 | IL36RN | -0.492 | 7.52E-05 |
| Nsp2 | NOS3 | -0.493 | 7.19E-05 |
| Nsp2 | RPTOR | -0.493 | 7.23E-05 |
| Nsp2 | MAPK4 | -0.493 | 7.18E-05 |
| Nsp3 | CACNA1S | -0.493 | 7.18E-05 |
| Nsp3 | CXCL2 | -0.493 | 7.20E-05 |
| Nsp4 | RAC3 | -0.493 | 7.24E-05 |
| Nsp4 | ID1 | -0.493 | 7.24E-05 |
| Nsp4 | NODAL | -0.493 | 7.41E-05 |
| Nsp4 | LCK | -0.493 | 7.21E-05 |
| Nsp4 | CACNA2D2 | -0.493 | 7.33E-05 |
| Nsp5 | THSD4 | -0.493 | 7.30E-05 |
| Nsp5 | DHX58 | -0.493 | 7.25E-05 |
| Nsp5 | ID3 | -0.493 | 7.38E-05 |
| Nsp7 | ELOB | -0.493 | 7.25E-05 |
| Nsp8 | LTB | -0.493 | 7.31E-05 |
| Nsp9 | SHC2 | -0.493 | 7.25E-05 |
| Nsp9 | IRS2 | -0.493 | 7.24E-05 |
| Nsp10 | JUN | -0.493 | 7.17E-05 |
| Nsp10 | CAPN1 | -0.493 | 7.36E-05 |
| Nsp12 | MAPK12 | -0.493 | 7.20E-05 |
| Nsp12 | PPP5C | -0.493 | 7.31E-05 |
| Nsp12 | INS | -0.493 | 7.26E-05 |
| Nsp13 | IL11 | -0.493 | 7.26E-05 |
| Nsp14 | ARAF | -0.493 | 7.40E-05 |
| Nsp15 | ANGPT4 | -0.493 | 7.33E-05 |
| Nsp15 | ITGA11 | -0.493 | 7.18E-05 |
| Nsp15 | FGF17 | -0.493 | 7.28E-05 |
| Nsp15 | PTPN5 | -0.493 | 7.18E-05 |
| Nsp15 | LCK | -0.493 | 7.34E-05 |
| Nsp16 | BMP4 | -0.493 | 7.39E-05 |
| Nsp16 | ITGA11 | -0.493 | 7.20E-05 |
| Nsp16 | CSH2 | -0.493 | 7.31E-05 |

| | | | |
|---|---|---|---|
| S | TNFRSF1B | -0.493 | 7.28E-05 |
| Orf8 | CACNG6 | -0.493 | 7.21E-05 |
| Orf8 | ID1 | -0.493 | 7.18E-05 |
| Orf8 | HSPA1B | -0.493 | 7.16E-05 |
| Orf8 | EGLN1 | -0.493 | 7.20E-05 |
| Nsp2 | DAB2IP | -0.494 | 7.04E-05 |
| Nsp2 | CALML5 | -0.494 | 6.93E-05 |
| Nsp3 | ENO3 | -0.494 | 7.01E-05 |
| Nsp3 | FOXP3 | -0.494 | 7.05E-05 |
| Nsp4 | BCAR1 | -0.494 | 6.88E-05 |
| Nsp4 | PFKL | -0.494 | 7.10E-05 |
| Nsp5 | RASGRF1 | -0.494 | 6.95E-05 |
| Nsp8 | MAPK12 | -0.494 | 7.01E-05 |
| Nsp8 | ITGA2B | -0.494 | 6.97E-05 |
| Nsp8 | THBS2 | -0.494 | 7.06E-05 |
| Nsp9 | CTF1 | -0.494 | 6.97E-05 |
| Nsp10 | PRR5 | -0.494 | 7.01E-05 |
| Nsp12 | AKT1 | -0.494 | 6.95E-05 |
| Nsp12 | THSD4 | -0.494 | 7.04E-05 |
| Nsp12 | ITGA3 | -0.494 | 7.09E-05 |
| Nsp12 | GRK6 | -0.494 | 7.12E-05 |
| Nsp12 | IKBKE | -0.494 | 7.03E-05 |
| Nsp12 | XCR1 | -0.494 | 7.08E-05 |
| Nsp12 | WNT5A | -0.494 | 6.98E-05 |
| Nsp13 | EDAR | -0.494 | 6.97E-05 |
| Nsp13 | TIMP1 | -0.494 | 7.13E-05 |
| Nsp14 | TNFSF9 | -0.494 | 7.03E-05 |
| Nsp15 | ACVRL1 | -0.494 | 7.00E-05 |
| Nsp16 | NFATC2 | -0.494 | 7.13E-05 |
| Nsp16 | ACTG1 | -0.494 | 6.98E-05 |
| Nsp16 | PRKCD | -0.494 | 7.01E-05 |
| Nsp16 | WNT10B | -0.494 | 7.12E-05 |
| Nsp16 | HK3 | -0.494 | 7.06E-05 |
| S | ACTG1 | -0.494 | 6.88E-05 |
| S | PDGFRB | -0.494 | 7.11E-05 |
| Orf6 | RPTOR | -0.494 | 7.07E-05 |
| Orf8 | CXCR5 | -0.494 | 7.07E-05 |
| Orf8 | HMOX1 | -0.494 | 6.99E-05 |
| Orf8 | SLC7A5 | -0.494 | 7.03E-05 |
| Orf8 | CTF1 | -0.494 | 7.10E-05 |
| Nsp3 | IRF3 | -0.495 | 6.77E-05 |
| Nsp3 | PDGFRB | -0.495 | 6.61E-05 |
| Nsp4 | TCL1A | -0.495 | 6.80E-05 |
| Nsp4 | LTB | -0.495 | 6.70E-05 |
| Nsp4 | FADD | -0.495 | 6.67E-05 |
| Nsp5 | FMOD | -0.495 | 6.62E-05 |
| Nsp5 | IL17C | -0.495 | 6.62E-05 |
| Nsp5 | FADD | -0.495 | 6.68E-05 |
| Nsp8 | CACNB1 | -0.495 | 6.62E-05 |
| Nsp8 | PFKP | -0.495 | 6.84E-05 |

| | | | |
|---|---|---|---|
| Nsp9 | WNT6 | -0.495 | 6.80E-05 |
| Nsp10 | MYC | -0.495 | 6.61E-05 |
| Nsp12 | NOS2 | -0.495 | 6.82E-05 |
| Nsp12 | IRAK1 | -0.495 | 6.71E-05 |
| Nsp12 | TNF | -0.495 | 6.80E-05 |
| Nsp13 | MAP4K2 | -0.495 | 6.81E-05 |
| Nsp13 | JAK3 | -0.495 | 6.63E-05 |
| Nsp14 | TYK2 | -0.495 | 6.69E-05 |
| Nsp15 | PRKCZ | -0.495 | 6.72E-05 |
| Nsp15 | IRAK1 | -0.495 | 6.61E-05 |
| Nsp16 | GNG13 | -0.495 | 6.81E-05 |
| Nsp16 | MAPK12 | -0.495 | 6.84E-05 |
| S | RAC3 | -0.495 | 6.85E-05 |
| S | INSR | -0.495 | 6.72E-05 |
| S | CACNG5 | -0.495 | 6.86E-05 |
| Nsp2 | MMP9 | -0.496 | 6.40E-05 |
| Nsp2 | GDF11 | -0.496 | 6.58E-05 |
| Nsp2 | JUND | -0.496 | 6.60E-05 |
| Nsp2 | FGFR3 | -0.496 | 6.55E-05 |
| Nsp4 | IL27RA | -0.496 | 6.59E-05 |
| Nsp4 | MAP3K11 | -0.496 | 6.49E-05 |
| Nsp5 | TNFSF14 | -0.496 | 6.57E-05 |
| Nsp5 | HCK | -0.496 | 6.39E-05 |
| Nsp5 | NGFR | -0.496 | 6.49E-05 |
| Nsp5 | RAC2 | -0.496 | 6.42E-05 |
| Nsp7 | ENDOG | -0.496 | 6.57E-05 |
| Nsp8 | CAMKK2 | -0.496 | 6.43E-05 |
| Nsp8 | TNFSF14 | -0.496 | 6.57E-05 |
| Nsp8 | ICAM1 | -0.496 | 6.46E-05 |
| Nsp8 | FOXO3 | -0.496 | 6.51E-05 |
| Nsp10 | VEGFA | -0.496 | 6.55E-05 |
| Nsp12 | NTRK1 | -0.496 | 6.48E-05 |
| Nsp12 | FGR | -0.496 | 6.43E-05 |
| Nsp12 | NGFR | -0.496 | 6.43E-05 |
| Nsp12 | TCL1A | -0.496 | 6.38E-05 |
| Nsp13 | TNF | -0.496 | 6.58E-05 |
| Nsp14 | MUC5B | -0.496 | 6.39E-05 |
| Nsp14 | TAB1 | -0.496 | 6.49E-05 |
| Nsp15 | CAMKK2 | -0.496 | 6.60E-05 |
| Nsp15 | NPRL2 | -0.496 | 6.49E-05 |
| Nsp16 | CACNA1S | -0.496 | 6.54E-05 |
| Nsp16 | IL12RB1 | -0.496 | 6.60E-05 |
| Nsp16 | INS | -0.496 | 6.44E-05 |
| S | TNFRSF25 | -0.496 | 6.36E-05 |
| S | ITGA3 | -0.496 | 6.36E-05 |
| Nsp2 | GRK2 | -0.497 | 6.26E-05 |
| Nsp2 | HSPA1B | -0.497 | 6.13E-05 |
| Nsp3 | ACTG1 | -0.497 | 6.24E-05 |
| Nsp3 | TNFRSF1B | -0.497 | 6.10E-05 |
| Nsp3 | NODAL | -0.497 | 6.29E-05 |

| | | | |
|---|---|---|---|
| Nsp4 | CHAD | -0.497 | 6.18E-05 |
| Nsp4 | CARD10 | -0.497 | 6.11E-05 |
| Nsp4 | CXCL2 | -0.497 | 6.28E-05 |
| Nsp5 | ADCY6 | -0.497 | 6.24E-05 |
| Nsp5 | HSPA1B | -0.497 | 6.13E-05 |
| Nsp8 | RAC3 | -0.497 | 6.17E-05 |
| Nsp8 | MUC5B | -0.497 | 6.33E-05 |
| Nsp8 | PLCG1 | -0.497 | 6.29E-05 |
| Nsp8 | JAK3 | -0.497 | 6.19E-05 |
| Nsp9 | HSPB1 | -0.497 | 6.34E-05 |
| Nsp10 | GNA12 | -0.497 | 6.14E-05 |
| Nsp12 | PIK3R5 | -0.497 | 6.31E-05 |
| Nsp12 | PLA2G4D | -0.497 | 6.18E-05 |
| Nsp13 | CXCL2 | -0.497 | 6.21E-05 |
| Nsp13 | FOXP3 | -0.497 | 6.20E-05 |
| Nsp14 | WNT7A | -0.497 | 6.13E-05 |
| Nsp16 | PLA2G4B | -0.497 | 6.14E-05 |
| Nsp16 | CSH1 | -0.497 | 6.22E-05 |
| Nsp16 | TNFRSF1B | -0.497 | 6.24E-05 |
| Nsp16 | GRK5 | -0.497 | 6.27E-05 |
| Nsp16 | EGLN2 | -0.497 | 6.19E-05 |
| Orf3a | MCL1 | -0.497 | 6.18E-05 |
| Orf3a | NCF1 | -0.497 | 6.22E-05 |
| Orf8 | TOLLIP | -0.497 | 6.29E-05 |
| Orf8 | CCR10 | -0.497 | 6.20E-05 |
| Orf8 | GDF11 | -0.497 | 6.13E-05 |
| Orf8 | PRKCG | -0.497 | 6.21E-05 |
| Nsp2 | HSPA2 | -0.498 | 5.88E-05 |
| Nsp2 | DUSP4 | -0.498 | 5.95E-05 |
| Nsp2 | DUSP9 | -0.498 | 6.04E-05 |
| Nsp3 | CHRD | -0.498 | 5.88E-05 |
| Nsp3 | CSF2RB | -0.498 | 6.00E-05 |
| Nsp3 | ACVR2B | -0.498 | 6.01E-05 |
| Nsp4 | ISG15 | -0.498 | 5.96E-05 |
| Nsp4 | ATP6V1B1 | -0.498 | 5.95E-05 |
| Nsp4 | PPP3R2 | -0.498 | 6.04E-05 |
| Nsp5 | CTSD | -0.498 | 5.95E-05 |
| Nsp5 | ADCY9 | -0.498 | 6.04E-05 |
| Nsp5 | GYS1 | -0.498 | 6.06E-05 |
| Nsp5 | CD14 | -0.498 | 5.94E-05 |
| Nsp5 | BCAR1 | -0.498 | 5.94E-05 |
| Nsp7 | FGF3 | -0.498 | 5.89E-05 |
| Nsp8 | IGF1R | -0.498 | 6.00E-05 |
| Nsp8 | MMP14 | -0.498 | 6.04E-05 |
| Nsp10 | TNFRSF8 | -0.498 | 6.07E-05 |
| Nsp12 | CXCR5 | -0.498 | 6.03E-05 |
| Nsp12 | EXOC7 | -0.498 | 5.96E-05 |
| Nsp12 | PFKP | -0.498 | 6.03E-05 |
| Nsp12 | LIF | -0.498 | 6.00E-05 |
| Nsp12 | PPP1R3E | -0.498 | 6.03E-05 |

| | | | |
|---|---|---|---|
| Nsp13 | CACNA2D4 | -0.498 | 6.05E-05 |
| Nsp13 | GFAP | -0.498 | 6.04E-05 |
| Nsp13 | ID3 | -0.498 | 5.88E-05 |
| Nsp13 | TYK2 | -0.498 | 5.92E-05 |
| Nsp13 | RAPGEF1 | -0.498 | 5.91E-05 |
| Nsp14 | RPS6KB2 | -0.498 | 5.88E-05 |
| Nsp15 | FGR | -0.498 | 5.89E-05 |
| Nsp15 | PFKP | -0.498 | 6.08E-05 |
| Nsp16 | ACTB | -0.498 | 6.10E-05 |
| Nsp16 | PHLPP1 | -0.498 | 6.03E-05 |
| Nsp16 | CNTFR | -0.498 | 5.87E-05 |
| Nsp16 | ITGB3 | -0.498 | 6.09E-05 |
| Nsp16 | G6PC3 | -0.498 | 5.93E-05 |
| Orf3a | WNT7B | -0.498 | 5.89E-05 |
| Orf3a | MAPK8IP2 | -0.498 | 5.97E-05 |
| Orf8 | TNFRSF25 | -0.498 | 5.90E-05 |
| Nsp2 | PPP3R2 | -0.499 | 5.81E-05 |
| Nsp2 | RASGRP2 | -0.499 | 5.83E-05 |
| Nsp4 | OSM | -0.499 | 5.67E-05 |
| Nsp5 | ADCY1 | -0.499 | 5.67E-05 |
| Nsp5 | ACVRL1 | -0.499 | 5.86E-05 |
| Nsp8 | MYC | -0.499 | 5.78E-05 |
| Nsp10 | DUSP8 | -0.499 | 5.66E-05 |
| Nsp12 | JMJD7-PLA2G | -0.499 | 5.78E-05 |
| Nsp12 | GREM1 | -0.499 | 5.73E-05 |
| Nsp12 | TNFRSF12A | -0.499 | 5.74E-05 |
| Nsp12 | BBC3 | -0.499 | 5.67E-05 |
| Nsp13 | NR4A1 | -0.499 | 5.66E-05 |
| Nsp13 | PKN3 | -0.499 | 5.82E-05 |
| Nsp13 | IFNL2 | -0.499 | 5.65E-05 |
| Nsp14 | TELO2 | -0.499 | 5.80E-05 |
| Nsp14 | PRKCG | -0.499 | 5.78E-05 |
| Nsp14 | LCK | -0.499 | 5.84E-05 |
| Nsp15 | FLNC | -0.499 | 5.71E-05 |
| Nsp16 | TNFRSF25 | -0.499 | 5.70E-05 |
| Nsp16 | ACSBG1 | -0.499 | 5.82E-05 |
| S | MUC5B | -0.499 | 5.82E-05 |
| S | ICAM1 | -0.499 | 5.69E-05 |
| S | CNTFR | -0.499 | 5.76E-05 |
| Orf8 | ITPR3 | -0.499 | 5.85E-05 |
| Orf8 | HSPA1A | -0.499 | 5.84E-05 |
| Nsp2 | GNA12 | -0.5 | 5.56E-05 |
| Nsp3 | TRIP10 | -0.5 | 5.46E-05 |
| Nsp3 | DHX58 | -0.5 | 5.61E-05 |
| Nsp3 | RAPGEF1 | -0.5 | 5.49E-05 |
| Nsp3 | LPAR2 | -0.5 | 5.49E-05 |
| Nsp4 | ITGA3 | -0.5 | 5.61E-05 |
| Nsp4 | TRIP10 | -0.5 | 5.58E-05 |
| Nsp4 | IL17C | -0.5 | 5.55E-05 |
| Nsp5 | CHRD | -0.5 | 5.44E-05 |

| | | | |
|---|---|---|---|
| Nsp5 | IL17RC | -0.5 | 5.59E-05 |
| Nsp5 | GRK6 | -0.5 | 5.51E-05 |
| Nsp5 | CACNG5 | -0.5 | 5.50E-05 |
| Nsp10 | LRP5 | -0.5 | 5.43E-05 |
| Nsp10 | TNF | -0.5 | 5.63E-05 |
| Nsp12 | IGF1R | -0.5 | 5.44E-05 |
| Nsp13 | TNFSF12 | -0.5 | 5.43E-05 |
| Nsp13 | ELAVL1 | -0.5 | 5.44E-05 |
| Nsp14 | NR4A1 | -0.5 | 5.46E-05 |
| Nsp14 | MAP4K2 | -0.5 | 5.55E-05 |
| Nsp14 | PFKL | -0.5 | 5.54E-05 |
| Nsp15 | TNFRSF25 | -0.5 | 5.51E-05 |
| Nsp15 | AKT1 | -0.5 | 5.43E-05 |
| Nsp15 | GFAP | -0.5 | 5.45E-05 |
| Nsp15 | PLA2G4D | -0.5 | 5.59E-05 |
| S | TNFRSF8 | -0.5 | 5.51E-05 |
| S | PPP1R3F | -0.5 | 5.50E-05 |
| S | ADCY4 | -0.5 | 5.54E-05 |
| Orf3a | SMAD7 | -0.5 | 5.55E-05 |
| Orf6 | EPOR | -0.5 | 5.55E-05 |
| Orf8 | CAPN1 | -0.5 | 5.62E-05 |
| Nsp2 | HSPA1A | -0.501 | 5.24E-05 |
| Nsp2 | ADCY5 | -0.501 | 5.32E-05 |
| Nsp4 | MAP4K2 | -0.501 | 5.34E-05 |
| Nsp5 | INSR | -0.501 | 5.33E-05 |
| Nsp7 | MAPK8IP2 | -0.501 | 5.33E-05 |
| Nsp8 | NFATC2 | -0.501 | 5.30E-05 |
| Nsp8 | FLCN | -0.501 | 5.32E-05 |
| Nsp8 | PDGFRB | -0.501 | 5.39E-05 |
| Nsp10 | NFATC2 | -0.501 | 5.22E-05 |
| Nsp12 | PRF1 | -0.501 | 5.38E-05 |
| Nsp12 | WNT3A | -0.501 | 5.23E-05 |
| Nsp12 | WNT10B | -0.501 | 5.26E-05 |
| Nsp12 | GNAI2 | -0.501 | 5.21E-05 |
| Nsp12 | ITGA2B | -0.501 | 5.22E-05 |
| Nsp13 | AKT1 | -0.501 | 5.32E-05 |
| Nsp13 | TNFSF14 | -0.501 | 5.42E-05 |
| Nsp13 | RAC3 | -0.501 | 5.35E-05 |
| Nsp13 | SESN2 | -0.501 | 5.24E-05 |
| Nsp13 | PFKP | -0.501 | 5.35E-05 |
| Nsp13 | ERBB2 | -0.501 | 5.22E-05 |
| Nsp14 | IRF3 | -0.501 | 5.26E-05 |
| Nsp14 | TNFRSF13C | -0.501 | 5.24E-05 |
| Nsp15 | FGF19 | -0.501 | 5.22E-05 |
| Nsp15 | NPRL3 | -0.501 | 5.31E-05 |
| Nsp15 | WNT7A | -0.501 | 5.41E-05 |
| Nsp15 | PIN1 | -0.501 | 5.34E-05 |
| Nsp16 | PRKCZ | -0.501 | 5.26E-05 |
| Nsp16 | RORC | -0.501 | 5.38E-05 |
| S | CREB3L1 | -0.501 | 5.27E-05 |

| | | | |
|---|---|---|---|
| Orf8 | IL34 | -0.501 | 5.29E-05 |
| Orf8 | MAPK11 | -0.501 | 5.25E-05 |
| Orf8 | PKN1 | -0.501 | 5.22E-05 |
| Orf8 | PLCB3 | -0.501 | 5.38E-05 |
| Nsp3 | PPP2R5D | -0.502 | 5.05E-05 |
| Nsp3 | VHL | -0.502 | 5.08E-05 |
| Nsp4 | PLCB2 | -0.502 | 5.15E-05 |
| Nsp4 | SRF | -0.502 | 5.06E-05 |
| Nsp4 | PLCG1 | -0.502 | 5.14E-05 |
| Nsp5 | LIF | -0.502 | 5.10E-05 |
| Nsp8 | CACNA1C | -0.502 | 5.06E-05 |
| Nsp8 | ACSBG1 | -0.502 | 5.14E-05 |
| Nsp8 | BCAR1 | -0.502 | 5.18E-05 |
| Nsp9 | ZAP70 | -0.502 | 5.16E-05 |
| Nsp10 | INHBB | -0.502 | 5.04E-05 |
| Nsp12 | ACTG1 | -0.502 | 5.14E-05 |
| Nsp13 | CACNA2D2 | -0.502 | 5.11E-05 |
| Nsp14 | ISG15 | -0.502 | 5.14E-05 |
| Nsp14 | EDAR | -0.502 | 5.15E-05 |
| Nsp14 | GDF5 | -0.502 | 5.14E-05 |
| Nsp15 | IGF1R | -0.502 | 5.16E-05 |
| Nsp15 | GREM1 | -0.502 | 5.16E-05 |
| Nsp15 | SRF | -0.502 | 5.11E-05 |
| S | EXOC7 | -0.502 | 5.13E-05 |
| Orf8 | SOCS1 | -0.502 | 5.16E-05 |
| Orf8 | EDAR | -0.502 | 5.04E-05 |
| Orf8 | AKT2 | -0.502 | 5.03E-05 |
| Orf8 | COMP | -0.502 | 5.09E-05 |
| Nsp2 | BMP8A | -0.503 | 4.81E-05 |
| Nsp2 | MAPK11 | -0.503 | 4.96E-05 |
| Nsp2 | MAP2K7 | -0.503 | 4.98E-05 |
| Nsp2 | CARD11 | -0.503 | 4.91E-05 |
| Nsp3 | PDGFA | -0.503 | 4.92E-05 |
| Nsp3 | GRK5 | -0.503 | 4.93E-05 |
| Nsp4 | DVL3 | -0.503 | 5.00E-05 |
| Nsp4 | TNFRSF8 | -0.503 | 4.97E-05 |
| Nsp4 | RPS6KA2 | -0.503 | 4.88E-05 |
| Nsp4 | WNT7A | -0.503 | 4.94E-05 |
| Nsp5 | INHBA | -0.503 | 4.92E-05 |
| Nsp5 | WNT7A | -0.503 | 4.91E-05 |
| Nsp5 | HSPA1A | -0.503 | 4.90E-05 |
| Nsp5 | FLNC | -0.503 | 4.90E-05 |
| Nsp7 | SMAD7 | -0.503 | 4.90E-05 |
| Nsp7 | NOG | -0.503 | 4.94E-05 |
| Nsp8 | FGF6 | -0.503 | 4.85E-05 |
| Nsp8 | TNFSF9 | -0.503 | 4.99E-05 |
| Nsp8 | PHLPP1 | -0.503 | 4.81E-05 |
| Nsp9 | BMP7 | -0.503 | 4.91E-05 |
| Nsp10 | STK11 | -0.503 | 4.93E-05 |
| Nsp12 | DUSP2 | -0.503 | 4.82E-05 |

| | | | |
|---|---|---|---|
| Nsp12 | NR4A1 | -0.503 | 4.84E-05 |
| Nsp12 | HSPA6 | -0.503 | 5.00E-05 |
| Nsp12 | WNT10A | -0.503 | 4.86E-05 |
| Nsp12 | LTB | -0.503 | 4.91E-05 |
| Nsp12 | CSH2 | -0.503 | 4.86E-05 |
| Nsp13 | PTK2B | -0.503 | 4.95E-05 |
| Nsp13 | SREBF1 | -0.503 | 4.95E-05 |
| Nsp14 | RRAS | -0.503 | 4.94E-05 |
| Nsp14 | FGF3 | -0.503 | 5.00E-05 |
| Nsp15 | ITGB5 | -0.503 | 4.83E-05 |
| Nsp16 | IL27 | -0.503 | 4.90E-05 |
| Nsp16 | PTPN5 | -0.503 | 4.90E-05 |
| S | CHRD | -0.503 | 4.97E-05 |
| S | PRF1 | -0.503 | 4.99E-05 |
| S | TRIM25 | -0.503 | 4.99E-05 |
| Orf3a | FZD9 | -0.503 | 4.88E-05 |
| Orf8 | VAV2 | -0.503 | 4.94E-05 |
| Orf8 | FMOD | -0.503 | 4.94E-05 |
| Orf8 | CACNA2D4 | -0.503 | 4.93E-05 |
| Orf8 | SMAD3 | -0.503 | 4.93E-05 |
| Orf8 | WNT1 | -0.503 | 4.96E-05 |
| Orf8 | LAMA5 | -0.503 | 4.93E-05 |
| Orf8 | THBS2 | -0.503 | 4.85E-05 |
| Nsp2 | PRKAR1B | -0.504 | 4.74E-05 |
| Nsp3 | TNF | -0.504 | 4.75E-05 |
| Nsp3 | NPRL3 | -0.504 | 4.72E-05 |
| Nsp5 | NR4A1 | -0.504 | 4.70E-05 |
| Nsp5 | PFKP | -0.504 | 4.64E-05 |
| Nsp5 | GNA12 | -0.504 | 4.73E-05 |
| Nsp8 | DUSP3 | -0.504 | 4.63E-05 |
| Nsp8 | LIF | -0.504 | 4.64E-05 |
| Nsp8 | IGF2 | -0.504 | 4.74E-05 |
| Nsp10 | INSR | -0.504 | 4.67E-05 |
| Nsp10 | RXRA | -0.504 | 4.75E-05 |
| Nsp12 | PTK2B | -0.504 | 4.65E-05 |
| Nsp12 | PRKAB1 | -0.504 | 4.72E-05 |
| Nsp12 | CXCL2 | -0.504 | 4.67E-05 |
| Nsp12 | NPRL3 | -0.504 | 4.74E-05 |
| Nsp13 | VHL | -0.504 | 4.62E-05 |
| Nsp14 | TNXB | -0.504 | 4.71E-05 |
| Nsp14 | MAPK3 | -0.504 | 4.66E-05 |
| Nsp14 | BAD | -0.504 | 4.73E-05 |
| Nsp15 | IL17B | -0.504 | 4.71E-05 |
| Nsp15 | JAK3 | -0.504 | 4.77E-05 |
| Nsp16 | CSF2RB | -0.504 | 4.79E-05 |
| Nsp16 | RAC3 | -0.504 | 4.66E-05 |
| Nsp16 | ECSIT | -0.504 | 4.76E-05 |
| Nsp16 | PRKACA | -0.504 | 4.74E-05 |
| Nsp16 | ITGA2B | -0.504 | 4.64E-05 |
| Nsp16 | CXCL14 | -0.504 | 4.70E-05 |

| | | | |
|---|---|---|---|
| S | PLCB2 | -0.504 | 4.79E-05 |
| Orf3a | RPS6KA4 | -0.504 | 4.71E-05 |
| Orf3a | LEFTY1 | -0.504 | 4.65E-05 |
| Orf3a | SRF | -0.504 | 4.79E-05 |
| Orf8 | PLCB2 | -0.504 | 4.63E-05 |
| Orf8 | GRK1 | -0.504 | 4.72E-05 |
| Orf8 | MAPK8IP3 | -0.504 | 4.74E-05 |
| Nsp2 | INHBB | -0.505 | 4.45E-05 |
| Nsp2 | COL6A1 | -0.505 | 4.46E-05 |
| Nsp3 | JMJD7-PLA2G | -0.505 | 4.49E-05 |
| Nsp3 | DUSP2 | -0.505 | 4.53E-05 |
| Nsp3 | GH2 | -0.505 | 4.53E-05 |
| Nsp4 | FGFR4 | -0.505 | 4.61E-05 |
| Nsp4 | RAC2 | -0.505 | 4.59E-05 |
| Nsp4 | FLCN | -0.505 | 4.50E-05 |
| Nsp5 | GNAI2 | -0.505 | 4.55E-05 |
| Nsp8 | AKT1 | -0.505 | 4.44E-05 |
| Nsp8 | PIK3R5 | -0.505 | 4.45E-05 |
| Nsp8 | ATP6V1B1 | -0.505 | 4.49E-05 |
| Nsp8 | CAPN2 | -0.505 | 4.44E-05 |
| Nsp8 | AKT2 | -0.505 | 4.52E-05 |
| Nsp9 | DUSP7 | -0.505 | 4.47E-05 |
| Nsp12 | TNFSF12 | -0.505 | 4.56E-05 |
| Nsp12 | TNFRSF1B | -0.505 | 4.45E-05 |
| Nsp12 | PRKACG | -0.505 | 4.48E-05 |
| Nsp12 | SOCS3 | -0.505 | 4.45E-05 |
| Nsp13 | CPT1B | -0.505 | 4.54E-05 |
| Nsp14 | AKT1 | -0.505 | 4.48E-05 |
| Nsp15 | NR4A1 | -0.505 | 4.54E-05 |
| Nsp15 | NGFR | -0.505 | 4.44E-05 |
| Nsp15 | RPS6KB2 | -0.505 | 4.47E-05 |
| Nsp15 | SOCS3 | -0.505 | 4.43E-05 |
| Nsp16 | CCL14 | -0.505 | 4.47E-05 |
| Nsp16 | PDGFB | -0.505 | 4.60E-05 |
| Nsp16 | FOXP3 | -0.505 | 4.60E-05 |
| S | CXCR5 | -0.505 | 4.57E-05 |
| S | GADD45A | -0.505 | 4.58E-05 |
| S | CACNB1 | -0.505 | 4.58E-05 |
| S | IL17C | -0.505 | 4.56E-05 |
| Orf3a | GDF1 | -0.505 | 4.59E-05 |
| Orf8 | NGFR | -0.505 | 4.57E-05 |
| Orf8 | ADCY3 | -0.505 | 4.58E-05 |
| Nsp2 | BMP6 | -0.506 | 4.28E-05 |
| Nsp3 | EDAR | -0.506 | 4.35E-05 |
| Nsp4 | CXCR5 | -0.506 | 4.42E-05 |
| Nsp5 | IRF7 | -0.506 | 4.38E-05 |
| Nsp8 | FOSL1 | -0.506 | 4.27E-05 |
| Nsp8 | CNTFR | -0.506 | 4.41E-05 |
| Nsp9 | FZD9 | -0.506 | 4.43E-05 |
| Nsp10 | MAP2K7 | -0.506 | 4.42E-05 |

| | | | |
|---|---|---|---|
| Nsp12 | MAP4K2 | -0.506 | 4.30E-05 |
| Nsp12 | ECSIT | -0.506 | 4.32E-05 |
| Nsp14 | CREB3L1 | -0.506 | 4.27E-05 |
| Nsp15 | MAPK15 | -0.506 | 4.26E-05 |
| Nsp16 | JMJD7-PLA2G | -0.506 | 4.30E-05 |
| Nsp16 | PPP2R1A | -0.506 | 4.29E-05 |
| Nsp16 | NPRL3 | -0.506 | 4.33E-05 |
| S | ATP6V1B1 | -0.506 | 4.39E-05 |
| S | HSPA6 | -0.506 | 4.38E-05 |
| S | RPS6KB2 | -0.506 | 4.27E-05 |
| Orf3a | EFNA2 | -0.506 | 4.33E-05 |
| Orf8 | PRF1 | -0.506 | 4.34E-05 |
| Orf8 | MAPK3 | -0.506 | 4.36E-05 |
| Orf8 | CACNG4 | -0.506 | 4.32E-05 |
| Orf8 | TNFRSF1B | -0.506 | 4.43E-05 |
| Orf8 | RUNX1 | -0.506 | 4.32E-05 |
| Nsp2 | CDKN2B | -0.507 | 4.13E-05 |
| Nsp3 | CCR7 | -0.507 | 4.20E-05 |
| Nsp3 | PLCG2 | -0.507 | 4.14E-05 |
| Nsp4 | PTK2B | -0.507 | 4.20E-05 |
| Nsp4 | CXCL3 | -0.507 | 4.20E-05 |
| Nsp5 | CHAD | -0.507 | 4.20E-05 |
| Nsp5 | FGF17 | -0.507 | 4.24E-05 |
| Nsp8 | PRF1 | -0.507 | 4.14E-05 |
| Nsp8 | ITGA3 | -0.507 | 4.25E-05 |
| Nsp8 | CHRM1 | -0.507 | 4.23E-05 |
| Nsp12 | PRKCZ | -0.507 | 4.13E-05 |
| Nsp12 | ITGB5 | -0.507 | 4.25E-05 |
| Nsp12 | BCAR1 | -0.507 | 4.15E-05 |
| Nsp12 | CPT1B | -0.507 | 4.20E-05 |
| Nsp12 | ACVRL1 | -0.507 | 4.13E-05 |
| Nsp12 | RAPGEF1 | -0.507 | 4.10E-05 |
| Nsp13 | PLA2G4B | -0.507 | 4.18E-05 |
| Nsp13 | LMNA | -0.507 | 4.22E-05 |
| Nsp13 | FLNC | -0.507 | 4.09E-05 |
| Nsp14 | NFKBIB | -0.507 | 4.23E-05 |
| Nsp14 | HCK | -0.507 | 4.23E-05 |
| Nsp14 | MAP3K11 | -0.507 | 4.16E-05 |
| Nsp14 | FGF17 | -0.507 | 4.21E-05 |
| S | AKT1 | -0.507 | 4.09E-05 |
| S | TNFSF9 | -0.507 | 4.22E-05 |
| S | TRAF2 | -0.507 | 4.12E-05 |
| S | GNB2 | -0.507 | 4.21E-05 |
| S | TICAM1 | -0.507 | 4.21E-05 |
| Orf8 | BMP6 | -0.507 | 4.17E-05 |
| Orf8 | CDKN2B | -0.507 | 4.19E-05 |
| Orf8 | FGF17 | -0.507 | 4.13E-05 |
| Orf8 | COL6A1 | -0.507 | 4.20E-05 |
| Nsp2 | ISG15 | -0.508 | 3.94E-05 |
| Nsp2 | GATA3 | -0.508 | 3.96E-05 |

| | | | |
|---|---|---|---|
| Nsp2 | TNFRSF4 | -0.508 | 4.08E-05 |
| Nsp3 | ITGA3 | -0.508 | 4.04E-05 |
| Nsp3 | TSC2 | -0.508 | 4.02E-05 |
| Nsp3 | FOSL1 | -0.508 | 3.94E-05 |
| Nsp4 | PFKP | -0.508 | 3.97E-05 |
| Nsp5 | CAMKK2 | -0.508 | 4.00E-05 |
| Nsp5 | RGMA | -0.508 | 4.06E-05 |
| Nsp5 | BCL3 | -0.508 | 4.03E-05 |
| Nsp8 | ACVR2B | -0.508 | 4.01E-05 |
| Nsp8 | GADD45A | -0.508 | 4.03E-05 |
| Nsp10 | GDF10 | -0.508 | 4.09E-05 |
| Nsp10 | IRS2 | -0.508 | 4.06E-05 |
| Nsp10 | TGFB1 | -0.508 | 3.94E-05 |
| Nsp12 | TLR9 | -0.508 | 4.03E-05 |
| Nsp12 | CSH1 | -0.508 | 3.97E-05 |
| Nsp12 | TYK2 | -0.508 | 4.05E-05 |
| Nsp12 | FADD | -0.508 | 3.97E-05 |
| Nsp13 | CCR7 | -0.508 | 3.98E-05 |
| Nsp15 | CSF2RB | -0.508 | 4.02E-05 |
| Nsp15 | LMNA | -0.508 | 3.93E-05 |
| Nsp15 | WNT4 | -0.508 | 4.07E-05 |
| Nsp15 | CXCL14 | -0.508 | 4.09E-05 |
| Nsp15 | TNFRSF13C | -0.508 | 4.08E-05 |
| Nsp16 | EDAR | -0.508 | 3.94E-05 |
| Nsp16 | RUNX1 | -0.508 | 4.04E-05 |
| S | NTRK1 | -0.508 | 3.99E-05 |
| S | RUNX1 | -0.508 | 4.08E-05 |
| Orf8 | MKNK2 | -0.508 | 4.05E-05 |
| Nsp2 | CACNA1I | -0.509 | 3.77E-05 |
| Nsp3 | TNFSF9 | -0.509 | 3.87E-05 |
| Nsp3 | CACNA2D2 | -0.509 | 3.87E-05 |
| Nsp3 | PIN1 | -0.509 | 3.88E-05 |
| Nsp4 | GRK6 | -0.509 | 3.77E-05 |
| Nsp4 | NGFR | -0.509 | 3.80E-05 |
| Nsp4 | GNB2 | -0.509 | 3.88E-05 |
| Nsp4 | GDF5 | -0.509 | 3.81E-05 |
| Nsp5 | TNXB | -0.509 | 3.89E-05 |
| Nsp5 | ARAF | -0.509 | 3.82E-05 |
| Nsp5 | LAMC3 | -0.509 | 3.80E-05 |
| Nsp5 | MYC | -0.509 | 3.84E-05 |
| Nsp8 | TNFRSF8 | -0.509 | 3.77E-05 |
| Nsp8 | NFKBIB | -0.509 | 3.80E-05 |
| Nsp8 | GRK6 | -0.509 | 3.90E-05 |
| Nsp8 | PLCB2 | -0.509 | 3.78E-05 |
| Nsp8 | ARAF | -0.509 | 3.77E-05 |
| Nsp8 | TRIM25 | -0.509 | 3.83E-05 |
| Nsp12 | ISG15 | -0.509 | 3.81E-05 |
| Nsp13 | NFKBIB | -0.509 | 3.78E-05 |
| Nsp13 | SRF | -0.509 | 3.78E-05 |
| Nsp13 | WNT10A | -0.509 | 3.91E-05 |

| | | | |
|---|---|---|---|
| Nsp13 | IL12RB1 | -0.509 | 3.82E-05 |
| Nsp13 | PPP1R3F | -0.509 | 3.89E-05 |
| Nsp13 | PDGFRB | -0.509 | 3.92E-05 |
| Nsp14 | PFKFB3 | -0.509 | 3.80E-05 |
| Nsp15 | CDKN2B | -0.509 | 3.80E-05 |
| Nsp15 | TYK2 | -0.509 | 3.78E-05 |
| Nsp15 | IL21R | -0.509 | 3.78E-05 |
| Nsp16 | PLA2G4D | -0.509 | 3.84E-05 |
| Nsp16 | HK2 | -0.509 | 3.77E-05 |
| S | PLCG1 | -0.509 | 3.82E-05 |
| Orf8 | CACNA1G | -0.509 | 3.78E-05 |
| Orf8 | DUSP4 | -0.509 | 3.83E-05 |
| Orf8 | GDF1 | -0.509 | 3.89E-05 |
| Orf8 | IL21R | -0.509 | 3.81E-05 |
| Orf8 | LCK | -0.509 | 3.78E-05 |
| Nsp2 | BMP8B | -0.51 | 3.68E-05 |
| Nsp3 | NFATC2 | -0.51 | 3.76E-05 |
| Nsp3 | PPP1R3F | -0.51 | 3.69E-05 |
| Nsp4 | NR4A1 | -0.51 | 3.71E-05 |
| Nsp4 | TAB1 | -0.51 | 3.74E-05 |
| Nsp5 | NTRK1 | -0.51 | 3.71E-05 |
| Nsp5 | PPP5C | -0.51 | 3.71E-05 |
| Nsp8 | PRKCZ | -0.51 | 3.75E-05 |
| Nsp8 | EDAR | -0.51 | 3.76E-05 |
| Nsp8 | WNT5A | -0.51 | 3.74E-05 |
| Nsp10 | CEBPB | -0.51 | 3.68E-05 |
| Nsp10 | RPS6KA2 | -0.51 | 3.63E-05 |
| Nsp12 | SMAD3 | -0.51 | 3.62E-05 |
| Nsp12 | RPS6KB2 | -0.51 | 3.74E-05 |
| Nsp12 | IL12RB1 | -0.51 | 3.67E-05 |
| Nsp13 | PLCG2 | -0.51 | 3.66E-05 |
| Nsp13 | PFKL | -0.51 | 3.64E-05 |
| Nsp13 | ADCY4 | -0.51 | 3.68E-05 |
| Nsp14 | CACNA1C | -0.51 | 3.74E-05 |
| Nsp14 | CD70 | -0.51 | 3.65E-05 |
| Nsp14 | COL6A2 | -0.51 | 3.67E-05 |
| Nsp15 | CACNA1C | -0.51 | 3.67E-05 |
| Nsp15 | MAPK8IP1 | -0.51 | 3.69E-05 |
| Nsp15 | EDAR | -0.51 | 3.70E-05 |
| Nsp15 | PDGFB | -0.51 | 3.74E-05 |
| Nsp15 | RASGRF1 | -0.51 | 3.76E-05 |
| Nsp16 | PPP3R2 | -0.51 | 3.73E-05 |
| Nsp16 | FGR | -0.51 | 3.67E-05 |
| Nsp16 | CPT1C | -0.51 | 3.71E-05 |
| S | CAPN2 | -0.51 | 3.74E-05 |
| S | LCK | -0.51 | 3.65E-05 |
| S | RASGRF1 | -0.51 | 3.68E-05 |
| Orf8 | ITGB4 | -0.51 | 3.69E-05 |
| Orf8 | ARAF | -0.51 | 3.73E-05 |
| Orf8 | CSH1 | -0.51 | 3.64E-05 |

| | | | |
|---|---|---|---|
| Orf8 | POMC | -0.51 | 3.73E-05 |
| Orf8 | TNFRSF13C | -0.51 | 3.72E-05 |
| Orf8 | SOCS3 | -0.51 | 3.67E-05 |
| Nsp2 | SLC7A5 | -0.511 | 3.55E-05 |
| Nsp3 | GREM1 | -0.511 | 3.60E-05 |
| Nsp3 | TCL1A | -0.511 | 3.50E-05 |
| Nsp5 | TRIP10 | -0.511 | 3.55E-05 |
| Nsp8 | FGFR4 | -0.511 | 3.61E-05 |
| Nsp8 | FGF18 | -0.511 | 3.61E-05 |
| Nsp8 | DUSP2 | -0.511 | 3.60E-05 |
| Nsp10 | FGF8 | -0.511 | 3.56E-05 |
| Nsp12 | GFAP | -0.511 | 3.61E-05 |
| Nsp13 | CCND3 | -0.511 | 3.55E-05 |
| Nsp13 | IRF7 | -0.511 | 3.49E-05 |
| Nsp13 | DUSP3 | -0.511 | 3.53E-05 |
| Nsp13 | ACTG1 | -0.511 | 3.47E-05 |
| Nsp13 | ID2 | -0.511 | 3.52E-05 |
| Nsp13 | ENO3 | -0.511 | 3.61E-05 |
| Nsp14 | WNT9B | -0.511 | 3.54E-05 |
| Nsp15 | FGFR4 | -0.511 | 3.47E-05 |
| Nsp15 | DUSP3 | -0.511 | 3.49E-05 |
| Nsp15 | CARD10 | -0.511 | 3.60E-05 |
| Nsp15 | DHX58 | -0.511 | 3.51E-05 |
| Nsp15 | GRK5 | -0.511 | 3.51E-05 |
| Nsp15 | PFKL | -0.511 | 3.61E-05 |
| Nsp15 | RAPGEF1 | -0.511 | 3.61E-05 |
| Nsp16 | TNFRSF8 | -0.511 | 3.47E-05 |
| Nsp16 | PFKP | -0.511 | 3.52E-05 |
| Nsp16 | CLCF1 | -0.511 | 3.53E-05 |
| Nsp16 | GFAP | -0.511 | 3.59E-05 |
| Nsp16 | IL21R | -0.511 | 3.49E-05 |
| S | CAMKK2 | -0.511 | 3.48E-05 |
| S | LAMC3 | -0.511 | 3.58E-05 |
| S | ECSIT | -0.511 | 3.55E-05 |
| S | RAC2 | -0.511 | 3.58E-05 |
| S | GNAI2 | -0.511 | 3.60E-05 |
| S | TCL1A | -0.511 | 3.50E-05 |
| Orf8 | CACNA1B | -0.511 | 3.58E-05 |
| Nsp2 | DUSP7 | -0.512 | 3.41E-05 |
| Nsp2 | SRF | -0.512 | 3.39E-05 |
| Nsp2 | RARA | -0.512 | 3.38E-05 |
| Nsp2 | NFATC1 | -0.512 | 3.38E-05 |
| Nsp3 | CCND3 | -0.512 | 3.41E-05 |
| Nsp3 | GDF5 | -0.512 | 3.44E-05 |
| Nsp4 | CHRD | -0.512 | 3.41E-05 |
| Nsp4 | WDR24 | -0.512 | 3.35E-05 |
| Nsp4 | JAK3 | -0.512 | 3.37E-05 |
| Nsp5 | DUSP2 | -0.512 | 3.36E-05 |
| Nsp5 | PDGFRB | -0.512 | 3.37E-05 |
| Nsp5 | CRK | -0.512 | 3.43E-05 |

| | | | |
|---|---|---|---|
| Nsp5 | BAD | -0.512 | 3.46E-05 |
| Nsp8 | MAP4K2 | -0.512 | 3.42E-05 |
| Nsp8 | CD70 | -0.512 | 3.39E-05 |
| Nsp8 | VHL | -0.512 | 3.45E-05 |
| Nsp8 | TICAM1 | -0.512 | 3.42E-05 |
| Nsp10 | NFATC1 | -0.512 | 3.46E-05 |
| Nsp12 | DUSP3 | -0.512 | 3.35E-05 |
| Nsp12 | ARAF | -0.512 | 3.46E-05 |
| Nsp12 | EDAR | -0.512 | 3.33E-05 |
| Nsp12 | CACNA2D4 | -0.512 | 3.34E-05 |
| Nsp13 | IFNL3 | -0.512 | 3.36E-05 |
| Nsp13 | ISG15 | -0.512 | 3.46E-05 |
| Nsp13 | CSH1 | -0.512 | 3.34E-05 |
| Nsp13 | PLCG1 | -0.512 | 3.38E-05 |
| Nsp14 | PITX2 | -0.512 | 3.45E-05 |
| Nsp14 | FGF18 | -0.512 | 3.40E-05 |
| Nsp14 | AKT2 | -0.512 | 3.45E-05 |
| Nsp14 | MKNK2 | -0.512 | 3.41E-05 |
| Nsp14 | CXCL14 | -0.512 | 3.40E-05 |
| Nsp15 | CD14 | -0.512 | 3.44E-05 |
| Nsp15 | BCAR1 | -0.512 | 3.45E-05 |
| Nsp16 | TRAF1 | -0.512 | 3.44E-05 |
| Nsp16 | ATP6V1B1 | -0.512 | 3.43E-05 |
| Nsp16 | HSPA6 | -0.512 | 3.47E-05 |
| Nsp16 | RAPGEF1 | -0.512 | 3.45E-05 |
| S | HCK | -0.512 | 3.43E-05 |
| S | NR4A1 | -0.512 | 3.46E-05 |
| S | ARAF | -0.512 | 3.33E-05 |
| S | FLCN | -0.512 | 3.40E-05 |
| Orf3a | JUN | -0.512 | 3.39E-05 |
| Orf8 | HSPA2 | -0.512 | 3.44E-05 |
| Orf8 | MAPK4 | -0.512 | 3.42E-05 |
| Orf8 | PIK3R2 | -0.512 | 3.40E-05 |
| Orf8 | PIK3CD | -0.512 | 3.35E-05 |
| Nsp2 | PKN1 | -0.513 | 3.26E-05 |
| Nsp3 | PTK2B | -0.513 | 3.29E-05 |
| Nsp3 | ECSIT | -0.513 | 3.25E-05 |
| Nsp4 | CAMKK2 | -0.513 | 3.25E-05 |
| Nsp4 | EPOR | -0.513 | 3.24E-05 |
| Nsp4 | GYS1 | -0.513 | 3.28E-05 |
| Nsp4 | CACNB1 | -0.513 | 3.25E-05 |
| Nsp4 | TSC2 | -0.513 | 3.28E-05 |
| Nsp4 | PPP5C | -0.513 | 3.24E-05 |
| Nsp4 | TYK2 | -0.513 | 3.24E-05 |
| Nsp5 | ADCY4 | -0.513 | 3.31E-05 |
| Nsp8 | TNFRSF25 | -0.513 | 3.31E-05 |
| Nsp8 | TBX21 | -0.513 | 3.32E-05 |
| Nsp8 | CXCL14 | -0.513 | 3.23E-05 |
| Nsp9 | ICAM1 | -0.513 | 3.20E-05 |
| Nsp12 | CACNB1 | -0.513 | 3.21E-05 |

| | | | |
|---|---|---|---|
| Nsp12 | WNT4 | -0.513 | 3.27E-05 |
| Nsp13 | ARAF | -0.513 | 3.32E-05 |
| Nsp13 | PLA2G4D | -0.513 | 3.20E-05 |
| Nsp13 | NPRL3 | -0.513 | 3.32E-05 |
| Nsp13 | BCAR1 | -0.513 | 3.24E-05 |
| Nsp14 | ADCY4 | -0.513 | 3.27E-05 |
| Nsp15 | PTK2B | -0.513 | 3.19E-05 |
| Nsp15 | CCR10 | -0.513 | 3.21E-05 |
| Nsp15 | TRIP10 | -0.513 | 3.21E-05 |
| Nsp15 | WDR24 | -0.513 | 3.32E-05 |
| Nsp15 | SMAD3 | -0.513 | 3.26E-05 |
| Nsp15 | IL12RB1 | -0.513 | 3.24E-05 |
| Nsp16 | THSD4 | -0.513 | 3.31E-05 |
| Nsp16 | DUSP2 | -0.513 | 3.26E-05 |
| Nsp16 | FLT3LG | -0.513 | 3.28E-05 |
| Nsp16 | TRIP10 | -0.513 | 3.28E-05 |
| S | MAPK12 | -0.513 | 3.21E-05 |
| S | PHLPP1 | -0.513 | 3.21E-05 |
| Orf8 | PHLPP1 | -0.513 | 3.32E-05 |
| Nsp3 | CXCL3 | -0.514 | 3.17E-05 |
| Nsp3 | BBC3 | -0.514 | 3.16E-05 |
| Nsp4 | PRF1 | -0.514 | 3.16E-05 |
| Nsp4 | PPP2R1A | -0.514 | 3.11E-05 |
| Nsp4 | GRK5 | -0.514 | 3.10E-05 |
| Nsp5 | FGFR4 | -0.514 | 3.12E-05 |
| Nsp5 | FLCN | -0.514 | 3.13E-05 |
| Nsp5 | GRK1 | -0.514 | 3.12E-05 |
| Nsp8 | FMOD | -0.514 | 3.14E-05 |
| Nsp8 | CACNA2D4 | -0.514 | 3.14E-05 |
| Nsp8 | CPT1C | -0.514 | 3.17E-05 |
| Nsp12 | WNT9A | -0.514 | 3.10E-05 |
| Nsp12 | WDR24 | -0.514 | 3.11E-05 |
| Nsp13 | STAT5A | -0.514 | 3.13E-05 |
| Nsp13 | CSF2RB | -0.514 | 3.14E-05 |
| Nsp13 | WNT4 | -0.514 | 3.11E-05 |
| Nsp13 | TNFRSF1B | -0.514 | 3.18E-05 |
| Nsp14 | MAPK8IP1 | -0.514 | 3.18E-05 |
| Nsp14 | CXCR3 | -0.514 | 3.16E-05 |
| Nsp14 | FGF19 | -0.514 | 3.11E-05 |
| Nsp14 | ID4 | -0.514 | 3.15E-05 |
| Nsp14 | WNT10A | -0.514 | 3.07E-05 |
| Nsp14 | CACNG5 | -0.514 | 3.15E-05 |
| Nsp15 | TNFSF14 | -0.514 | 3.15E-05 |
| Nsp15 | BAD | -0.514 | 3.16E-05 |
| Nsp15 | CSH2 | -0.514 | 3.10E-05 |
| S | FGFR4 | -0.514 | 3.17E-05 |
| S | ADCY1 | -0.514 | 3.13E-05 |
| S | FMOD | -0.514 | 3.10E-05 |
| S | TRIP10 | -0.514 | 3.11E-05 |
| S | TBX21 | -0.514 | 3.09E-05 |

| | | | |
|---|---|---|---|
| S | TYK2 | -0.514 | 3.11E-05 |
| S | CXCL14 | -0.514 | 3.18E-05 |
| S | TNFRSF13C | -0.514 | 3.06E-05 |
| Orf8 | CCND3 | -0.514 | 3.13E-05 |
| Orf8 | CAPN2 | -0.514 | 3.10E-05 |
| Nsp2 | MAP2K2 | -0.515 | 2.96E-05 |
| Nsp3 | NTRK1 | -0.515 | 3.02E-05 |
| Nsp3 | MAP4K2 | -0.515 | 2.94E-05 |
| Nsp3 | IGF1R | -0.515 | 3.06E-05 |
| Nsp3 | LCK | -0.515 | 3.06E-05 |
| Nsp3 | CXCL14 | -0.515 | 3.02E-05 |
| Nsp4 | NFKBIB | -0.515 | 2.95E-05 |
| Nsp5 | MUC5B | -0.515 | 3.03E-05 |
| Nsp8 | FGF3 | -0.515 | 3.05E-05 |
| Nsp8 | PDGFB | -0.515 | 2.95E-05 |
| Nsp10 | CALML5 | -0.515 | 3.02E-05 |
| Nsp12 | EPOR | -0.515 | 3.03E-05 |
| Nsp12 | FLCN | -0.515 | 2.99E-05 |
| Nsp12 | GNB2 | -0.515 | 3.02E-05 |
| Nsp12 | MKNK2 | -0.515 | 3.00E-05 |
| Nsp12 | RUNX1 | -0.515 | 3.02E-05 |
| Nsp13 | CXCL3 | -0.515 | 3.03E-05 |
| Nsp13 | RUNX1 | -0.515 | 3.01E-05 |
| Nsp13 | WNT5A | -0.515 | 3.03E-05 |
| Nsp14 | CACNG4 | -0.515 | 2.95E-05 |
| Nsp14 | MAPK13 | -0.515 | 3.05E-05 |
| Nsp15 | IL27 | -0.515 | 3.06E-05 |
| Nsp15 | ITPR3 | -0.515 | 2.94E-05 |
| Nsp15 | GRK6 | -0.515 | 3.01E-05 |
| Nsp16 | TSC2 | -0.515 | 2.96E-05 |
| Nsp16 | FADD | -0.515 | 3.00E-05 |
| S | TNFSF14 | -0.515 | 2.94E-05 |
| Orf3a | LPAR5 | -0.515 | 2.95E-05 |
| Orf8 | CTSD | -0.515 | 2.99E-05 |
| Orf8 | LAMC3 | -0.515 | 3.02E-05 |
| Orf8 | GDF6 | -0.515 | 2.95E-05 |
| Orf8 | TNFRSF4 | -0.515 | 2.96E-05 |
| Orf8 | FZD2 | -0.515 | 2.99E-05 |
| Nsp2 | TBKBP1 | -0.516 | 2.93E-05 |
| Nsp2 | MAPK8IP1 | -0.516 | 2.87E-05 |
| Nsp3 | TNFSF14 | -0.516 | 2.91E-05 |
| Nsp3 | PPP3R2 | -0.516 | 2.85E-05 |
| Nsp3 | EXOC7 | -0.516 | 2.92E-05 |
| Nsp4 | EPHA2 | -0.516 | 2.89E-05 |
| Nsp5 | DUSP1 | -0.516 | 2.84E-05 |
| Nsp5 | MAPK13 | -0.516 | 2.83E-05 |
| Nsp5 | PDGFB | -0.516 | 2.86E-05 |
| Nsp5 | JAK3 | -0.516 | 2.93E-05 |
| Nsp8 | NTRK1 | -0.516 | 2.91E-05 |
| Nsp8 | LMNA | -0.516 | 2.85E-05 |

| | | | |
|---|---|---|---|
| Nsp9 | EFNA2 | -0.516 | 2.89E-05 |
| Nsp12 | DVL3 | -0.516 | 2.88E-05 |
| Nsp13 | JMJD7-PLA2G | -0.516 | 2.85E-05 |
| Nsp13 | HCK | -0.516 | 2.89E-05 |
| Nsp13 | CACNB1 | -0.516 | 2.90E-05 |
| Nsp14 | ITPR3 | -0.516 | 2.84E-05 |
| Nsp14 | CACNG1 | -0.516 | 2.92E-05 |
| Nsp15 | LAMC3 | -0.516 | 2.84E-05 |
| Nsp16 | CD14 | -0.516 | 2.83E-05 |
| Nsp16 | SREBF1 | -0.516 | 2.88E-05 |
| S | NFKBIB | -0.516 | 2.87E-05 |
| Orf8 | CSF2RB | -0.516 | 2.88E-05 |
| Orf8 | FGFR3 | -0.516 | 2.84E-05 |
| Nsp2 | SOCS1 | -0.517 | 2.81E-05 |
| Nsp2 | COMP | -0.517 | 2.76E-05 |
| Nsp2 | VEGFA | -0.517 | 2.75E-05 |
| Nsp3 | RAC2 | -0.517 | 2.74E-05 |
| Nsp3 | RUNX1 | -0.517 | 2.72E-05 |
| Nsp4 | LAMC3 | -0.517 | 2.80E-05 |
| Nsp4 | RPS6KB2 | -0.517 | 2.73E-05 |
| Nsp4 | RUNX1 | -0.517 | 2.75E-05 |
| Nsp5 | YWHAG | -0.517 | 2.71E-05 |
| Nsp8 | DVL3 | -0.517 | 2.81E-05 |
| Nsp8 | CCND2 | -0.517 | 2.73E-05 |
| Nsp8 | LAMC3 | -0.517 | 2.76E-05 |
| Nsp8 | MCL1 | -0.517 | 2.73E-05 |
| Nsp8 | SRF | -0.517 | 2.80E-05 |
| Nsp8 | WNT4 | -0.517 | 2.72E-05 |
| Nsp10 | THSD4 | -0.517 | 2.74E-05 |
| Nsp10 | CACNG8 | -0.517 | 2.77E-05 |
| Nsp12 | CCND3 | -0.517 | 2.80E-05 |
| Nsp12 | NFATC2 | -0.517 | 2.74E-05 |
| Nsp12 | RAC3 | -0.517 | 2.72E-05 |
| Nsp12 | POMC | -0.517 | 2.80E-05 |
| Nsp13 | PDGFA | -0.517 | 2.77E-05 |
| Nsp13 | ITGA3 | -0.517 | 2.76E-05 |
| Nsp14 | CPT1C | -0.517 | 2.75E-05 |
| Nsp15 | PLCB2 | -0.517 | 2.74E-05 |
| Nsp15 | CACNA2D2 | -0.517 | 2.74E-05 |
| Nsp16 | CACNB1 | -0.517 | 2.72E-05 |
| Nsp16 | FGF3 | -0.517 | 2.80E-05 |
| S | ACKR3 | -0.517 | 2.79E-05 |
| S | PFKL | -0.517 | 2.76E-05 |
| Orf3a | GDF11 | -0.517 | 2.77E-05 |
| Orf8 | BCL3 | -0.517 | 2.80E-05 |
| Orf8 | ACKR3 | -0.517 | 2.80E-05 |
| Nsp3 | LMNA | -0.518 | 2.66E-05 |
| Nsp4 | CDC25B | -0.518 | 2.68E-05 |
| Nsp4 | GSK3A | -0.518 | 2.64E-05 |
| Nsp4 | NFATC2 | -0.518 | 2.60E-05 |

| | | | |
|---|---|---|---|
| Nsp5 | PLCB3 | -0.518 | 2.63E-05 |
| Nsp8 | GNB2 | -0.518 | 2.62E-05 |
| Nsp8 | MLST8 | -0.518 | 2.67E-05 |
| Nsp8 | PIN1 | -0.518 | 2.64E-05 |
| Nsp10 | ACKR3 | -0.518 | 2.65E-05 |
| Nsp12 | MAP3K6 | -0.518 | 2.67E-05 |
| Nsp13 | IL27RA | -0.518 | 2.61E-05 |
| Nsp13 | INHBA | -0.518 | 2.69E-05 |
| Nsp13 | OSM | -0.518 | 2.64E-05 |
| Nsp13 | PIN1 | -0.518 | 2.64E-05 |
| Nsp14 | CAMKK2 | -0.518 | 2.61E-05 |
| Nsp14 | PRF1 | -0.518 | 2.61E-05 |
| Nsp14 | IL17RC | -0.518 | 2.69E-05 |
| Nsp14 | ADCY9 | -0.518 | 2.67E-05 |
| Nsp14 | TRAF2 | -0.518 | 2.62E-05 |
| Nsp14 | BCL3 | -0.518 | 2.67E-05 |
| Nsp14 | PPP1R3E | -0.518 | 2.63E-05 |
| Nsp16 | IRF3 | -0.518 | 2.59E-05 |
| Nsp16 | MAPK13 | -0.518 | 2.68E-05 |
| Nsp16 | CACNA2D2 | -0.518 | 2.59E-05 |
| Nsp16 | VEGFA | -0.518 | 2.61E-05 |
| S | AKT2 | -0.518 | 2.61E-05 |
| Orf3a | LEFTY2 | -0.518 | 2.65E-05 |
| Orf8 | ENDOG | -0.518 | 2.64E-05 |
| Orf8 | PREX1 | -0.518 | 2.63E-05 |
| Orf8 | RASGRP2 | -0.518 | 2.65E-05 |
| Nsp2 | PIAS4 | -0.519 | 2.55E-05 |
| Nsp3 | GSK3A | -0.519 | 2.51E-05 |
| Nsp3 | PRKCZ | -0.519 | 2.50E-05 |
| Nsp3 | CHRM1 | -0.519 | 2.57E-05 |
| Nsp3 | SMAD3 | -0.519 | 2.55E-05 |
| Nsp3 | CPT1B | -0.519 | 2.56E-05 |
| Nsp4 | HCK | -0.519 | 2.57E-05 |
| Nsp5 | FGF19 | -0.519 | 2.49E-05 |
| Nsp5 | RPS6KB2 | -0.519 | 2.49E-05 |
| Nsp8 | EXOC7 | -0.519 | 2.51E-05 |
| Nsp8 | FLNC | -0.519 | 2.58E-05 |
| Nsp12 | PLCG2 | -0.519 | 2.58E-05 |
| Nsp12 | MAP3K11 | -0.519 | 2.53E-05 |
| Nsp12 | IL17C | -0.519 | 2.56E-05 |
| Nsp13 | CXCR5 | -0.519 | 2.51E-05 |
| Nsp13 | GSK3A | -0.519 | 2.52E-05 |
| Nsp13 | NGFR | -0.519 | 2.59E-05 |
| Nsp13 | CACNG2 | -0.519 | 2.54E-05 |
| Nsp14 | IRF7 | -0.519 | 2.49E-05 |
| Nsp14 | RPTOR | -0.519 | 2.52E-05 |
| Nsp15 | TRAF2 | -0.519 | 2.56E-05 |
| Nsp15 | SESN2 | -0.519 | 2.50E-05 |
| Nsp15 | FGF3 | -0.519 | 2.49E-05 |
| Nsp16 | CAPN2 | -0.519 | 2.50E-05 |

| | | | |
|---|---|---|---|
| Nsp16 | SOCS3 | -0.519 | 2.57E-05 |
| S | IL34 | -0.519 | 2.49E-05 |
| S | MYC | -0.519 | 2.55E-05 |
| Orf8 | FZD7 | -0.519 | 2.55E-05 |
| Orf8 | DUSP5 | -0.519 | 2.52E-05 |
| Orf8 | PTPN6 | -0.519 | 2.49E-05 |
| Orf8 | WNT9B | -0.519 | 2.50E-05 |
| Nsp2 | EGLN1 | -0.52 | 2.48E-05 |
| Nsp3 | ISG15 | -0.52 | 2.45E-05 |
| Nsp3 | HSPA6 | -0.52 | 2.44E-05 |
| Nsp3 | CACNB1 | -0.52 | 2.47E-05 |
| Nsp4 | WNT9A | -0.52 | 2.42E-05 |
| Nsp4 | FGF19 | -0.52 | 2.47E-05 |
| Nsp4 | LIF | -0.52 | 2.39E-05 |
| Nsp4 | SOCS3 | -0.52 | 2.47E-05 |
| Nsp8 | RPTOR | -0.52 | 2.47E-05 |
| Nsp8 | PFKL | -0.52 | 2.39E-05 |
| Nsp10 | GREM2 | -0.52 | 2.48E-05 |
| Nsp12 | ID4 | -0.52 | 2.39E-05 |
| Nsp12 | CXCL3 | -0.52 | 2.40E-05 |
| Nsp12 | GRK5 | -0.52 | 2.48E-05 |
| Nsp14 | INSR | -0.52 | 2.40E-05 |
| Nsp14 | FOXO3 | -0.52 | 2.42E-05 |
| Nsp14 | ADCY3 | -0.52 | 2.44E-05 |
| Nsp14 | PLCB3 | -0.52 | 2.41E-05 |
| Nsp15 | TOLLIP | -0.52 | 2.39E-05 |
| Nsp15 | GNB2 | -0.52 | 2.43E-05 |
| Nsp16 | IRS1 | -0.52 | 2.46E-05 |
| Nsp16 | ADCY6 | -0.52 | 2.39E-05 |
| Nsp16 | THBS2 | -0.52 | 2.45E-05 |
| Orf8 | OSM | -0.52 | 2.42E-05 |
| Orf8 | MAP2K2 | -0.52 | 2.42E-05 |
| Nsp2 | TELO2 | -0.521 | 2.38E-05 |
| Nsp2 | DUSP6 | -0.521 | 2.32E-05 |
| Nsp4 | IRF7 | -0.521 | 2.36E-05 |
| Nsp4 | RRAS | -0.521 | 2.35E-05 |
| Nsp5 | SRF | -0.521 | 2.36E-05 |
| Nsp8 | TNFRSF13C | -0.521 | 2.28E-05 |
| Nsp10 | CARD14 | -0.521 | 2.33E-05 |
| Nsp12 | FGF6 | -0.521 | 2.31E-05 |
| Nsp12 | THBS2 | -0.521 | 2.36E-05 |
| Nsp13 | CAMKK2 | -0.521 | 2.31E-05 |
| Nsp13 | CD14 | -0.521 | 2.35E-05 |
| Nsp13 | LTB | -0.521 | 2.33E-05 |
| Nsp14 | IL27 | -0.521 | 2.34E-05 |
| Nsp14 | DUSP5 | -0.521 | 2.38E-05 |
| Nsp14 | WNT9A | -0.521 | 2.36E-05 |
| Nsp15 | MCL1 | -0.521 | 2.34E-05 |
| Nsp15 | LIF | -0.521 | 2.35E-05 |
| Nsp15 | THBS2 | -0.521 | 2.38E-05 |

| | | | |
|---|---|---|---|
| Nsp16 | PIDD1 | -0.521 | 2.31E-05 |
| Nsp16 | VAV2 | -0.521 | 2.31E-05 |
| S | FLNC | -0.521 | 2.29E-05 |
| Orf8 | CACNG1 | -0.521 | 2.29E-05 |
| Orf8 | WNT3A | -0.521 | 2.31E-05 |
| Orf8 | TNFRSF18 | -0.521 | 2.30E-05 |
| Orf8 | MMP9 | -0.521 | 2.35E-05 |
| Orf8 | CACNG5 | -0.521 | 2.38E-05 |
| Nsp2 | CACNG6 | -0.522 | 2.21E-05 |
| Nsp3 | PFKL | -0.522 | 2.27E-05 |
| Nsp4 | FMOD | -0.522 | 2.27E-05 |
| Nsp4 | CD70 | -0.522 | 2.19E-05 |
| Nsp4 | PRKACG | -0.522 | 2.20E-05 |
| Nsp5 | EPHA2 | -0.522 | 2.23E-05 |
| Nsp5 | ULK1 | -0.522 | 2.22E-05 |
| Nsp8 | PITX2 | -0.522 | 2.20E-05 |
| Nsp8 | RRAS | -0.522 | 2.21E-05 |
| Nsp8 | CD14 | -0.522 | 2.27E-05 |
| Nsp10 | GDF15 | -0.522 | 2.19E-05 |
| Nsp10 | CAPN2 | -0.522 | 2.21E-05 |
| Nsp12 | FGFR4 | -0.522 | 2.25E-05 |
| Nsp12 | GSK3A | -0.522 | 2.24E-05 |
| Nsp12 | PXN | -0.522 | 2.22E-05 |
| Nsp13 | GRK6 | -0.522 | 2.27E-05 |
| Nsp13 | DHX58 | -0.522 | 2.21E-05 |
| Nsp13 | CSH2 | -0.522 | 2.28E-05 |
| Nsp14 | DVL3 | -0.522 | 2.21E-05 |
| Nsp14 | TNFRSF8 | -0.522 | 2.19E-05 |
| Nsp14 | GYS1 | -0.522 | 2.25E-05 |
| Nsp14 | COL6A1 | -0.522 | 2.20E-05 |
| Nsp15 | YWHAG | -0.522 | 2.24E-05 |
| Nsp15 | CXCR3 | -0.522 | 2.24E-05 |
| Nsp15 | CX3CL1 | -0.522 | 2.19E-05 |
| Nsp16 | CACNA1C | -0.522 | 2.20E-05 |
| Nsp16 | PKN3 | -0.522 | 2.25E-05 |
| Nsp16 | ACVRL1 | -0.522 | 2.19E-05 |
| S | LMNA | -0.522 | 2.23E-05 |
| Orf8 | SMAD7 | -0.522 | 2.22E-05 |
| Orf8 | NOS3 | -0.522 | 2.19E-05 |
| Orf8 | TNFRSF6B | -0.522 | 2.20E-05 |
| Nsp2 | ID4 | -0.523 | 2.13E-05 |
| Nsp3 | MAPK13 | -0.523 | 2.12E-05 |
| Nsp3 | WNT7A | -0.523 | 2.13E-05 |
| Nsp4 | WNT9B | -0.523 | 2.14E-05 |
| Nsp5 | PTPN6 | -0.523 | 2.15E-05 |
| Nsp5 | PREX1 | -0.523 | 2.16E-05 |
| Nsp8 | WNT11 | -0.523 | 2.17E-05 |
| Nsp12 | FGF17 | -0.523 | 2.18E-05 |
| Nsp12 | RASGRF1 | -0.523 | 2.14E-05 |
| Nsp13 | MAPK12 | -0.523 | 2.18E-05 |

| | | | |
|---|---|---|---|
| Nsp13 | FLCN | -0.523 | 2.14E-05 |
| Nsp13 | IKBKE | -0.523 | 2.13E-05 |
| Nsp13 | PTPN7 | -0.523 | 2.14E-05 |
| Nsp14 | SRC | -0.523 | 2.17E-05 |
| Nsp14 | SLC7A5 | -0.523 | 2.10E-05 |
| Nsp15 | RPS6KA2 | -0.523 | 2.14E-05 |
| Nsp15 | WNT11 | -0.523 | 2.18E-05 |
| Nsp15 | CPT1C | -0.523 | 2.13E-05 |
| Nsp16 | NTRK1 | -0.523 | 2.18E-05 |
| Nsp16 | TELO2 | -0.523 | 2.18E-05 |
| Nsp16 | LCK | -0.523 | 2.18E-05 |
| S | IL17RC | -0.523 | 2.16E-05 |
| S | CARD10 | -0.523 | 2.17E-05 |
| S | BAD | -0.523 | 2.13E-05 |
| S | JAK3 | -0.523 | 2.17E-05 |
| S | CPT1C | -0.523 | 2.17E-05 |
| Orf8 | TRAF2 | -0.523 | 2.18E-05 |
| Orf8 | TELO2 | -0.523 | 2.10E-05 |
| Nsp2 | TNFRSF13C | -0.524 | 2.09E-05 |
| Nsp4 | MAPK13 | -0.524 | 2.06E-05 |
| Nsp4 | WNT4 | -0.524 | 2.02E-05 |
| Nsp5 | CREB3L1 | -0.524 | 2.06E-05 |
| Nsp8 | CREB3L1 | -0.524 | 2.03E-05 |
| Nsp8 | PPP5C | -0.524 | 2.04E-05 |
| Nsp10 | WNT7B | -0.524 | 2.06E-05 |
| Nsp10 | PIM1 | -0.524 | 2.08E-05 |
| Nsp12 | ICAM1 | -0.524 | 2.07E-05 |
| Nsp12 | TRIM25 | -0.524 | 2.04E-05 |
| Nsp13 | PRF1 | -0.524 | 2.05E-05 |
| Nsp13 | RRAS | -0.524 | 2.09E-05 |
| Nsp13 | PRKACA | -0.524 | 2.09E-05 |
| Nsp13 | GDF5 | -0.524 | 2.05E-05 |
| Nsp14 | TOLLIP | -0.524 | 2.04E-05 |
| Nsp14 | MYC | -0.524 | 2.01E-05 |
| Nsp14 | PXN | -0.524 | 2.07E-05 |
| Nsp15 | ID1 | -0.524 | 2.04E-05 |
| Nsp16 | RPS6KA2 | -0.524 | 2.08E-05 |
| Nsp16 | RASGRF1 | -0.524 | 2.02E-05 |
| S | ADCY9 | -0.524 | 2.07E-05 |
| S | PPP2R1A | -0.524 | 2.08E-05 |
| Orf8 | DAB2IP | -0.524 | 2.02E-05 |
| Orf8 | RASGRF1 | -0.524 | 2.08E-05 |
| Nsp2 | RPS6KA2 | -0.525 | 1.98E-05 |
| Nsp2 | RXRA | -0.525 | 1.98E-05 |
| Nsp3 | CXCR5 | -0.525 | 1.97E-05 |
| Nsp3 | GRK6 | -0.525 | 1.94E-05 |
| Nsp3 | PPP2R1A | -0.525 | 1.96E-05 |
| Nsp3 | OSM | -0.525 | 1.94E-05 |
| Nsp3 | TBX21 | -0.525 | 1.99E-05 |
| Nsp3 | BCAR1 | -0.525 | 1.96E-05 |

| | | | |
|---|---|---|---|
| Nsp4 | PPP1R3E | -0.525 | 1.99E-05 |
| Nsp4 | BAD | -0.525 | 1.99E-05 |
| Nsp4 | RASGRF1 | -0.525 | 1.98E-05 |
| Nsp6 | GDF7 | -0.525 | 2.01E-05 |
| Nsp8 | NGFR | -0.525 | 1.97E-05 |
| Nsp8 | PTPN6 | -0.525 | 2.00E-05 |
| Nsp10 | FZD10 | -0.525 | 1.98E-05 |
| Nsp12 | FLT3LG | -0.525 | 1.98E-05 |
| Nsp12 | CCR10 | -0.525 | 1.99E-05 |
| Nsp12 | GDF5 | -0.525 | 1.99E-05 |
| Nsp12 | VEGFA | -0.525 | 1.93E-05 |
| Nsp13 | WNT9B | -0.525 | 1.94E-05 |
| Nsp13 | MAP3K6 | -0.525 | 1.98E-05 |
| Nsp14 | CHAD | -0.525 | 2.00E-05 |
| Nsp14 | TBX21 | -0.525 | 1.93E-05 |
| Nsp16 | RELT | -0.525 | 2.00E-05 |
| Nsp16 | STAT5A | -0.525 | 1.94E-05 |
| Nsp16 | SMAD3 | -0.525 | 1.94E-05 |
| Nsp16 | TRIM25 | -0.525 | 1.94E-05 |
| S | MAP3K11 | -0.525 | 1.96E-05 |
| S | CD14 | -0.525 | 1.98E-05 |
| S | FOSL1 | -0.525 | 1.97E-05 |
| Orf8 | GNA12 | -0.525 | 1.95E-05 |
| Nsp2 | OSM | -0.526 | 1.89E-05 |
| Nsp2 | FZD5 | -0.526 | 1.88E-05 |
| Nsp3 | CARD10 | -0.526 | 1.89E-05 |
| Nsp3 | FLNC | -0.526 | 1.92E-05 |
| Nsp3 | THBS2 | -0.526 | 1.90E-05 |
| Nsp4 | CACNG4 | -0.526 | 1.91E-05 |
| Nsp4 | TRAF2 | -0.526 | 1.88E-05 |
| Nsp4 | MLST8 | -0.526 | 1.92E-05 |
| Nsp4 | TRIM25 | -0.526 | 1.92E-05 |
| Nsp5 | NFATC2 | -0.526 | 1.89E-05 |
| Nsp5 | RELB | -0.526 | 1.89E-05 |
| Nsp5 | PPP2R2C | -0.526 | 1.86E-05 |
| Nsp5 | WNT4 | -0.526 | 1.85E-05 |
| Nsp5 | WNT11 | -0.526 | 1.88E-05 |
| Nsp5 | TNFRSF1B | -0.526 | 1.84E-05 |
| Nsp8 | CARD10 | -0.526 | 1.91E-05 |
| Nsp9 | FZD10 | -0.526 | 1.89E-05 |
| Nsp12 | RGMA | -0.526 | 1.90E-05 |
| Nsp12 | CXCR3 | -0.526 | 1.88E-05 |
| Nsp13 | FGFR4 | -0.526 | 1.88E-05 |
| Nsp14 | MCL1 | -0.526 | 1.86E-05 |
| Nsp14 | MAP3K6 | -0.526 | 1.88E-05 |
| Nsp14 | JUND | -0.526 | 1.89E-05 |
| Nsp15 | CDC25B | -0.526 | 1.85E-05 |
| Nsp15 | PPP1R3E | -0.526 | 1.92E-05 |
| Nsp16 | NPRL2 | -0.526 | 1.88E-05 |
| Nsp16 | IGF2 | -0.526 | 1.87E-05 |

| | | | |
|---|---|---|---|
| S | RPS6KA2 | -0.526 | 1.91E-05 |
| S | MLST8 | -0.526 | 1.90E-05 |
| S | PRKACG | -0.526 | 1.87E-05 |
| Orf8 | CCR7 | -0.526 | 1.89E-05 |
| Orf8 | DUSP1 | -0.526 | 1.91E-05 |
| Orf8 | BAD | -0.526 | 1.89E-05 |
| Nsp2 | IL17D | -0.527 | 1.80E-05 |
| Nsp2 | FLT4 | -0.527 | 1.83E-05 |
| Nsp3 | PRF1 | -0.527 | 1.80E-05 |
| Nsp3 | RPS6KB2 | -0.527 | 1.77E-05 |
| Nsp4 | XCR1 | -0.527 | 1.83E-05 |
| Nsp5 | MAP3K6 | -0.527 | 1.81E-05 |
| Nsp8 | ACTG1 | -0.527 | 1.79E-05 |
| Nsp9 | GDF7 | -0.527 | 1.80E-05 |
| Nsp12 | INHBA | -0.527 | 1.82E-05 |
| Nsp14 | CACNG6 | -0.527 | 1.82E-05 |
| Nsp14 | DUSP1 | -0.527 | 1.77E-05 |
| Nsp14 | JUNB | -0.527 | 1.78E-05 |
| Nsp14 | CACNB1 | -0.527 | 1.81E-05 |
| Nsp15 | ISG15 | -0.527 | 1.79E-05 |
| Nsp15 | INSR | -0.527 | 1.84E-05 |
| Nsp15 | TNFRSF12A | -0.527 | 1.76E-05 |
| Nsp16 | CCR10 | -0.527 | 1.79E-05 |
| Nsp16 | EXOC7 | -0.527 | 1.77E-05 |
| S | FADD | -0.527 | 1.77E-05 |
| Orf3a | ENDOG | -0.527 | 1.83E-05 |
| Orf8 | CACNG7 | -0.527 | 1.77E-05 |
| Orf8 | JUND | -0.527 | 1.82E-05 |
| Nsp2 | LRP5 | -0.528 | 1.76E-05 |
| Nsp2 | WNT5B | -0.528 | 1.76E-05 |
| Nsp2 | IL17RA | -0.528 | 1.71E-05 |
| Nsp3 | INSR | -0.528 | 1.72E-05 |
| Nsp4 | TNXB | -0.528 | 1.74E-05 |
| Nsp4 | ACKR3 | -0.528 | 1.69E-05 |
| Nsp4 | PIN1 | -0.528 | 1.75E-05 |
| Nsp8 | ID1 | -0.528 | 1.76E-05 |
| Nsp8 | SMAD3 | -0.528 | 1.75E-05 |
| Nsp8 | PXN | -0.528 | 1.72E-05 |
| Nsp12 | CXCL14 | -0.528 | 1.74E-05 |
| Nsp12 | CTF1 | -0.528 | 1.76E-05 |
| Nsp13 | TAB1 | -0.528 | 1.72E-05 |
| Nsp13 | SMAD3 | -0.528 | 1.73E-05 |
| Nsp13 | PPP1R3E | -0.528 | 1.70E-05 |
| Nsp14 | ADCY1 | -0.528 | 1.70E-05 |
| Nsp14 | CCR10 | -0.528 | 1.69E-05 |
| Nsp14 | VAV2 | -0.528 | 1.72E-05 |
| Nsp14 | CALML5 | -0.528 | 1.72E-05 |
| Nsp14 | EGLN1 | -0.528 | 1.74E-05 |
| Nsp15 | PRF1 | -0.528 | 1.72E-05 |
| Nsp15 | PPP2R2C | -0.528 | 1.76E-05 |

| | | | |
|---|---|---|---|
| Nsp15 | RUNX1 | -0.528 | 1.74E-05 |
| S | CD70 | -0.528 | 1.75E-05 |
| S | MKNK2 | -0.528 | 1.76E-05 |
| Orf8 | CXCR3 | -0.528 | 1.69E-05 |
| Orf8 | PPP1R3D | -0.528 | 1.75E-05 |
| Orf8 | SH2B2 | -0.528 | 1.70E-05 |
| Orf8 | TICAM1 | -0.528 | 1.71E-05 |
| Nsp2 | DVL1 | -0.529 | 1.67E-05 |
| Nsp3 | PIK3R5 | -0.529 | 1.64E-05 |
| Nsp5 | RRAS | -0.529 | 1.66E-05 |
| Nsp5 | LEFTY2 | -0.529 | 1.68E-05 |
| Nsp5 | PKN1 | -0.529 | 1.68E-05 |
| Nsp5 | COL6A1 | -0.529 | 1.65E-05 |
| Nsp8 | MAPK13 | -0.529 | 1.65E-05 |
| Nsp8 | WDR24 | -0.529 | 1.62E-05 |
| Nsp12 | NFKBIB | -0.529 | 1.63E-05 |
| Nsp13 | PRKAB1 | -0.529 | 1.66E-05 |
| Nsp13 | ID4 | -0.529 | 1.64E-05 |
| Nsp13 | PPP5C | -0.529 | 1.62E-05 |
| Nsp13 | NODAL | -0.529 | 1.64E-05 |
| Nsp14 | CACNG7 | -0.529 | 1.66E-05 |
| Nsp14 | ICAM1 | -0.529 | 1.63E-05 |
| Nsp15 | EPHA2 | -0.529 | 1.62E-05 |
| Nsp15 | MAP3K11 | -0.529 | 1.64E-05 |
| Nsp15 | XCR1 | -0.529 | 1.68E-05 |
| Orf8 | FGF3 | -0.529 | 1.63E-05 |
| Orf8 | NCF1 | -0.529 | 1.63E-05 |
| Orf8 | CARD11 | -0.529 | 1.63E-05 |
| Nsp2 | BMP7 | -0.53 | 1.60E-05 |
| Nsp2 | GDF7 | -0.53 | 1.57E-05 |
| Nsp2 | TGFB1 | -0.53 | 1.62E-05 |
| Nsp3 | GNAI2 | -0.53 | 1.60E-05 |
| Nsp3 | TYK2 | -0.53 | 1.59E-05 |
| Nsp3 | FADD | -0.53 | 1.56E-05 |
| Nsp4 | TLR9 | -0.53 | 1.61E-05 |
| Nsp4 | ITPR3 | -0.53 | 1.61E-05 |
| Nsp4 | MAPK3 | -0.53 | 1.60E-05 |
| Nsp4 | PTPN6 | -0.53 | 1.56E-05 |
| Nsp4 | GNA12 | -0.53 | 1.61E-05 |
| Nsp5 | SRC | -0.53 | 1.55E-05 |
| Nsp5 | TRADD | -0.53 | 1.55E-05 |
| Nsp5 | AKT2 | -0.53 | 1.57E-05 |
| Nsp5 | TNFRSF4 | -0.53 | 1.57E-05 |
| Nsp8 | GSK3A | -0.53 | 1.55E-05 |
| Nsp8 | PRKACG | -0.53 | 1.59E-05 |
| Nsp10 | GATA3 | -0.53 | 1.60E-05 |
| Nsp12 | EPHA2 | -0.53 | 1.56E-05 |
| Nsp12 | HCK | -0.53 | 1.56E-05 |
| Nsp13 | CHAD | -0.53 | 1.55E-05 |
| Nsp13 | NFATC2 | -0.53 | 1.56E-05 |

| | | | |
|---|---|---|---|
| Nsp14 | MMP9 | -0.53 | 1.60E-05 |
| Nsp14 | DUSP6 | -0.53 | 1.61E-05 |
| Nsp16 | IRAK1 | -0.53 | 1.60E-05 |
| Nsp16 | SESN2 | -0.53 | 1.59E-05 |
| Nsp16 | PDGFRB | -0.53 | 1.57E-05 |
| S | CTSD | -0.53 | 1.60E-05 |
| S | FLT3LG | -0.53 | 1.57E-05 |
| S | PDGFB | -0.53 | 1.58E-05 |
| Orf3a | SHC2 | -0.53 | 1.59E-05 |
| Orf8 | RPS6KA4 | -0.53 | 1.58E-05 |
| Orf8 | EFNA3 | -0.53 | 1.55E-05 |
| Orf8 | ULK1 | -0.53 | 1.60E-05 |
| Orf8 | PRKACG | -0.53 | 1.57E-05 |
| Nsp2 | FGF3 | -0.531 | 1.48E-05 |
| Nsp3 | SRF | -0.531 | 1.51E-05 |
| Nsp4 | CACNA1C | -0.531 | 1.54E-05 |
| Nsp4 | MUC5B | -0.531 | 1.51E-05 |
| Nsp4 | DUSP6 | -0.531 | 1.51E-05 |
| Nsp5 | CDC25B | -0.531 | 1.53E-05 |
| Nsp5 | WNT3A | -0.531 | 1.50E-05 |
| Nsp8 | IL17RC | -0.531 | 1.54E-05 |
| Nsp9 | CACNG8 | -0.531 | 1.50E-05 |
| Nsp12 | PLCB2 | -0.531 | 1.49E-05 |
| Nsp12 | MUC5B | -0.531 | 1.50E-05 |
| Nsp12 | LCK | -0.531 | 1.51E-05 |
| Nsp13 | NTRK1 | -0.531 | 1.48E-05 |
| Nsp14 | ITGB4 | -0.531 | 1.53E-05 |
| Nsp14 | CDKN2B | -0.531 | 1.55E-05 |
| Nsp14 | PKN1 | -0.531 | 1.51E-05 |
| Nsp14 | ULK1 | -0.531 | 1.55E-05 |
| Nsp15 | WNT9B | -0.531 | 1.49E-05 |
| Nsp15 | TBX21 | -0.531 | 1.52E-05 |
| Nsp16 | ADCY9 | -0.531 | 1.49E-05 |
| Nsp16 | INSR | -0.531 | 1.50E-05 |
| Nsp16 | PRKACG | -0.531 | 1.53E-05 |
| S | TNXB | -0.531 | 1.51E-05 |
| Orf8 | FOSB | -0.531 | 1.52E-05 |
| Nsp2 | NCF1 | -0.532 | 1.45E-05 |
| Nsp2 | BCL2 | -0.532 | 1.43E-05 |
| Nsp3 | CAMKK2 | -0.532 | 1.48E-05 |
| Nsp3 | ATP6V1B1 | -0.532 | 1.42E-05 |
| Nsp3 | DUSP3 | -0.532 | 1.44E-05 |
| Nsp3 | ID4 | -0.532 | 1.48E-05 |
| Nsp4 | IL27 | -0.532 | 1.42E-05 |
| Nsp4 | FGF18 | -0.532 | 1.46E-05 |
| Nsp4 | PLCB3 | -0.532 | 1.47E-05 |
| Nsp5 | ITPR3 | -0.532 | 1.42E-05 |
| Nsp8 | TNXB | -0.532 | 1.46E-05 |
| Nsp8 | CALML5 | -0.532 | 1.46E-05 |
| Nsp10 | GDF7 | -0.532 | 1.42E-05 |

| | | | |
|---|---|---|---|
| Nsp12 | SRF | -0.532 | 1.44E-05 |
| Nsp13 | EXOC7 | -0.532 | 1.44E-05 |
| Nsp15 | TNFRSF8 | -0.532 | 1.43E-05 |
| Nsp15 | NFATC2 | -0.532 | 1.47E-05 |
| Nsp16 | MAP4K2 | -0.532 | 1.46E-05 |
| S | WNT9B | -0.532 | 1.43E-05 |
| Orf8 | RELB | -0.532 | 1.46E-05 |
| Orf8 | PPP2R2C | -0.532 | 1.43E-05 |
| Nsp3 | AKT1 | -0.533 | 1.40E-05 |
| Nsp3 | MUC5B | -0.533 | 1.37E-05 |
| Nsp3 | WNT9B | -0.533 | 1.42E-05 |
| Nsp4 | TRADD | -0.533 | 1.40E-05 |
| Nsp4 | PDGFB | -0.533 | 1.40E-05 |
| Nsp5 | CXCR3 | -0.533 | 1.41E-05 |
| Nsp8 | ISG15 | -0.533 | 1.38E-05 |
| Nsp10 | INHBA | -0.533 | 1.39E-05 |
| Nsp12 | JUND | -0.533 | 1.36E-05 |
| Nsp12 | CACNG5 | -0.533 | 1.37E-05 |
| Nsp13 | CARD10 | -0.533 | 1.37E-05 |
| Nsp14 | DUSP4 | -0.533 | 1.41E-05 |
| Nsp14 | PREX1 | -0.533 | 1.37E-05 |
| Nsp14 | TNFRSF4 | -0.533 | 1.41E-05 |
| Nsp15 | TRADD | -0.533 | 1.41E-05 |
| Nsp15 | HCK | -0.533 | 1.37E-05 |
| Nsp15 | CTSD | -0.533 | 1.38E-05 |
| Nsp15 | MAPK3 | -0.533 | 1.39E-05 |
| Nsp15 | MUC5B | -0.533 | 1.41E-05 |
| Nsp15 | TSC2 | -0.533 | 1.38E-05 |
| Nsp15 | PXN | -0.533 | 1.37E-05 |
| Nsp16 | PITX2 | -0.533 | 1.39E-05 |
| Nsp16 | CXCR5 | -0.533 | 1.37E-05 |
| Nsp16 | ICAM1 | -0.533 | 1.42E-05 |
| Orf8 | PRR5 | -0.533 | 1.39E-05 |
| Orf8 | LIF | -0.533 | 1.41E-05 |
| Nsp2 | GREM2 | -0.534 | 1.32E-05 |
| Nsp2 | HRK | -0.534 | 1.31E-05 |
| Nsp3 | PITX2 | -0.534 | 1.33E-05 |
| Nsp3 | ARAF | -0.534 | 1.32E-05 |
| Nsp3 | SOCS3 | -0.534 | 1.34E-05 |
| Nsp4 | GADD45B | -0.534 | 1.31E-05 |
| Nsp5 | FGF18 | -0.534 | 1.34E-05 |
| Nsp5 | MAPK4 | -0.534 | 1.30E-05 |
| Nsp5 | PXN | -0.534 | 1.30E-05 |
| Nsp8 | PREX1 | -0.534 | 1.30E-05 |
| Nsp12 | FGF18 | -0.534 | 1.33E-05 |
| Nsp12 | TOLLIP | -0.534 | 1.33E-05 |
| Nsp12 | PLCG1 | -0.534 | 1.31E-05 |
| Nsp12 | VHL | -0.534 | 1.35E-05 |
| Nsp13 | CXCR3 | -0.534 | 1.31E-05 |
| Nsp13 | PLCB2 | -0.534 | 1.30E-05 |

| | | | |
|---|---|---|---|
| Nsp13 | EGLN1 | -0.534 | 1.32E-05 |
| Nsp14 | CTSD | -0.534 | 1.31E-05 |
| Nsp14 | GRK1 | -0.534 | 1.34E-05 |
| Nsp14 | RARA | -0.534 | 1.33E-05 |
| Nsp15 | TLR9 | -0.534 | 1.30E-05 |
| Nsp15 | CACNG4 | -0.534 | 1.34E-05 |
| Nsp15 | CAPN2 | -0.534 | 1.31E-05 |
| Nsp15 | FLCN | -0.534 | 1.34E-05 |
| Nsp15 | OSM | -0.534 | 1.32E-05 |
| Nsp15 | MAP3K6 | -0.534 | 1.33E-05 |
| Nsp15 | TRIM25 | -0.534 | 1.33E-05 |
| Nsp15 | PLCB3 | -0.534 | 1.33E-05 |
| Nsp16 | CHRD | -0.534 | 1.35E-05 |
| Nsp16 | ISG15 | -0.534 | 1.33E-05 |
| Nsp16 | CHRM1 | -0.534 | 1.35E-05 |
| Nsp16 | TRAF2 | -0.534 | 1.30E-05 |
| S | TELO2 | -0.534 | 1.35E-05 |
| Orf3a | NOG | -0.534 | 1.31E-05 |
| Orf8 | TBKBP1 | -0.534 | 1.35E-05 |
| Orf8 | CDC25B | -0.534 | 1.32E-05 |
| Nsp3 | NR4A1 | -0.535 | 1.28E-05 |
| Nsp3 | MCL1 | -0.535 | 1.27E-05 |
| Nsp3 | ICAM1 | -0.535 | 1.28E-05 |
| Nsp3 | TRIM25 | -0.535 | 1.28E-05 |
| Nsp4 | YWHAG | -0.535 | 1.27E-05 |
| Nsp4 | BCL3 | -0.535 | 1.27E-05 |
| Nsp4 | MAP3K6 | -0.535 | 1.27E-05 |
| Nsp4 | VHL | -0.535 | 1.27E-05 |
| Nsp5 | TELO2 | -0.535 | 1.29E-05 |
| Nsp8 | MAP3K6 | -0.535 | 1.27E-05 |
| Nsp8 | ADCY4 | -0.535 | 1.25E-05 |
| Nsp8 | SOCS3 | -0.535 | 1.24E-05 |
| Nsp10 | SHC2 | -0.535 | 1.28E-05 |
| Nsp12 | TNXB | -0.535 | 1.28E-05 |
| Nsp12 | ELAVL1 | -0.535 | 1.29E-05 |
| Nsp12 | CDKN2B | -0.535 | 1.28E-05 |
| Nsp14 | GDF11 | -0.535 | 1.24E-05 |
| Nsp14 | NCF1 | -0.535 | 1.26E-05 |
| Nsp15 | PITX2 | -0.535 | 1.25E-05 |
| Nsp15 | AKT2 | -0.535 | 1.28E-05 |
| Nsp15 | ID4 | -0.535 | 1.26E-05 |
| Nsp15 | SH2B2 | -0.535 | 1.26E-05 |
| S | GYS1 | -0.535 | 1.29E-05 |
| Orf8 | PIAS4 | -0.535 | 1.25E-05 |
| Nsp2 | CARD14 | -0.536 | 1.24E-05 |
| Nsp2 | JUNB | -0.536 | 1.22E-05 |
| Nsp2 | FZD1 | -0.536 | 1.23E-05 |
| Nsp3 | ID1 | -0.536 | 1.20E-05 |
| Nsp3 | IL17C | -0.536 | 1.19E-05 |
| Nsp4 | TICAM1 | -0.536 | 1.23E-05 |

| | | | |
|---|---|---|---|
| Nsp5 | FOSB | -0.536 | 1.24E-05 |
| Nsp5 | ITGB4 | -0.536 | 1.21E-05 |
| Nsp5 | FLT3LG | -0.536 | 1.22E-05 |
| Nsp5 | CD70 | -0.536 | 1.21E-05 |
| Nsp8 | HCK | -0.536 | 1.22E-05 |
| Nsp8 | ADCY9 | -0.536 | 1.19E-05 |
| Nsp8 | RAC2 | -0.536 | 1.20E-05 |
| Nsp8 | TELO2 | -0.536 | 1.19E-05 |
| Nsp12 | LAMC3 | -0.536 | 1.21E-05 |
| Nsp12 | TAB1 | -0.536 | 1.22E-05 |
| Nsp13 | MAPK13 | -0.536 | 1.24E-05 |
| Nsp13 | HSPA6 | -0.536 | 1.20E-05 |
| Nsp14 | FZD2 | -0.536 | 1.22E-05 |
| Nsp14 | MAPK8IP3 | -0.536 | 1.23E-05 |
| Nsp14 | SH2B2 | -0.536 | 1.21E-05 |
| Nsp15 | PIK3R5 | -0.536 | 1.21E-05 |
| Nsp16 | ID1 | -0.536 | 1.23E-05 |
| Nsp16 | ID4 | -0.536 | 1.20E-05 |
| Nsp16 | WNT5A | -0.536 | 1.21E-05 |
| Orf8 | PRKAR1B | -0.536 | 1.23E-05 |
| Orf8 | ADCY9 | -0.536 | 1.21E-05 |
| Nsp3 | PLCB2 | -0.537 | 1.17E-05 |
| Nsp3 | MAP3K11 | -0.537 | 1.19E-05 |
| Nsp3 | NGFR | -0.537 | 1.14E-05 |
| Nsp3 | RPS6KA2 | -0.537 | 1.17E-05 |
| Nsp3 | FGF19 | -0.537 | 1.16E-05 |
| Nsp4 | TOLLIP | -0.537 | 1.18E-05 |
| Nsp4 | PREX1 | -0.537 | 1.14E-05 |
| Nsp4 | PKN1 | -0.537 | 1.18E-05 |
| Nsp5 | PITX2 | -0.537 | 1.16E-05 |
| Nsp5 | TOLLIP | -0.537 | 1.17E-05 |
| Nsp5 | ID4 | -0.537 | 1.19E-05 |
| Nsp5 | VHL | -0.537 | 1.19E-05 |
| Nsp8 | CHAD | -0.537 | 1.15E-05 |
| Nsp8 | GYS1 | -0.537 | 1.14E-05 |
| Nsp8 | FGF17 | -0.537 | 1.15E-05 |
| Nsp10 | BMP6 | -0.537 | 1.16E-05 |
| Nsp12 | YWHAG | -0.537 | 1.18E-05 |
| Nsp12 | MAPK15 | -0.537 | 1.16E-05 |
| Nsp12 | MAPK13 | -0.537 | 1.17E-05 |
| Nsp13 | ATP6V1B1 | -0.537 | 1.15E-05 |
| Nsp13 | TRIM25 | -0.537 | 1.15E-05 |
| Nsp14 | MAPK11 | -0.537 | 1.16E-05 |
| Nsp14 | PTPN6 | -0.537 | 1.17E-05 |
| Nsp14 | LAMA5 | -0.537 | 1.18E-05 |
| Nsp15 | ADCY1 | -0.537 | 1.16E-05 |
| Nsp15 | NODAL | -0.537 | 1.16E-05 |
| Nsp15 | IL17C | -0.537 | 1.15E-05 |
| Nsp16 | PPP5C | -0.537 | 1.15E-05 |
| Nsp16 | FOSL1 | -0.537 | 1.16E-05 |

| | | | |
|---|---|---|---|
| S | FOSB | -0.537 | 1.16E-05 |
| S | MAPK13 | -0.537 | 1.14E-05 |
| S | WNT11 | -0.537 | 1.14E-05 |
| S | WNT5A | -0.537 | 1.14E-05 |
| Nsp3 | HCK | -0.538 | 1.14E-05 |
| Nsp3 | VAV2 | -0.538 | 1.13E-05 |
| Nsp3 | CD14 | -0.538 | 1.09E-05 |
| Nsp3 | PPP5C | -0.538 | 1.13E-05 |
| Nsp4 | EDAR | -0.538 | 1.12E-05 |
| Nsp4 | INSR | -0.538 | 1.12E-05 |
| Nsp4 | VEGFA | -0.538 | 1.09E-05 |
| Nsp5 | ELAVL1 | -0.538 | 1.09E-05 |
| Nsp5 | RPS6KA2 | -0.538 | 1.14E-05 |
| Nsp8 | TLR9 | -0.538 | 1.11E-05 |
| Nsp8 | WNT9B | -0.538 | 1.10E-05 |
| Nsp8 | GRK1 | -0.538 | 1.09E-05 |
| Nsp8 | MKNK2 | -0.538 | 1.11E-05 |
| Nsp9 | CEBPB | -0.538 | 1.10E-05 |
| Nsp12 | FGF19 | -0.538 | 1.12E-05 |
| Nsp12 | MLST8 | -0.538 | 1.10E-05 |
| Nsp13 | TNXB | -0.538 | 1.09E-05 |
| Nsp13 | PIK3R5 | -0.538 | 1.09E-05 |
| Nsp14 | DUSP9 | -0.538 | 1.09E-05 |
| Nsp14 | BMP8B | -0.538 | 1.11E-05 |
| Nsp14 | POMC | -0.538 | 1.12E-05 |
| Nsp15 | WNT3A | -0.538 | 1.10E-05 |
| Nsp15 | GYS1 | -0.538 | 1.10E-05 |
| Nsp15 | WNT5A | -0.538 | 1.09E-05 |
| Nsp16 | CCR7 | -0.538 | 1.09E-05 |
| Nsp16 | GREM1 | -0.538 | 1.12E-05 |
| Nsp16 | RELA | -0.538 | 1.11E-05 |
| S | MCL1 | -0.538 | 1.11E-05 |
| Orf8 | WNT7B | -0.538 | 1.10E-05 |
| Orf8 | DVL1 | -0.538 | 1.11E-05 |
| Orf8 | ICAM1 | -0.538 | 1.10E-05 |
| Orf8 | GADD45B | -0.538 | 1.10E-05 |
| Orf8 | NOG | -0.538 | 1.10E-05 |
| Nsp2 | LEFTY1 | -0.539 | 1.07E-05 |
| Nsp3 | TRAF2 | -0.539 | 1.08E-05 |
| Nsp4 | VAV2 | -0.539 | 1.08E-05 |
| Nsp4 | PRKCG | -0.539 | 1.07E-05 |
| Nsp4 | CPT1C | -0.539 | 1.06E-05 |
| Nsp4 | PTPN7 | -0.539 | 1.05E-05 |
| Nsp5 | PRKAR1B | -0.539 | 1.05E-05 |
| Nsp5 | TLR9 | -0.539 | 1.07E-05 |
| Nsp5 | MAPK11 | -0.539 | 1.05E-05 |
| Nsp5 | FLT4 | -0.539 | 1.04E-05 |
| Nsp5 | MLST8 | -0.539 | 1.08E-05 |
| Nsp5 | WDR24 | -0.539 | 1.05E-05 |
| Nsp8 | CTSD | -0.539 | 1.06E-05 |

| | | | |
|---|---|---|---|
| Nsp8 | TNFRSF4 | -0.539 | 1.08E-05 |
| Nsp12 | CARD10 | -0.539 | 1.06E-05 |
| Nsp12 | PPP2R2C | -0.539 | 1.05E-05 |
| Nsp13 | WNT3A | -0.539 | 1.06E-05 |
| Nsp13 | ID1 | -0.539 | 1.08E-05 |
| Nsp14 | CACNA1G | -0.539 | 1.06E-05 |
| Nsp15 | DVL3 | -0.539 | 1.08E-05 |
| Nsp15 | NFKBIB | -0.539 | 1.09E-05 |
| Nsp15 | PLCG1 | -0.539 | 1.04E-05 |
| Nsp15 | PKN1 | -0.539 | 1.06E-05 |
| Nsp16 | ITGA3 | -0.539 | 1.08E-05 |
| Nsp16 | ACVR2B | -0.539 | 1.06E-05 |
| Nsp16 | LAMC3 | -0.539 | 1.05E-05 |
| S | RRAS | -0.539 | 1.04E-05 |
| S | PREX1 | -0.539 | 1.06E-05 |
| Orf8 | CACNA1C | -0.539 | 1.06E-05 |
| Orf8 | DUSP9 | -0.539 | 1.06E-05 |
| Orf8 | BCL2 | -0.539 | 1.08E-05 |
| Nsp3 | FGFR4 | -0.54 | 1.03E-05 |
| Nsp3 | RRAS | -0.54 | 1.02E-05 |
| Nsp3 | TELO2 | -0.54 | 9.99E-06 |
| Nsp3 | PHLPP1 | -0.54 | 1.02E-05 |
| Nsp3 | TICAM1 | -0.54 | 1.00E-05 |
| Nsp4 | ADCY1 | -0.54 | 1.02E-05 |
| Nsp4 | CCR10 | -0.54 | 9.99E-06 |
| Nsp4 | TELO2 | -0.54 | 1.00E-05 |
| Nsp4 | TBX21 | -0.54 | 1.03E-05 |
| Nsp4 | WNT10A | -0.54 | 9.97E-06 |
| Nsp4 | CACNG5 | -0.54 | 1.02E-05 |
| Nsp5 | NFKBIB | -0.54 | 1.01E-05 |
| Nsp5 | LEFTY1 | -0.54 | 1.01E-05 |
| Nsp5 | WNT10A | -0.54 | 1.03E-05 |
| Nsp8 | TRAF2 | -0.54 | 1.03E-05 |
| Nsp8 | FGF19 | -0.54 | 9.99E-06 |
| Nsp8 | PLCB3 | -0.54 | 1.01E-05 |
| Nsp12 | ACKR3 | -0.54 | 1.00E-05 |
| Nsp12 | GADD45B | -0.54 | 1.03E-05 |
| Nsp12 | FOXO3 | -0.54 | 9.95E-06 |
| Nsp13 | MAPK15 | -0.54 | 1.03E-05 |
| Nsp13 | TRAF2 | -0.54 | 1.02E-05 |
| Nsp14 | FLT4 | -0.54 | 1.04E-05 |
| Nsp14 | PIK3CD | -0.54 | 1.02E-05 |
| Nsp14 | CCND1 | -0.54 | 9.96E-06 |
| Nsp14 | PIN1 | -0.54 | 9.94E-06 |
| Nsp15 | ADCY9 | -0.54 | 1.03E-05 |
| Nsp15 | TAB1 | -0.54 | 1.01E-05 |
| Nsp15 | ACKR3 | -0.54 | 1.03E-05 |
| Nsp15 | CCND1 | -0.54 | 1.03E-05 |
| Nsp15 | EGLN1 | -0.54 | 1.00E-05 |
| Nsp16 | TBX21 | -0.54 | 1.01E-05 |

| | | | |
|---|---|---|---|
| S | PITX2 | -0.54 | 1.03E-05 |
| S | GNA12 | -0.54 | 9.99E-06 |
| S | CALML5 | -0.54 | 1.02E-05 |
| S | BCAR1 | -0.54 | 1.04E-05 |
| Orf8 | YWHAG | -0.54 | 1.00E-05 |
| Nsp2 | GDF6 | -0.541 | 9.81E-06 |
| Nsp3 | IL34 | -0.541 | 9.85E-06 |
| Nsp3 | LAMC3 | -0.541 | 9.59E-06 |
| Nsp4 | MCL1 | -0.541 | 9.82E-06 |
| Nsp5 | DAB2IP | -0.541 | 9.71E-06 |
| Nsp5 | CACNA1G | -0.541 | 9.90E-06 |
| Nsp5 | ICAM1 | -0.541 | 9.89E-06 |
| Nsp8 | FOSB | -0.541 | 9.49E-06 |
| Nsp12 | IL27 | -0.541 | 9.84E-06 |
| Nsp12 | ITPR3 | -0.541 | 9.78E-06 |
| Nsp12 | WNT9B | -0.541 | 9.89E-06 |
| Nsp13 | IGF1R | -0.541 | 9.87E-06 |
| Nsp13 | GNB2 | -0.541 | 9.91E-06 |
| Nsp13 | TICAM1 | -0.541 | 9.82E-06 |
| Nsp13 | THBS2 | -0.541 | 9.86E-06 |
| Nsp15 | SRC | -0.541 | 9.64E-06 |
| Nsp15 | PTPN6 | -0.541 | 9.70E-06 |
| Nsp16 | PPP2R5D | -0.541 | 9.72E-06 |
| Nsp16 | LMNA | -0.541 | 9.67E-06 |
| Orf8 | IL17D | -0.541 | 9.56E-06 |
| Orf8 | RGMA | -0.541 | 9.71E-06 |
| Orf8 | FLT4 | -0.541 | 9.52E-06 |
| Orf8 | FGF19 | -0.541 | 9.61E-06 |
| Nsp2 | MAPK8IP2 | -0.542 | 9.43E-06 |
| Nsp3 | NFKBIB | -0.542 | 9.35E-06 |
| Nsp3 | WNT5A | -0.542 | 9.11E-06 |
| Nsp4 | CTSD | -0.542 | 9.44E-06 |
| Nsp4 | FOXO3 | -0.542 | 9.24E-06 |
| Nsp8 | EPHA2 | -0.542 | 9.25E-06 |
| Nsp8 | EPOR | -0.542 | 9.07E-06 |
| Nsp8 | CACNA1G | -0.542 | 9.28E-06 |
| Nsp12 | PPP2R1A | -0.542 | 9.45E-06 |
| Nsp12 | TELO2 | -0.542 | 9.11E-06 |
| Nsp12 | PDGFB | -0.542 | 9.09E-06 |
| Nsp13 | CDKN2B | -0.542 | 9.10E-06 |
| Nsp13 | TSC2 | -0.542 | 9.22E-06 |
| Nsp14 | YWHAG | -0.542 | 9.09E-06 |
| Nsp14 | CACNA1A | -0.542 | 9.23E-06 |
| Nsp14 | PTPN7 | -0.542 | 9.30E-06 |
| Nsp15 | ATP6V1B1 | -0.542 | 9.39E-06 |
| Nsp15 | DUSP4 | -0.542 | 9.28E-06 |
| Nsp15 | MAPK11 | -0.542 | 9.32E-06 |
| Nsp16 | TOLLIP | -0.542 | 9.35E-06 |
| Nsp16 | NGFR | -0.542 | 9.25E-06 |
| S | PFKFB3 | -0.542 | 9.30E-06 |

| | | | |
|---|---|---|---|
| Orf8 | LEFTY1 | -0.542 | 9.39E-06 |
| Orf8 | ADCY5 | -0.542 | 9.26E-06 |
| Nsp2 | FZD8 | -0.543 | 8.81E-06 |
| Nsp2 | LMNB2 | -0.543 | 8.97E-06 |
| Nsp3 | GNB2 | -0.543 | 8.99E-06 |
| Nsp3 | CALML5 | -0.543 | 8.66E-06 |
| Nsp3 | TNFRSF13C | -0.543 | 8.76E-06 |
| Nsp4 | ID4 | -0.543 | 8.67E-06 |
| Nsp4 | WNT11 | -0.543 | 8.78E-06 |
| Nsp4 | ADCY4 | -0.543 | 8.69E-06 |
| Nsp4 | MKNK2 | -0.543 | 9.04E-06 |
| Nsp5 | WNT9A | -0.543 | 8.75E-06 |
| Nsp8 | WNT7A | -0.543 | 8.82E-06 |
| Nsp8 | CACNA1B | -0.543 | 8.74E-06 |
| Nsp12 | CTSD | -0.543 | 8.92E-06 |
| Nsp12 | MAPK11 | -0.543 | 8.85E-06 |
| Nsp12 | ULK1 | -0.543 | 8.68E-06 |
| Nsp13 | LIF | -0.543 | 8.79E-06 |
| Nsp13 | BAD | -0.543 | 8.82E-06 |
| Nsp14 | PPP2R2C | -0.543 | 8.74E-06 |
| Nsp14 | CACNA1B | -0.543 | 8.75E-06 |
| Nsp15 | ICAM1 | -0.543 | 8.95E-06 |
| Nsp16 | PRF1 | -0.543 | 8.87E-06 |
| Nsp16 | FOSB | -0.543 | 8.88E-06 |
| Nsp16 | PLCB2 | -0.543 | 9.03E-06 |
| Nsp16 | MUC5B | -0.543 | 8.81E-06 |
| Nsp16 | CARD10 | -0.543 | 8.90E-06 |
| S | CACNG4 | -0.543 | 8.95E-06 |
| Orf8 | WNT6 | -0.543 | 8.69E-06 |
| Nsp2 | LEFTY2 | -0.544 | 8.33E-06 |
| Nsp3 | TAB1 | -0.544 | 8.30E-06 |
| Nsp3 | JAK3 | -0.544 | 8.37E-06 |
| Nsp4 | FLT3LG | -0.544 | 8.58E-06 |
| Nsp8 | CXCR5 | -0.544 | 8.42E-06 |
| Nsp8 | CACNG1 | -0.544 | 8.63E-06 |
| Nsp8 | PFKFB3 | -0.544 | 8.28E-06 |
| Nsp12 | TSC2 | -0.544 | 8.34E-06 |
| Nsp13 | EPHA2 | -0.544 | 8.65E-06 |
| Nsp13 | GYS1 | -0.544 | 8.65E-06 |
| Nsp13 | RPS6KA2 | -0.544 | 8.31E-06 |
| Nsp14 | GADD45B | -0.544 | 8.30E-06 |
| Nsp14 | WNT1 | -0.544 | 8.61E-06 |
| Nsp14 | ADCY5 | -0.544 | 8.28E-06 |
| Nsp15 | TNFRSF4 | -0.544 | 8.40E-06 |
| Nsp16 | GNB2 | -0.544 | 8.28E-06 |
| Nsp16 | SRF | -0.544 | 8.56E-06 |
| Nsp16 | IL17C | -0.544 | 8.59E-06 |
| S | RPTOR | -0.544 | 8.47E-06 |
| S | GRK1 | -0.544 | 8.55E-06 |
| S | WNT7A | -0.544 | 8.49E-06 |

| | | | |
|---|---|---|---|
| S | XCR1 | -0.544 | 8.48E-06 |
| Orf8 | CACNA1H | -0.544 | 8.37E-06 |
| Orf8 | GDF10 | -0.544 | 8.61E-06 |
| Orf8 | RXRA | -0.544 | 8.64E-06 |
| Orf8 | AMH | -0.544 | 8.35E-06 |
| Orf8 | IL17RA | -0.544 | 8.39E-06 |
| Nsp2 | MCL1 | -0.545 | 8.21E-06 |
| Nsp3 | FLCN | -0.545 | 8.07E-06 |
| Nsp3 | WNT4 | -0.545 | 7.90E-06 |
| Nsp3 | MAP3K6 | -0.545 | 8.21E-06 |
| Nsp4 | WNT3A | -0.545 | 8.04E-06 |
| Nsp5 | CARD14 | -0.545 | 8.14E-06 |
| Nsp5 | SMAD3 | -0.545 | 8.16E-06 |
| Nsp8 | ITPR3 | -0.545 | 8.21E-06 |
| Nsp8 | CACNG4 | -0.545 | 8.12E-06 |
| Nsp12 | TRAF2 | -0.545 | 7.94E-06 |
| Nsp12 | NODAL | -0.545 | 8.09E-06 |
| Nsp12 | SH2B2 | -0.545 | 8.03E-06 |
| Nsp13 | WDR24 | -0.545 | 7.98E-06 |
| Nsp14 | WNT3A | -0.545 | 8.06E-06 |
| Nsp14 | MAPK4 | -0.545 | 8.25E-06 |
| Nsp14 | ADCY7 | -0.545 | 8.04E-06 |
| Nsp15 | IRF7 | -0.545 | 7.96E-06 |
| Nsp15 | FADD | -0.545 | 8.23E-06 |
| Nsp16 | DUSP3 | -0.545 | 8.25E-06 |
| S | ISG15 | -0.545 | 7.98E-06 |
| Orf8 | GRK2 | -0.545 | 8.20E-06 |
| Orf8 | FZD5 | -0.545 | 8.14E-06 |
| Orf8 | RARA | -0.545 | 8.20E-06 |
| Nsp2 | RPS6KA4 | -0.546 | 7.58E-06 |
| Nsp3 | LIF | -0.546 | 7.89E-06 |
| Nsp3 | PRKACG | -0.546 | 7.63E-06 |
| Nsp4 | ULK1 | -0.546 | 7.69E-06 |
| Nsp5 | DUSP5 | -0.546 | 7.86E-06 |
| Nsp5 | CCR10 | -0.546 | 7.79E-06 |
| Nsp5 | IL17RA | -0.546 | 7.55E-06 |
| Nsp8 | CDKN2B | -0.546 | 7.84E-06 |
| Nsp8 | ID4 | -0.546 | 7.72E-06 |
| Nsp12 | FMOD | -0.546 | 7.85E-06 |
| Nsp12 | CPT1C | -0.546 | 7.63E-06 |
| Nsp13 | DVL3 | -0.546 | 7.88E-06 |
| Nsp13 | DUSP2 | -0.546 | 7.72E-06 |
| Nsp13 | CTF1 | -0.546 | 7.64E-06 |
| Nsp14 | IL17D | -0.546 | 7.79E-06 |
| Nsp14 | HSPA1B | -0.546 | 7.88E-06 |
| Nsp14 | CARD11 | -0.546 | 7.73E-06 |
| Nsp15 | GSK3A | -0.546 | 7.81E-06 |
| Nsp15 | GNAI2 | -0.546 | 7.71E-06 |
| Nsp15 | MKNK2 | -0.546 | 7.73E-06 |
| Nsp15 | ULK1 | -0.546 | 7.62E-06 |

| | | | |
|---|---|---|---|
| Nsp16 | CAMKK2 | -0.546 | 7.87E-06 |
| Nsp16 | PTK2B | -0.546 | 7.64E-06 |
| Nsp16 | ADCY1 | -0.546 | 7.55E-06 |
| Nsp16 | WNT9B | -0.546 | 7.80E-06 |
| S | SRF | -0.546 | 7.67E-06 |
| S | PLCB3 | -0.546 | 7.82E-06 |
| Orf8 | IL27 | -0.546 | 7.69E-06 |
| Orf8 | LEFTY2 | -0.546 | 7.55E-06 |
| Orf8 | TGFB1 | -0.546 | 7.59E-06 |
| Orf8 | CACNA1A | -0.546 | 7.75E-06 |
| Nsp2 | GDF10 | -0.547 | 7.23E-06 |
| Nsp3 | DVL3 | -0.547 | 7.52E-06 |
| Nsp3 | CSH1 | -0.547 | 7.39E-06 |
| Nsp3 | PLCG1 | -0.547 | 7.33E-06 |
| Nsp4 | DUSP1 | -0.547 | 7.47E-06 |
| Nsp4 | CREB3L1 | -0.547 | 7.23E-06 |
| Nsp5 | DVL3 | -0.547 | 7.22E-06 |
| Nsp5 | CACNA1A | -0.547 | 7.25E-06 |
| Nsp5 | XCR1 | -0.547 | 7.29E-06 |
| Nsp5 | SH2B2 | -0.547 | 7.32E-06 |
| Nsp5 | RUNX1 | -0.547 | 7.33E-06 |
| Nsp5 | CTF1 | -0.547 | 7.38E-06 |
| Nsp8 | RPS6KA2 | -0.547 | 7.29E-06 |
| Nsp9 | FGF4 | -0.547 | 7.54E-06 |
| Nsp12 | CACNG7 | -0.547 | 7.22E-06 |
| Nsp12 | TRADD | -0.547 | 7.38E-06 |
| Nsp12 | GYS1 | -0.547 | 7.35E-06 |
| Nsp12 | DUSP4 | -0.547 | 7.34E-06 |
| Nsp12 | AKT2 | -0.547 | 7.39E-06 |
| Nsp12 | CD70 | -0.547 | 7.42E-06 |
| Nsp12 | COL6A2 | -0.547 | 7.44E-06 |
| Nsp13 | EPOR | -0.547 | 7.47E-06 |
| Nsp13 | GNAI2 | -0.547 | 7.54E-06 |
| Nsp13 | GRK5 | -0.547 | 7.39E-06 |
| Nsp14 | WNT7B | -0.547 | 7.54E-06 |
| Nsp14 | WNT4 | -0.547 | 7.38E-06 |
| Nsp15 | FOSB | -0.547 | 7.43E-06 |
| Nsp15 | TELO2 | -0.547 | 7.30E-06 |
| Nsp15 | WNT10A | -0.547 | 7.48E-06 |
| Nsp15 | VEGFA | -0.547 | 7.49E-06 |
| Nsp15 | PTPN7 | -0.547 | 7.30E-06 |
| Nsp16 | CACNG6 | -0.547 | 7.24E-06 |
| S | FOXO3 | -0.547 | 7.25E-06 |
| S | PIN1 | -0.547 | 7.45E-06 |
| Orf8 | TNFRSF8 | -0.547 | 7.21E-06 |
| Orf8 | MAPK8IP2 | -0.547 | 7.29E-06 |
| Orf8 | DUSP6 | -0.547 | 7.33E-06 |
| Nsp2 | WNT6 | -0.548 | 6.94E-06 |
| Nsp3 | CACNA1C | -0.548 | 6.89E-06 |
| Nsp3 | FGF18 | -0.548 | 7.00E-06 |

| | | | |
|---|---|---|---|
| Nsp3 | CAPN2 | -0.548 | 7.14E-06 |
| Nsp4 | PHLPP1 | -0.548 | 6.92E-06 |
| Nsp4 | FGF3 | -0.548 | 7.06E-06 |
| Nsp4 | SH2B2 | -0.548 | 7.05E-06 |
| Nsp4 | PXN | -0.548 | 7.18E-06 |
| Nsp5 | CACNG1 | -0.548 | 6.91E-06 |
| Nsp5 | RPTOR | -0.548 | 7.10E-06 |
| Nsp5 | SOCS3 | -0.548 | 6.98E-06 |
| Nsp8 | WNT10A | -0.548 | 6.98E-06 |
| Nsp12 | CDC25B | -0.548 | 7.09E-06 |
| Nsp13 | FOXO3 | -0.548 | 6.88E-06 |
| Nsp13 | XCR1 | -0.548 | 7.04E-06 |
| Nsp13 | RASGRF1 | -0.548 | 6.95E-06 |
| Nsp14 | HRK | -0.548 | 6.91E-06 |
| Nsp15 | CHAD | -0.548 | 7.11E-06 |
| Nsp15 | FMOD | -0.548 | 6.98E-06 |
| Nsp15 | CTF1 | -0.548 | 7.14E-06 |
| Nsp16 | RRAS | -0.548 | 6.99E-06 |
| Nsp16 | CACNG4 | -0.548 | 7.15E-06 |
| S | EPHA2 | -0.548 | 7.20E-06 |
| S | GRK6 | -0.548 | 6.96E-06 |
| S | CACNA1G | -0.548 | 7.19E-06 |
| S | SOCS3 | -0.548 | 7.16E-06 |
| Orf8 | ADCY7 | -0.548 | 6.91E-06 |
| Nsp2 | PPP1R3D | -0.549 | 6.82E-06 |
| Nsp3 | TNFRSF8 | -0.549 | 6.64E-06 |
| Nsp3 | CREB3L1 | -0.549 | 6.64E-06 |
| Nsp3 | RASGRF1 | -0.549 | 6.84E-06 |
| Nsp4 | POMC | -0.549 | 6.61E-06 |
| Nsp5 | PIM1 | -0.549 | 6.81E-06 |
| Nsp8 | ADCY1 | -0.549 | 6.78E-06 |
| Nsp8 | EGLN1 | -0.549 | 6.72E-06 |
| Nsp8 | PTPN7 | -0.549 | 6.62E-06 |
| Nsp12 | CAMKK2 | -0.549 | 6.79E-06 |
| Nsp12 | ADCY4 | -0.549 | 6.59E-06 |
| Nsp13 | PRKCZ | -0.549 | 6.68E-06 |
| Nsp13 | MUC5B | -0.549 | 6.58E-06 |
| Nsp13 | GREM1 | -0.549 | 6.57E-06 |
| Nsp13 | ACKR3 | -0.549 | 6.68E-06 |
| Nsp14 | CAPN2 | -0.549 | 6.67E-06 |
| Nsp15 | TNXB | -0.549 | 6.82E-06 |
| Nsp15 | CACNG5 | -0.549 | 6.65E-06 |
| Nsp16 | CTSD | -0.549 | 6.59E-06 |
| Nsp16 | NODAL | -0.549 | 6.67E-06 |
| S | PPP5C | -0.549 | 6.76E-06 |
| Orf8 | GATA3 | -0.549 | 6.81E-06 |
| Orf8 | FZD9 | -0.549 | 6.76E-06 |
| Orf8 | STK11 | -0.549 | 6.84E-06 |
| Nsp3 | MKNK2 | -0.55 | 6.41E-06 |
| Nsp4 | CACNA1G | -0.55 | 6.33E-06 |

| | | | |
|---|---|---|---|
| Nsp4 | FGF17 | -0.55 | 6.28E-06 |
| Nsp4 | PPP2R2C | -0.55 | 6.52E-06 |
| Nsp4 | MAPK4 | -0.55 | 6.37E-06 |
| Nsp4 | JUND | -0.55 | 6.32E-06 |
| Nsp12 | PITX2 | -0.55 | 6.40E-06 |
| Nsp12 | CACNA1C | -0.55 | 6.53E-06 |
| Nsp13 | PDGFB | -0.55 | 6.49E-06 |
| Nsp14 | PRR5 | -0.55 | 6.56E-06 |
| Nsp14 | MAP2K2 | -0.55 | 6.47E-06 |
| Nsp14 | FGF8 | -0.55 | 6.51E-06 |
| Nsp16 | LIF | -0.55 | 6.31E-06 |
| Nsp16 | PIN1 | -0.55 | 6.56E-06 |
| S | CACNG1 | -0.55 | 6.56E-06 |
| Nsp2 | DDIT4 | -0.551 | 6.18E-06 |
| Nsp2 | TRADD | -0.551 | 5.99E-06 |
| Nsp2 | PGAM5 | -0.551 | 6.16E-06 |
| Nsp2 | STK11 | -0.551 | 6.07E-06 |
| Nsp3 | PDGFB | -0.551 | 6.07E-06 |
| Nsp4 | SRC | -0.551 | 6.13E-06 |
| Nsp4 | RPTOR | -0.551 | 6.07E-06 |
| Nsp4 | DUSP4 | -0.551 | 5.99E-06 |
| Nsp5 | POMC | -0.551 | 5.98E-06 |
| Nsp5 | STK11 | -0.551 | 6.05E-06 |
| Nsp8 | CXCR3 | -0.551 | 6.18E-06 |
| Nsp8 | PKN1 | -0.551 | 6.17E-06 |
| Nsp12 | COL6A1 | -0.551 | 6.13E-06 |
| Nsp13 | RGMA | -0.551 | 6.04E-06 |
| Nsp13 | LAMC3 | -0.551 | 6.06E-06 |
| Nsp13 | CD70 | -0.551 | 6.17E-06 |
| Nsp14 | TRADD | -0.551 | 6.21E-06 |
| Orf8 | EFNA2 | -0.551 | 6.21E-06 |
| Nsp2 | LTB | -0.552 | 5.81E-06 |
| Nsp3 | GYS1 | -0.552 | 5.90E-06 |
| Nsp3 | FGF3 | -0.552 | 5.96E-06 |
| Nsp3 | CSH2 | -0.552 | 5.76E-06 |
| Nsp4 | ADCY9 | -0.552 | 5.85E-06 |
| Nsp4 | HSPA1B | -0.552 | 5.96E-06 |
| Nsp5 | PIAS4 | -0.552 | 5.89E-06 |
| Nsp5 | PTPN7 | -0.552 | 5.92E-06 |
| Nsp8 | RASGRF1 | -0.552 | 5.83E-06 |
| Nsp12 | ATP6V1B1 | -0.552 | 5.85E-06 |
| Nsp12 | ITGB4 | -0.552 | 5.97E-06 |
| Nsp13 | TLR9 | -0.552 | 5.74E-06 |
| Nsp13 | YWHAG | -0.552 | 5.86E-06 |
| Nsp14 | PHLPP1 | -0.552 | 5.95E-06 |
| Nsp15 | IL34 | -0.552 | 5.84E-06 |
| Nsp15 | VAV2 | -0.552 | 5.86E-06 |
| Nsp16 | CDKN2B | -0.552 | 5.74E-06 |
| Nsp16 | MLST8 | -0.552 | 5.88E-06 |
| Orf8 | FZD8 | -0.552 | 5.80E-06 |

| | | | |
|---|---|---|---|
| Orf8 | LRP5 | -0.552 | 5.87E-06 |
| Nsp3 | CHAD | -0.553 | 5.57E-06 |
| Nsp3 | CACNG5 | -0.553 | 5.70E-06 |
| Nsp5 | GATA3 | -0.553 | 5.62E-06 |
| Nsp5 | CACNA1B | -0.553 | 5.49E-06 |
| Nsp5 | ADCY7 | -0.553 | 5.58E-06 |
| Nsp12 | IL17RC | -0.553 | 5.51E-06 |
| Nsp12 | CACNA1G | -0.553 | 5.59E-06 |
| Nsp12 | VAV2 | -0.553 | 5.57E-06 |
| Nsp12 | BCL3 | -0.553 | 5.47E-06 |
| Nsp12 | GNA12 | -0.553 | 5.47E-06 |
| Nsp12 | FGF8 | -0.553 | 5.60E-06 |
| Nsp13 | ITPR3 | -0.553 | 5.51E-06 |
| Nsp13 | ECSIT | -0.553 | 5.56E-06 |
| Nsp13 | CACNG5 | -0.553 | 5.64E-06 |
| Nsp13 | TNFRSF13C | -0.553 | 5.58E-06 |
| Nsp14 | HSPA1A | -0.553 | 5.57E-06 |
| Nsp14 | BCL2 | -0.553 | 5.64E-06 |
| Nsp15 | CREB3L1 | -0.553 | 5.66E-06 |
| Nsp15 | CACNA1A | -0.553 | 5.55E-06 |
| Nsp16 | AKT1 | -0.553 | 5.57E-06 |
| Nsp16 | TRADD | -0.553 | 5.63E-06 |
| Nsp16 | MAPK3 | -0.553 | 5.50E-06 |
| Nsp16 | WNT4 | -0.553 | 5.65E-06 |
| Nsp16 | WNT11 | -0.553 | 5.49E-06 |
| Nsp16 | CPT1B | -0.553 | 5.50E-06 |
| S | IRF7 | -0.553 | 5.51E-06 |
| S | CXCR3 | -0.553 | 5.47E-06 |
| S | GATA3 | -0.553 | 5.53E-06 |
| S | MAP3K6 | -0.553 | 5.57E-06 |
| Orf8 | HSPB1 | -0.553 | 5.53E-06 |
| Nsp2 | FZD10 | -0.554 | 5.20E-06 |
| Nsp2 | CACNG8 | -0.554 | 5.36E-06 |
| Nsp3 | FLT3LG | -0.554 | 5.27E-06 |
| Nsp3 | VEGFA | -0.554 | 5.21E-06 |
| Nsp4 | CACNG7 | -0.554 | 5.38E-06 |
| Nsp4 | ITGB4 | -0.554 | 5.34E-06 |
| Nsp4 | MMP9 | -0.554 | 5.32E-06 |
| Nsp4 | AKT2 | -0.554 | 5.25E-06 |
| Nsp8 | TOLLIP | -0.554 | 5.35E-06 |
| Nsp8 | RUNX1 | -0.554 | 5.21E-06 |
| Nsp12 | TNFRSF8 | -0.554 | 5.26E-06 |
| Nsp12 | WNT11 | -0.554 | 5.20E-06 |
| Nsp12 | LAMA5 | -0.554 | 5.35E-06 |
| Nsp13 | CAPN1 | -0.554 | 5.29E-06 |
| Nsp14 | DAB2IP | -0.554 | 5.22E-06 |
| Nsp14 | RASGRP2 | -0.554 | 5.27E-06 |
| Nsp15 | HSPB1 | -0.554 | 5.35E-06 |
| Nsp15 | PRR5 | -0.554 | 5.38E-06 |
| Nsp15 | PREX1 | -0.554 | 5.30E-06 |

| | | | |
|---|---|---|---|
| Nsp15 | DUSP6 | -0.554 | 5.27E-06 |
| Nsp16 | AKT2 | -0.554 | 5.40E-06 |
| Nsp16 | PFKFB3 | -0.554 | 5.25E-06 |
| S | CDC25B | -0.554 | 5.26E-06 |
| S | PTPN6 | -0.554 | 5.35E-06 |
| S | FGF19 | -0.554 | 5.26E-06 |
| S | PPP2R2C | -0.554 | 5.24E-06 |
| Orf8 | CARD14 | -0.554 | 5.41E-06 |
| Nsp2 | GDF1 | -0.555 | 5.04E-06 |
| Nsp3 | TRADD | -0.555 | 5.12E-06 |
| Nsp4 | IL17RC | -0.555 | 5.06E-06 |
| Nsp4 | CXCR3 | -0.555 | 4.98E-06 |
| Nsp5 | WNT3 | -0.555 | 5.07E-06 |
| Nsp5 | PRKCG | -0.555 | 5.18E-06 |
| Nsp8 | LCK | -0.555 | 5.11E-06 |
| Nsp8 | HSPA1B | -0.555 | 5.15E-06 |
| Nsp10 | FGF4 | -0.555 | 5.16E-06 |
| Nsp12 | PHLPP1 | -0.555 | 5.01E-06 |
| Nsp12 | TICAM1 | -0.555 | 4.96E-06 |
| Nsp13 | TRADD | -0.555 | 5.03E-06 |
| Nsp13 | TNFRSF6B | -0.555 | 5.13E-06 |
| Nsp14 | WNT3 | -0.555 | 5.09E-06 |
| Nsp15 | CACNA1G | -0.555 | 5.07E-06 |
| Nsp15 | FOXO3 | -0.555 | 4.96E-06 |
| Nsp15 | BCL2 | -0.555 | 5.07E-06 |
| Nsp16 | DUSP1 | -0.555 | 5.08E-06 |
| Nsp16 | FLNC | -0.555 | 5.07E-06 |
| S | DVL3 | -0.555 | 4.98E-06 |
| S | WNT4 | -0.555 | 5.01E-06 |
| Orf8 | GREM2 | -0.555 | 5.13E-06 |
| Orf8 | BMP8B | -0.555 | 4.96E-06 |
| Orf8 | ZAP70 | -0.555 | 5.00E-06 |
| Nsp4 | RGMA | -0.556 | 4.82E-06 |
| Nsp4 | PFKFB3 | -0.556 | 4.89E-06 |
| Nsp5 | ACKR3 | -0.556 | 4.78E-06 |
| Nsp5 | GADD45B | -0.556 | 4.88E-06 |
| Nsp5 | FOXO3 | -0.556 | 4.79E-06 |
| Nsp5 | COL6A2 | -0.556 | 4.87E-06 |
| Nsp5 | LAMA5 | -0.556 | 4.73E-06 |
| Nsp8 | ACKR3 | -0.556 | 4.94E-06 |
| Nsp12 | DUSP1 | -0.556 | 4.87E-06 |
| Nsp12 | PREX1 | -0.556 | 4.80E-06 |
| Nsp12 | CALML5 | -0.556 | 4.78E-06 |
| Nsp13 | FOSL1 | -0.556 | 4.84E-06 |
| Nsp14 | FZD7 | -0.556 | 4.83E-06 |
| Nsp14 | RELB | -0.556 | 4.91E-06 |
| Nsp14 | TICAM1 | -0.556 | 4.83E-06 |
| Nsp15 | ITGB4 | -0.556 | 4.90E-06 |
| Nsp16 | CALML5 | -0.556 | 4.74E-06 |
| Nsp16 | HSPA1B | -0.556 | 4.75E-06 |

| | | | |
|---|---|---|---|
| S | CHAD | -0.556 | 4.89E-06 |
| S | NGFR | -0.556 | 4.82E-06 |
| Nsp3 | EPHA2 | -0.557 | 4.54E-06 |
| Nsp3 | WNT9A | -0.557 | 4.69E-06 |
| Nsp4 | HSPA1A | -0.557 | 4.64E-06 |
| Nsp5 | MMP9 | -0.557 | 4.56E-06 |
| Nsp5 | BCL2 | -0.557 | 4.58E-06 |
| Nsp8 | MAPK11 | -0.557 | 4.61E-06 |
| Nsp8 | PPP1R3E | -0.557 | 4.53E-06 |
| Nsp12 | DAB2IP | -0.557 | 4.51E-06 |
| Nsp12 | PRKCG | -0.557 | 4.72E-06 |
| Nsp13 | GDF11 | -0.557 | 4.68E-06 |
| Nsp13 | VEGFA | -0.557 | 4.56E-06 |
| Nsp14 | ENDOG | -0.557 | 4.71E-06 |
| Nsp14 | WNT5B | -0.557 | 4.53E-06 |
| Nsp15 | RELB | -0.557 | 4.53E-06 |
| Nsp15 | WNT9A | -0.557 | 4.70E-06 |
| Nsp15 | MAPK8IP3 | -0.557 | 4.72E-06 |
| Nsp16 | PIK3R5 | -0.557 | 4.65E-06 |
| Nsp16 | MAPK8IP1 | -0.557 | 4.53E-06 |
| Nsp16 | MAP3K11 | -0.557 | 4.54E-06 |
| Nsp16 | GATA3 | -0.557 | 4.60E-06 |
| Nsp16 | FGF17 | -0.557 | 4.67E-06 |
| Nsp16 | CREB3L1 | -0.557 | 4.63E-06 |
| Nsp16 | EGLN1 | -0.557 | 4.54E-06 |
| Nsp16 | PLCB3 | -0.557 | 4.51E-06 |
| S | YWHAG | -0.557 | 4.55E-06 |
| S | MAPK3 | -0.557 | 4.53E-06 |
| Orf3a | FGF4 | -0.557 | 4.65E-06 |
| Nsp3 | IL17RC | -0.558 | 4.49E-06 |
| Nsp3 | WDR24 | -0.558 | 4.39E-06 |
| Nsp3 | PKN1 | -0.558 | 4.44E-06 |
| Nsp3 | PLCB3 | -0.558 | 4.48E-06 |
| Nsp4 | GDF11 | -0.558 | 4.45E-06 |
| Nsp4 | CALML5 | -0.558 | 4.39E-06 |
| Nsp4 | COL6A1 | -0.558 | 4.34E-06 |
| Nsp4 | COL6A2 | -0.558 | 4.50E-06 |
| Nsp5 | IL34 | -0.558 | 4.34E-06 |
| Nsp5 | CARD11 | -0.558 | 4.29E-06 |
| Nsp12 | HSPA1B | -0.558 | 4.45E-06 |
| Nsp13 | FLT3LG | -0.558 | 4.47E-06 |
| Nsp13 | SOCS3 | -0.558 | 4.39E-06 |
| Nsp14 | NOS3 | -0.558 | 4.44E-06 |
| Nsp14 | GATA3 | -0.558 | 4.40E-06 |
| Nsp15 | SLC7A5 | -0.558 | 4.31E-06 |
| Nsp15 | WNT1 | -0.558 | 4.34E-06 |
| Nsp15 | POMC | -0.558 | 4.41E-06 |
| Nsp15 | CACNA1B | -0.558 | 4.42E-06 |
| Nsp16 | FGF19 | -0.558 | 4.49E-06 |
| Nsp16 | MYC | -0.558 | 4.33E-06 |

| | | | |
|---|---|---|---|
| S | WDR24 | -0.558 | 4.44E-06 |
| S | PTPN7 | -0.558 | 4.46E-06 |
| Orf8 | JUN | -0.558 | 4.46E-06 |
| Orf8 | PPP2R3B | -0.558 | 4.37E-06 |
| Orf8 | FZD1 | -0.558 | 4.30E-06 |
| Orf8 | LTB | -0.558 | 4.30E-06 |
| Orf8 | FGF8 | -0.558 | 4.33E-06 |
| Nsp2 | IRS2 | -0.559 | 4.12E-06 |
| Nsp4 | DAB2IP | -0.559 | 4.29E-06 |
| Nsp4 | RARA | -0.559 | 4.18E-06 |
| Nsp5 | SMAD7 | -0.559 | 4.20E-06 |
| Nsp5 | TNFRSF6B | -0.559 | 4.27E-06 |
| Nsp8 | HSPA1A | -0.559 | 4.23E-06 |
| Nsp8 | XCR1 | -0.559 | 4.11E-06 |
| Nsp12 | RRAS | -0.559 | 4.10E-06 |
| Nsp13 | CTSD | -0.559 | 4.13E-06 |
| Nsp13 | FGF19 | -0.559 | 4.17E-06 |
| Nsp13 | POMC | -0.559 | 4.18E-06 |
| Nsp14 | BMP8A | -0.559 | 4.19E-06 |
| Nsp14 | PIK3R2 | -0.559 | 4.29E-06 |
| Nsp15 | CACNG6 | -0.559 | 4.25E-06 |
| Nsp15 | MLST8 | -0.559 | 4.15E-06 |
| Nsp15 | CPT1B | -0.559 | 4.22E-06 |
| Nsp16 | IL17RC | -0.559 | 4.19E-06 |
| Nsp16 | HSPA1A | -0.559 | 4.28E-06 |
| Nsp16 | TCL1A | -0.559 | 4.10E-06 |
| S | CACNG6 | -0.559 | 4.19E-06 |
| S | SRC | -0.559 | 4.21E-06 |
| Orf8 | LMNB2 | -0.559 | 4.15E-06 |
| Orf8 | WNT5B | -0.559 | 4.26E-06 |
| Nsp2 | SMAD6 | -0.56 | 3.93E-06 |
| Nsp3 | IL27 | -0.56 | 3.99E-06 |
| Nsp3 | CXCR3 | -0.56 | 4.00E-06 |
| Nsp3 | PFKFB3 | -0.56 | 4.09E-06 |
| Nsp3 | CPT1C | -0.56 | 3.92E-06 |
| Nsp4 | PRR5 | -0.56 | 3.98E-06 |
| Nsp4 | TNFRSF6B | -0.56 | 3.92E-06 |
| Nsp4 | LAMA5 | -0.56 | 3.91E-06 |
| Nsp5 | CACNA1H | -0.56 | 4.05E-06 |
| Nsp8 | CACNG6 | -0.56 | 4.04E-06 |
| Nsp8 | SRC | -0.56 | 3.94E-06 |
| Nsp8 | WNT9A | -0.56 | 3.92E-06 |
| Nsp12 | RARA | -0.56 | 4.03E-06 |
| Nsp12 | PKN1 | -0.56 | 4.01E-06 |
| Nsp12 | ZAP70 | -0.56 | 4.04E-06 |
| Nsp13 | TOLLIP | -0.56 | 4.04E-06 |
| Nsp13 | ZAP70 | -0.56 | 3.98E-06 |
| Nsp15 | DUSP1 | -0.56 | 4.06E-06 |
| Nsp15 | GRK2 | -0.56 | 3.91E-06 |
| Nsp15 | HRK | -0.56 | 4.01E-06 |

| | | | |
|---|---|---|---|
| Nsp16 | DHX58 | -0.56 | 4.01E-06 |
| Nsp16 | WNT10A | -0.56 | 3.94E-06 |
| S | WNT9A | -0.56 | 4.08E-06 |
| S | ULK1 | -0.56 | 4.04E-06 |
| Orf8 | BMP7 | -0.56 | 4.01E-06 |
| Nsp3 | TLR9 | -0.561 | 3.79E-06 |
| Nsp3 | PPP2R2C | -0.561 | 3.87E-06 |
| Nsp4 | CDKN2B | -0.561 | 3.83E-06 |
| Nsp5 | PRR5 | -0.561 | 3.84E-06 |
| Nsp5 | PFKFB3 | -0.561 | 3.75E-06 |
| Nsp5 | HRK | -0.561 | 3.89E-06 |
| Nsp8 | LEFTY1 | -0.561 | 3.90E-06 |
| Nsp12 | FZD2 | -0.561 | 3.82E-06 |
| Nsp13 | PITX2 | -0.561 | 3.87E-06 |
| Nsp13 | INSR | -0.561 | 3.83E-06 |
| Nsp14 | IL34 | -0.561 | 3.87E-06 |
| Nsp14 | IL17RA | -0.561 | 3.74E-06 |
| Nsp14 | FGFR3 | -0.561 | 3.79E-06 |
| Nsp15 | PFKFB3 | -0.561 | 3.85E-06 |
| Nsp15 | COL6A1 | -0.561 | 3.80E-06 |
| Nsp16 | TNFSF14 | -0.561 | 3.76E-06 |
| S | VEGFA | -0.561 | 3.74E-06 |
| Orf8 | DUSP7 | -0.561 | 3.74E-06 |
| Nsp2 | FZD9 | -0.562 | 3.61E-06 |
| Nsp3 | CCR10 | -0.562 | 3.71E-06 |
| Nsp3 | DUSP6 | -0.562 | 3.62E-06 |
| Nsp3 | ADCY4 | -0.562 | 3.69E-06 |
| Nsp3 | HSPA1B | -0.562 | 3.63E-06 |
| Nsp4 | MAPK8IP3 | -0.562 | 3.54E-06 |
| Nsp5 | MAPK3 | -0.562 | 3.61E-06 |
| Nsp5 | DUSP6 | -0.562 | 3.58E-06 |
| Nsp8 | IRF7 | -0.562 | 3.61E-06 |
| Nsp12 | ADCY9 | -0.562 | 3.63E-06 |
| Nsp14 | CACNA1H | -0.562 | 3.59E-06 |
| Nsp15 | GRK1 | -0.562 | 3.60E-06 |
| Nsp15 | CALML5 | -0.562 | 3.55E-06 |
| Nsp15 | FGFR3 | -0.562 | 3.57E-06 |
| Nsp16 | WNT1 | -0.562 | 3.70E-06 |
| Nsp16 | JAK3 | -0.562 | 3.62E-06 |
| S | ITPR3 | -0.562 | 3.56E-06 |
| Nsp2 | EFNA3 | -0.563 | 3.39E-06 |
| Nsp3 | ITPR3 | -0.563 | 3.40E-06 |
| Nsp3 | CACNA1G | -0.563 | 3.46E-06 |
| Nsp3 | ACKR3 | -0.563 | 3.39E-06 |
| Nsp4 | PITX2 | -0.563 | 3.50E-06 |
| Nsp4 | TNFRSF4 | -0.563 | 3.53E-06 |
| Nsp4 | FZD2 | -0.563 | 3.40E-06 |
| Nsp5 | CDKN2B | -0.563 | 3.50E-06 |
| Nsp8 | PPP2R2C | -0.563 | 3.48E-06 |
| Nsp8 | CTF1 | -0.563 | 3.40E-06 |

| | | | |
|---|---|---|---|
| Nsp12 | HSPA1A | -0.563 | 3.52E-06 |
| Nsp12 | BAD | -0.563 | 3.41E-06 |
| Nsp12 | PTPN7 | -0.563 | 3.39E-06 |
| Nsp13 | CXCL14 | -0.563 | 3.40E-06 |
| Nsp14 | RGMA | -0.563 | 3.44E-06 |
| Nsp14 | GDF1 | -0.563 | 3.46E-06 |
| Nsp14 | FGF22 | -0.563 | 3.43E-06 |
| Nsp15 | JUND | -0.563 | 3.49E-06 |
| Nsp15 | LAMA5 | -0.563 | 3.52E-06 |
| Nsp16 | NR4A1 | -0.563 | 3.41E-06 |
| Nsp16 | ARAF | -0.563 | 3.43E-06 |
| Nsp16 | FMOD | -0.563 | 3.49E-06 |
| Nsp16 | TYK2 | -0.563 | 3.43E-06 |
| Orf8 | JUNB | -0.563 | 3.51E-06 |
| Nsp3 | ADCY1 | -0.564 | 3.21E-06 |
| Nsp3 | CDKN2B | -0.564 | 3.27E-06 |
| Nsp3 | FGF17 | -0.564 | 3.29E-06 |
| Nsp3 | XCR1 | -0.564 | 3.30E-06 |
| Nsp5 | COMP | -0.564 | 3.29E-06 |
| Nsp12 | SRC | -0.564 | 3.24E-06 |
| Nsp12 | TNFRSF6B | -0.564 | 3.27E-06 |
| Nsp12 | TBX21 | -0.564 | 3.26E-06 |
| Nsp12 | TNFRSF4 | -0.564 | 3.22E-06 |
| Nsp12 | PLCB3 | -0.564 | 3.33E-06 |
| Nsp13 | DUSP5 | -0.564 | 3.33E-06 |
| Nsp13 | VAV2 | -0.564 | 3.27E-06 |
| Nsp13 | COL6A1 | -0.564 | 3.22E-06 |
| Nsp14 | GRK2 | -0.564 | 3.31E-06 |
| Nsp15 | CACNG1 | -0.564 | 3.31E-06 |
| Nsp15 | NOS3 | -0.564 | 3.36E-06 |
| Nsp15 | JUNB | -0.564 | 3.28E-06 |
| Nsp15 | CD70 | -0.564 | 3.28E-06 |
| Nsp15 | PIK3CD | -0.564 | 3.22E-06 |
| Nsp15 | HSPA1A | -0.564 | 3.28E-06 |
| Nsp15 | HSPA1B | -0.564 | 3.36E-06 |
| Nsp15 | TICAM1 | -0.564 | 3.28E-06 |
| Nsp16 | GRK1 | -0.564 | 3.23E-06 |
| Nsp16 | ACKR3 | -0.564 | 3.31E-06 |
| S | CACNA1A | -0.564 | 3.27E-06 |
| Orf8 | LPAR5 | -0.564 | 3.22E-06 |
| Orf8 | PIM1 | -0.564 | 3.24E-06 |
| Orf8 | CACNA1I | -0.564 | 3.23E-06 |
| Nsp3 | FMOD | -0.565 | 3.17E-06 |
| Nsp3 | GRK1 | -0.565 | 3.08E-06 |
| Nsp4 | RELB | -0.565 | 3.10E-06 |
| Nsp4 | MAPK11 | -0.565 | 3.08E-06 |
| Nsp4 | ICAM1 | -0.565 | 3.15E-06 |
| Nsp4 | EGLN1 | -0.565 | 3.15E-06 |
| Nsp4 | TNFRSF13C | -0.565 | 3.18E-06 |
| Nsp5 | MAPK8IP1 | -0.565 | 3.07E-06 |

| | | | |
|---|---|---|---|
| Nsp5 | EPOR | -0.565 | 3.11E-06 |
| Nsp5 | NOS3 | -0.565 | 3.16E-06 |
| Nsp5 | PPP1R3E | -0.565 | 3.08E-06 |
| Nsp5 | RARA | -0.565 | 3.17E-06 |
| Nsp8 | DUSP1 | -0.565 | 3.15E-06 |
| Nsp8 | MAP2K2 | -0.565 | 3.12E-06 |
| Nsp12 | MAPK4 | -0.565 | 3.06E-06 |
| Nsp12 | EGLN1 | -0.565 | 3.08E-06 |
| Nsp13 | CREB3L1 | -0.565 | 3.19E-06 |
| Nsp14 | PPP1R3D | -0.565 | 3.12E-06 |
| Nsp15 | PPP3R2 | -0.565 | 3.18E-06 |
| Nsp15 | PRKCG | -0.565 | 3.17E-06 |
| Nsp15 | RARA | -0.565 | 3.15E-06 |
| Nsp16 | BCAR1 | -0.565 | 3.17E-06 |
| Nsp16 | PFKL | -0.565 | 3.08E-06 |
| Nsp16 | CACNG5 | -0.565 | 3.11E-06 |
| S | TLR9 | -0.565 | 3.13E-06 |
| Orf8 | DDIT4 | -0.565 | 3.16E-06 |
| Nsp3 | CACNG6 | -0.566 | 2.95E-06 |
| Nsp3 | IRF7 | -0.566 | 3.05E-06 |
| Nsp3 | YWHAG | -0.566 | 3.01E-06 |
| Nsp3 | MAPK3 | -0.566 | 3.05E-06 |
| Nsp3 | PXN | -0.566 | 2.94E-06 |
| Nsp4 | GRK1 | -0.566 | 2.96E-06 |
| Nsp4 | ADCY3 | -0.566 | 3.04E-06 |
| Nsp5 | CAPN1 | -0.566 | 2.92E-06 |
| Nsp5 | EGLN1 | -0.566 | 3.01E-06 |
| Nsp8 | ITGB4 | -0.566 | 2.96E-06 |
| Nsp8 | NOS3 | -0.566 | 3.03E-06 |
| Nsp8 | LEFTY2 | -0.566 | 3.00E-06 |
| Nsp13 | SRC | -0.566 | 3.04E-06 |
| Nsp13 | DAB2IP | -0.566 | 3.04E-06 |
| Nsp13 | CCR10 | -0.566 | 3.01E-06 |
| Nsp14 | COMP | -0.566 | 2.97E-06 |
| Nsp15 | ENDOG | -0.566 | 2.94E-06 |
| Nsp16 | WNT7A | -0.566 | 2.93E-06 |
| Nsp16 | HRK | -0.566 | 3.01E-06 |
| Nsp16 | CCND1 | -0.566 | 3.02E-06 |
| S | ITGB4 | -0.566 | 3.05E-06 |
| S | COL6A1 | -0.566 | 2.99E-06 |
| Orf8 | SMAD6 | -0.566 | 3.04E-06 |
| Orf8 | HRK | -0.566 | 3.01E-06 |
| Nsp3 | ADCY9 | -0.567 | 2.77E-06 |
| Nsp3 | PTPN7 | -0.567 | 2.88E-06 |
| Nsp8 | CCR10 | -0.567 | 2.84E-06 |
| Nsp10 | PPP2R3B | -0.567 | 2.82E-06 |
| Nsp12 | GDF11 | -0.567 | 2.91E-06 |
| Nsp12 | MAPK8IP3 | -0.567 | 2.87E-06 |
| Nsp14 | DUSP7 | -0.567 | 2.83E-06 |
| Nsp14 | AMH | -0.567 | 2.82E-06 |

| | | | |
|---|---|---|---|
| Nsp15 | ADCY4 | -0.567 | 2.86E-06 |
| Nsp15 | FGF22 | -0.567 | 2.88E-06 |
| Nsp16 | FGFR4 | -0.567 | 2.79E-06 |
| Nsp16 | YWHAG | -0.567 | 2.91E-06 |
| S | DUSP1 | -0.567 | 2.78E-06 |
| S | ID1 | -0.567 | 2.84E-06 |
| S | PXN | -0.567 | 2.80E-06 |
| Orf8 | FZD10 | -0.567 | 2.80E-06 |
| Orf8 | PGAM5 | -0.567 | 2.79E-06 |
| Orf8 | NFATC1 | -0.567 | 2.86E-06 |
| Nsp3 | SRC | -0.568 | 2.67E-06 |
| Nsp3 | HSPA1A | -0.568 | 2.64E-06 |
| Nsp3 | FOXO3 | -0.568 | 2.75E-06 |
| Nsp4 | IL34 | -0.568 | 2.72E-06 |
| Nsp4 | MAPK8IP1 | -0.568 | 2.72E-06 |
| Nsp5 | CCND1 | -0.568 | 2.69E-06 |
| Nsp8 | MAPK8IP1 | -0.568 | 2.77E-06 |
| Nsp12 | CACNG4 | -0.568 | 2.70E-06 |
| Nsp14 | HSPA2 | -0.568 | 2.70E-06 |
| Nsp14 | FZD8 | -0.568 | 2.64E-06 |
| Nsp14 | ZAP70 | -0.568 | 2.69E-06 |
| Nsp15 | ADCY3 | -0.568 | 2.71E-06 |
| Nsp15 | COL6A2 | -0.568 | 2.76E-06 |
| Nsp16 | FGF18 | -0.568 | 2.73E-06 |
| Nsp16 | JUND | -0.568 | 2.68E-06 |
| S | BCL3 | -0.568 | 2.69E-06 |
| S | LIF | -0.568 | 2.71E-06 |
| S | ID4 | -0.568 | 2.66E-06 |
| S | CACNA1B | -0.568 | 2.69E-06 |
| Orf8 | BMP8A | -0.568 | 2.71E-06 |
| Nsp2 | ENDOG | -0.569 | 2.58E-06 |
| Nsp3 | CDC25B | -0.569 | 2.58E-06 |
| Nsp3 | DUSP1 | -0.569 | 2.63E-06 |
| Nsp3 | CTSD | -0.569 | 2.58E-06 |
| Nsp3 | GNA12 | -0.569 | 2.53E-06 |
| Nsp4 | DUSP9 | -0.569 | 2.63E-06 |
| Nsp4 | WNT1 | -0.569 | 2.52E-06 |
| Nsp5 | ADCY3 | -0.569 | 2.54E-06 |
| Nsp8 | CARD14 | -0.569 | 2.53E-06 |
| Nsp8 | DAB2IP | -0.569 | 2.55E-06 |
| Nsp8 | PRKCG | -0.569 | 2.61E-06 |
| Nsp8 | ADCY3 | -0.569 | 2.52E-06 |
| Nsp12 | CAPN2 | -0.569 | 2.61E-06 |
| Nsp12 | FGF3 | -0.569 | 2.54E-06 |
| Nsp12 | TNFRSF13C | -0.569 | 2.60E-06 |
| Nsp13 | PPP3R2 | -0.569 | 2.58E-06 |
| Nsp13 | MLST8 | -0.569 | 2.56E-06 |
| Nsp13 | WNT11 | -0.569 | 2.53E-06 |
| Nsp13 | MKNK2 | -0.569 | 2.63E-06 |
| Nsp13 | LAMA5 | -0.569 | 2.53E-06 |

| | | | |
|---|---|---|---|
| Nsp14 | SOCS1 | -0.569 | 2.57E-06 |
| Nsp15 | DUSP9 | -0.569 | 2.53E-06 |
| Nsp16 | NFKBIB | -0.569 | 2.58E-06 |
| Nsp16 | DUSP6 | -0.569 | 2.58E-06 |
| Nsp16 | BCL2 | -0.569 | 2.54E-06 |
| S | ADCY3 | -0.569 | 2.62E-06 |
| Nsp2 | LPAR5 | -0.57 | 2.39E-06 |
| Nsp2 | NOG | -0.57 | 2.45E-06 |
| Nsp3 | WNT10A | -0.57 | 2.49E-06 |
| Nsp4 | FOSB | -0.57 | 2.43E-06 |
| Nsp5 | CACNG6 | -0.57 | 2.46E-06 |
| Nsp5 | MAP2K2 | -0.57 | 2.46E-06 |
| Nsp8 | RASGRP2 | -0.57 | 2.43E-06 |
| Nsp12 | FGF22 | -0.57 | 2.48E-06 |
| Nsp13 | PPP2R2C | -0.57 | 2.40E-06 |
| Nsp14 | CTF1 | -0.57 | 2.40E-06 |
| Nsp15 | DAB2IP | -0.57 | 2.48E-06 |
| Nsp15 | IL17RC | -0.57 | 2.50E-06 |
| Nsp15 | EPOR | -0.57 | 2.41E-06 |
| Nsp15 | PIAS4 | -0.57 | 2.39E-06 |
| Nsp15 | ADCY7 | -0.57 | 2.47E-06 |
| Nsp16 | IL34 | -0.57 | 2.49E-06 |
| S | FGF3 | -0.57 | 2.45E-06 |
| Orf8 | MAP2K7 | -0.57 | 2.45E-06 |
| Nsp2 | DUSP8 | -0.571 | 2.31E-06 |
| Nsp3 | FOSB | -0.571 | 2.30E-06 |
| Nsp3 | MAPK8IP1 | -0.571 | 2.31E-06 |
| Nsp3 | CACNG4 | -0.571 | 2.29E-06 |
| Nsp3 | TOLLIP | -0.571 | 2.29E-06 |
| Nsp3 | PPP1R3E | -0.571 | 2.37E-06 |
| Nsp3 | POMC | -0.571 | 2.36E-06 |
| Nsp12 | RELB | -0.571 | 2.30E-06 |
| Nsp13 | CACNA1G | -0.571 | 2.27E-06 |
| Nsp13 | TBX21 | -0.571 | 2.38E-06 |
| Nsp14 | TNFRSF18 | -0.571 | 2.30E-06 |
| Nsp14 | CACNA1I | -0.571 | 2.33E-06 |
| Nsp15 | SOCS1 | -0.571 | 2.33E-06 |
| Nsp15 | RGMA | -0.571 | 2.31E-06 |
| Nsp15 | LMNB2 | -0.571 | 2.30E-06 |
| Nsp15 | PIK3R2 | -0.571 | 2.31E-06 |
| Nsp15 | CARD11 | -0.571 | 2.33E-06 |
| Nsp16 | FLCN | -0.571 | 2.30E-06 |
| Nsp16 | PREX1 | -0.571 | 2.30E-06 |
| Nsp16 | SH2B2 | -0.571 | 2.35E-06 |
| S | SLC7A5 | -0.571 | 2.32E-06 |
| S | PRKCG | -0.571 | 2.30E-06 |
| S | MAP2K2 | -0.571 | 2.35E-06 |
| Nsp4 | GRK2 | -0.572 | 2.17E-06 |
| Nsp5 | PGAM5 | -0.572 | 2.21E-06 |
| Nsp5 | DUSP4 | -0.572 | 2.24E-06 |

| | | | |
|---|---|---|---|
| Nsp8 | CDC25B | -0.572 | 2.22E-06 |
| Nsp12 | MCL1 | -0.572 | 2.16E-06 |
| Nsp12 | PFKFB3 | -0.572 | 2.17E-06 |
| Nsp12 | CREB3L1 | -0.572 | 2.26E-06 |
| Nsp12 | WNT1 | -0.572 | 2.18E-06 |
| Nsp13 | ULK1 | -0.572 | 2.23E-06 |
| Nsp15 | GADD45B | -0.572 | 2.21E-06 |
| Nsp16 | EPHA2 | -0.572 | 2.18E-06 |
| Nsp16 | SLC7A5 | -0.572 | 2.27E-06 |
| Nsp16 | GNB3 | -0.572 | 2.17E-06 |
| S | RELB | -0.572 | 2.20E-06 |
| Orf8 | DUSP8 | -0.572 | 2.25E-06 |
| Nsp4 | NOS3 | -0.573 | 2.06E-06 |
| Nsp5 | WNT7B | -0.573 | 2.10E-06 |
| Nsp12 | PPP3R2 | -0.573 | 2.13E-06 |
| Nsp12 | RPS6KA2 | -0.573 | 2.07E-06 |
| Nsp13 | CPT1C | -0.573 | 2.10E-06 |
| Nsp13 | PLCB3 | -0.573 | 2.16E-06 |
| Nsp14 | PGAM5 | -0.573 | 2.11E-06 |
| Nsp14 | HSPB1 | -0.573 | 2.12E-06 |
| Nsp14 | GREM2 | -0.573 | 2.16E-06 |
| Nsp14 | DVL1 | -0.573 | 2.07E-06 |
| Nsp15 | WNT3 | -0.573 | 2.11E-06 |
| Nsp16 | MAPK11 | -0.573 | 2.15E-06 |
| Nsp16 | GNA12 | -0.573 | 2.09E-06 |
| Nsp16 | ULK1 | -0.573 | 2.12E-06 |
| Nsp16 | TICAM1 | -0.573 | 2.10E-06 |
| S | DUSP2 | -0.573 | 2.07E-06 |
| S | DUSP5 | -0.573 | 2.08E-06 |
| Orf8 | WNT3 | -0.573 | 2.13E-06 |
| Orf8 | IRS2 | -0.573 | 2.15E-06 |
| Nsp3 | ULK1 | -0.574 | 2.04E-06 |
| Nsp4 | CACNA1H | -0.574 | 1.96E-06 |
| Nsp4 | DUSP5 | -0.574 | 2.05E-06 |
| Nsp4 | FLT4 | -0.574 | 2.04E-06 |
| Nsp8 | CACNA1A | -0.574 | 1.98E-06 |
| Nsp8 | LAMA5 | -0.574 | 1.99E-06 |
| Nsp12 | IL34 | -0.574 | 2.00E-06 |
| Nsp12 | WNT5B | -0.574 | 2.05E-06 |
| Nsp13 | IL27 | -0.574 | 1.99E-06 |
| Nsp13 | GADD45B | -0.574 | 2.04E-06 |
| Nsp14 | TNFRSF6B | -0.574 | 1.96E-06 |
| Nsp15 | TBKBP1 | -0.574 | 2.03E-06 |
| Nsp15 | CACNG7 | -0.574 | 2.02E-06 |
| Nsp15 | PGAM5 | -0.574 | 1.98E-06 |
| Nsp15 | BCL3 | -0.574 | 2.04E-06 |
| Nsp16 | GNAI2 | -0.574 | 2.00E-06 |
| S | NOS3 | -0.574 | 1.96E-06 |
| S | WNT10A | -0.574 | 2.03E-06 |
| S | ADCY7 | -0.574 | 2.02E-06 |

| | | | |
|---|---|---|---|
| S | HSPA1B | -0.574 | 2.03E-06 |
| Nsp2 | CEBPB | -0.575 | 1.93E-06 |
| Nsp3 | TNXB | -0.575 | 1.92E-06 |
| Nsp3 | DUSP4 | -0.575 | 1.95E-06 |
| Nsp3 | PREX1 | -0.575 | 1.89E-06 |
| Nsp4 | HRK | -0.575 | 1.88E-06 |
| Nsp5 | CACNG7 | -0.575 | 1.95E-06 |
| Nsp5 | FGF3 | -0.575 | 1.92E-06 |
| Nsp5 | VEGFA | -0.575 | 1.91E-06 |
| Nsp5 | FGF8 | -0.575 | 1.87E-06 |
| Nsp8 | BMP8A | -0.575 | 1.89E-06 |
| Nsp8 | BCL2 | -0.575 | 1.90E-06 |
| Nsp12 | JUNB | -0.575 | 1.86E-06 |
| Nsp13 | CACNA1C | -0.575 | 1.88E-06 |
| Nsp13 | FGF18 | -0.575 | 1.87E-06 |
| Nsp13 | PTPN6 | -0.575 | 1.92E-06 |
| Nsp14 | CARD14 | -0.575 | 1.90E-06 |
| Nsp16 | ITPR3 | -0.575 | 1.88E-06 |
| Nsp16 | GRK6 | -0.575 | 1.93E-06 |
| Nsp16 | FGF22 | -0.575 | 1.90E-06 |
| S | DAB2IP | -0.575 | 1.93E-06 |
| S | TNFRSF4 | -0.575 | 1.91E-06 |
| S | LAMA5 | -0.575 | 1.87E-06 |
| Nsp3 | MLST8 | -0.576 | 1.85E-06 |
| Nsp3 | TNFRSF4 | -0.576 | 1.80E-06 |
| Nsp3 | PRKCG | -0.576 | 1.79E-06 |
| Nsp3 | BAD | -0.576 | 1.83E-06 |
| Nsp4 | ADCY5 | -0.576 | 1.77E-06 |
| Nsp8 | CACNG7 | -0.576 | 1.81E-06 |
| Nsp12 | MAPK3 | -0.576 | 1.84E-06 |
| Nsp12 | LMNB2 | -0.576 | 1.82E-06 |
| Nsp13 | PPP2R1A | -0.576 | 1.84E-06 |
| Nsp13 | RELB | -0.576 | 1.82E-06 |
| Nsp13 | FGF17 | -0.576 | 1.81E-06 |
| Nsp13 | MAPK8IP3 | -0.576 | 1.77E-06 |
| Nsp15 | PRKAR1B | -0.576 | 1.79E-06 |
| Nsp15 | MAP2K2 | -0.576 | 1.84E-06 |
| Nsp16 | SMAD7 | -0.576 | 1.80E-06 |
| Nsp16 | TLR9 | -0.576 | 1.80E-06 |
| Nsp16 | WDR24 | -0.576 | 1.81E-06 |
| Nsp16 | CACNA1B | -0.576 | 1.82E-06 |
| Nsp16 | CTF1 | -0.576 | 1.80E-06 |
| S | MMP9 | -0.576 | 1.78E-06 |
| Nsp2 | AMH | -0.577 | 1.70E-06 |
| Nsp3 | WNT11 | -0.577 | 1.68E-06 |
| Nsp4 | CCND1 | -0.577 | 1.73E-06 |
| Nsp8 | GNA12 | -0.577 | 1.71E-06 |
| Nsp8 | POMC | -0.577 | 1.70E-06 |
| Nsp8 | ULK1 | -0.577 | 1.68E-06 |
| Nsp12 | GRK2 | -0.577 | 1.70E-06 |

| | | | |
|---|---|---|---|
| Nsp12 | DUSP6 | -0.577 | 1.72E-06 |
| Nsp13 | TNFRSF8 | -0.577 | 1.75E-06 |
| Nsp13 | ADCY1 | -0.577 | 1.70E-06 |
| Nsp13 | PKN1 | -0.577 | 1.70E-06 |
| Nsp13 | PXN | -0.577 | 1.72E-06 |
| Nsp14 | LEFTY2 | -0.577 | 1.74E-06 |
| Nsp14 | PIAS4 | -0.577 | 1.75E-06 |
| Nsp14 | FZD5 | -0.577 | 1.69E-06 |
| Nsp15 | FLT4 | -0.577 | 1.73E-06 |
| Nsp15 | FGF8 | -0.577 | 1.71E-06 |
| Nsp16 | CDC25B | -0.577 | 1.72E-06 |
| Nsp16 | BCL3 | -0.577 | 1.70E-06 |
| Nsp16 | COL6A1 | -0.577 | 1.71E-06 |
| Orf8 | ADCY1 | -0.577 | 1.74E-06 |
| Nsp3 | ITGB4 | -0.578 | 1.63E-06 |
| Nsp3 | AKT2 | -0.578 | 1.62E-06 |
| Nsp4 | CACNG1 | -0.578 | 1.64E-06 |
| Nsp4 | CAPN2 | -0.578 | 1.66E-06 |
| Nsp4 | CARD11 | -0.578 | 1.67E-06 |
| Nsp5 | MAPK8IP3 | -0.578 | 1.62E-06 |
| Nsp5 | FGFR3 | -0.578 | 1.61E-06 |
| Nsp8 | YWHAG | -0.578 | 1.60E-06 |
| Nsp8 | MMP9 | -0.578 | 1.64E-06 |
| Nsp8 | MAPK4 | -0.578 | 1.63E-06 |
| Nsp12 | GRK1 | -0.578 | 1.66E-06 |
| Nsp13 | IL17RC | -0.578 | 1.61E-06 |
| Nsp13 | MAPK3 | -0.578 | 1.62E-06 |
| Nsp15 | STK11 | -0.578 | 1.64E-06 |
| Nsp16 | GSK3A | -0.578 | 1.60E-06 |
| Nsp16 | RPS6KB2 | -0.578 | 1.63E-06 |
| Nsp16 | CACNA1A | -0.578 | 1.62E-06 |
| Nsp16 | PKN1 | -0.578 | 1.61E-06 |
| Nsp16 | PTPN7 | -0.578 | 1.61E-06 |
| S | WNT3 | -0.578 | 1.67E-06 |
| S | FLT4 | -0.578 | 1.65E-06 |
| S | FGF17 | -0.578 | 1.62E-06 |
| S | PKN1 | -0.578 | 1.66E-06 |
| Orf3a | GDF15 | -0.578 | 1.62E-06 |
| Nsp3 | CTF1 | -0.579 | 1.52E-06 |
| Nsp4 | CACNG6 | -0.579 | 1.52E-06 |
| Nsp4 | WNT7B | -0.579 | 1.57E-06 |
| Nsp4 | FGFR3 | -0.579 | 1.52E-06 |
| Nsp8 | BMP8B | -0.579 | 1.53E-06 |
| Nsp8 | ADCY7 | -0.579 | 1.56E-06 |
| Nsp8 | CARD11 | -0.579 | 1.59E-06 |
| Nsp12 | IL17D | -0.579 | 1.55E-06 |
| Nsp12 | MAPK8IP1 | -0.579 | 1.59E-06 |
| Nsp13 | ITGB4 | -0.579 | 1.52E-06 |
| Nsp13 | FMOD | -0.579 | 1.57E-06 |
| Nsp14 | TBKBP1 | -0.579 | 1.52E-06 |

| | | | |
|---|---|---|---|
| Nsp14 | SMAD7 | -0.579 | 1.56E-06 |
| Nsp15 | WNT7B | -0.579 | 1.56E-06 |
| Nsp15 | CAPN1 | -0.579 | 1.57E-06 |
| Nsp16 | CACNA1G | -0.579 | 1.54E-06 |
| S | MAPK8IP1 | -0.579 | 1.56E-06 |
| S | TOLLIP | -0.579 | 1.52E-06 |
| Nsp3 | MAPK11 | -0.58 | 1.50E-06 |
| Nsp4 | LMNB2 | -0.58 | 1.45E-06 |
| Nsp4 | CACNA1B | -0.58 | 1.50E-06 |
| Nsp4 | CTF1 | -0.58 | 1.47E-06 |
| Nsp8 | TRADD | -0.58 | 1.51E-06 |
| Nsp8 | WNT5B | -0.58 | 1.47E-06 |
| Nsp12 | NOS3 | -0.58 | 1.46E-06 |
| Nsp12 | NCF1 | -0.58 | 1.45E-06 |
| Nsp12 | RASGRP2 | -0.58 | 1.51E-06 |
| Nsp13 | PREX1 | -0.58 | 1.46E-06 |
| Nsp13 | BCL3 | -0.58 | 1.49E-06 |
| Nsp16 | RAC2 | -0.58 | 1.51E-06 |
| Nsp16 | PPP2R2C | -0.58 | 1.50E-06 |
| Nsp16 | MAP3K6 | -0.58 | 1.51E-06 |
| S | HSPA1A | -0.58 | 1.47E-06 |
| Nsp2 | SHC2 | -0.581 | 1.38E-06 |
| Nsp2 | SMAD7 | -0.581 | 1.40E-06 |
| Nsp3 | EPOR | -0.581 | 1.41E-06 |
| Nsp4 | JUNB | -0.581 | 1.44E-06 |
| Nsp4 | PIAS4 | -0.581 | 1.37E-06 |
| Nsp6 | GDF15 | -0.581 | 1.42E-06 |
| Nsp8 | WNT3A | -0.581 | 1.40E-06 |
| Nsp8 | RELB | -0.581 | 1.41E-06 |
| Nsp8 | DUSP9 | -0.581 | 1.42E-06 |
| Nsp8 | PIK3CD | -0.581 | 1.44E-06 |
| Nsp12 | RPTOR | -0.581 | 1.37E-06 |
| Nsp13 | IL34 | -0.581 | 1.38E-06 |
| Nsp13 | PRKCG | -0.581 | 1.38E-06 |
| Nsp13 | PRKACG | -0.581 | 1.43E-06 |
| Nsp14 | LEFTY1 | -0.581 | 1.42E-06 |
| Nsp15 | MMP9 | -0.581 | 1.42E-06 |
| Nsp15 | MAPK4 | -0.581 | 1.44E-06 |
| Nsp16 | CACNG1 | -0.581 | 1.40E-06 |
| Nsp16 | HCK | -0.581 | 1.44E-06 |
| Nsp16 | GYS1 | -0.581 | 1.39E-06 |
| Nsp16 | TNFRSF4 | -0.581 | 1.38E-06 |
| Nsp16 | PXN | -0.581 | 1.42E-06 |
| S | FGF18 | -0.581 | 1.38E-06 |
| Nsp3 | RARA | -0.582 | 1.31E-06 |
| Nsp4 | PIK3R2 | -0.582 | 1.36E-06 |
| Nsp4 | ADCY7 | -0.582 | 1.30E-06 |
| Nsp5 | FZD2 | -0.582 | 1.35E-06 |
| Nsp8 | DUSP4 | -0.582 | 1.31E-06 |
| Nsp8 | COL6A2 | -0.582 | 1.34E-06 |

| | | | |
|---|---|---|---|
| Nsp12 | FZD7 | -0.582 | 1.35E-06 |
| Nsp12 | PIAS4 | -0.582 | 1.32E-06 |
| Nsp13 | WNT7B | -0.582 | 1.35E-06 |
| Nsp13 | PIK3R2 | -0.582 | 1.35E-06 |
| Nsp13 | FGF3 | -0.582 | 1.31E-06 |
| Nsp13 | FGF8 | -0.582 | 1.31E-06 |
| Nsp14 | EFNA2 | -0.582 | 1.30E-06 |
| Nsp15 | LEFTY2 | -0.582 | 1.35E-06 |
| Nsp15 | INHBB | -0.582 | 1.34E-06 |
| Nsp15 | DVL1 | -0.582 | 1.35E-06 |
| Nsp15 | GDF11 | -0.582 | 1.31E-06 |
| Nsp16 | SOCS1 | -0.582 | 1.35E-06 |
| Nsp3 | DAB2IP | -0.583 | 1.24E-06 |
| Nsp3 | NOS3 | -0.583 | 1.29E-06 |
| Nsp3 | PTPN6 | -0.583 | 1.29E-06 |
| Nsp3 | COL6A1 | -0.583 | 1.28E-06 |
| Nsp4 | COMP | -0.583 | 1.26E-06 |
| Nsp8 | CACNA1H | -0.583 | 1.28E-06 |
| Nsp12 | CACNA1H | -0.583 | 1.27E-06 |
| Nsp12 | HSPB1 | -0.583 | 1.27E-06 |
| Nsp12 | PPP1R3D | -0.583 | 1.24E-06 |
| Nsp12 | CAPN1 | -0.583 | 1.24E-06 |
| Nsp13 | TELO2 | -0.583 | 1.29E-06 |
| Nsp13 | PFKFB3 | -0.583 | 1.29E-06 |
| Nsp15 | RPTOR | -0.583 | 1.25E-06 |
| Nsp16 | CXCR3 | -0.583 | 1.26E-06 |
| S | CARD11 | -0.583 | 1.27E-06 |
| Orf8 | GDF15 | -0.583 | 1.24E-06 |
| Nsp3 | GADD45B | -0.584 | 1.18E-06 |
| Nsp4 | FZD7 | -0.584 | 1.17E-06 |
| Nsp5 | HSPA2 | -0.584 | 1.21E-06 |
| Nsp5 | LRP5 | -0.584 | 1.20E-06 |
| Nsp5 | TGFB1 | -0.584 | 1.23E-06 |
| Nsp10 | MCL1 | -0.584 | 1.18E-06 |
| Nsp12 | ENDOG | -0.584 | 1.22E-06 |
| Nsp13 | CDC25B | -0.584 | 1.23E-06 |
| Nsp13 | MAPK11 | -0.584 | 1.22E-06 |
| Nsp14 | PRKAR1B | -0.584 | 1.23E-06 |
| Nsp14 | TGFB1 | -0.584 | 1.23E-06 |
| Nsp15 | EFNA3 | -0.584 | 1.23E-06 |
| Nsp15 | TGFB1 | -0.584 | 1.21E-06 |
| Nsp16 | IRF7 | -0.584 | 1.22E-06 |
| Nsp16 | POMC | -0.584 | 1.21E-06 |
| Nsp16 | PRKCG | -0.584 | 1.21E-06 |
| S | RGMA | -0.584 | 1.19E-06 |
| S | EPOR | -0.584 | 1.22E-06 |
| Orf8 | SHC2 | -0.584 | 1.17E-06 |
| Nsp3 | HRK | -0.585 | 1.15E-06 |
| Nsp4 | SLC7A5 | -0.585 | 1.11E-06 |
| Nsp4 | CACNA1A | -0.585 | 1.12E-06 |

| | | | |
|---|---|---|---|
| Nsp4 | ZAP70 | -0.585 | 1.13E-06 |
| Nsp4 | FGF8 | -0.585 | 1.13E-06 |
| Nsp5 | IL17D | -0.585 | 1.14E-06 |
| Nsp8 | COMP | -0.585 | 1.12E-06 |
| Nsp12 | CARD11 | -0.585 | 1.17E-06 |
| Nsp13 | CACNG6 | -0.585 | 1.13E-06 |
| Nsp13 | DUSP4 | -0.585 | 1.12E-06 |
| Nsp16 | SRC | -0.585 | 1.15E-06 |
| Nsp16 | JUNB | -0.585 | 1.14E-06 |
| S | LEFTY2 | -0.585 | 1.12E-06 |
| Orf8 | INHBB | -0.585 | 1.14E-06 |
| Nsp3 | WNT3A | -0.586 | 1.07E-06 |
| Nsp3 | RPTOR | -0.586 | 1.09E-06 |
| Nsp4 | WNT3 | -0.586 | 1.06E-06 |
| Nsp4 | IL17RA | -0.586 | 1.09E-06 |
| Nsp5 | RPS6KA4 | -0.586 | 1.09E-06 |
| Nsp5 | RXRA | -0.586 | 1.10E-06 |
| Nsp5 | ADCY5 | -0.586 | 1.07E-06 |
| Nsp8 | BCL3 | -0.586 | 1.06E-06 |
| Nsp12 | ADCY1 | -0.586 | 1.06E-06 |
| Nsp13 | GNA12 | -0.586 | 1.11E-06 |
| Nsp13 | HRK | -0.586 | 1.11E-06 |
| Nsp14 | SMAD6 | -0.586 | 1.07E-06 |
| Nsp14 | LMNB2 | -0.586 | 1.08E-06 |
| Nsp14 | BMP7 | -0.586 | 1.07E-06 |
| Nsp14 | NFATC1 | -0.586 | 1.11E-06 |
| Nsp15 | GATA3 | -0.586 | 1.09E-06 |
| Nsp15 | GDF1 | -0.586 | 1.07E-06 |
| Nsp16 | PRR5 | -0.586 | 1.10E-06 |
| S | LEFTY1 | -0.586 | 1.09E-06 |
| S | COL6A2 | -0.586 | 1.06E-06 |
| Nsp3 | LAMA5 | -0.587 | 1.02E-06 |
| Nsp3 | EGLN1 | -0.587 | 1.04E-06 |
| Nsp4 | IL17D | -0.587 | 1.03E-06 |
| Nsp4 | ENDOG | -0.587 | 1.03E-06 |
| Nsp4 | CAPN1 | -0.587 | 1.05E-06 |
| Nsp5 | LMNB2 | -0.587 | 1.05E-06 |
| Nsp8 | PRR5 | -0.587 | 1.02E-06 |
| Nsp8 | IL17RA | -0.587 | 1.02E-06 |
| Nsp8 | FGFR3 | -0.587 | 1.04E-06 |
| Nsp12 | FLT4 | -0.587 | 1.03E-06 |
| Nsp12 | COMP | -0.587 | 1.04E-06 |
| Nsp12 | ADCY3 | -0.587 | 1.03E-06 |
| Nsp15 | FZD7 | -0.587 | 1.05E-06 |
| Nsp15 | FZD2 | -0.587 | 1.04E-06 |
| Nsp16 | DUSP4 | -0.587 | 1.05E-06 |
| Nsp16 | MKNK2 | -0.587 | 1.02E-06 |
| Orf8 | FGF4 | -0.587 | 1.04E-06 |
| Nsp2 | PPP2R3B | -0.588 | 9.73E-07 |
| Nsp3 | MMP9 | -0.588 | 9.94E-07 |

| | | | |
|---|---|---|---|
| Nsp3 | CACNA1B | -0.588 | 9.56E-07 |
| Nsp4 | STK11 | -0.588 | 9.50E-07 |
| Nsp5 | GRK2 | -0.588 | 9.67E-07 |
| Nsp5 | GDF6 | -0.588 | 9.53E-07 |
| Nsp12 | LEFTY2 | -0.588 | 9.73E-07 |
| Nsp12 | PTPN6 | -0.588 | 9.61E-07 |
| Nsp13 | DUSP1 | -0.588 | 9.74E-07 |
| Nsp13 | RARA | -0.588 | 9.52E-07 |
| Nsp15 | RPS6KA4 | -0.588 | 9.75E-07 |
| Nsp15 | NCF1 | -0.588 | 9.81E-07 |
| Nsp15 | ZAP70 | -0.588 | 9.98E-07 |
| Nsp16 | RPTOR | -0.588 | 9.56E-07 |
| Nsp16 | RELB | -0.588 | 9.87E-07 |
| S | BMP8B | -0.588 | 9.60E-07 |
| S | IL17RA | -0.588 | 9.63E-07 |
| S | BCL2 | -0.588 | 9.80E-07 |
| Nsp3 | MAP2K2 | -0.589 | 9.07E-07 |
| Nsp3 | BCL2 | -0.589 | 9.46E-07 |
| Nsp4 | FZD8 | -0.589 | 9.12E-07 |
| Nsp5 | JUNB | -0.589 | 9.18E-07 |
| Nsp8 | GATA3 | -0.589 | 9.24E-07 |
| Nsp12 | HSPA2 | -0.589 | 9.15E-07 |
| Nsp12 | DUSP5 | -0.589 | 9.37E-07 |
| Nsp12 | PIK3R2 | -0.589 | 9.39E-07 |
| Nsp12 | DUSP9 | -0.589 | 9.43E-07 |
| Nsp12 | ADCY7 | -0.589 | 9.26E-07 |
| Nsp13 | NOS3 | -0.589 | 9.27E-07 |
| Nsp13 | DUSP6 | -0.589 | 9.19E-07 |
| Nsp15 | NOG | -0.589 | 9.12E-07 |
| Nsp16 | CACNG7 | -0.589 | 9.42E-07 |
| Nsp16 | CHAD | -0.589 | 9.13E-07 |
| Nsp16 | MAP2K2 | -0.589 | 9.02E-07 |
| Nsp16 | FGF8 | -0.589 | 9.43E-07 |
| S | CARD14 | -0.589 | 9.41E-07 |
| Orf8 | GDF7 | -0.589 | 9.33E-07 |
| Nsp3 | RGMA | -0.59 | 8.72E-07 |
| Nsp4 | PPP1R3D | -0.59 | 8.61E-07 |
| Nsp4 | MYC | -0.59 | 8.82E-07 |
| Nsp4 | PIK3CD | -0.59 | 8.65E-07 |
| Nsp4 | NCF1 | -0.59 | 8.77E-07 |
| Nsp4 | BCL2 | -0.59 | 8.68E-07 |
| Nsp5 | FZD7 | -0.59 | 8.94E-07 |
| Nsp5 | PIK3R2 | -0.59 | 8.77E-07 |
| Nsp5 | WNT5B | -0.59 | 8.76E-07 |
| Nsp5 | JUND | -0.59 | 8.88E-07 |
| Nsp8 | SH2B2 | -0.59 | 8.54E-07 |
| Nsp8 | COL6A1 | -0.59 | 8.74E-07 |
| Nsp12 | FZD5 | -0.59 | 8.88E-07 |
| Nsp12 | GDF1 | -0.59 | 8.59E-07 |
| Nsp13 | JUND | -0.59 | 8.91E-07 |

| | | | |
|---|---|---|---|
| Nsp13 | ADCY3 | -0.59 | 8.78E-07 |
| Nsp13 | FGF22 | -0.59 | 8.90E-07 |
| Nsp14 | RXRA | -0.59 | 8.98E-07 |
| Nsp14 | CAPN1 | -0.59 | 8.93E-07 |
| Nsp15 | CACNA1H | -0.59 | 8.77E-07 |
| Nsp15 | LEFTY1 | -0.59 | 8.54E-07 |
| Nsp15 | RASGRP2 | -0.59 | 8.86E-07 |
| Nsp15 | ADCY5 | -0.59 | 8.99E-07 |
| Nsp16 | PGAM5 | -0.59 | 8.94E-07 |
| Nsp16 | ADCY4 | -0.59 | 8.91E-07 |
| S | SMAD7 | -0.59 | 8.75E-07 |
| Nsp3 | SLC7A5 | -0.591 | 8.27E-07 |
| Nsp3 | CACNA1A | -0.591 | 8.48E-07 |
| Nsp4 | GDF6 | -0.591 | 8.51E-07 |
| Nsp5 | GDF10 | -0.591 | 8.23E-07 |
| Nsp5 | PIK3CD | -0.591 | 8.10E-07 |
| Nsp5 | WNT1 | -0.591 | 8.19E-07 |
| Nsp8 | DUSP6 | -0.591 | 8.43E-07 |
| Nsp12 | CACNG1 | -0.591 | 8.23E-07 |
| Nsp12 | INSR | -0.591 | 8.17E-07 |
| Nsp12 | CACNA1B | -0.591 | 8.29E-07 |
| Nsp12 | CCND1 | -0.591 | 8.30E-07 |
| Nsp13 | COL6A2 | -0.591 | 8.18E-07 |
| Nsp14 | GDF6 | -0.591 | 8.47E-07 |
| Nsp14 | MAPK8IP2 | -0.591 | 8.34E-07 |
| Nsp14 | PPP2R3B | -0.591 | 8.24E-07 |
| Nsp14 | FZD9 | -0.591 | 8.10E-07 |
| S | MAPK11 | -0.591 | 8.11E-07 |
| S | DUSP6 | -0.591 | 8.50E-07 |
| Orf8 | MCL1 | -0.591 | 8.21E-07 |
| Nsp4 | PGAM5 | -0.592 | 8.03E-07 |
| Nsp4 | MAP2K2 | -0.592 | 7.72E-07 |
| Nsp5 | BMP8B | -0.592 | 7.78E-07 |
| Nsp8 | RGMA | -0.592 | 7.68E-07 |
| Nsp8 | BMP6 | -0.592 | 7.97E-07 |
| Nsp8 | GDF10 | -0.592 | 7.79E-07 |
| Nsp8 | FLT4 | -0.592 | 8.06E-07 |
| Nsp8 | GADD45B | -0.592 | 7.99E-07 |
| Nsp8 | VEGFA | -0.592 | 7.81E-07 |
| Nsp12 | LEFTY1 | -0.592 | 7.95E-07 |
| Nsp13 | ENDOG | -0.592 | 7.75E-07 |
| Nsp13 | GRK1 | -0.592 | 8.08E-07 |
| Nsp13 | SH2B2 | -0.592 | 7.99E-07 |
| Nsp14 | DUSP8 | -0.592 | 7.83E-07 |
| Nsp16 | IL17D | -0.592 | 7.82E-07 |
| S | CCR10 | -0.592 | 7.84E-07 |
| Nsp3 | PRR5 | -0.593 | 7.31E-07 |
| Nsp3 | FGF22 | -0.593 | 7.44E-07 |
| Nsp5 | CACNA1I | -0.593 | 7.40E-07 |
| Nsp5 | ZAP70 | -0.593 | 7.47E-07 |

| | | | |
|---|---|---|---|
| Nsp8 | PIAS4 | -0.593 | 7.37E-07 |
| Nsp8 | MAPK8IP3 | -0.593 | 7.54E-07 |
| Nsp8 | CAPN1 | -0.593 | 7.38E-07 |
| Nsp12 | FZD8 | -0.593 | 7.27E-07 |
| Nsp12 | WNT3 | -0.593 | 7.40E-07 |
| Nsp12 | BCL2 | -0.593 | 7.63E-07 |
| Nsp14 | RPS6KA4 | -0.593 | 7.29E-07 |
| Nsp14 | NOG | -0.593 | 7.32E-07 |
| Nsp15 | GDF6 | -0.593 | 7.27E-07 |
| Nsp15 | TNFRSF6B | -0.593 | 7.47E-07 |
| Nsp16 | WNT5B | -0.593 | 7.54E-07 |
| Nsp16 | ADCY7 | -0.593 | 7.47E-07 |
| S | CACNA1H | -0.593 | 7.59E-07 |
| S | PIM1 | -0.593 | 7.64E-07 |
| Nsp3 | RELB | -0.594 | 7.00E-07 |
| Nsp3 | COL6A2 | -0.594 | 7.05E-07 |
| Nsp5 | GDF11 | -0.594 | 6.95E-07 |
| Nsp8 | MAPK3 | -0.594 | 7.07E-07 |
| Nsp8 | SLC7A5 | -0.594 | 6.99E-07 |
| Nsp12 | WNT7B | -0.594 | 7.07E-07 |
| Nsp12 | DUSP7 | -0.594 | 7.10E-07 |
| Nsp13 | LMNB2 | -0.594 | 7.23E-07 |
| Nsp14 | DDIT4 | -0.594 | 6.99E-07 |
| Nsp15 | BMP7 | -0.594 | 7.12E-07 |
| Nsp16 | XCR1 | -0.594 | 7.04E-07 |
| S | PRR5 | -0.594 | 7.02E-07 |
| S | POMC | -0.594 | 7.18E-07 |
| Orf8 | CACNG8 | -0.594 | 7.25E-07 |
| Nsp3 | PIAS4 | -0.595 | 6.62E-07 |
| Nsp3 | SH2B2 | -0.595 | 6.62E-07 |
| Nsp4 | FZD5 | -0.595 | 6.86E-07 |
| Nsp5 | FZD8 | -0.595 | 6.74E-07 |
| Nsp8 | LMNB2 | -0.595 | 6.67E-07 |
| Nsp8 | DUSP5 | -0.595 | 6.79E-07 |
| Nsp8 | WNT1 | -0.595 | 6.72E-07 |
| Nsp8 | FGF8 | -0.595 | 6.63E-07 |
| Nsp12 | MMP9 | -0.595 | 6.66E-07 |
| Nsp13 | MAPK8IP1 | -0.595 | 6.69E-07 |
| Nsp15 | PPP1R3D | -0.595 | 6.64E-07 |
| Nsp16 | PLCG1 | -0.595 | 6.53E-07 |
| S | WNT3A | -0.595 | 6.83E-07 |
| Nsp2 | EFNA2 | -0.596 | 6.19E-07 |
| Nsp3 | CD70 | -0.596 | 6.46E-07 |
| Nsp5 | EFNA3 | -0.596 | 6.39E-07 |
| Nsp5 | INHBB | -0.596 | 6.35E-07 |
| Nsp5 | AMH | -0.596 | 6.34E-07 |
| Nsp12 | SOCS1 | -0.596 | 6.50E-07 |
| Nsp12 | MYC | -0.596 | 6.23E-07 |
| Nsp13 | CAPN2 | -0.596 | 6.45E-07 |
| Nsp14 | WNT6 | -0.596 | 6.43E-07 |

| | | | |
|---|---|---|---|
| Nsp15 | GNA12 | -0.596 | 6.27E-07 |
| Nsp15 | BMP8B | -0.596 | 6.38E-07 |
| Nsp16 | HSPB1 | -0.596 | 6.39E-07 |
| Nsp16 | DUSP5 | -0.596 | 6.34E-07 |
| Nsp16 | MMP9 | -0.596 | 6.48E-07 |
| Nsp16 | FGFR3 | -0.596 | 6.48E-07 |
| Nsp16 | CARD11 | -0.596 | 6.23E-07 |
| Nsp3 | GATA3 | -0.597 | 5.90E-07 |
| Nsp4 | WNT5B | -0.597 | 6.05E-07 |
| Nsp8 | CCND1 | -0.597 | 6.12E-07 |
| Nsp13 | TNFRSF4 | -0.597 | 6.00E-07 |
| Nsp15 | COMP | -0.597 | 6.14E-07 |
| Nsp15 | WNT5B | -0.597 | 6.04E-07 |
| Nsp15 | IL17RA | -0.597 | 5.87E-07 |
| Nsp16 | PRKAR1B | -0.597 | 5.96E-07 |
| Nsp16 | WNT3A | -0.597 | 6.05E-07 |
| S | BMP8A | -0.597 | 6.00E-07 |
| S | PIK3CD | -0.597 | 6.10E-07 |
| Nsp3 | BCL3 | -0.598 | 5.59E-07 |
| Nsp4 | HSPA2 | -0.598 | 5.81E-07 |
| Nsp4 | CARD14 | -0.598 | 5.74E-07 |
| Nsp4 | RPS6KA4 | -0.598 | 5.80E-07 |
| Nsp4 | DVL1 | -0.598 | 5.70E-07 |
| Nsp4 | AMH | -0.598 | 5.82E-07 |
| Nsp4 | RASGRP2 | -0.598 | 5.85E-07 |
| Nsp4 | FGF22 | -0.598 | 5.75E-07 |
| Nsp5 | FZD5 | -0.598 | 5.78E-07 |
| Nsp5 | FGF22 | -0.598 | 5.77E-07 |
| Nsp8 | DVL1 | -0.598 | 5.58E-07 |
| Nsp12 | CACNA1A | -0.598 | 5.59E-07 |
| Nsp13 | GRK2 | -0.598 | 5.54E-07 |
| Nsp13 | MMP9 | -0.598 | 5.62E-07 |
| Nsp13 | COMP | -0.598 | 5.67E-07 |
| Nsp14 | INHBB | -0.598 | 5.65E-07 |
| Nsp15 | BMP6 | -0.598 | 5.63E-07 |
| Nsp4 | SMAD7 | -0.599 | 5.53E-07 |
| Nsp5 | NCF1 | -0.599 | 5.36E-07 |
| Nsp8 | JUND | -0.599 | 5.48E-07 |
| Nsp12 | PRR5 | -0.599 | 5.37E-07 |
| Nsp12 | IL17RA | -0.599 | 5.32E-07 |
| Nsp12 | ADCY5 | -0.599 | 5.32E-07 |
| Nsp13 | CACNA1H | -0.599 | 5.30E-07 |
| Nsp13 | CACNG7 | -0.599 | 5.29E-07 |
| Nsp13 | WNT5B | -0.599 | 5.53E-07 |
| Nsp13 | CALML5 | -0.599 | 5.35E-07 |
| Nsp14 | STK11 | -0.599 | 5.36E-07 |
| Nsp15 | IL17D | -0.599 | 5.28E-07 |
| Nsp16 | ITGB4 | -0.599 | 5.44E-07 |
| Nsp16 | FLT4 | -0.599 | 5.32E-07 |
| Nsp16 | MAPK4 | -0.599 | 5.36E-07 |

| | | | |
|---|---|---|---|
| S | RASGRP2 | -0.599 | 5.44E-07 |
| Nsp3 | CACNG1 | -0.6 | 4.98E-07 |
| Nsp3 | GDF6 | -0.6 | 5.01E-07 |
| Nsp3 | ADCY7 | -0.6 | 5.11E-07 |
| Nsp5 | GDF7 | -0.6 | 5.16E-07 |
| Nsp5 | PPP1R3D | -0.6 | 5.19E-07 |
| Nsp8 | HRK | -0.6 | 5.08E-07 |
| Nsp12 | CARD14 | -0.6 | 5.09E-07 |
| Nsp13 | HSPA1B | -0.6 | 5.10E-07 |
| Nsp14 | CEBPB | -0.6 | 5.12E-07 |
| Nsp15 | HSPA2 | -0.6 | 5.15E-07 |
| Nsp3 | CCND1 | -0.601 | 4.71E-07 |
| Nsp3 | IL17RA | -0.601 | 4.71E-07 |
| Nsp5 | BMP8A | -0.601 | 4.84E-07 |
| Nsp5 | DUSP9 | -0.601 | 4.75E-07 |
| Nsp14 | CACNG8 | -0.601 | 4.96E-07 |
| Nsp16 | RPS6KA4 | -0.601 | 4.79E-07 |
| Nsp16 | NOS3 | -0.601 | 4.78E-07 |
| Nsp16 | FOXO3 | -0.601 | 4.91E-07 |
| S | DUSP4 | -0.601 | 4.90E-07 |
| Nsp3 | FLT4 | -0.602 | 4.46E-07 |
| Nsp3 | MYC | -0.602 | 4.59E-07 |
| Nsp3 | FGFR3 | -0.602 | 4.58E-07 |
| Nsp4 | BMP8B | -0.602 | 4.48E-07 |
| Nsp8 | TNFRSF6B | -0.602 | 4.57E-07 |
| Nsp13 | CACNG4 | -0.602 | 4.50E-07 |
| Nsp16 | TAB1 | -0.602 | 4.45E-07 |
| Nsp16 | NFATC1 | -0.602 | 4.60E-07 |
| Nsp16 | IL17RA | -0.602 | 4.65E-07 |
| Nsp3 | LMNB2 | -0.603 | 4.26E-07 |
| Nsp3 | DUSP9 | -0.603 | 4.41E-07 |
| Nsp5 | ENDOG | -0.603 | 4.36E-07 |
| Nsp5 | RASGRP2 | -0.603 | 4.45E-07 |
| Nsp8 | ENDOG | -0.603 | 4.32E-07 |
| Nsp8 | TBKBP1 | -0.603 | 4.27E-07 |
| Nsp13 | PHLPP1 | -0.603 | 4.30E-07 |
| Nsp14 | FZD10 | -0.603 | 4.41E-07 |
| Nsp14 | LPAR5 | -0.603 | 4.23E-07 |
| Nsp14 | JUN | -0.603 | 4.21E-07 |
| Nsp14 | FZD1 | -0.603 | 4.42E-07 |
| Nsp15 | LRP5 | -0.603 | 4.23E-07 |
| Nsp16 | LEFTY2 | -0.603 | 4.41E-07 |
| S | PRKAR1B | -0.603 | 4.35E-07 |
| S | CACNG7 | -0.603 | 4.31E-07 |
| Nsp4 | GREM2 | -0.604 | 4.08E-07 |
| Nsp8 | SMAD7 | -0.604 | 4.08E-07 |
| Nsp12 | BMP8A | -0.604 | 4.06E-07 |
| Nsp12 | HRK | -0.604 | 4.00E-07 |
| Nsp13 | FOSB | -0.604 | 4.01E-07 |
| Nsp13 | CACNG1 | -0.604 | 4.00E-07 |

| | | | |
|---|---|---|---|
| Nsp13 | CACNA1A | -0.604 | 4.13E-07 |
| Nsp14 | EFNA3 | -0.604 | 4.10E-07 |
| Nsp14 | LRP5 | -0.604 | 4.02E-07 |
| Nsp16 | PTPN6 | -0.604 | 3.98E-07 |
| S | GADD45B | -0.604 | 3.99E-07 |
| S | MAPK8IP3 | -0.604 | 4.07E-07 |
| S | FGF8 | -0.604 | 3.99E-07 |
| Nsp3 | PIK3R2 | -0.605 | 3.94E-07 |
| Nsp3 | CARD11 | -0.605 | 3.93E-07 |
| Nsp4 | INHBB | -0.605 | 3.83E-07 |
| Nsp4 | CACNA1I | -0.605 | 3.85E-07 |
| Nsp8 | RARA | -0.605 | 3.93E-07 |
| Nsp12 | DVL1 | -0.605 | 3.81E-07 |
| Nsp13 | RPTOR | -0.605 | 3.80E-07 |
| Nsp13 | PPP1R3D | -0.605 | 3.93E-07 |
| Nsp13 | HSPA1A | -0.605 | 3.86E-07 |
| Nsp13 | ADCY7 | -0.605 | 3.80E-07 |
| Nsp14 | PIM1 | -0.605 | 3.79E-07 |
| Nsp15 | SMAD7 | -0.605 | 3.77E-07 |
| Nsp15 | FZD5 | -0.605 | 3.77E-07 |
| Nsp16 | PPP1R3E | -0.605 | 3.97E-07 |
| Nsp16 | COL6A2 | -0.605 | 3.92E-07 |
| Nsp16 | LAMA5 | -0.605 | 3.97E-07 |
| Nsp2 | FGF4 | -0.606 | 3.72E-07 |
| Nsp3 | CACNG7 | -0.606 | 3.59E-07 |
| Nsp3 | TNFRSF6B | -0.606 | 3.63E-07 |
| Nsp4 | DDIT4 | -0.606 | 3.57E-07 |
| Nsp4 | LEFTY2 | -0.606 | 3.76E-07 |
| Nsp4 | BMP7 | -0.606 | 3.62E-07 |
| Nsp5 | WNT6 | -0.606 | 3.66E-07 |
| Nsp5 | BMP6 | -0.606 | 3.56E-07 |
| Nsp5 | GDF1 | -0.606 | 3.65E-07 |
| Nsp12 | CACNA1I | -0.606 | 3.76E-07 |
| Nsp13 | ADCY9 | -0.606 | 3.68E-07 |
| Nsp13 | AKT2 | -0.606 | 3.60E-07 |
| Nsp13 | WNT1 | -0.606 | 3.63E-07 |
| Nsp15 | CACNA1I | -0.606 | 3.71E-07 |
| Nsp16 | ENDOG | -0.606 | 3.74E-07 |
| Nsp16 | ADCY3 | -0.606 | 3.57E-07 |
| S | PIK3R2 | -0.606 | 3.58E-07 |
| S | PIAS4 | -0.606 | 3.71E-07 |
| S | CCND1 | -0.606 | 3.56E-07 |
| S | FGFR3 | -0.606 | 3.67E-07 |
| Nsp2 | TNFRSF18 | -0.607 | 3.39E-07 |
| Nsp3 | MAPK8IP3 | -0.607 | 3.55E-07 |
| Nsp4 | HSPB1 | -0.607 | 3.40E-07 |
| Nsp8 | ADCY5 | -0.607 | 3.46E-07 |
| Nsp12 | PIK3CD | -0.607 | 3.52E-07 |
| Nsp12 | AMH | -0.607 | 3.43E-07 |
| Nsp13 | PIK3CD | -0.607 | 3.52E-07 |

| | | | |
|---|---|---|---|
| Nsp16 | DAB2IP | -0.607 | 3.55E-07 |
| Nsp16 | LMNB2 | -0.607 | 3.52E-07 |
| Nsp16 | LEFTY1 | -0.607 | 3.37E-07 |
| S | CDKN2B | -0.607 | 3.44E-07 |
| Nsp3 | ADCY3 | -0.608 | 3.29E-07 |
| Nsp8 | PIK3R2 | -0.608 | 3.36E-07 |
| Nsp12 | BMP7 | -0.608 | 3.27E-07 |
| Nsp12 | BMP8B | -0.608 | 3.24E-07 |
| Nsp12 | PIM1 | -0.608 | 3.20E-07 |
| Nsp13 | FZD7 | -0.608 | 3.30E-07 |
| Nsp13 | WNT3 | -0.608 | 3.26E-07 |
| Nsp13 | FLT4 | -0.608 | 3.27E-07 |
| Nsp13 | CACNA1B | -0.608 | 3.30E-07 |
| Nsp13 | RASGRP2 | -0.608 | 3.28E-07 |
| Nsp15 | GREM2 | -0.608 | 3.23E-07 |
| Nsp16 | PIAS4 | -0.608 | 3.20E-07 |
| S | IL17D | -0.608 | 3.25E-07 |
| S | PGAM5 | -0.608 | 3.27E-07 |
| S | DUSP9 | -0.608 | 3.22E-07 |
| Orf8 | CEBPB | -0.608 | 3.29E-07 |
| Nsp4 | DUSP7 | -0.609 | 3.08E-07 |
| Nsp4 | TGFB1 | -0.609 | 3.09E-07 |
| Nsp12 | WNT6 | -0.609 | 3.12E-07 |
| Nsp12 | INHBB | -0.609 | 3.04E-07 |
| Nsp12 | STK11 | -0.609 | 3.03E-07 |
| Nsp15 | SMAD6 | -0.609 | 3.10E-07 |
| Nsp15 | DUSP5 | -0.609 | 3.11E-07 |
| Nsp15 | DUSP7 | -0.609 | 3.01E-07 |
| S | ADCY5 | -0.609 | 3.09E-07 |
| Nsp3 | CAPN1 | -0.61 | 2.97E-07 |
| Nsp3 | STK11 | -0.61 | 2.92E-07 |
| Nsp4 | LEFTY1 | -0.61 | 2.87E-07 |
| Nsp8 | IL17D | -0.61 | 2.89E-07 |
| Nsp13 | IL17RA | -0.61 | 3.00E-07 |
| Nsp14 | MAP2K7 | -0.61 | 2.87E-07 |
| Nsp15 | BMP8A | -0.61 | 2.86E-07 |
| Nsp16 | PIK3R2 | -0.61 | 2.87E-07 |
| S | COMP | -0.61 | 2.84E-07 |
| S | RARA | -0.61 | 2.89E-07 |
| S | SH2B2 | -0.61 | 2.97E-07 |
| Nsp3 | CACNA1H | -0.611 | 2.74E-07 |
| Nsp3 | SMAD7 | -0.611 | 2.80E-07 |
| Nsp3 | PGAM5 | -0.611 | 2.83E-07 |
| Nsp3 | PPP1R3D | -0.611 | 2.83E-07 |
| Nsp3 | PIK3CD | -0.611 | 2.81E-07 |
| Nsp4 | TBKBP1 | -0.611 | 2.71E-07 |
| Nsp4 | GATA3 | -0.611 | 2.73E-07 |
| Nsp4 | GDF1 | -0.611 | 2.69E-07 |
| Nsp5 | DUSP7 | -0.611 | 2.69E-07 |
| Nsp8 | LRP5 | -0.611 | 2.83E-07 |

| | | | |
|---|---|---|---|
| Nsp13 | FZD2 | -0.611 | 2.69E-07 |
| Nsp16 | PPP1R3D | -0.611 | 2.73E-07 |
| Nsp16 | PIM1 | -0.611 | 2.69E-07 |
| Nsp16 | MAP2K7 | -0.611 | 2.71E-07 |
| S | TNFRSF6B | -0.611 | 2.77E-07 |
| Nsp3 | FZD7 | -0.612 | 2.58E-07 |
| Nsp3 | JUND | -0.612 | 2.55E-07 |
| Nsp3 | RASGRP2 | -0.612 | 2.67E-07 |
| Nsp8 | WNT3 | -0.612 | 2.57E-07 |
| Nsp8 | PIM1 | -0.612 | 2.65E-07 |
| Nsp8 | NFATC1 | -0.612 | 2.60E-07 |
| Nsp12 | LRP5 | -0.612 | 2.53E-07 |
| Nsp15 | AMH | -0.612 | 2.58E-07 |
| Nsp16 | BMP7 | -0.612 | 2.54E-07 |
| Nsp16 | GDF1 | -0.612 | 2.68E-07 |
| Nsp16 | ADCY5 | -0.612 | 2.63E-07 |
| S | WNT5B | -0.612 | 2.57E-07 |
| S | PPP1R3E | -0.612 | 2.63E-07 |
| Nsp3 | JUNB | -0.613 | 2.53E-07 |
| Nsp3 | GDF11 | -0.613 | 2.43E-07 |
| Nsp3 | ADCY5 | -0.613 | 2.51E-07 |
| Nsp4 | SOCS1 | -0.613 | 2.52E-07 |
| Nsp8 | GRK2 | -0.613 | 2.49E-07 |
| Nsp8 | GDF6 | -0.613 | 2.46E-07 |
| Nsp8 | JUN | -0.613 | 2.48E-07 |
| Nsp8 | NCF1 | -0.613 | 2.41E-07 |
| Nsp8 | ZAP70 | -0.613 | 2.41E-07 |
| Nsp13 | HSPA2 | -0.613 | 2.41E-07 |
| Nsp13 | PIAS4 | -0.613 | 2.50E-07 |
| Nsp14 | IRS2 | -0.613 | 2.44E-07 |
| Nsp16 | EFNA3 | -0.613 | 2.48E-07 |
| S | EGLN1 | -0.613 | 2.47E-07 |
| Nsp3 | LEFTY1 | -0.614 | 2.33E-07 |
| Nsp13 | IL17D | -0.614 | 2.35E-07 |
| Nsp13 | PRR5 | -0.614 | 2.31E-07 |
| Nsp13 | BMP8B | -0.614 | 2.36E-07 |
| Nsp13 | SLC7A5 | -0.614 | 2.30E-07 |
| Nsp15 | MYC | -0.614 | 2.27E-07 |
| S | RXRA | -0.614 | 2.31E-07 |
| Nsp3 | IL17D | -0.615 | 2.13E-07 |
| Nsp3 | LEFTY2 | -0.615 | 2.21E-07 |
| Nsp3 | INHBB | -0.615 | 2.21E-07 |
| Nsp3 | MAPK4 | -0.615 | 2.15E-07 |
| Nsp4 | NFATC1 | -0.615 | 2.16E-07 |
| Nsp4 | NOG | -0.615 | 2.21E-07 |
| Nsp5 | DDIT4 | -0.615 | 2.25E-07 |
| Nsp5 | EFNA2 | -0.615 | 2.16E-07 |
| Nsp15 | CARD14 | -0.615 | 2.14E-07 |
| Nsp16 | TNXB | -0.615 | 2.25E-07 |
| Nsp16 | WNT9A | -0.615 | 2.13E-07 |

| | | | |
|---|---|---|---|
| Nsp16 | FZD2 | -0.615 | 2.14E-07 |
| S | WNT7B | -0.615 | 2.20E-07 |
| Nsp5 | HSPB1 | -0.616 | 2.01E-07 |
| Nsp12 | EFNA3 | -0.616 | 2.09E-07 |
| Nsp13 | CARD11 | -0.616 | 2.08E-07 |
| Nsp14 | BMP6 | -0.616 | 2.11E-07 |
| Nsp16 | CARD14 | -0.616 | 2.11E-07 |
| Nsp16 | BMP6 | -0.616 | 2.07E-07 |
| Nsp16 | DUSP9 | -0.616 | 2.08E-07 |
| Nsp16 | PPP2R3B | -0.616 | 2.11E-07 |
| Nsp3 | TBKBP1 | -0.617 | 1.94E-07 |
| Nsp3 | WNT5B | -0.617 | 2.00E-07 |
| Nsp4 | BMP8A | -0.617 | 1.89E-07 |
| Nsp4 | PIM1 | -0.617 | 1.94E-07 |
| Nsp5 | DUSP8 | -0.617 | 1.98E-07 |
| Nsp5 | IRS2 | -0.617 | 1.98E-07 |
| Nsp5 | SLC7A5 | -0.617 | 1.89E-07 |
| Nsp5 | FZD1 | -0.617 | 1.94E-07 |
| Nsp12 | EFNA2 | -0.617 | 1.91E-07 |
| Nsp13 | FZD5 | -0.617 | 1.98E-07 |
| Nsp13 | GDF1 | -0.617 | 1.95E-07 |
| Nsp16 | BMP8A | -0.617 | 1.92E-07 |
| S | MAP2K7 | -0.617 | 2.01E-07 |
| S | STK11 | -0.617 | 1.98E-07 |
| S | FGF22 | -0.617 | 1.92E-07 |
| Nsp5 | PPP2R3B | -0.618 | 1.80E-07 |
| Nsp5 | NFATC1 | -0.618 | 1.84E-07 |
| Nsp5 | MAP2K7 | -0.618 | 1.79E-07 |
| Nsp8 | BMP7 | -0.618 | 1.81E-07 |
| Nsp13 | MAPK4 | -0.618 | 1.85E-07 |
| Nsp15 | FZD9 | -0.618 | 1.88E-07 |
| Nsp16 | CACNA1H | -0.618 | 1.80E-07 |
| Nsp16 | BMP8B | -0.618 | 1.85E-07 |
| S | HRK | -0.618 | 1.88E-07 |
| S | NFATC1 | -0.618 | 1.88E-07 |
| S | CAPN1 | -0.618 | 1.80E-07 |
| Nsp3 | WNT7B | -0.619 | 1.78E-07 |
| Nsp4 | GDF7 | -0.619 | 1.77E-07 |
| Nsp8 | FZD7 | -0.619 | 1.69E-07 |
| Nsp8 | JUNB | -0.619 | 1.72E-07 |
| Nsp8 | MAP2K7 | -0.619 | 1.70E-07 |
| Nsp12 | RPS6KA4 | -0.619 | 1.75E-07 |
| Nsp13 | INHBB | -0.619 | 1.70E-07 |
| Nsp16 | TBKBP1 | -0.619 | 1.75E-07 |
| Nsp16 | GDF10 | -0.619 | 1.75E-07 |
| Nsp16 | NCF1 | -0.619 | 1.78E-07 |
| Nsp16 | RARA | -0.619 | 1.76E-07 |
| Nsp16 | NOG | -0.619 | 1.78E-07 |
| Nsp2 | JUN | -0.62 | 1.68E-07 |
| Nsp3 | ENDOG | -0.62 | 1.68E-07 |

| | | | |
|---|---|---|---|
| Nsp3 | FZD2 | -0.62 | 1.65E-07 |
| Nsp4 | SMAD6 | -0.62 | 1.68E-07 |
| Nsp8 | GDF11 | -0.62 | 1.59E-07 |
| Nsp12 | TBKBP1 | -0.62 | 1.62E-07 |
| Nsp12 | FOSB | -0.62 | 1.68E-07 |
| Nsp12 | GDF7 | -0.62 | 1.60E-07 |
| Nsp13 | LEFTY2 | -0.62 | 1.63E-07 |
| Nsp13 | CCND1 | -0.62 | 1.60E-07 |
| Nsp15 | MAP2K7 | -0.62 | 1.60E-07 |
| Nsp16 | COMP | -0.62 | 1.64E-07 |
| S | BMP6 | -0.62 | 1.66E-07 |
| S | GDF6 | -0.62 | 1.61E-07 |
| S | INHBB | -0.62 | 1.65E-07 |
| S | MAPK4 | -0.62 | 1.60E-07 |
| Nsp3 | BMP8B | -0.621 | 1.53E-07 |
| Nsp3 | FGF8 | -0.621 | 1.54E-07 |
| Nsp4 | EFNA2 | -0.621 | 1.55E-07 |
| Nsp5 | DVL1 | -0.621 | 1.57E-07 |
| Nsp8 | HSPA2 | -0.621 | 1.54E-07 |
| Nsp8 | TNFRSF18 | -0.621 | 1.58E-07 |
| Nsp13 | GDF6 | -0.621 | 1.55E-07 |
| Nsp15 | LPAR5 | -0.621 | 1.58E-07 |
| Nsp16 | FZD8 | -0.621 | 1.55E-07 |
| Nsp16 | INHBB | -0.621 | 1.51E-07 |
| S | GDF11 | -0.621 | 1.55E-07 |
| Nsp4 | PRKAR1B | -0.622 | 1.46E-07 |
| Nsp4 | LRP5 | -0.622 | 1.46E-07 |
| Nsp5 | SOCS1 | -0.622 | 1.47E-07 |
| Nsp5 | TBKBP1 | -0.622 | 1.48E-07 |
| Nsp5 | NOG | -0.622 | 1.43E-07 |
| Nsp8 | PRKAR1B | -0.622 | 1.47E-07 |
| Nsp12 | FZD9 | -0.622 | 1.43E-07 |
| Nsp12 | MAP2K7 | -0.622 | 1.47E-07 |
| Nsp13 | MCL1 | -0.622 | 1.47E-07 |
| Nsp13 | DVL1 | -0.622 | 1.49E-07 |
| Nsp13 | DUSP9 | -0.622 | 1.45E-07 |
| Nsp13 | FGFR3 | -0.622 | 1.45E-07 |
| Nsp16 | FZD7 | -0.622 | 1.44E-07 |
| S | FZD7 | -0.622 | 1.47E-07 |
| S | HSPB1 | -0.622 | 1.44E-07 |
| Nsp4 | EFNA3 | -0.623 | 1.35E-07 |
| Nsp4 | FZD9 | -0.623 | 1.38E-07 |
| Nsp5 | BMP7 | -0.623 | 1.38E-07 |
| Nsp15 | WNT6 | -0.623 | 1.35E-07 |
| Nsp15 | PIM1 | -0.623 | 1.37E-07 |
| Nsp16 | TNFRSF18 | -0.623 | 1.38E-07 |
| Nsp16 | GADD45B | -0.623 | 1.35E-07 |
| Nsp16 | PIK3CD | -0.623 | 1.36E-07 |
| Nsp3 | GRK2 | -0.624 | 1.28E-07 |
| Nsp8 | RPS6KA4 | -0.624 | 1.30E-07 |

| | | | |
|---|---|---|---|
| Nsp8 | FZD2 | -0.624 | 1.26E-07 |
| Nsp12 | MAP2K2 | -0.624 | 1.31E-07 |
| Nsp13 | HSPB1 | -0.624 | 1.26E-07 |
| Nsp13 | DUSP7 | -0.624 | 1.26E-07 |
| Nsp14 | SHC2 | -0.624 | 1.32E-07 |
| Nsp15 | MAPK8IP2 | -0.624 | 1.27E-07 |
| Nsp16 | EPOR | -0.624 | 1.27E-07 |
| Nsp16 | WNT3 | -0.624 | 1.32E-07 |
| Nsp16 | CD70 | -0.624 | 1.32E-07 |
| Nsp16 | STK11 | -0.624 | 1.26E-07 |
| S | TBKBP1 | -0.624 | 1.31E-07 |
| S | CACNA1I | -0.624 | 1.31E-07 |
| Nsp3 | DUSP5 | -0.625 | 1.25E-07 |
| Nsp4 | FZD1 | -0.625 | 1.23E-07 |
| Nsp5 | GREM2 | -0.625 | 1.24E-07 |
| Nsp12 | FGFR3 | -0.625 | 1.20E-07 |
| Nsp13 | LEFTY1 | -0.625 | 1.21E-07 |
| Nsp15 | FZD8 | -0.625 | 1.19E-07 |
| Nsp16 | DVL3 | -0.625 | 1.25E-07 |
| Nsp16 | DUSP7 | -0.625 | 1.23E-07 |
| S | GDF10 | -0.625 | 1.21E-07 |
| Nsp3 | HSPB1 | -0.626 | 1.11E-07 |
| Nsp3 | BMP6 | -0.626 | 1.17E-07 |
| Nsp3 | WNT3 | -0.626 | 1.14E-07 |
| Nsp5 | LPAR5 | -0.626 | 1.13E-07 |
| Nsp8 | SOCS1 | -0.626 | 1.16E-07 |
| Nsp8 | PGAM5 | -0.626 | 1.13E-07 |
| Nsp8 | INHBB | -0.626 | 1.14E-07 |
| Nsp12 | SMAD6 | -0.626 | 1.14E-07 |
| Nsp13 | SOCS1 | -0.626 | 1.15E-07 |
| Nsp16 | MAPK8IP3 | -0.626 | 1.13E-07 |
| Nsp3 | PRKAR1B | -0.627 | 1.05E-07 |
| Nsp3 | RPS6KA4 | -0.627 | 1.08E-07 |
| Nsp3 | EFNA3 | -0.627 | 1.10E-07 |
| Nsp3 | COMP | -0.627 | 1.10E-07 |
| Nsp4 | MAPK8IP2 | -0.627 | 1.10E-07 |
| Nsp8 | SHC2 | -0.627 | 1.08E-07 |
| Nsp8 | STK11 | -0.627 | 1.06E-07 |
| Nsp12 | GDF10 | -0.627 | 1.06E-07 |
| Nsp12 | GDF6 | -0.627 | 1.10E-07 |
| Nsp12 | DUSP8 | -0.627 | 1.08E-07 |
| Nsp13 | RPS6KA4 | -0.627 | 1.05E-07 |
| Nsp13 | CACNA1I | -0.627 | 1.05E-07 |
| Nsp13 | MAP2K7 | -0.627 | 1.05E-07 |
| Nsp15 | GDF10 | -0.627 | 1.05E-07 |
| Nsp15 | NFATC1 | -0.627 | 1.09E-07 |
| Nsp16 | GRK2 | -0.627 | 1.07E-07 |
| S | JUNB | -0.627 | 1.05E-07 |
| S | CTF1 | -0.627 | 1.10E-07 |
| Nsp3 | ZAP70 | -0.628 | 1.01E-07 |

| | | | |
|---|---|---|---|
| Nsp12 | SLC7A5 | -0.628 | 1.04E-07 |
| Nsp12 | FZD1 | -0.628 | 9.88E-08 |
| Nsp13 | BMP8A | -0.628 | 1.02E-07 |
| Nsp3 | WNT1 | -0.629 | 9.54E-08 |
| Nsp4 | RXRA | -0.629 | 9.71E-08 |
| Nsp4 | IRS2 | -0.629 | 9.75E-08 |
| Nsp5 | MAPK8IP2 | -0.629 | 9.79E-08 |
| Nsp8 | CACNA1I | -0.629 | 9.40E-08 |
| Nsp12 | TGFB1 | -0.629 | 9.51E-08 |
| Nsp15 | DDIT4 | -0.629 | 9.43E-08 |
| Nsp16 | HSPA2 | -0.629 | 9.57E-08 |
| Nsp16 | RGMA | -0.629 | 9.81E-08 |
| Nsp16 | GDF6 | -0.629 | 9.51E-08 |
| Nsp3 | GREM2 | -0.63 | 9.14E-08 |
| Nsp4 | DUSP8 | -0.63 | 8.94E-08 |
| Nsp5 | SMAD6 | -0.63 | 9.24E-08 |
| Nsp8 | DUSP7 | -0.63 | 8.85E-08 |
| Nsp8 | RXRA | -0.63 | 8.79E-08 |
| Nsp9 | GDF15 | -0.63 | 8.78E-08 |
| Nsp15 | DUSP8 | -0.63 | 9.19E-08 |
| Nsp15 | EFNA2 | -0.63 | 8.76E-08 |
| Nsp15 | CACNG8 | -0.63 | 8.74E-08 |
| Nsp16 | SMAD6 | -0.63 | 9.09E-08 |
| Nsp16 | CACNG8 | -0.63 | 9.24E-08 |
| S | HSPA2 | -0.63 | 9.01E-08 |
| Nsp3 | CARD14 | -0.631 | 8.25E-08 |
| Nsp3 | GDF1 | -0.631 | 8.54E-08 |
| Nsp3 | NFATC1 | -0.631 | 8.29E-08 |
| Nsp8 | GDF1 | -0.631 | 8.27E-08 |
| Nsp13 | JUNB | -0.631 | 8.70E-08 |
| Nsp13 | LRP5 | -0.631 | 8.67E-08 |
| Nsp16 | FZD9 | -0.631 | 8.27E-08 |
| Nsp3 | HSPA2 | -0.632 | 7.98E-08 |
| Nsp4 | CEBPB | -0.632 | 7.89E-08 |
| Nsp4 | WNT6 | -0.632 | 7.87E-08 |
| Nsp8 | FGF22 | -0.632 | 7.95E-08 |
| Nsp13 | WNT6 | -0.632 | 7.90E-08 |
| Nsp13 | TGFB1 | -0.632 | 7.81E-08 |
| Nsp14 | GDF7 | -0.632 | 7.81E-08 |
| Nsp16 | FZD5 | -0.632 | 7.91E-08 |
| Nsp16 | ZAP70 | -0.632 | 7.76E-08 |
| S | ZAP70 | -0.632 | 7.71E-08 |
| Nsp4 | MAP2K7 | -0.633 | 7.68E-08 |
| Nsp8 | PPP2R3B | -0.633 | 7.44E-08 |
| Nsp8 | NOG | -0.633 | 7.51E-08 |
| Nsp12 | PRKAR1B | -0.633 | 7.55E-08 |
| Nsp12 | MAPK8IP2 | -0.633 | 7.55E-08 |
| Nsp13 | CARD14 | -0.633 | 7.54E-08 |
| Nsp13 | ADCY5 | -0.633 | 7.67E-08 |
| Nsp16 | LRP5 | -0.633 | 7.70E-08 |

| | | | |
|---|---|---|---|
| Nsp16 | JUN | -0.633 | 7.66E-08 |
| Nsp16 | TGFB1 | -0.633 | 7.49E-08 |
| Nsp16 | RASGRP2 | -0.633 | 7.63E-08 |
| S | GREM2 | -0.633 | 7.52E-08 |
| S | JUND | -0.633 | 7.39E-08 |
| Nsp3 | AMH | -0.634 | 7.04E-08 |
| Nsp12 | CACNG6 | -0.634 | 6.83E-08 |
| Nsp12 | CEBPB | -0.634 | 7.13E-08 |
| Nsp13 | GATA3 | -0.634 | 7.00E-08 |
| Nsp14 | FGF4 | -0.634 | 6.99E-08 |
| Nsp16 | GDF11 | -0.634 | 7.01E-08 |
| S | GRK2 | -0.634 | 7.02E-08 |
| S | FZD2 | -0.634 | 6.95E-08 |
| Nsp3 | BMP8A | -0.635 | 6.68E-08 |
| Nsp4 | BMP6 | -0.635 | 6.73E-08 |
| Nsp8 | WNT7B | -0.635 | 6.67E-08 |
| Nsp13 | MAP2K2 | -0.635 | 6.61E-08 |
| S | TRADD | -0.635 | 6.81E-08 |
| S | PPP2R3B | -0.635 | 6.56E-08 |
| Nsp3 | DVL1 | -0.636 | 6.24E-08 |
| Nsp3 | TGFB1 | -0.636 | 6.06E-08 |
| Nsp5 | CACNG8 | -0.636 | 6.12E-08 |
| Nsp12 | GREM2 | -0.636 | 6.35E-08 |
| Nsp13 | NCF1 | -0.636 | 6.08E-08 |
| Nsp13 | BCL2 | -0.636 | 6.04E-08 |
| Nsp14 | GDF10 | -0.636 | 6.28E-08 |
| Nsp15 | GDF7 | -0.636 | 6.39E-08 |
| Nsp16 | WNT7B | -0.636 | 6.26E-08 |
| Nsp16 | GDF7 | -0.636 | 6.03E-08 |
| S | LMNB2 | -0.636 | 6.10E-08 |
| Nsp12 | GATA3 | -0.637 | 5.72E-08 |
| Nsp13 | GREM2 | -0.637 | 5.93E-08 |
| S | LRP5 | -0.637 | 5.86E-08 |
| Nsp3 | DDIT4 | -0.638 | 5.32E-08 |
| Nsp5 | FZD9 | -0.638 | 5.38E-08 |
| Nsp15 | FZD10 | -0.638 | 5.52E-08 |
| S | DVL1 | -0.638 | 5.47E-08 |
| Nsp12 | RXRA | -0.639 | 5.14E-08 |
| Nsp13 | FZD8 | -0.639 | 5.18E-08 |
| Nsp13 | NOG | -0.639 | 5.24E-08 |
| Nsp13 | STK11 | -0.639 | 5.02E-08 |
| Nsp16 | DDIT4 | -0.639 | 5.04E-08 |
| Nsp16 | MAPK8IP2 | -0.639 | 5.16E-08 |
| S | TGFB1 | -0.639 | 5.06E-08 |
| Nsp3 | PIM1 | -0.64 | 4.73E-08 |
| Nsp12 | IRS2 | -0.64 | 4.77E-08 |
| Nsp15 | FGF4 | -0.64 | 4.91E-08 |
| Nsp16 | SHC2 | -0.64 | 4.88E-08 |
| Nsp16 | AMH | -0.64 | 4.83E-08 |
| Nsp16 | CACNA1I | -0.64 | 4.82E-08 |

| | | | |
|---|---|---|---|
| S | GDF7 | -0.64 | 4.74E-08 |
| Nsp3 | FZD8 | -0.641 | 4.53E-08 |
| Nsp3 | LRP5 | -0.641 | 4.63E-08 |
| Nsp5 | CEBPB | -0.641 | 4.55E-08 |
| Nsp5 | FGF4 | -0.641 | 4.46E-08 |
| Nsp8 | GDF7 | -0.641 | 4.56E-08 |
| Nsp13 | BMP7 | -0.641 | 4.48E-08 |
| Nsp13 | PIM1 | -0.641 | 4.60E-08 |
| Nsp15 | RXRA | -0.641 | 4.47E-08 |
| Nsp15 | PPP2R3B | -0.641 | 4.59E-08 |
| Nsp16 | LPAR5 | -0.641 | 4.47E-08 |
| Nsp3 | SOCS1 | -0.642 | 4.14E-08 |
| Nsp8 | EFNA3 | -0.642 | 4.15E-08 |
| Nsp12 | DDIT4 | -0.642 | 4.30E-08 |
| Nsp12 | NFATC1 | -0.642 | 4.35E-08 |
| Nsp12 | NOG | -0.642 | 4.27E-08 |
| Nsp16 | GREM2 | -0.642 | 4.19E-08 |
| Nsp3 | CACNA1I | -0.643 | 4.09E-08 |
| Nsp8 | PPP1R3D | -0.643 | 4.12E-08 |
| Nsp8 | FZD9 | -0.643 | 4.00E-08 |
| Nsp13 | DDIT4 | -0.643 | 3.98E-08 |
| Nsp13 | RXRA | -0.643 | 4.09E-08 |
| S | NCF1 | -0.643 | 3.97E-08 |
| Nsp12 | PGAM5 | -0.644 | 3.68E-08 |
| Nsp13 | GDF7 | -0.644 | 3.63E-08 |
| Nsp13 | MYC | -0.644 | 3.71E-08 |
| Nsp16 | GDF15 | -0.644 | 3.80E-08 |
| Nsp16 | DUSP8 | -0.644 | 3.73E-08 |
| S | ENDOG | -0.644 | 3.69E-08 |
| S | RPS6KA4 | -0.644 | 3.69E-08 |
| Nsp3 | BMP7 | -0.645 | 3.41E-08 |
| Nsp3 | FZD5 | -0.645 | 3.58E-08 |
| Nsp4 | LPAR5 | -0.645 | 3.45E-08 |
| Nsp8 | FZD8 | -0.645 | 3.41E-08 |
| Nsp12 | SMAD7 | -0.645 | 3.47E-08 |
| Nsp12 | FZD10 | -0.645 | 3.59E-08 |
| Nsp12 | LPAR5 | -0.645 | 3.44E-08 |
| Nsp13 | AMH | -0.645 | 3.58E-08 |
| Nsp15 | FZD1 | -0.645 | 3.53E-08 |
| S | DUSP7 | -0.645 | 3.42E-08 |
| S | PPP1R3D | -0.645 | 3.56E-08 |
| Nsp3 | SMAD6 | -0.646 | 3.20E-08 |
| Nsp4 | PPP2R3B | -0.646 | 3.31E-08 |
| Nsp8 | AMH | -0.646 | 3.33E-08 |
| Nsp8 | TGFB1 | -0.646 | 3.27E-08 |
| Nsp15 | TNFRSF18 | -0.646 | 3.25E-08 |
| Nsp16 | EFNA2 | -0.646 | 3.22E-08 |
| Nsp13 | SMAD7 | -0.647 | 3.15E-08 |
| Nsp13 | EFNA3 | -0.647 | 3.13E-08 |
| Nsp15 | CEBPB | -0.647 | 3.17E-08 |

| | | | |
|---|---|---|---|
| S | DUSP8 | -0.647 | 3.08E-08 |
| S | WNT1 | -0.647 | 3.01E-08 |
| Nsp3 | DUSP7 | -0.648 | 2.88E-08 |
| Nsp3 | NCF1 | -0.648 | 2.85E-08 |
| Nsp3 | MAP2K7 | -0.648 | 2.94E-08 |
| Nsp4 | CACNG8 | -0.648 | 2.91E-08 |
| Nsp8 | HSPB1 | -0.648 | 2.98E-08 |
| Nsp8 | FZD5 | -0.648 | 2.82E-08 |
| Nsp12 | CACNG8 | -0.649 | 2.62E-08 |
| Nsp13 | FZD1 | -0.649 | 2.66E-08 |
| Nsp13 | FZD9 | -0.649 | 2.75E-08 |
| Nsp4 | FGF4 | -0.65 | 2.46E-08 |
| Nsp8 | GREM2 | -0.65 | 2.55E-08 |
| Nsp15 | IRS2 | -0.65 | 2.54E-08 |
| Nsp8 | DDIT4 | -0.651 | 2.32E-08 |
| Nsp15 | JUN | -0.651 | 2.33E-08 |
| Nsp16 | DVL1 | -0.651 | 2.40E-08 |
| Nsp16 | RXRA | -0.651 | 2.36E-08 |
| Nsp16 | TNFRSF6B | -0.651 | 2.32E-08 |
| S | BMP7 | -0.651 | 2.31E-08 |
| Nsp3 | NOG | -0.652 | 2.26E-08 |
| Nsp5 | SHC2 | -0.652 | 2.28E-08 |
| Nsp8 | WNT6 | -0.652 | 2.22E-08 |
| Nsp13 | BMP6 | -0.652 | 2.28E-08 |
| Nsp13 | DUSP8 | -0.652 | 2.20E-08 |
| Nsp16 | CEBPB | -0.652 | 2.29E-08 |
| S | EFNA3 | -0.652 | 2.18E-08 |
| S | GDF1 | -0.652 | 2.18E-08 |
| Nsp3 | GDF10 | -0.653 | 2.03E-08 |
| Nsp3 | FZD1 | -0.653 | 2.03E-08 |
| Nsp4 | FZD10 | -0.653 | 2.05E-08 |
| Nsp8 | IRS2 | -0.653 | 2.02E-08 |
| Nsp13 | SMAD6 | -0.653 | 2.11E-08 |
| Nsp16 | CAPN1 | -0.653 | 2.13E-08 |
| Nsp3 | GDF7 | -0.654 | 1.99E-08 |
| Nsp4 | TNFRSF18 | -0.654 | 1.91E-08 |
| Nsp13 | EFNA2 | -0.654 | 1.92E-08 |
| Nsp16 | WNT6 | -0.654 | 2.00E-08 |
| Nsp16 | FGF4 | -0.654 | 1.89E-08 |
| Nsp3 | TNFRSF18 | -0.655 | 1.76E-08 |
| Nsp3 | DUSP8 | -0.655 | 1.84E-08 |
| Nsp8 | SMAD6 | -0.655 | 1.77E-08 |
| Nsp13 | NFATC1 | -0.655 | 1.86E-08 |
| Nsp5 | JUN | -0.656 | 1.74E-08 |
| Nsp13 | PRKAR1B | -0.656 | 1.71E-08 |
| Nsp13 | GDF10 | -0.656 | 1.71E-08 |
| S | SOCS1 | -0.656 | 1.76E-08 |
| S | FZD8 | -0.656 | 1.65E-08 |
| S | JUN | -0.656 | 1.70E-08 |
| Nsp12 | PPP2R3B | -0.657 | 1.58E-08 |

| | | | |
|---|---|---|---|
| Nsp15 | SHC2 | -0.657 | 1.61E-08 |
| Nsp3 | LPAR5 | -0.658 | 1.50E-08 |
| Nsp3 | MAPK8IP2 | -0.658 | 1.48E-08 |
| Nsp13 | FGF4 | -0.658 | 1.48E-08 |
| S | AMH | -0.658 | 1.44E-08 |
| Nsp3 | RXRA | -0.659 | 1.35E-08 |
| Nsp5 | FZD10 | -0.659 | 1.40E-08 |
| Nsp8 | LPAR5 | -0.659 | 1.44E-08 |
| Nsp12 | JUN | -0.659 | 1.41E-08 |
| Nsp13 | TBKBP1 | -0.659 | 1.42E-08 |
| Nsp2 | GDF15 | -0.66 | 1.26E-08 |
| Nsp4 | SHC2 | -0.66 | 1.26E-08 |
| Nsp12 | SHC2 | -0.66 | 1.30E-08 |
| S | FZD5 | -0.66 | 1.28E-08 |
| Nsp3 | PPP2R3B | -0.661 | 1.21E-08 |
| Nsp8 | DUSP8 | -0.662 | 1.14E-08 |
| Nsp4 | GDF10 | -0.663 | 1.02E-08 |
| Nsp8 | MAPK8IP2 | -0.663 | 1.04E-08 |
| Nsp12 | FGF4 | -0.663 | 1.09E-08 |
| Nsp16 | FZD1 | -0.663 | 1.03E-08 |
| Nsp13 | PGAM5 | -0.664 | 9.72E-09 |
| Nsp13 | MAPK8IP2 | -0.664 | 9.79E-09 |
| S | SMAD6 | -0.664 | 9.72E-09 |
| Nsp3 | WNT6 | -0.665 | 9.15E-09 |
| Nsp3 | FZD9 | -0.665 | 9.50E-09 |
| Nsp8 | FZD1 | -0.665 | 8.94E-09 |
| Nsp13 | FZD10 | -0.665 | 9.53E-09 |
| Nsp15 | GDF15 | -0.665 | 9.45E-09 |
| S | SHC2 | -0.665 | 9.03E-09 |
| Nsp3 | EFNA2 | -0.666 | 8.80E-09 |
| Nsp5 | TNFRSF18 | -0.666 | 8.66E-09 |
| Nsp16 | IRS2 | -0.666 | 8.74E-09 |
| S | FZD1 | -0.667 | 8.06E-09 |
| Nsp8 | FGF4 | -0.668 | 7.52E-09 |
| Nsp13 | CEBPB | -0.669 | 6.81E-09 |
| Nsp16 | FZD10 | -0.669 | 6.87E-09 |
| Nsp3 | SHC2 | -0.67 | 6.60E-09 |
| Nsp3 | IRS2 | -0.67 | 6.63E-09 |
| Nsp8 | GDF15 | -0.67 | 6.50E-09 |
| Nsp14 | GDF15 | -0.67 | 6.56E-09 |
| Nsp4 | JUN | -0.671 | 5.98E-09 |
| S | FZD10 | -0.671 | 5.99E-09 |
| S | MAPK8IP2 | -0.671 | 5.89E-09 |
| S | WNT6 | -0.672 | 5.81E-09 |
| S | NOG | -0.672 | 5.60E-09 |
| Nsp8 | CACNG8 | -0.673 | 5.25E-09 |
| S | TNFRSF18 | -0.673 | 5.09E-09 |
| Nsp3 | FGF4 | -0.675 | 4.51E-09 |
| Nsp13 | PPP2R3B | -0.675 | 4.45E-09 |
| S | FZD9 | -0.675 | 4.56E-09 |

| | | | |
|---|---|---|---|
| Nsp5 | GDF15 | -0.676 | 4.16E-09 |
| Nsp3 | FZD10 | -0.677 | 3.85E-09 |
| Nsp8 | EFNA2 | -0.677 | 3.88E-09 |
| Nsp12 | TNFRSF18 | -0.677 | 4.07E-09 |
| Nsp13 | JUN | -0.677 | 4.03E-09 |
| Nsp13 | GDF15 | -0.678 | 3.57E-09 |
| Nsp3 | JUN | -0.679 | 3.29E-09 |
| Nsp13 | LPAR5 | -0.679 | 3.49E-09 |
| Nsp13 | IRS2 | -0.679 | 3.30E-09 |
| S | LPAR5 | -0.679 | 3.50E-09 |
| S | CACNG8 | -0.679 | 3.34E-09 |
| S | DDIT4 | -0.68 | 3.13E-09 |
| S | IRS2 | -0.68 | 3.24E-09 |
| Nsp8 | FZD10 | -0.682 | 2.74E-09 |
| Nsp8 | CEBPB | -0.682 | 2.76E-09 |
| Nsp12 | BMP6 | -0.685 | 2.12E-09 |
| Nsp3 | CEBPB | -0.686 | 2.02E-09 |
| Nsp13 | SHC2 | -0.686 | 2.02E-09 |
| Nsp13 | CACNG8 | -0.686 | 1.98E-09 |
| Nsp3 | CACNG8 | -0.687 | 1.88E-09 |
| S | FGF4 | -0.69 | 1.54E-09 |
| Nsp13 | TNFRSF18 | -0.692 | 1.32E-09 |
| S | EFNA2 | -0.695 | 1.03E-09 |
| Nsp3 | GDF15 | -0.698 | 8.19E-10 |
| Nsp4 | GDF15 | -0.7 | 6.71E-10 |
| S | CEBPB | -0.703 | 5.45E-10 |
| S | GDF15 | -0.71 | 2.93E-10 |
| Nsp12 | GDF15 | -0.734 | 3.87E-11 |
