## Supplementary-C for "Impact Analysis of SARS-CoV2 on Signaling Pathways during COVID19 Pathogenesis using Codon Usage Assisted Host-Viral Protein Interactions"

**Table S3: The list of top score 100 central host protein. For each protein, centrality score, correlation, interacting viral protein, and involved pathways are shown in different columns.**

| Host protein | Centrality score | Correlation | Viral protein count | Viral protein list | Pathway-count | Pathway list |
|---|---|---|---|---|---|---|
| MYC | 2843 | -0.545 | 13 | Nsp2,Nsp3,Nsp4,Nsp5,Nsp8,Nsp10,Nsp12,Nsp13,Nsp14,Nsp15,Nsp16,S,Orf8 | 4 | MAPK signaling pathway, PI3K-Akt signaling pathway, Jak-STAT signaling pathway, TGF-beta signaling pathway |
| TRIM25 | 2656 | -0.505 | 13 | Nsp2,Nsp3,Nsp4,Nsp5,Nsp8,Nsp10,Nsp12,Nsp13,Nsp14,Nsp15,Nsp16,S,Orf8 | 2 | NF-kappa B signaling pathway, RIG-I-like receptor signaling pathway |
| EGFR | 2452 | -0.435 | 8 | Nsp3,Nsp4,Nsp5,Nsp12,Nsp13,Nsp15,Nsp16,Orf8 | 4 | MAPK signaling pathway, PI3K-Akt signaling pathway, Jak-STAT signaling pathway, HIF-1 signaling pathway |
| BRCA1 | 2236 | 0.648 | 17 | Nsp2,Nsp3,Nsp4,Nsp5,Nsp6,Nsp8,Nsp9,Nsp12,Nsp13,Nsp14,Nsp15,Nsp16,S,N,Orf3a,Orf7a,Orf8 | 1 | PI3K-Akt signaling pathway |
| MDM2 | 2219 | 0.591 | 20 | Nsp2,Nsp3,Nsp4,Nsp5,Nsp6,Nsp7,Nsp8,Nsp9,Nsp12,Nsp13,Nsp14,Nsp15,Nsp16,S,M,N,Orf3a,Orf7a,Orf7b,Orf8 | 1 | PI3K-Akt signaling pathway |
| NTRK1 | 2030 | -0.484 | 14 | Nsp2,Nsp3,Nsp4,Nsp5,Nsp7,Nsp8,Nsp12,Nsp13,Nsp14,Nsp15,Nsp16,S,Orf3a,Orf8 | 3 | MAPK signaling pathway, PI3K-Akt signaling pathway, Apoptosis |
| KRAS | 1944 | 0.558 | 15 | Nsp3,Nsp4,Nsp6,Nsp8,Nsp9,Nsp12,Nsp13,Nsp14,Nsp15,Nsp16,S,N,Orf3a,Orf7a,Orf8 | 6 | MAPK signaling pathway, Chemokine signaling pathway, PI3K-Akt signaling pathway, Apoptosis, Insulin signaling pathway, mTOR signaling pathway |
| ELAVL1 | 1914 | -0.460 | 13 | Nsp2,Nsp3,Nsp4,Nsp5,Nsp8,Nsp9,Nsp10,Nsp12,Nsp13,Nsp14,Nsp15,Nsp16,S | 1 | IL-17 signaling pathway |
| HSP90AA1 | 1734 | 0.459 | 7 | Nsp2,Nsp3,Nsp4,Nsp6,Nsp8,S,Orf7a | 3 | IL-17 signaling pathway, PI3K-Akt signaling pathway, Th17 cell differentiation |
| CUL1 | 1503 | 0.442 | 2 | N,Orf3a | 1 | TGF-beta signaling pathway |

| | | | | | | |
|---|---|---|---|---|---|---|
| BIRC3 | 1483 | 0.662 | 17 | Nsp2,Nsp3,Nsp4,Nsp5,Nsp6,Nsp8,Nsp9,Nsp12,Nsp13,Nsp14,Nsp15,Nsp16,S,N,Orf3a,Orf7a,Orf8 | 3 | NF-kappa B signaling pathway, TNF signaling pathway, Apoptosis |
| HSPA8 | 1317 | 0.475 | 5 | Nsp1,Nsp2,S,M,N | 1 | MAPK signaling pathway |
| TRAF2 | 1163 | -0.515 | 13 | Nsp2,Nsp3,Nsp4,Nsp5,Nsp8,Nsp10,Nsp12,Nsp13,Nsp14,Nsp15,Nsp16,S,Orf8 | 7 | NF-kappa B signaling pathway, TNF signaling pathway, IL-17 signaling pathway, RIG-I-like receptor signaling pathway, MAPK signaling pathway, Apoptosis, Adipocytokine signaling pathway |
| HIF1A | 1090 | 0.623 | 18 | Nsp2,Nsp3,Nsp4,Nsp5,Nsp6,Nsp8,Nsp9,Nsp12,Nsp13,Nsp14,Nsp15,Nsp16,S,M,N,Orf3a,Orf7a,Orf8 | 2 | Th17 cell differentiationHIF-1 signaling pathway |
| CREBBP | 1060 | -0.437 | 5 | Nsp3,Nsp4,Nsp8,Nsp12,Nsp16 | 3 | Jak-STAT signaling pathway, TGF-beta signaling pathway, HIF-1 signaling pathway |
| LMNA | 1021 | -0.499 | 11 | Nsp3,Nsp4,Nsp5,Nsp8,Nsp12,Nsp13,Nsp14,Nsp15,Nsp16,S,Orf8 | 1 | Apoptosis |
| SKP1 | 996 | 0.470 | 2 | E,Orf3a | 1 | TGF-beta signaling pathway |
| TRAF6 | 988 | 0.427 | 3 | Nsp3,Nsp8,S | 5 | NF-kappa B signaling pathway, IL-17 signaling pathway, RIG-I-like receptor signaling pathway, MAPK signaling pathway, Toll-like receptor signaling pathway |
| IKBKG | 959 | -0.431 | 9 | Nsp3,Nsp8,Nsp12,Nsp13,Nsp14,Nsp15,Nsp16,S,Orf8 | 11 | NF-kappa B signaling pathway, TNF signaling pathway, IL-17 signaling pathway, RIG-I-like receptor signaling pathway, MAPK signaling pathway, Chemokine signaling pathway, PI3K-Akt signaling pathway, Th17 cell differentiation, Toll-like receptor signaling pathway, Apoptosis, Adipocytokine signaling pathway |
| VHL | 958 | -0.484 | 12 | Nsp2,Nsp3,Nsp4,Nsp5,Nsp7,Nsp8,Nsp12,Nsp13,Nsp14,Nsp15,Nsp16,S | 1 | HIF-1 signaling pathway |
| CYLD | 945 | 0.583 | 16 | Nsp2,Nsp3,Nsp4,Nsp5,Nsp6,Nsp8,Nsp9,Nsp12,Nsp13,Nsp14,Nsp15,Nsp16,S,Orf3a,Orf7a,Orf8 | 2 | NF-kappa B signaling pathway, RIG-I-like receptor signaling pathway |
| YWHAZ | 891 | 0.455 | 6 | Nsp2,Nsp5,Nsp13,S,N,Orf7a | 1 | PI3K-Akt signaling pathway |
| AKT1 | 886 | -0.500 | 11 | Nsp3,Nsp4,Nsp5,Nsp8,Nsp12,Nsp13,Nsp14,Nsp15,Nsp16,S,Orf8 | 11 | TNF signaling pathway, MAPK signaling pathway, Chemokine signaling pathway, PI3K-Akt signaling pathway, Jak-STAT signaling pathway, Toll-like receptor signaling pathway, HIF-1 signaling pathway, Apoptosis, Insulin signaling pathway, mTOR signaling pathway, Adipocytokine signaling pathway |
| HRAS | 885 | -0.451 | 10 | Nsp3,Nsp4,Nsp5,Nsp8,Nsp12,Nsp13,Nsp14,Nsp16,S,Orf8 | 7 | MAPK signaling pathway, Chemokine signaling pathway, PI3K-Akt signaling pathway, Jak-STAT signaling pathway, Apoptosis, Insulin signaling pathway, mTOR signaling pathway, |
| RELA | 859 | -0.455 | 10 | Nsp3,Nsp4,Nsp5,Nsp8,Nsp12,Nsp13,Nsp15,Nsp16,S,Orf6 | 12 | NF-kappa B signaling pathway, TNF signaling pathway, IL-17 signaling pathway, RIG-I-like receptor signaling pathway, MAPK signaling pathway, Chemokine signaling pathway, PI3K-Akt signaling pathway, Th17 cell differentiation, Toll-like receptor signaling pathway, HIF-1 signaling pathway, Apoptosis, Adipocytokine signaling pathway |
| PPP2R1A | 800 | -0.506 | 11 | Nsp3,Nsp4,Nsp5,Nsp8,Nsp12,Nsp13,Nsp14,Nsp15,Nsp16,S,Orf8 | 2 | PI3K-Akt signaling pathway, TGF-beta signaling pathway |
| SMAD3 | 799 | -0.493 | 14 | Nsp2,Nsp3,Nsp4,Nsp5,Nsp7,Nsp8,Nsp10,Nsp12,Nsp13,Nsp14,Nsp15,Nsp16,S,Orf8 | 2 | Th17 cell differentiation, TGF-beta signaling pathway |
| CDKN1A | 768 | -0.444 | 4 | Nsp15,Nsp16,Orf6,Orf10 | 3 | PI3K-Akt signaling pathway, Jak-STAT signaling pathway, HIF-1 signaling pathway |

| | | | | | | |
|---|---|---|---|---|---|---|
| CUL2 | 767 | 0.615 | 19 | Nsp2,Nsp3,Nsp4,Nsp5,Nsp6,Nsp7,Nsp8,Nsp9,Nsp12,Nsp13,Nsp14,Nsp15,Nsp16,S,M,N,Orf3a,Orf7a,Orf8 | 1 | HIF-1 signaling pathway |
| SRC | 712 | -0.527 | 14 | Nsp2,Nsp3,Nsp4,Nsp5,Nsp8,Nsp10,Nsp12,Nsp13,Nsp14,Nsp15,Nsp16,S,Orf3a,Orf8 | 1 | Chemokine signaling pathway |
| HSPB1 | 685 | -0.557 | 17 | Nsp2,Nsp3,Nsp4,Nsp5,Nsp6,Nsp7,Nsp8,Nsp9,Nsp10,Nsp12,Nsp13,Nsp14,Nsp15,Nsp16,S,Orf3a,Orf8 | 1 | MAPK signaling pathway |
| PIK3R1 | 684 | 0.451 | 5 | Nsp3,Nsp16,S,N,Orf3a | 9 | TNF signaling pathway, Chemokine signaling pathway, PI3K-Akt signaling pathway, Jak-STAT signaling pathway, Toll-like receptor signaling pathway, HIF-1 signaling pathway, Apoptosis, Insulin signaling pathway, mTOR signaling pathway |
| RPS6 | 680 | 0.459 | 4 | Nsp2,Nsp4,S,N | 7 | TNF signaling pathway, MAPK signaling pathway, PI3K-Akt signaling pathway, TGF-beta signaling pathway, HIF-1 signaling pathway, Insulin signaling pathway, mTOR signaling pathway |
| PPP2CA | 658 | 0.527 | 14 | Nsp2,Nsp3,Nsp4,Nsp5,Nsp6,Nsp8,Nsp12,Nsp13,Nsp14,Nsp15,Nsp16,S,M,Orf7a | 2 | PI3K-Akt signaling pathway, TGF-beta signaling pathway |
| JUN | 655 | -0.584 | 18 | Nsp2,Nsp3,Nsp4,Nsp5,Nsp6,Nsp7,Nsp8,Nsp9,Nsp10,Nsp12,Nsp13,Nsp14,Nsp15,Nsp16,S,Orf3a,Orf6,Orf8 | 6 | TNF signaling pathway, IL-17 signaling pathway, MAPK signaling pathway, Th17 cell differentiation, Toll-like receptor signaling pathway, Apoptosis |
| CDC37 | 636 | -0.459 | 11 | Nsp3,Nsp4,Nsp5,Nsp8,Nsp12,Nsp13,Nsp14,Nsp15,Nsp16,S,Orf8 | 1 | PI3K-Akt signaling pathway |
| ACTB | 601 | -0.464 | 9 | Nsp3,Nsp4,Nsp5,Nsp8,Nsp12,Nsp13,Nsp15,Nsp16,S | 1 | Apoptosis |
| XIAP | 579 | 0.640 | 18 | Nsp2,Nsp3,Nsp4,Nsp5,Nsp6,Nsp7,Nsp8,Nsp9,Nsp12,Nsp13,Nsp14,Nsp15,Nsp16,S,M,Orf3a,Orf7a,Orf8 | 2 | NF-kappa B signaling pathway, Apoptosis |
| YWHAG | 558 | -0.510 | 17 | Nsp2,Nsp3,Nsp4,Nsp5,Nsp6,Nsp7,Nsp8,Nsp9,Nsp10,Nsp12,Nsp13,Nsp14,Nsp15,Nsp16,S,Orf3a,Orf8 | 1 | PI3K-Akt signaling pathway |
| IKBKB | 552 | -0.426 | 2 | Nsp12,Nsp13 | 13 | NF-kappa B signaling pathway, TNF signaling pathway, IL-17 signaling pathway, RIG-I-like receptor signaling pathway, MAPK signaling pathway, Chemokine signaling pathway, PI3K-Akt signaling pathway, Th17 cell differentiation, Toll-like receptor signaling pathway, Apoptosis, Insulin signaling pathway, mTOR signaling pathway, Adipocytokine signaling pathway |
| SMAD4 | 546 | 0.517 | 17 | Nsp2,Nsp3,Nsp4,Nsp5,Nsp6,Nsp8,Nsp9,Nsp10,Nsp12,Nsp13,Nsp14,Nsp15,Nsp16,S,M,Orf3a,Orf8 | 2 | Th17 cell differentiation, TGF-beta signaling pathway |
| ITCH | 538 | 0.643 | 17 | Nsp2,Nsp3,Nsp4,Nsp5,Nsp6,Nsp8,Nsp9,Nsp12,Nsp13,Nsp14,Nsp15,Nsp16,S,N,Orf3a,Orf7a,Orf8 | 1 | TNF signaling pathway |
| SMAD2 | 538 | 0.477 | 13 | Nsp3,Nsp4,Nsp5,Nsp8,Nsp9,Nsp12,Nsp13,Nsp14,Nsp15,Nsp16,S,Orf3a,Orf8 | 2 | Th17 cell differentiation, TGF-beta signaling pathway |
| ITGA4 | 530 | 0.558 | 14 | Nsp3,Nsp4,Nsp5,Nsp6,Nsp8,Nsp12,Nsp13,Nsp14,Nsp15,Nsp16,S,N,Orf3a,Orf7a | 1 | PI3K-Akt signaling pathway |

| | | | | | | |
|---|---|---|---|---|---|---|
| ARRB2 | 512 | -0.448 | 9 | Nsp3,Nsp4,Nsp8,Nsp12,Nsp13,Nsp15,Nsp16,S,Orf8 | 2 | MAPK signaling pathway, Chemokine signaling pathway |
| NFKBIA | 501 | -0.450 | 8 | Nsp3,Nsp4,Nsp5,Nsp8,Nsp13,Nsp15,Nsp16,S | 9 | NF-kappa B signaling pathway, TNF signaling pathway, IL-17 signaling pathway, RIG-I-like receptor signaling pathway, Chemokine signaling pathway, Th17 cell differentiation, Toll-like receptor signaling pathway, Apoptosis, Adipocytokine signaling pathway |
| RPTOR | 501 | -0.532 | 15 | Nsp2,Nsp3,Nsp4,Nsp5,Nsp7,Nsp8,Nsp10,Nsp12,Nsp13,Nsp14,Nsp15,Nsp16,S,Orf6,Orf8 | 3 | PI3K-Akt signaling pathway, Insulin signaling pathway, mTOR signaling pathway |
| TNF | 497 | -0.474 | 11 | Nsp2,Nsp3,Nsp4,Nsp8,Nsp10,Nsp12,Nsp13,Nsp15,Nsp16,S,Orf8 | 11 | NF-kappa B signaling pathway, Cytokine-cytokine receptor interaction, TNF signaling pathway, IL-17 signaling pathway, RIG-I-like receptor signaling pathway, MAPK signaling pathway, TGF-beta signaling pathway, Toll-like receptor signaling pathway, Apoptosis, mTOR signaling pathway, Adipocytokine signaling pathway |
| TRAF1 | 493 | -0.470 | 11 | Nsp3,Nsp4,Nsp5,Nsp8,Nsp10,Nsp12,Nsp13,Nsp14,Nsp15,Nsp16,Orf8 | 3 | NF-kappa B signaling pathway, TNF signaling pathway, Apoptosis |
| ELOB | 491 | -0.446 | 10 | Nsp2,Nsp3,Nsp4,Nsp5,Nsp7,Nsp8,Nsp12,Nsp13,Nsp16,S | 1 | HIF-1 signaling pathway |
| BIRC2 | 490 | 0.653 | 19 | Nsp2,Nsp3,Nsp4,Nsp5,Nsp6,Nsp7,Nsp8,Nsp9,Nsp12,Nsp13,Nsp14,Nsp15,Nsp16,S,M,N,Orf3a,Orf7a,Orf8 | 3 | NF-kappa B signaling pathway, TNF signaling pathway, Apoptosis |
| MAPT | 489 | -0.423 | 1 | Nsp13 | 1 | MAPK signaling pathway |
| ERBB2 | 488 | -0.456 | 11 | Nsp3,Nsp4,Nsp5,Nsp8,Nsp12,Nsp13,Nsp14,Nsp15,Nsp16,S,Orf8 | 3 | MAPK signaling pathway, PI3K-Akt signaling pathway, HIF-1 signaling pathway |
| MAP3K7 | 483 | 0.558 | 17 | Nsp2,Nsp3,Nsp4,Nsp5,Nsp6,Nsp8,Nsp9,Nsp12,Nsp13,Nsp14,Nsp15,Nsp16,S,M,N,Orf3a,Orf8 | 6 | NF-kappa B signaling pathway, TNF signaling pathway, IL-17 signaling pathway, RIG-I-like receptor signaling pathway, MAPK signaling pathway, Toll-like receptor signaling pathway |
| MAPK6 | 483 | 0.589 | 19 | Nsp2,Nsp3,Nsp4,Nsp5,Nsp6,Nsp8,Nsp9,Nsp12,Nsp13,Nsp14,Nsp15,Nsp16,S,M,N,Orf3a,Orf6,Orf7a,Orf8 | 1 | IL-17 signaling pathway |
| ELOC | 472 | 0.440 | 1 | Nsp13 | 1 | HIF-1 signaling pathway |
| PIN1 | 471 | -0.501 | 13 | Nsp2,Nsp3,Nsp4,Nsp5,Nsp8,Nsp10,Nsp12,Nsp13,Nsp14,Nsp15,Nsp16,S,Orf8 | 2 | RIG-I-like receptor signaling pathway, mTOR signaling pathway |
| PTPN11 | 467 | 0.429 | 4 | Nsp6,Nsp14,Nsp16,Orf7a | 2 | Jak-STAT signaling pathway, Adipocytokine signaling pathway |
| VCAM1 | 463 | 0.499 | 13 | Nsp3,Nsp4,Nsp5,Nsp6,Nsp8,Nsp9,Nsp12,Nsp13,Nsp14,Nsp15,Nsp16,S,Orf7a | 2 | NF-kappa B signaling pathway, TNF signaling pathway |
| CHUK | 462 | 0.489 | 11 | Nsp2,Nsp3,Nsp4,Nsp6,Nsp8,Nsp12,Nsp13,Nsp14,Nsp15,Nsp16,S | 12 | NF-kappa B signaling pathway, TNF signaling pathway, IL-17 signaling pathway, RIG-I-like receptor signaling pathway, MAPK signaling pathway, Chemokine signaling pathway, PI3K-Akt signaling pathway, Th17 cell differentiation, Toll-like receptor signaling pathway, Apoptosis, mTOR signaling pathway, Adipocytokine signaling pathway |
| MAPK8 | 444 | 0.472 | 12 | Nsp3,Nsp4,Nsp5,Nsp6,Nsp8,Nsp12,Nsp14,Nsp15,Nsp16,S,Orf3a,Orf7a | 9 | TNF signaling pathway, IL-17 signaling pathway, RIG-I-like receptor signaling pathway, MAPK signaling pathway, Th17 cell differentiation, Toll-like receptor signaling pathway, Apoptosis, Insulin signaling pathway, Adipocytokine signaling pathway |
| CRK | 438 | -0.462 | 8 | Nsp2,Nsp4,Nsp5,Nsp12,Nsp13,Nsp15,Nsp16,S | 3 | MAPK signaling pathway, Chemokine signaling pathway, Insulin signaling pathway, |

| | | | | | | |
|---|---|---|---|---|---|---|
| CCND1 | 435 | -0.547 | 14 | Nsp2,Nsp3,Nsp4,Nsp5,Nsp6,Nsp8,Nsp12,Nsp13,Nsp14,Nsp15,Nsp16,S,Orf3a,Orf8 | 2 | PI3K-Akt signaling pathway, Jak-STAT signaling pathway |
| ATM | 426 | 0.682 | 18 | Nsp2,Nsp3,Nsp4,Nsp5,Nsp6,Nsp8,Nsp9,Nsp12,Nsp13,Nsp14,Nsp15,Nsp16,S,M,N,Orf3a,Orf7a,Orf8 | 2 | NF-kappa B signaling pathway, Apoptosis |
| HSPA1A | 421 | -0.529 | 15 | Nsp2,Nsp3,Nsp4,Nsp5,Nsp8,Nsp10,Nsp12,Nsp13,Nsp14,Nsp15,Nsp16,S,Orf3a,Orf8,Orf10 | 1 | MAPK signaling pathway |
| CDC42 | 419 | 0.550 | 15 | Nsp2,Nsp3,Nsp4,Nsp5,Nsp6,Nsp7,Nsp8,Nsp12,Nsp13,Nsp14,Nsp15,Nsp16,S,Orf3a,Orf7a | 2 | MAPK signaling pathway, Chemokine signaling pathway |
| TBK1 | 416 | 0.663 | 18 | Nsp2,Nsp3,Nsp4,Nsp5,Nsp6,Nsp8,Nsp9,Nsp10,Nsp12,Nsp13,Nsp14,Nsp15,Nsp16,S,M,Orf3a,Orf7a,Orf8 | 3 | IL-17 signaling pathway, RIG-I-like receptor signaling pathway, Toll-like receptor signaling pathway |
| AIFM1 | 407 | 0.469 | 3 | Nsp6,N,Orf7a | 1 | Apoptosis |
| BCL2L1 | 396 | -0.421 | 1 | Nsp5 | 4 | NF-kappa B signaling pathway, PI3K-Akt signaling pathway, Jak-STAT signaling pathway, Apoptosis |
| CDKN1B | 391 | -0.435 | 1 | Nsp10 | 2 | PI3K-Akt signaling pathway, HIF-1 signaling pathway |
| TNFRSF1A | 384 | -0.456 | 12 | Nsp3,Nsp4,Nsp5,Nsp8,Nsp10,Nsp12,Nsp13,Nsp14,Nsp15,Nsp16,S,Orf8 | 7 | NF-kappa B signaling pathway, Cytokine-cytokine receptor interaction, TNF signaling pathway, MAPK signaling pathway, Apoptosis, mTOR signaling pathway, Adipocytokine signaling pathway |
| YWHAH | 383 | -0.424 | 1 | Nsp13 | 1 | PI3K-Akt signaling pathway |
| MAX | 380 | -0.444 | 4 | Nsp3,Nsp4,Nsp15,Nsp16 | 1 | MAPK signaling pathway |
| ARAF | 372 | -0.504 | 12 | Nsp3,Nsp4,Nsp5,Nsp8,Nsp10,Nsp12,Nsp13,Nsp14,Nsp15,Nsp16,S,Orf8 | 2 | MAPK signaling pathway, Insulin signaling pathway |
| TRAF3 | 370 | -0.430 | 3 | Nsp7,Nsp12,S | 5 | NF-kappa B signaling pathway, TNF signaling pathway, IL-17 signaling pathway, RIG-I-like receptor signaling pathway, Toll-like receptor signaling pathway |
| ARRB1 | 367 | -0.430 | 4 | Nsp3,Nsp12,Nsp13,Nsp16 | 2 | MAPK signaling pathway, Chemokine signaling pathway |
| DDX3X | 362 | 0.497 | 14 | Nsp1,Nsp2,Nsp3,Nsp4,Nsp8,Nsp12,Nsp13,Nsp14,Nsp15,S,M,N,Orf3a,Orf8 | 1 | RIG-I-like receptor signaling pathway |
| RARA | 356 | -0.539 | 16 | Nsp2,Nsp3,Nsp4,Nsp5,Nsp8,Nsp9,Nsp10,Nsp12,Nsp13,Nsp14,Nsp15,Nsp16,S,Orf3a,Orf6,Orf8 | 1 | Th17 cell differentiation |
| FGFR1 | 353 | -0.450 | 8 | Nsp3,Nsp4,Nsp5,Nsp12,Nsp13,Nsp15,Nsp16,S | 2 | MAPK signaling pathway, PI3K-Akt signaling pathway |
| RXRA | 353 | -0.569 | 17 | Nsp2,Nsp3,Nsp4,Nsp5,Nsp6,Nsp8,Nsp9,Nsp10,Nsp12,Nsp13,Nsp14,Nsp15,Nsp16,S,Orf3a,Orf6,Orf8 | 3 | PI3K-Akt signaling pathway, Th17 cell differentiation, Adipocytokine signaling pathway |
| SMURF2 | 352 | 0.544 | 15 | Nsp2,Nsp3,Nsp4,Nsp5,Nsp6,Nsp8,Nsp12,Nsp13,Nsp14,Nsp15,Nsp16,S,Orf3a,Orf7a,Orf8 | 1 | TGF-beta signaling pathway |
| ATF2 | 347 | 0.635 | 18 | Nsp2,Nsp3,Nsp4,Nsp5,Nsp6,Nsp8,Nsp9,Nsp12,Nsp13,Nsp14,Nsp15,Nsp16,S,M,N,Orf3a,Orf7a,Orf8 | 3 | TNF signaling pathway, MAPK signaling pathway, PI3K-Akt signaling pathway, |
| PPP2CB | 342 | 0.515 | 11 | Nsp2,Nsp3,Nsp4,Nsp5,Nsp12,Nsp13,Nsp15,S,M,Orf7a,Orf8 | 2 | PI3K-Akt signaling pathway, TGF-beta signaling pathway, |
| GAPDH | 338 | -0.427 | 1 | Nsp16 | 1 | HIF-1 signaling pathway, |

| | | | | | | |
|---|---|---|---|---|---|---|
| MAPK3 | 337 | -0.521 | 16 | Nsp2,Nsp3,Nsp4,Nsp5,Nsp7,Nsp8,Nsp9,Nsp10,Nsp12,Nsp13,Nsp14,Nsp15,Nsp16,S,Orf3a,Orf8 | 12 | TNF signaling pathway, IL-17 signaling pathway, MAPK signaling pathway, Chemokine signaling pathway, PI3K-Akt signaling pathway, Th17 cell differentiation, TGF-beta signaling pathway, Toll-like receptor signaling pathway, HIF-1 signaling pathway, Apoptosis, Insulin signaling pathway, mTOR signaling pathway |
| RUNX1 | 334 | -0.508 | 12 | Nsp2,Nsp3,Nsp4,Nsp5,Nsp8,Nsp12,Nsp13,Nsp14,Nsp15,Nsp16,S,Orf8 | 1 | Th17 cell differentiation |
| HSP90B1 | 331 | 0.577 | 19 | Nsp2,Nsp3,Nsp4,Nsp5,Nsp6,Nsp7,Nsp8,Nsp12,Nsp13,Nsp14,Nsp15,Nsp16,S,E,M,N,Orf3a,Orf7a,Orf8 | 2 | IL-17 signaling pathway, PI3K-Akt signaling pathway |
| PDK1 | 330 | 0.488 | 3 | N,Orf3a,Orf7a | 1 | HIF-1 signaling pathway |
| BCL2 | 329 | -0.548 | 15 | Nsp2,Nsp3,Nsp4,Nsp5,Nsp7,Nsp8,Nsp10,Nsp12,Nsp13,Nsp14,Nsp15,Nsp16,S,Orf3a,Orf8 | 5 | NF-kappa B signaling pathway, PI3K-Akt signaling pathway, Jak-STAT signaling pathway, HIF-1 signaling pathway, Apoptosis |
| TGFBR1 | 327 | 0.563 | 18 | Nsp2,Nsp3,Nsp4,Nsp5,Nsp6,Nsp7,Nsp8,Nsp12,Nsp13,Nsp14,Nsp15,Nsp16,S,N,Orf3a,Orf7a,Orf7b,Orf8 | 4 | Cytokine-cytokine receptor interaction, MAPK signaling pathway, Th17 cell differentiation, TGF-beta signaling pathway |
| PRKCZ | 319 | -0.493 | 11 | Nsp3,Nsp4,Nsp8,Nsp10,Nsp12,Nsp13,Nsp14,Nsp15,Nsp16,S,Orf8 | 2 | Chemokine signaling pathway, Insulin signaling pathway, |
| DVL2 | 317 | -0.436 | 7 | Nsp3,Nsp8,Nsp12,Nsp13,Nsp16,Orf8,Orf10 | 1 | mTOR signaling pathway, |
| IRAK1 | 313 | -0.477 | 11 | Nsp3,Nsp4,Nsp5,Nsp8,Nsp12,Nsp13,Nsp14,Nsp15,Nsp16,S,Orf8 | 3 | NF-kappa B signaling pathway, MAPK signaling pathway, Toll-like receptor signaling pathway, |
| MAVS | 306 | -0.430 | 1 | Nsp16 | 1 | RIG-I-like receptor signaling pathway, |
| RICTOR | 304 | 0.587 | 18 | Nsp2,Nsp3,Nsp4,Nsp5,Nsp6,Nsp8,Nsp9,Nsp10,Nsp12,Nsp13,Nsp14,Nsp15,Nsp16,S,N,Orf3a,Orf7a,Orf8 | 1 | mTOR signaling pathway |
| PTK2 | 300 | 0.425 | 3 | Nsp3,Nsp13,S | 2 | Chemokine signaling pathway, PI3K-Akt signaling pathway |
| FADD | 294 | -0.487 | 13 | Nsp2,Nsp3,Nsp4,Nsp5,Nsp8,Nsp12,Nsp13,Nsp14,Nsp15,Nsp16,S,Orf3a,Orf6 | 5 | TNF signaling pathway, IL-17 signaling pathway, RIG-I-like receptor signaling pathway, Toll-like receptor signaling pathway, Apoptosis |
| PLCG1 | 291 | -0.509 | 12 | Nsp3,Nsp4,Nsp5,Nsp8,Nsp10,Nsp12,Nsp13,Nsp14,Nsp15,Nsp16,S,Orf8 | 3 | NF-kappa B signaling pathway, Th17 cell differentiationHIF-1 signaling pathway |
| TANK | 286 | 0.488 | 9 | Nsp3,Nsp6,Nsp12,Nsp13,Nsp16,S,N,Orf3a,Orf7a | 1 | RIG-I-like receptor signaling pathway |
| TAB2 | 285 | 0.571 | 20 | Nsp2,Nsp3,Nsp4,Nsp5,Nsp6,Nsp7,Nsp8,Nsp9,Nsp12,Nsp13,Nsp14,Nsp15,Nsp16,S,E,N,Orf3a,Orf7a,Orf7b,Orf8 | 5 | NF-kappa B signaling pathway, TNF signaling pathway, IL-17 signaling pathway, MAPK signaling pathway, Toll-like receptor signaling pathway |
